# The Effects of Functional Training on Muscle Strength in Athletes: A Meta-Analysis

**DOI:** 10.1101/2024.06.01.596934

**Authors:** Junyan Liu, Lei Shang, Hongjun Yu

## Abstract

The study aimed to analyze the effects of functional training (FT) on athletes muscle strength compared to traditional resistance training (TRT). A systematic search was conducted in 6 databases from inception to January 2024. Baseline and outcome measures from randomized controlled trials (RCT) were assessed for the impact of FT on athlete muscle strength, across sex, age, and sports levels. Hedge’s g effect sizes were calculated using a random-effects model, and subgroup and single training factor analyses were performed to address potential sources of heterogeneity, along with a meta-regression analysis. The inclusion of 67 studies involving 1718 athletes revealed significant moderate to large effects of FT on maximum strength (k=11; ES=2.68; p≤ 0.001), power (k=12; ES=0.68; p<0.001), as well as muscle endurance(k=12; ES=4.13; p<0.001).In conclusion, FT appears to offer potential benefits for enhancing muscle strength in athletes, with considerations for individual differences in sex, age, and training programs. Findings suggest that longer training sessions may be associated with improvements in maximum strength and muscle endurance, whereas shorter sessions might be more conducive to power development. Additionally, preliminary evidence hints that younger athletes may experience more pronounced training benefits, though this observation requires further investigation to establish a more definitive correlation.

## Introduction

Athletic performance is shaped by a multitude of factors, including genetics, mental and physical health, and the training regimens employed[48,69,88,101].While genetics play a significant role, the potential for enhancement through focused training efforts is equally important[91]. Technical skill is a critical component, yet it is just one of many elements that contribute to optimal performance[33,45]. Strengthening athletic abilities is essential for sports excellence, impacting a range of physical attributes necessary for success[115]. Notably, muscle strength is a pivotal factor that not only elevates performance but also mitigates injury risk[32,58,67,73,79,120]. Elite athletes often exhibit superior muscle strength, a key attribute for their sport-specific proficiency[34].

Muscle strength, defined as the capacity to exert force, is influenced by a combination of physical and neurological factors[104,113], as well as an individual’s fitness level and training type[10,25]. Among the various training methodologies, functional training (FT) aims to enhance overall movement efficiency and power through diverse, multi-directional exercises with an emphasis on injury prevention. FT has demonstrated positive impacts on strength [2,5,23,26,29,31,37,42,47,51,59,70,81,87,96,106,107,121,127,130], power[5,57,59,119,121,122,133], physical abilities[41,131,132], and injury prevention [20,43,93,94,98] making it a staple in comprehensive athletic training approaches.

Gambetta’s research underscores the value of FT in improving coordination across multiple joints and refining proprioception[44]. Boyle advocates for a training approach that emphasizes balance and proprioception, favoring dynamic bodyweight exercises over stationary equipment[24]. Silva-Grigoletto proposes that FT integrates various skills to enhance the effectiveness and safety of performance in specific sports through balanced training[105]. For example, the Bulgarian split squat—a common FT exercise—demands stability and body awareness to engage multiple joints and move in different planes, unlike the seated leg curl, which is a basic traditional resistance training (TRT) exercise targeting a single muscle group without the need for balance[44,105].

Despite the extensive research highlighting the benefits of FT, there remains a lack of comprehensive studies directly comparing the effects of TRT and FT on muscle strength. This study seeks to address this gap by evaluating the impact of these two training modalities on muscle strength, an area that, while not entirely uncharted, has not been exhaustively explored in comparative terms. Additionally, this research will explore how variables such as training frequency, timing, and personal attributes like age and fitness level might influence the effectiveness of FT.

Through a thorough meta-analysis, the objective is to offer practical insights for crafting training programs that effectively enhance muscle strength in athletes. While there are many studies on the effects of both TRT and FT individually, few meta-analyses have directly compared the two. By conducting this analysis, we aim to provide a more nuanced understanding of how these training approaches can be optimized for athletic performance.

## 2. Methods

The study adhered to the PRISMA (Preferred Reporting Items for Systematic Reviews and Meta-Analyses) guidelines for literature search, data extraction, and analysis[76]. It was also pre-registered on the PROSPERO (International Prospective Register of Systematic Reviews) website under registration number CRD42024509103.

### Search Strategy

We systematically searched 6 databases (Web of Science, Google Scholar, Baidu Scholar, EBSCOhost SPORTDiscus with Full Text, Pubmed, Wanfang Data Knowledge Service Platform, and China National Knowledge Infrastructure (CNKI)), from their inception until January 2024. The search terms used in Pubmed included (((functional training[Title] OR functional exercise*[Title] OR core training[Title] OR neuromuscular training[Title] OR plyometric training[Title] OR balance training[Title]) OR swiss ball[Title] OR suspension Training[Title] OR proprioceptive training[Title] OR TRX[Title]) AND (athlete*[Title/Abstract] OR player*[Title/Abstract] OR sport*[Title/Abstract] OR runner*[Title/Abstract] OR rower*[Title/Abstract] OR boxer*[Title/Abstract] OR swimmer*[Title/Abstract] OR jumper*[Title/Abstract] OR shooter*[Title/Abstract] OR throw[Title/Abstract] OR cyclist*[Title/Abstract] OR trained[Title/Abstract] OR archer*[Title/Abstract] OR diver*[Title/Abstract] OR skater*[Title/Abstract] OR gymnast*[Title/Abstract] OR climber*[Title/Abstract] OR athletic[Title/Abstract] OR fence*[Title/Abstract] OR surfer*[Title/Abstract] OR sprinter*[Title/Abstract] OR kayaker*[Title/Abstract] OR triathlete*[Title/Abstract] OR wrestler*[Title/Abstract] OR skater*[Title/Abstract])) AND (effect*[Title/Abstract] OR impact*[Title/Abstract] OR influence[Title/Abstract] OR affect[Title/Abstract])) NOT (patient*[Title] OR old*[Title] OR aged*[Title] OR elderly[Title] OR review[Title] OR meta[Title]). The remaining databases used a combination of the same search terms. The search terms in Chinese were “functional training” and “athletes”, located in the article title.

### Inclusion and Exclusion Criteria

Inclusion criteria encompassed RCT with the control group receiving TRT or other routine training, distinct training plan, intervention duration-4 weeks, and study participants being healthy athletes engaged in regular training.The exclusion criteria include: non-RCTs, lack of training (encompassing TRT or routine training integrating TRT elements) in the control group, intervention duration <4 weeks, subjects not regular athletes, belonging to patient/recovery categories, and missing data.

### Data Extraction

Key data were extracted from the studies by the researchers, encompassing information such as the first author’s name and the publication year, details about the participants including their sex, athletic level, and age, as well as the sample size.

Additionally, intervention specifics were recorded, including the duration of the intervention, training frequency, training program and the study’s outcomes. The outcomes measured spanned across various indicators of strength and power, including the one-repetition maximum (1RM) for deep and half squat (DP/SP), grip strength (GS), and bench press (BP) for maximal strength[124]. Power was assessed through tests like the standing long jump (SLJ)[3,27], countermovement jump (CMJ) [9,71,100] and medicine ball throw(TMBP)[112]. Muscular endurance was evaluated by the number of 1-minute sit-ups (SU)[17,38], number of push-ups (PSU)[13,50], and number of pull-ups (PLU)[12].

### Quality Assessment

Two researchers independently assessed the quality of the included studies using the Revised Cochrane risk-of-bias tool for randomized trials (RoB 2)[111]. The risk of bias was scrutinized across six domains and seven specific evaluation criteria. Each criterion was categorized as presenting a “low risk,” “unclear,” or “high risk” of bias, determined by the context provided by the literature. To investigate the potential for publication bias among the studies, a funnel plot was employed. Any asymmetry observed in the funnel plot might suggest the influence of publication bias[110].

## 3. Statistical Analyses

We performed a meta-analysis to evaluate the comparative impact of FT versus TRT on outcomes related to muscle strength. The systematic data analysis was facilitated by RevMan software (version 5; Cochrane Collaboration, Oxford, UK)[103] and Stata (version 15; StataCorp, LP, College Station, TX)[109]. In instances where data was not readily available, efforts were made to contact the authors for the required information; studies that did not respond were excluded from the analysis. The random-effects model[21] was applied to compute and synthesize the effect sizes (either mean difference [MD] or standardized mean difference [SMD]) between the experimental and control groups[8]. The significance level was set at p < 0.05, with a 95% confidence interval. Effect sizes were interpreted as very small (MD/SMD < 0.2), small (MD/SMD = 0.2-0.5), medium (MD/SMD = 0.5-0.8), large (MD/SMD > 0.8[102]. Heterogeneity among the studies was assessed using the I^2^ statistic, with thresholds of 25%, 50%, and 75% representing low, moderate, and high levels of heterogeneity, respectively[54]. To evaluate the stability of significant findings against publication bias, a fail-safety N analysis was conducted, where higher values suggest greater robustness[83]. This analysis, in conjunction with a sensitivity analysis plot, offers a thorough assessment of the reliability and robustness of the study outcomes. Additionally, meta-regression and subgroup analyses were employed to pinpoint the primary sources of heterogeneity[97,118]. Publication bias was investigated using Egger’s test, with p<0.05 considered indicative of significant bias[84].

## 4. Results

After implementing the search strategy, we initially identified 15,050 articles from English databases and 3,126 from Chinese databases. After removing duplicates, 7,551 articles were left for further consideration. Screening of titles and abstracts narrowed down the pool to 670 relevant articles. Upon further evaluation, 595 articles were excluded due to factors such as the nature of the intervention, the health status of participants, and incomplete data. This process culminated in 75 articles fulfilling the inclusion criteria for quantitative analysis (Figure 1). The meta-analysis encompassed 67 articles with a total of 1,708 subjects, with 862 engaging in FT and 846 in TRT. The participants included 1,376 male and 332 female athletes. Additional details regarding the subjects and interventions are provided in Table 1.

**Fig. 1.**
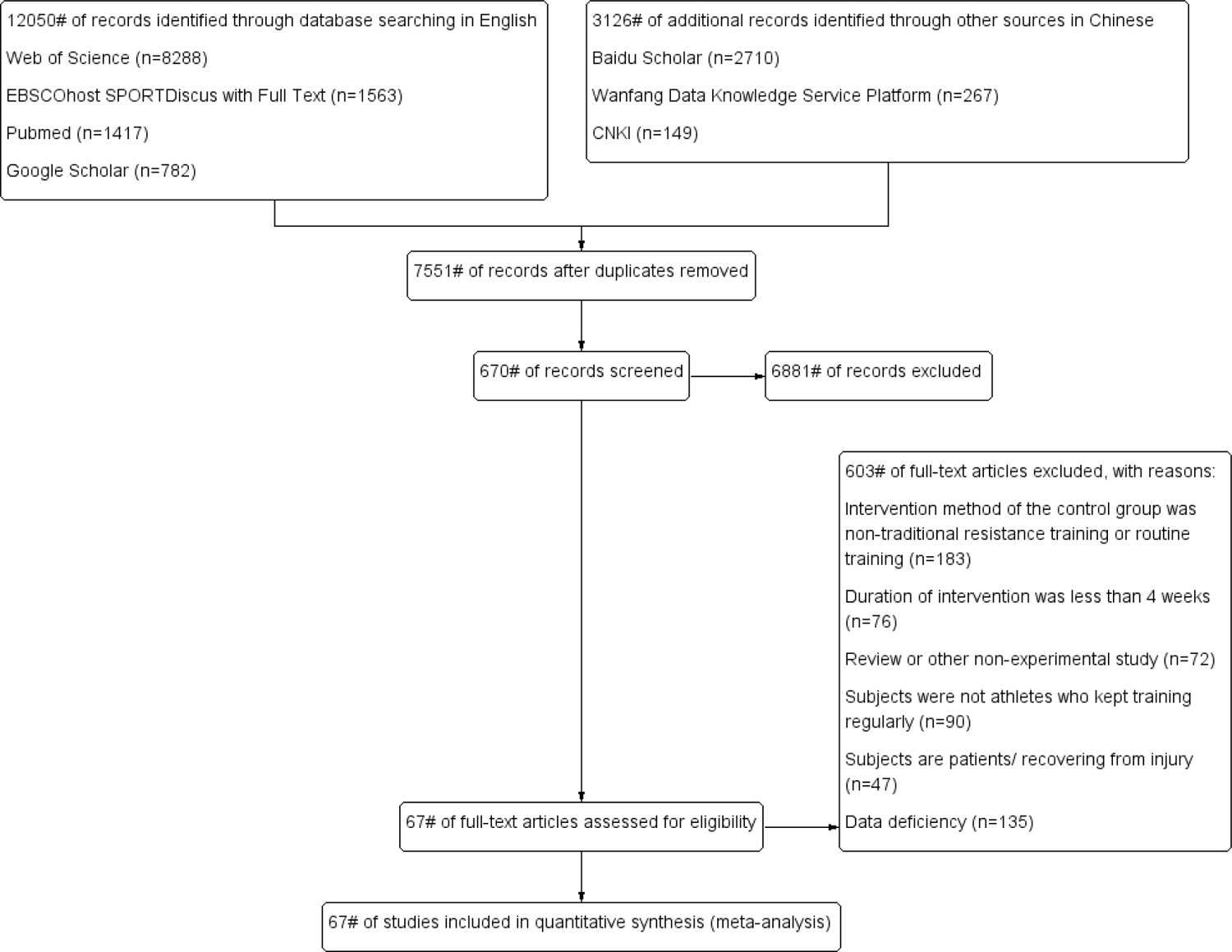
The PRISMA flow diagram shows the article selection process..

**Table 1.**
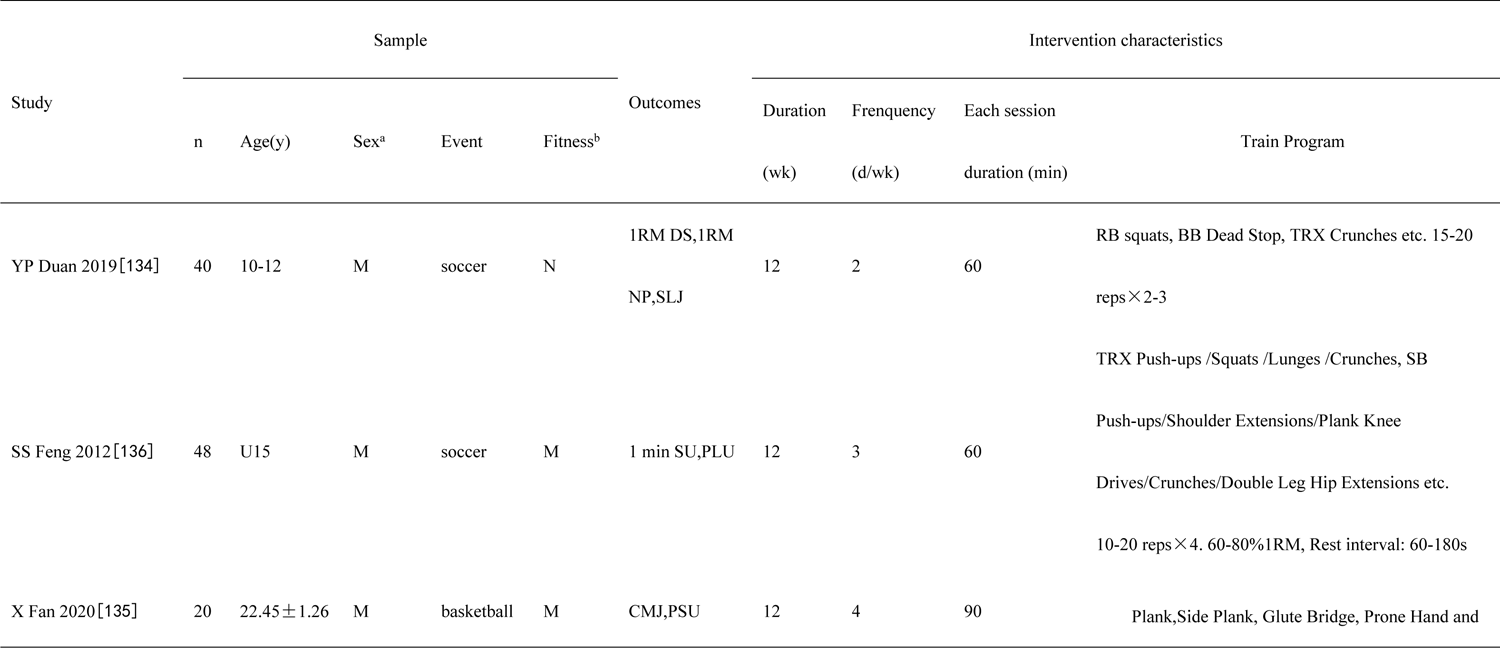
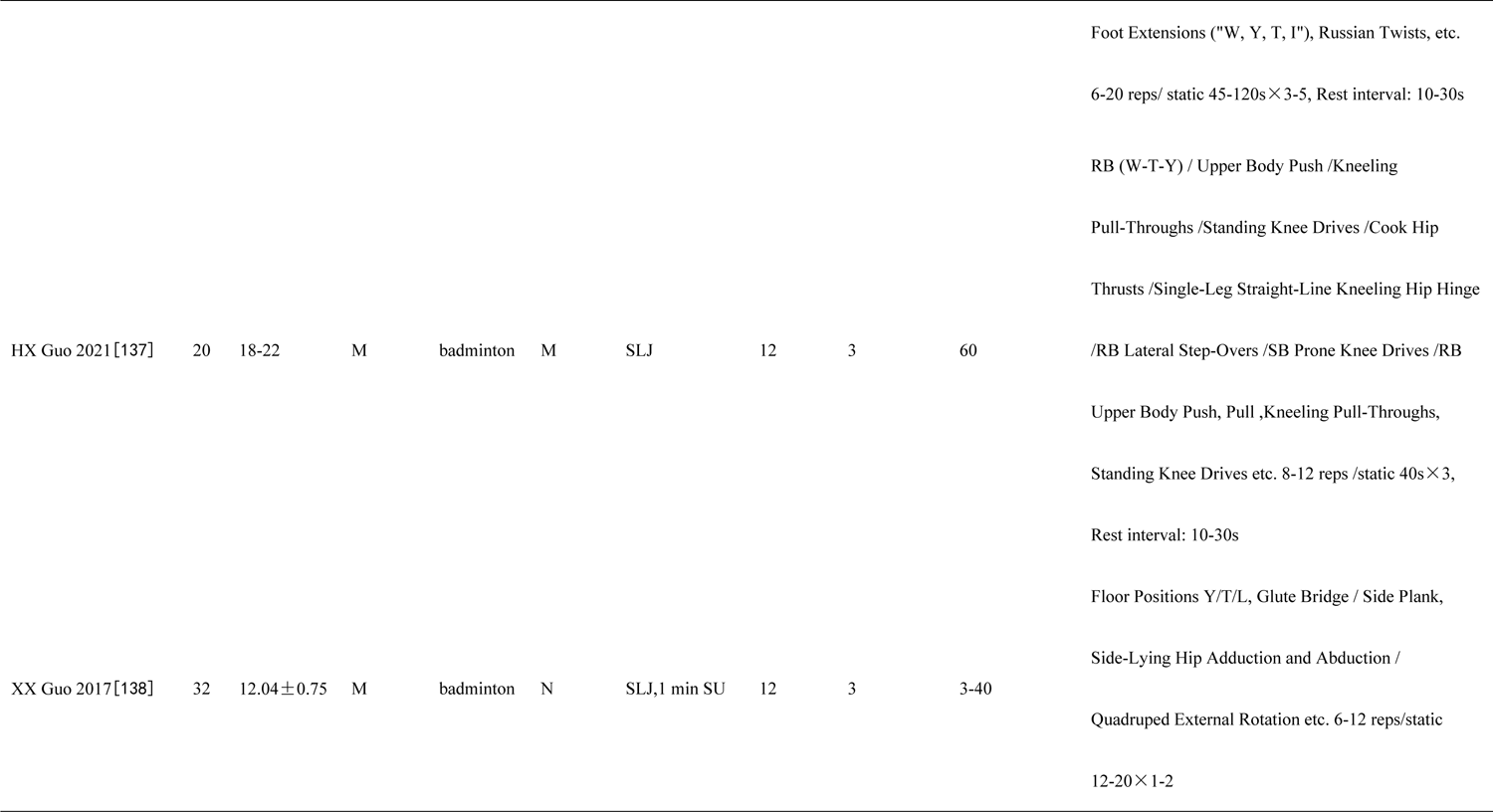
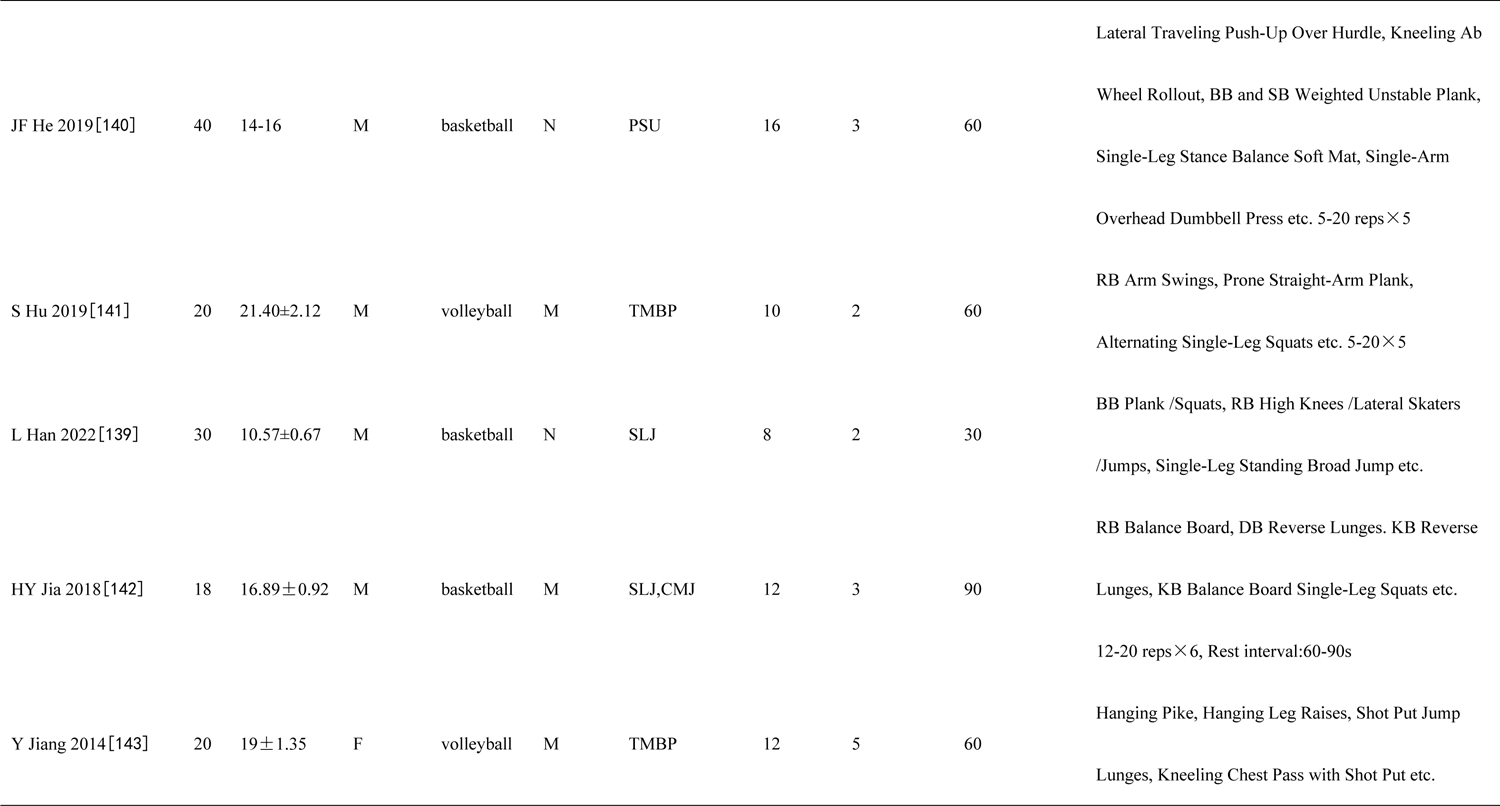
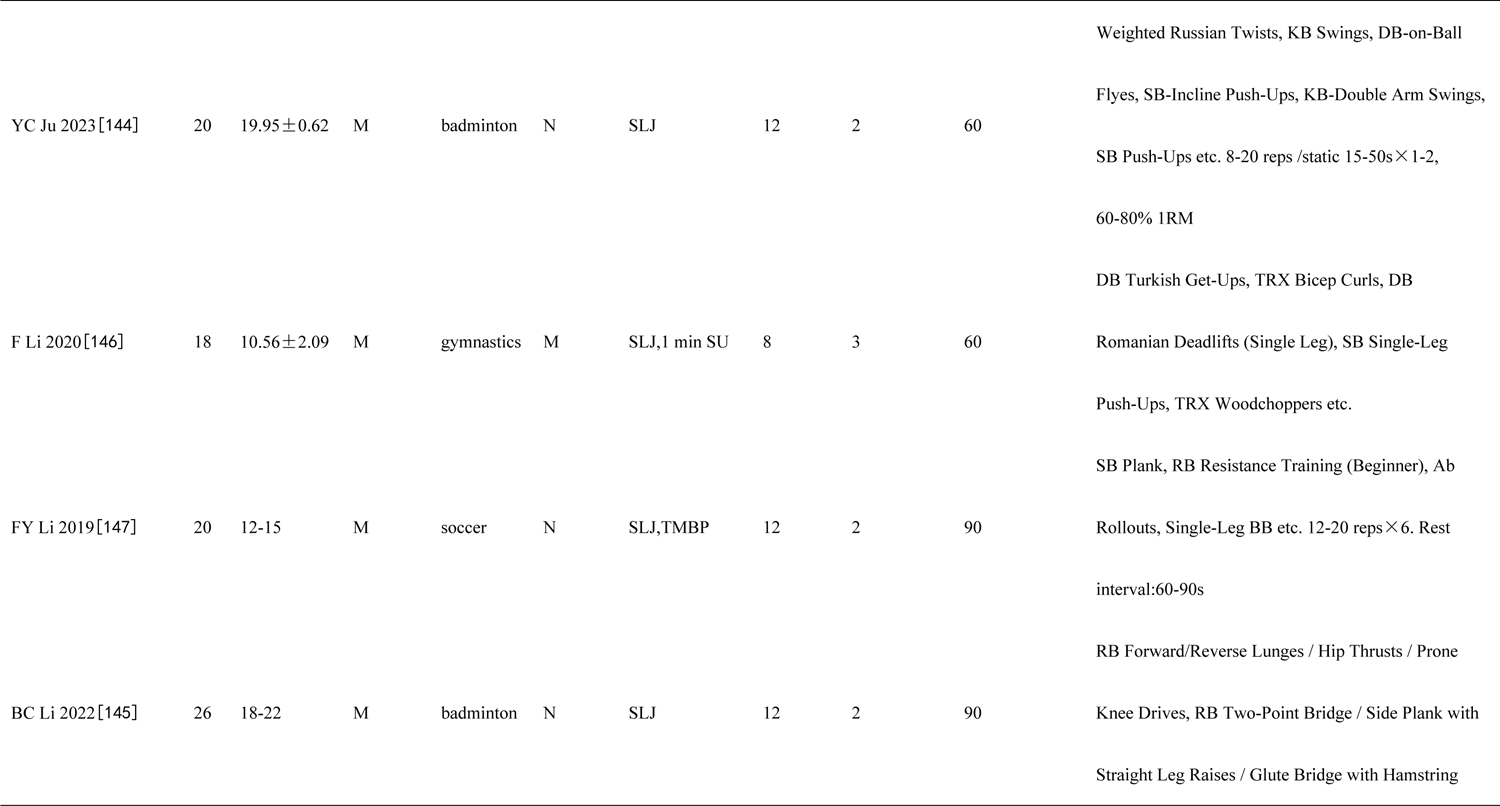
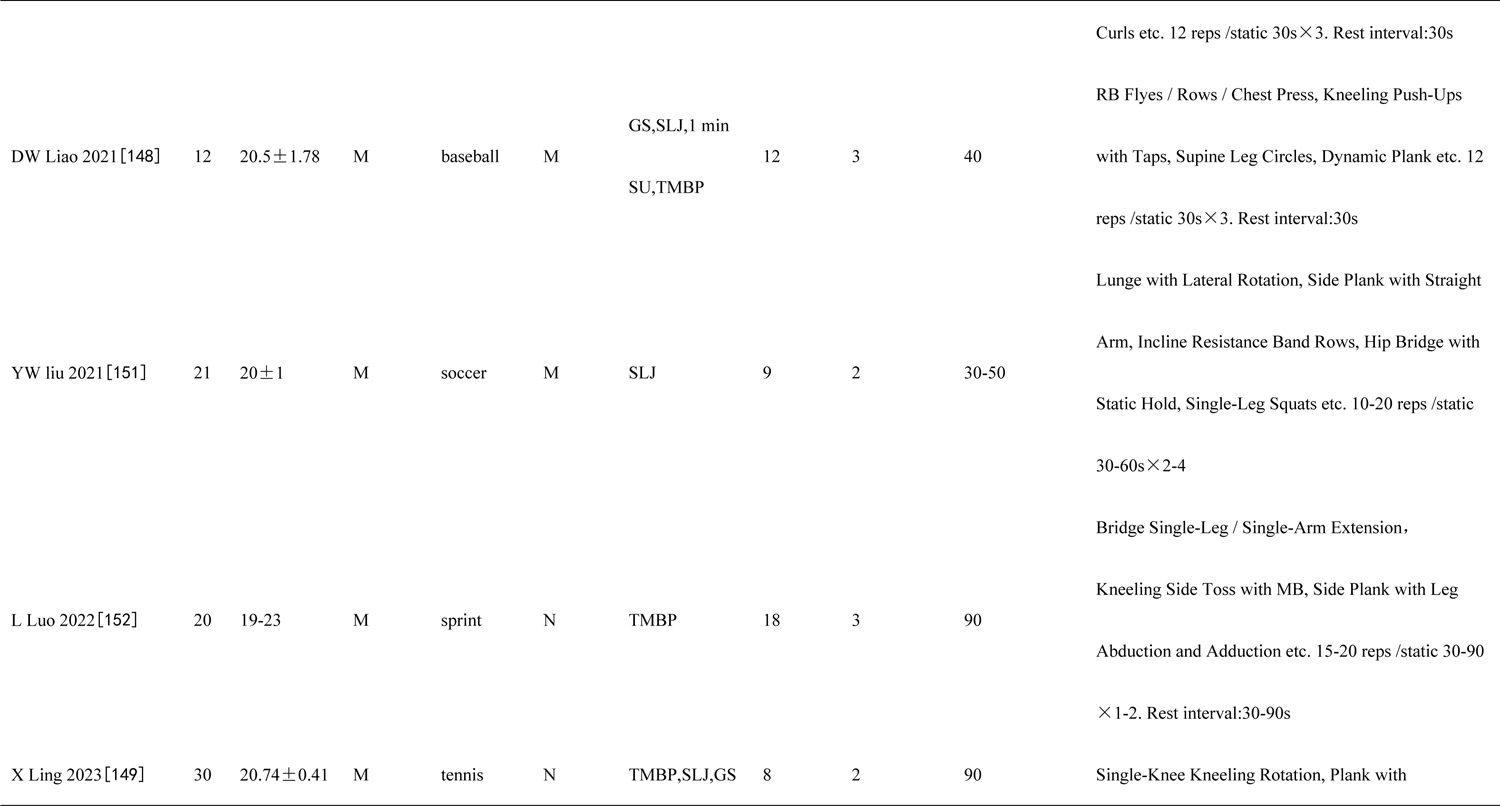
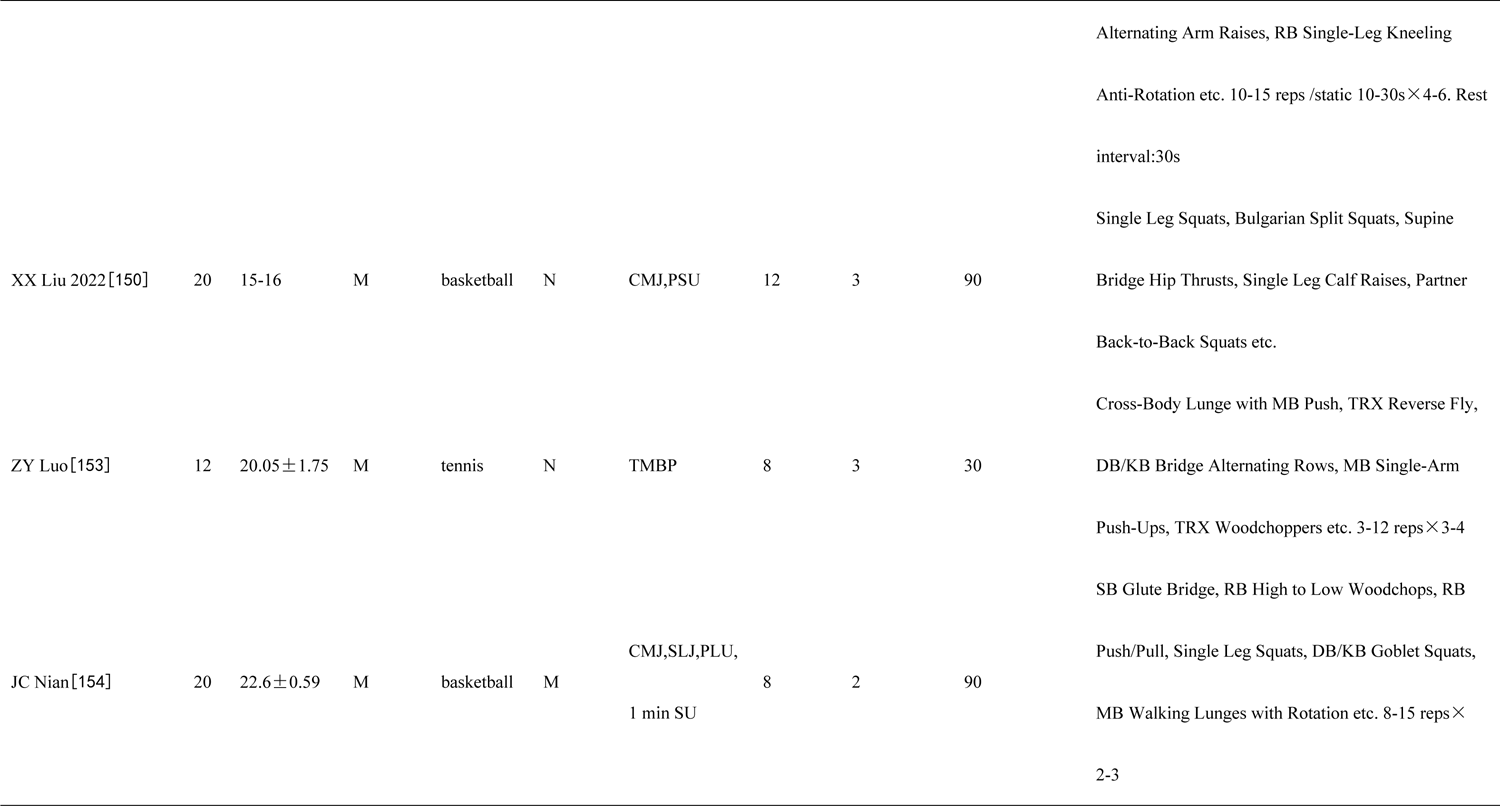
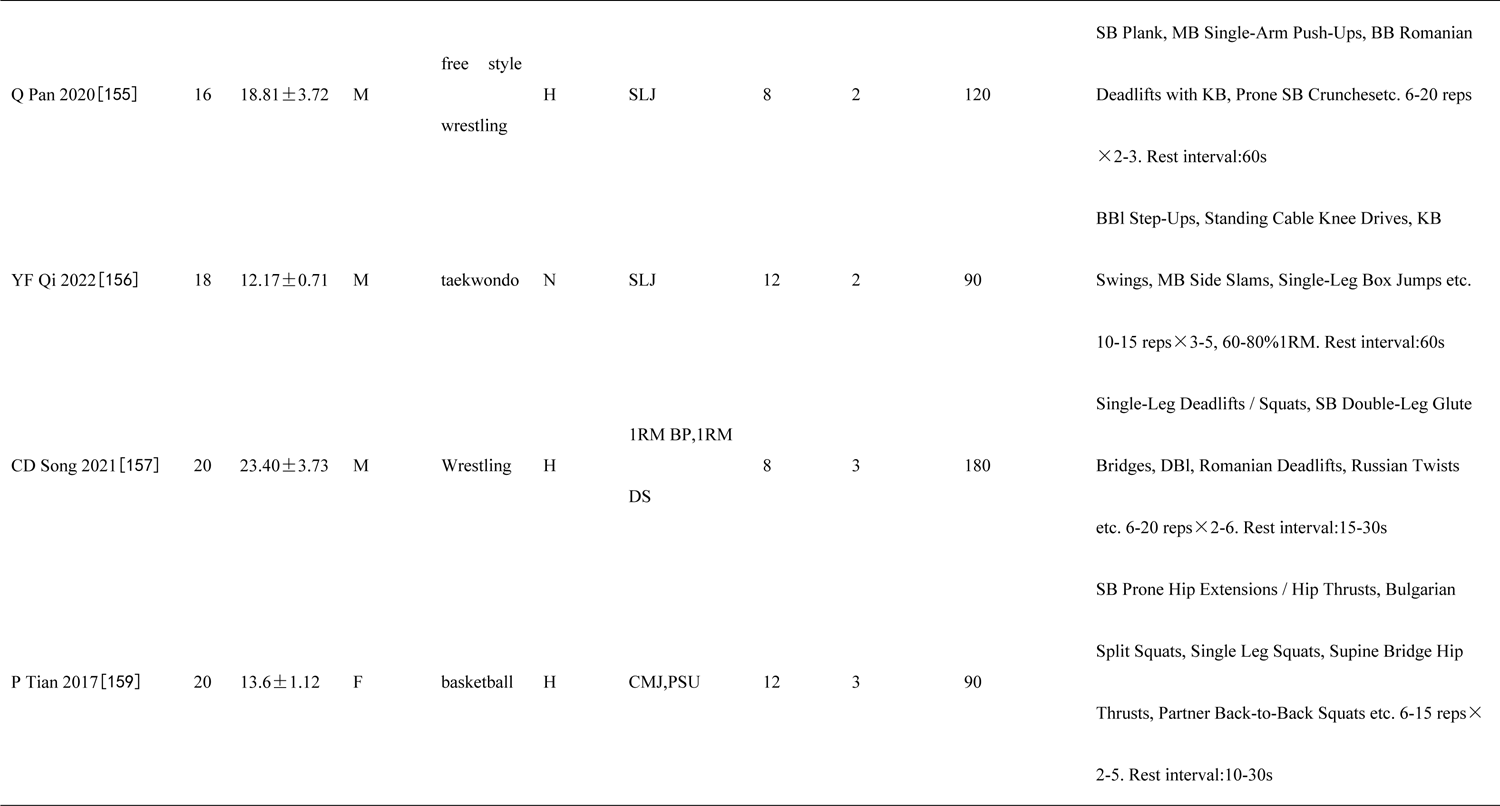
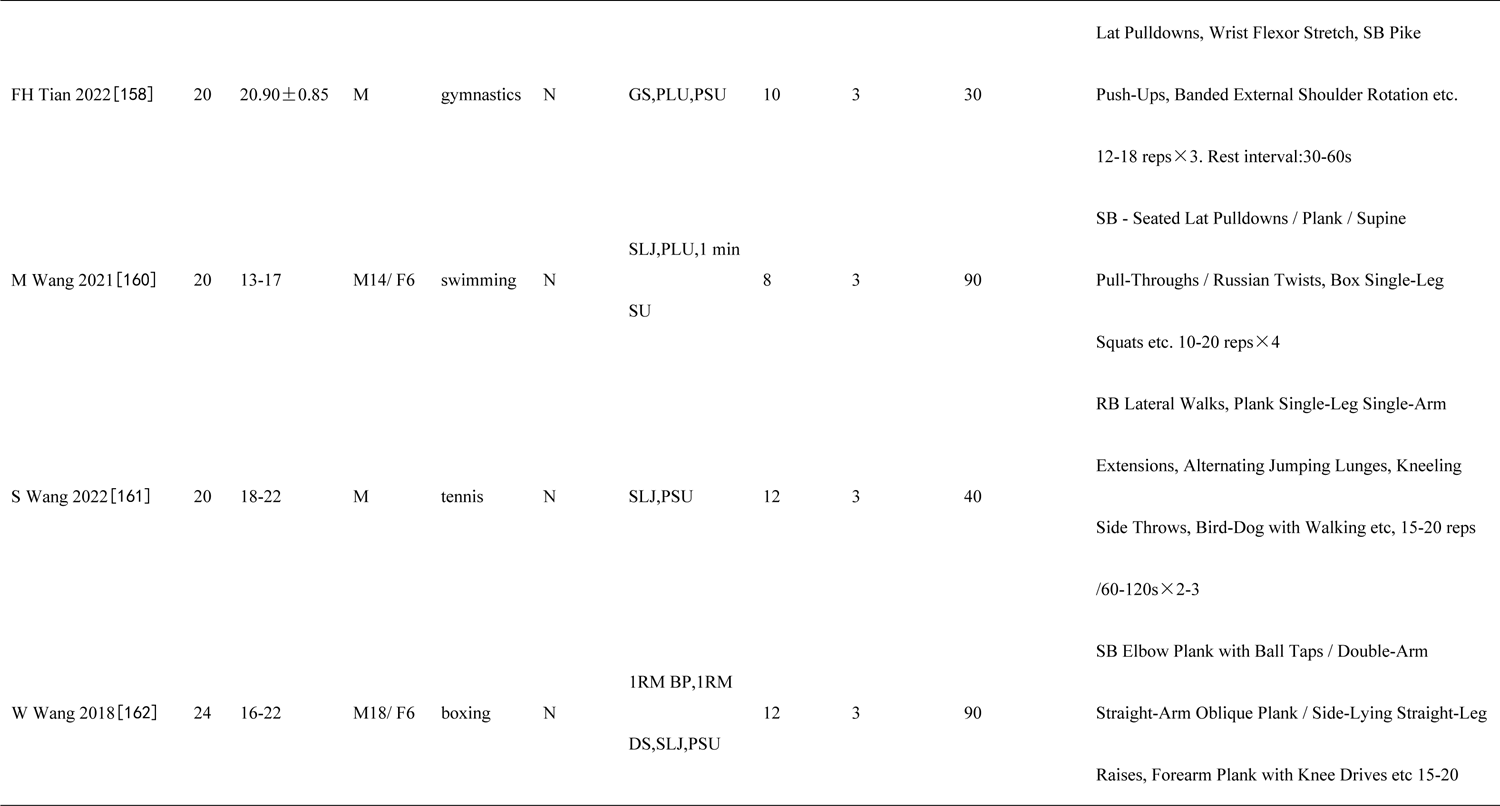
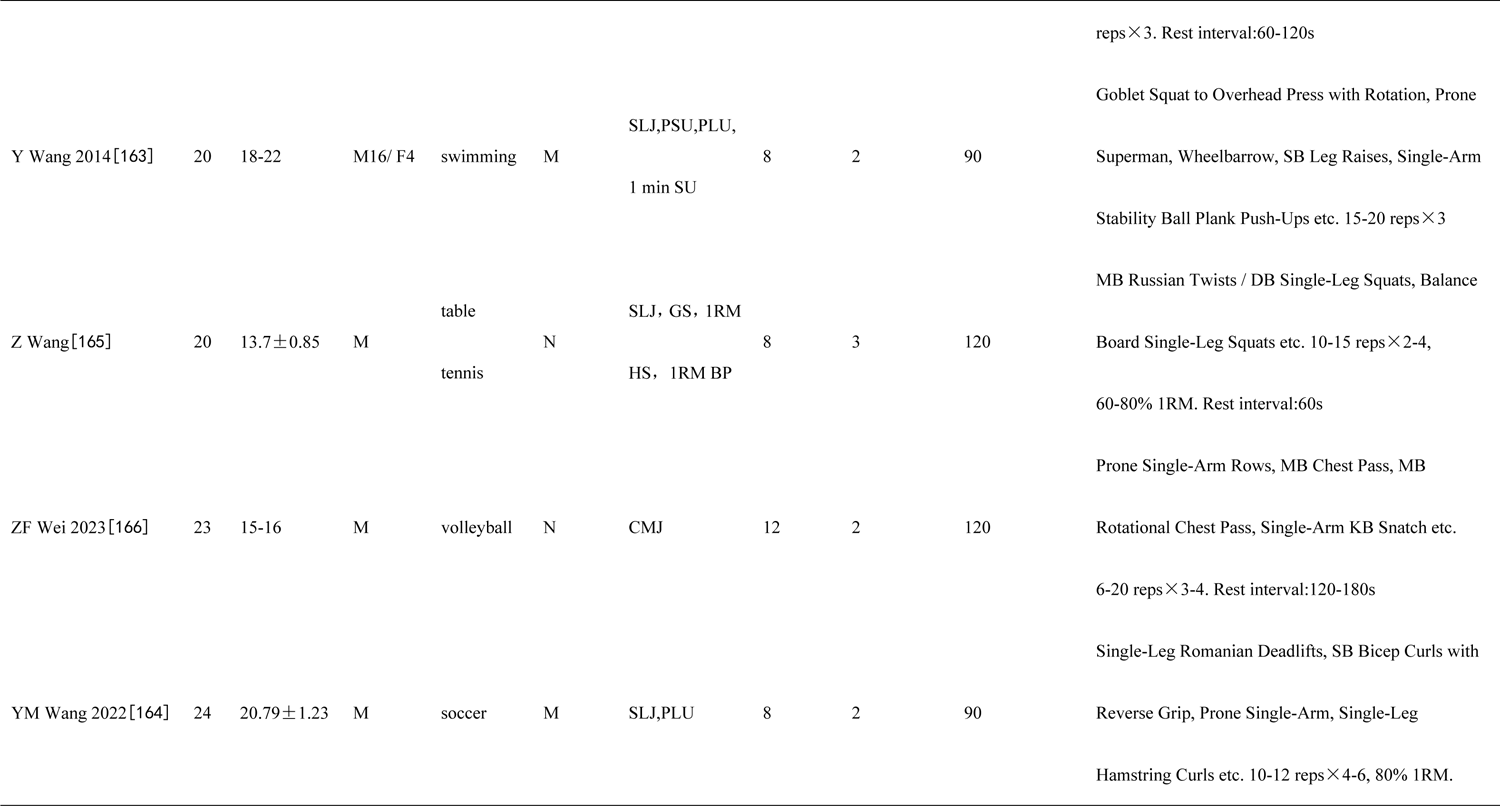
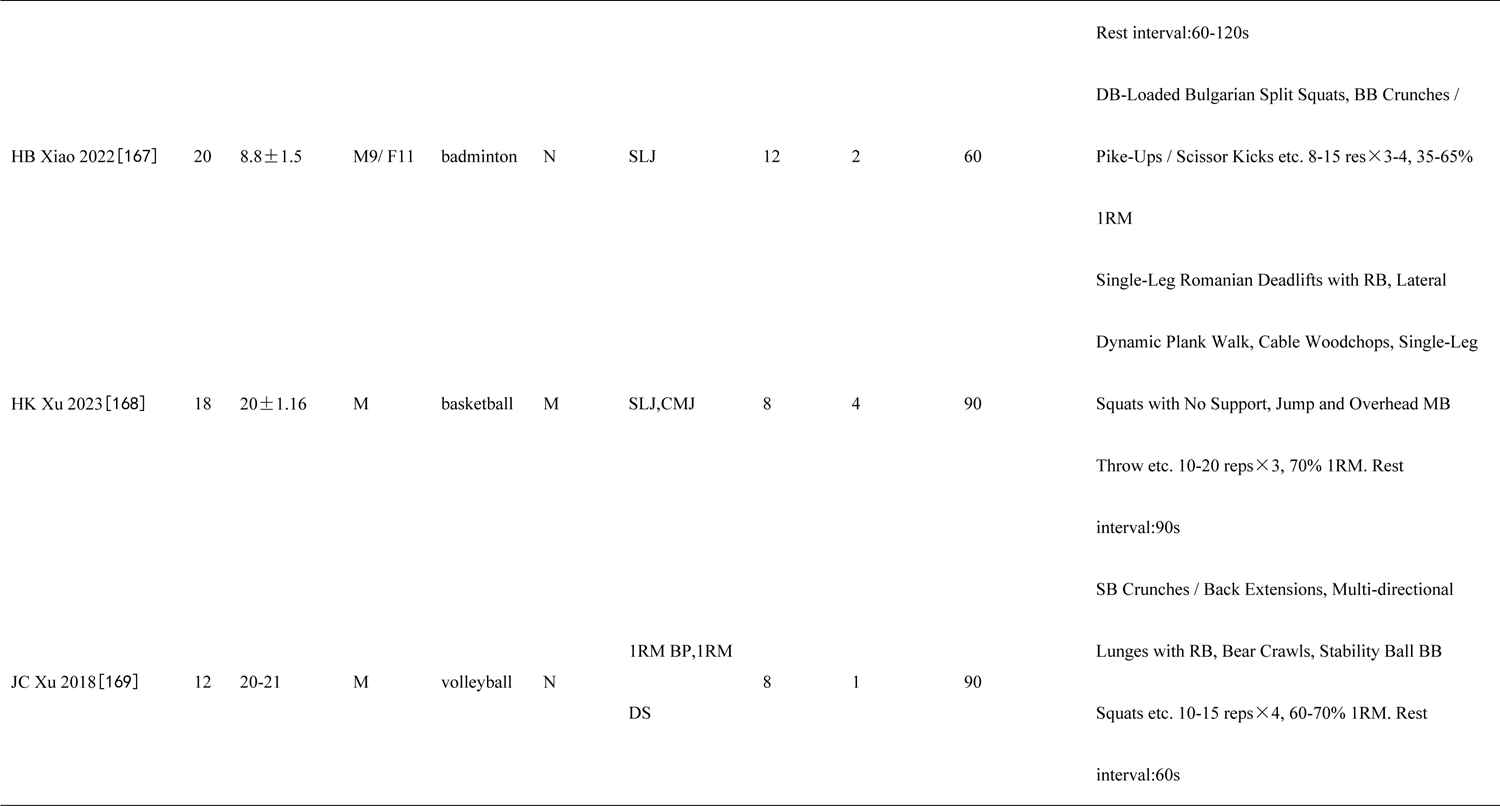
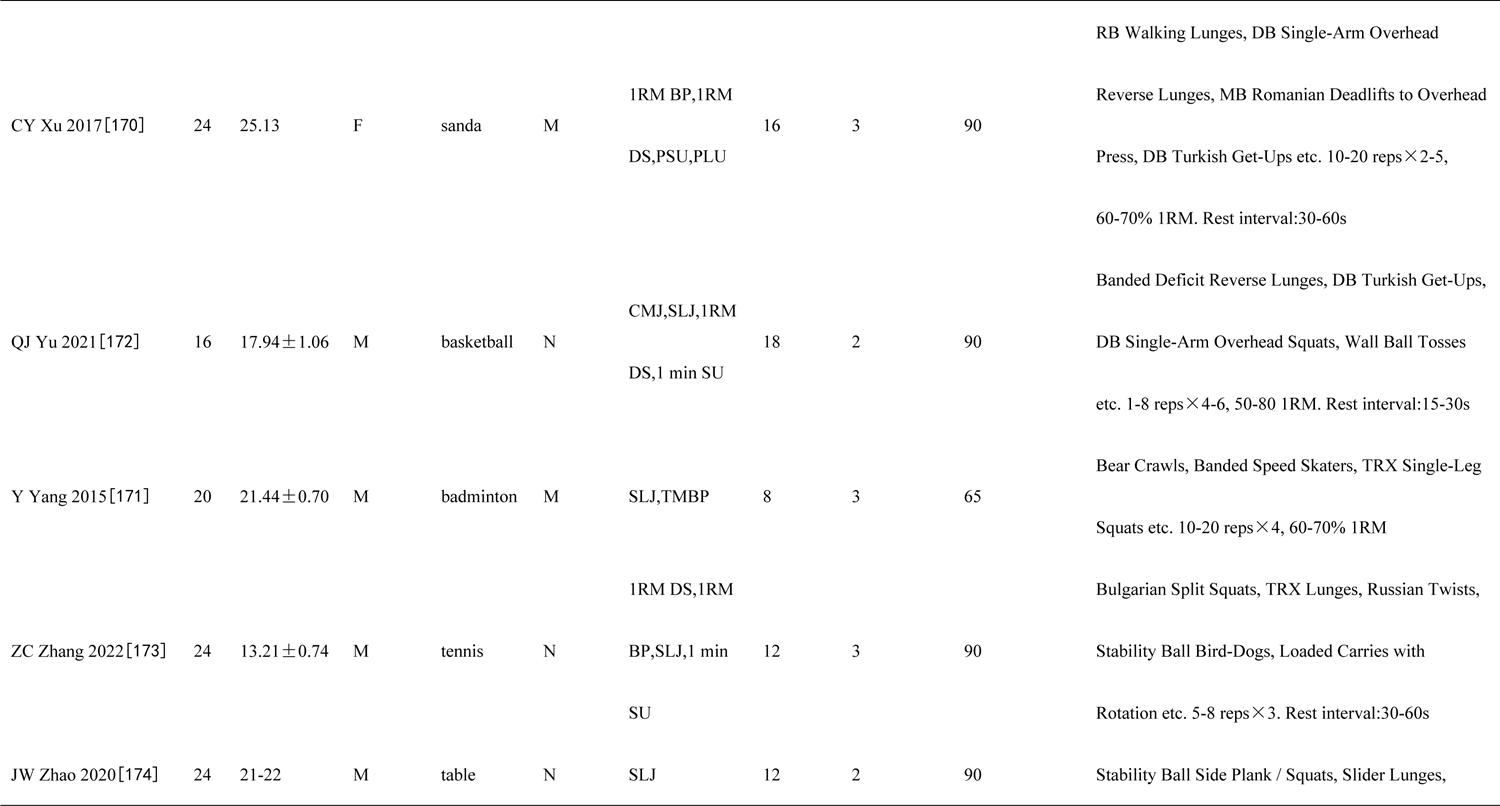
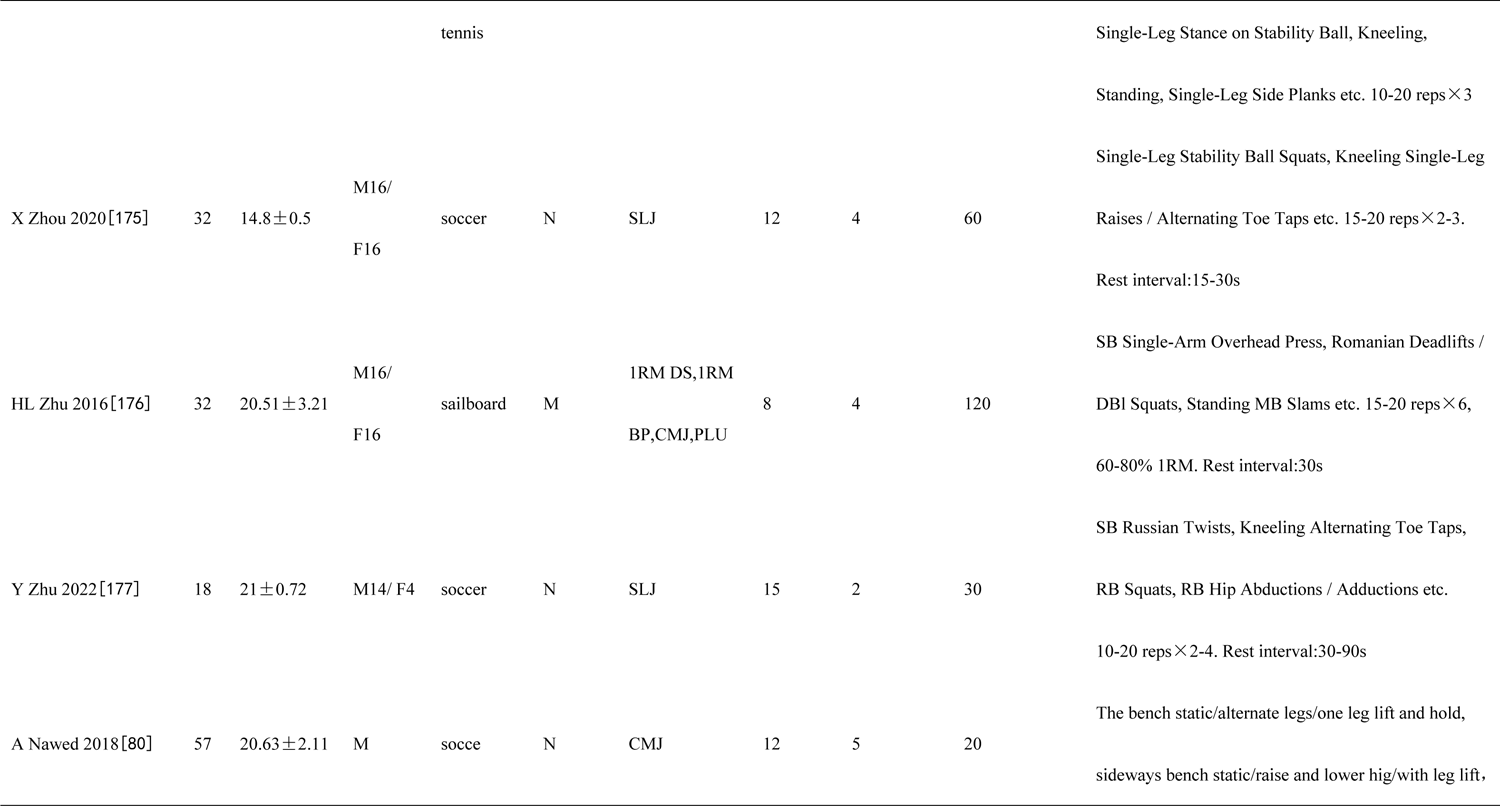
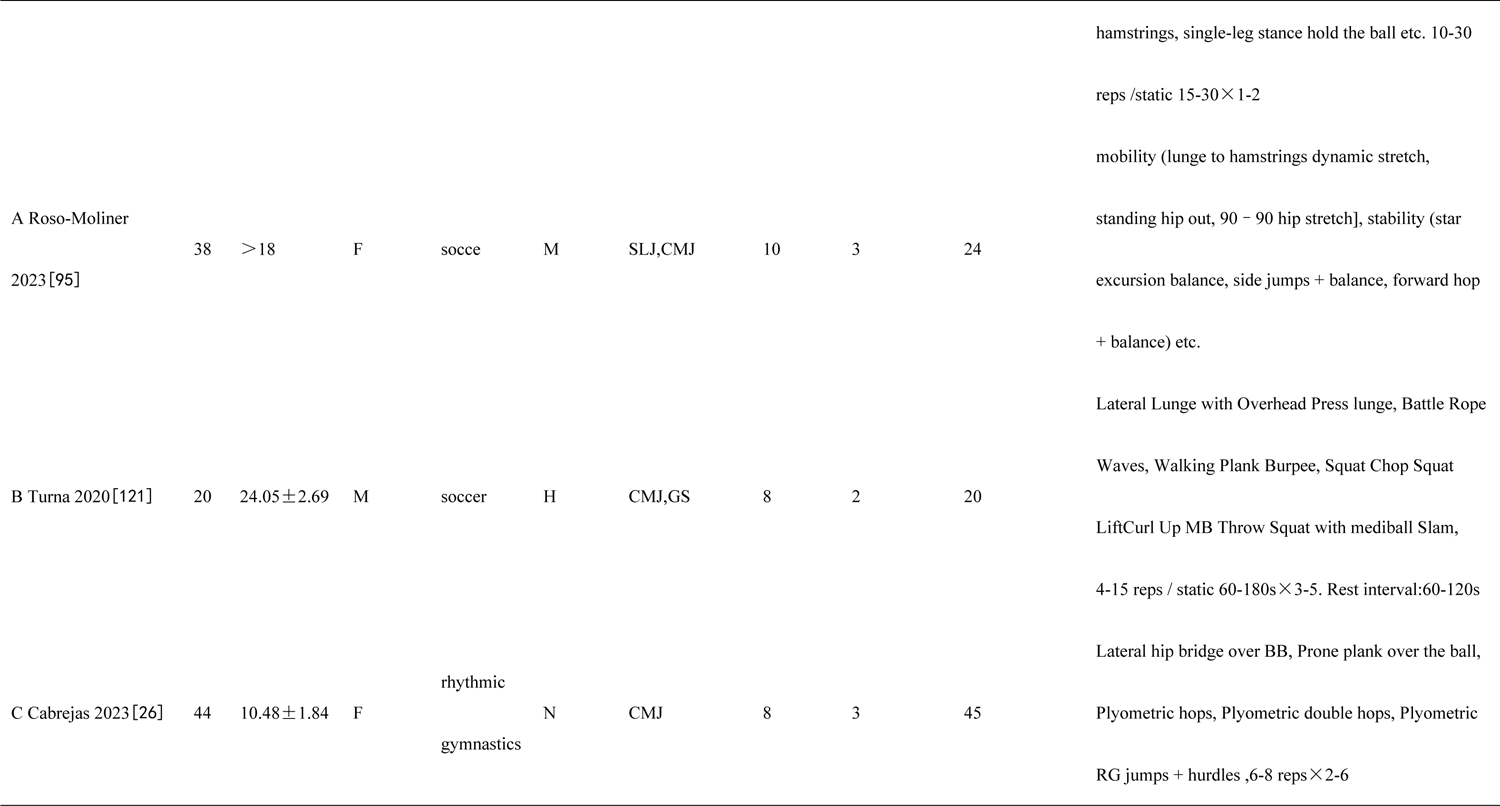
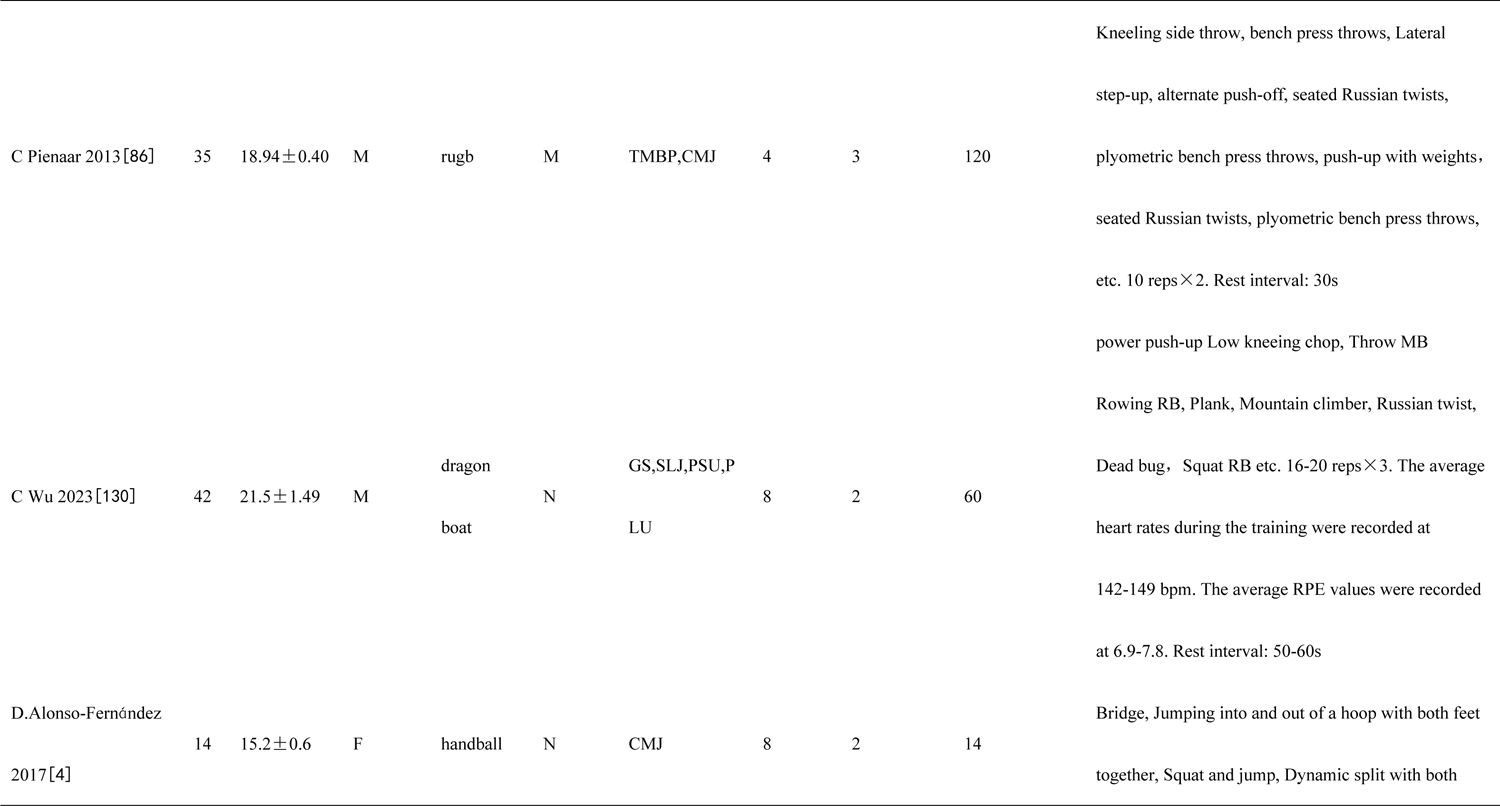
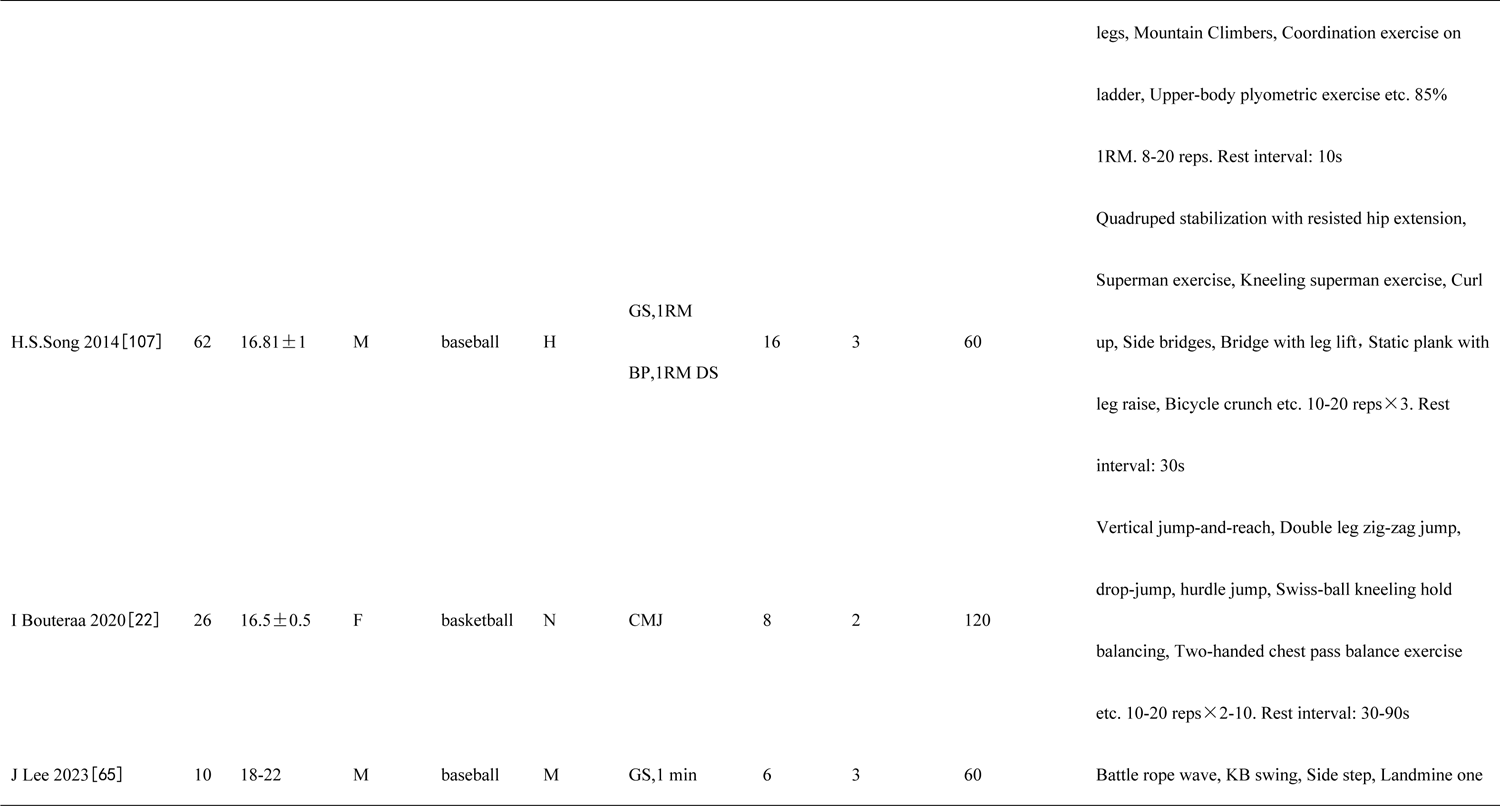
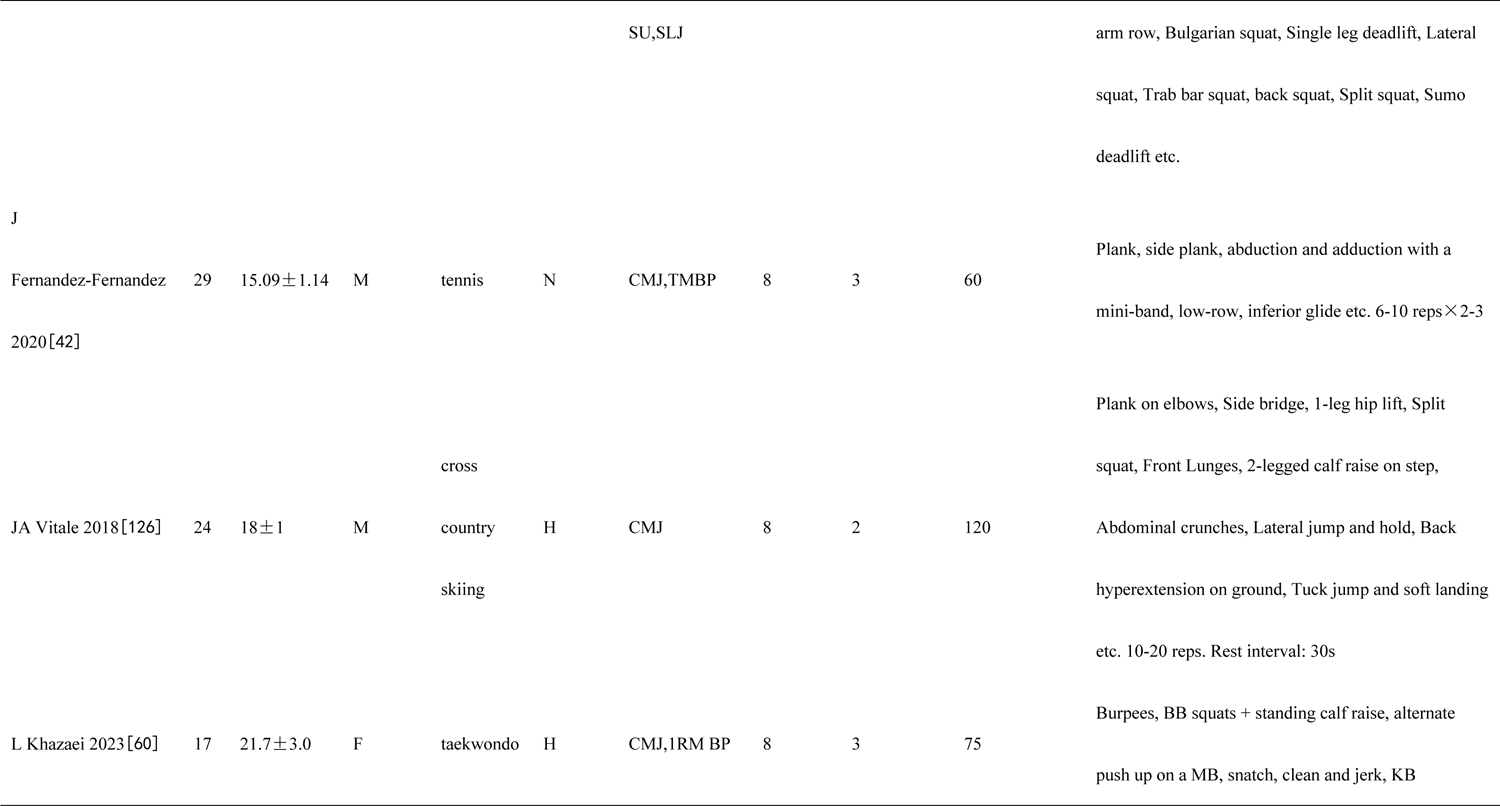
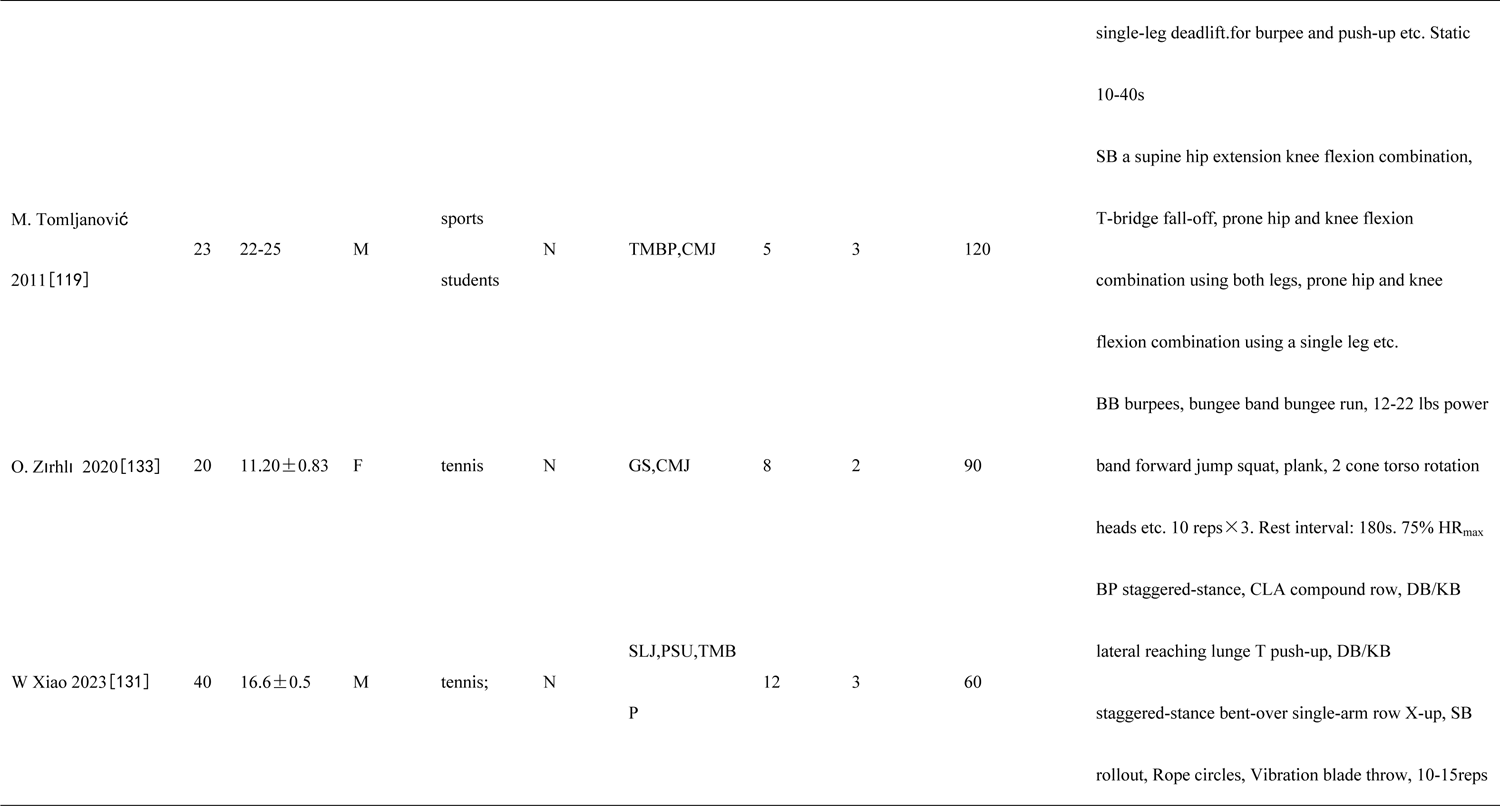
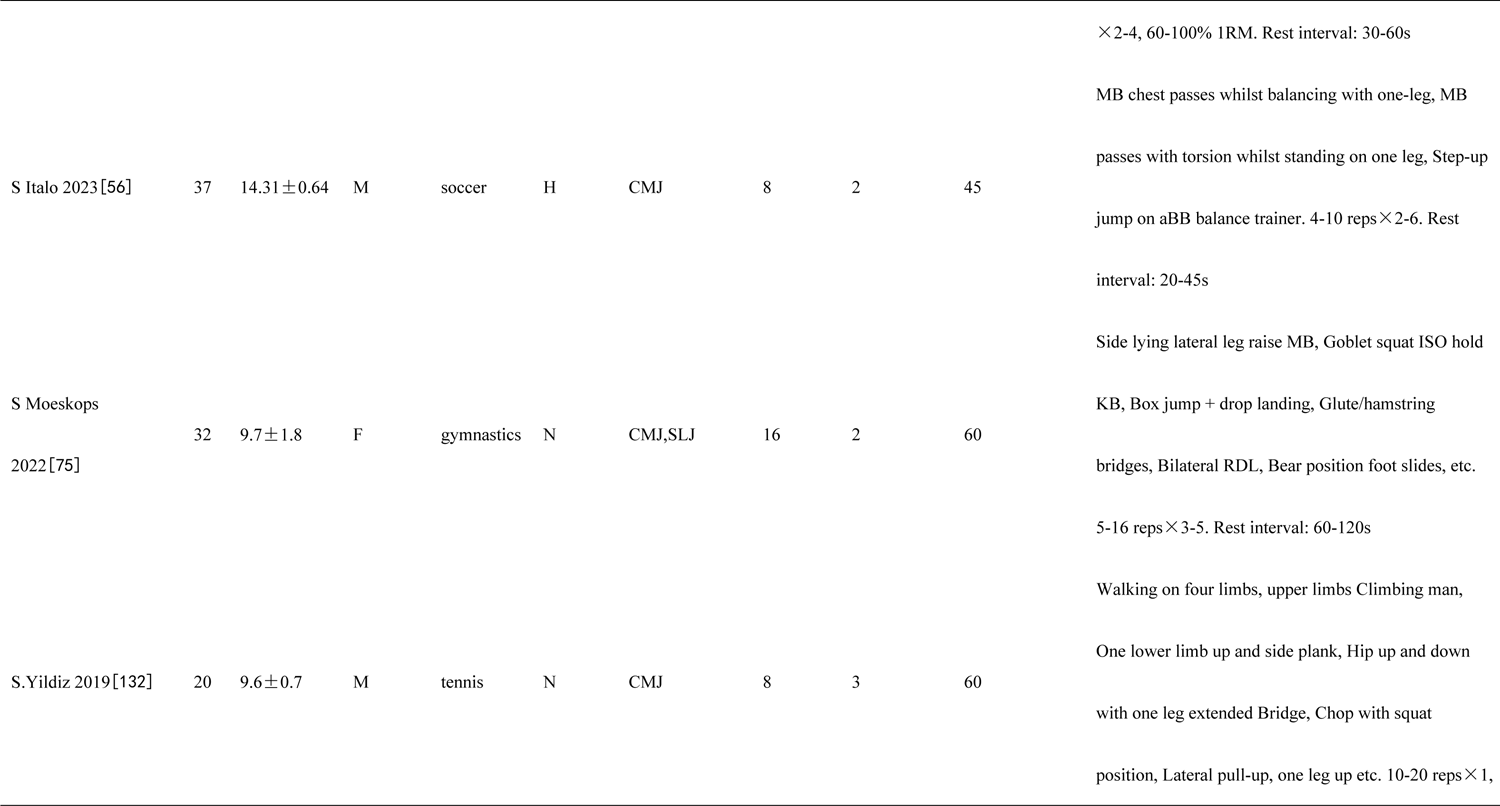
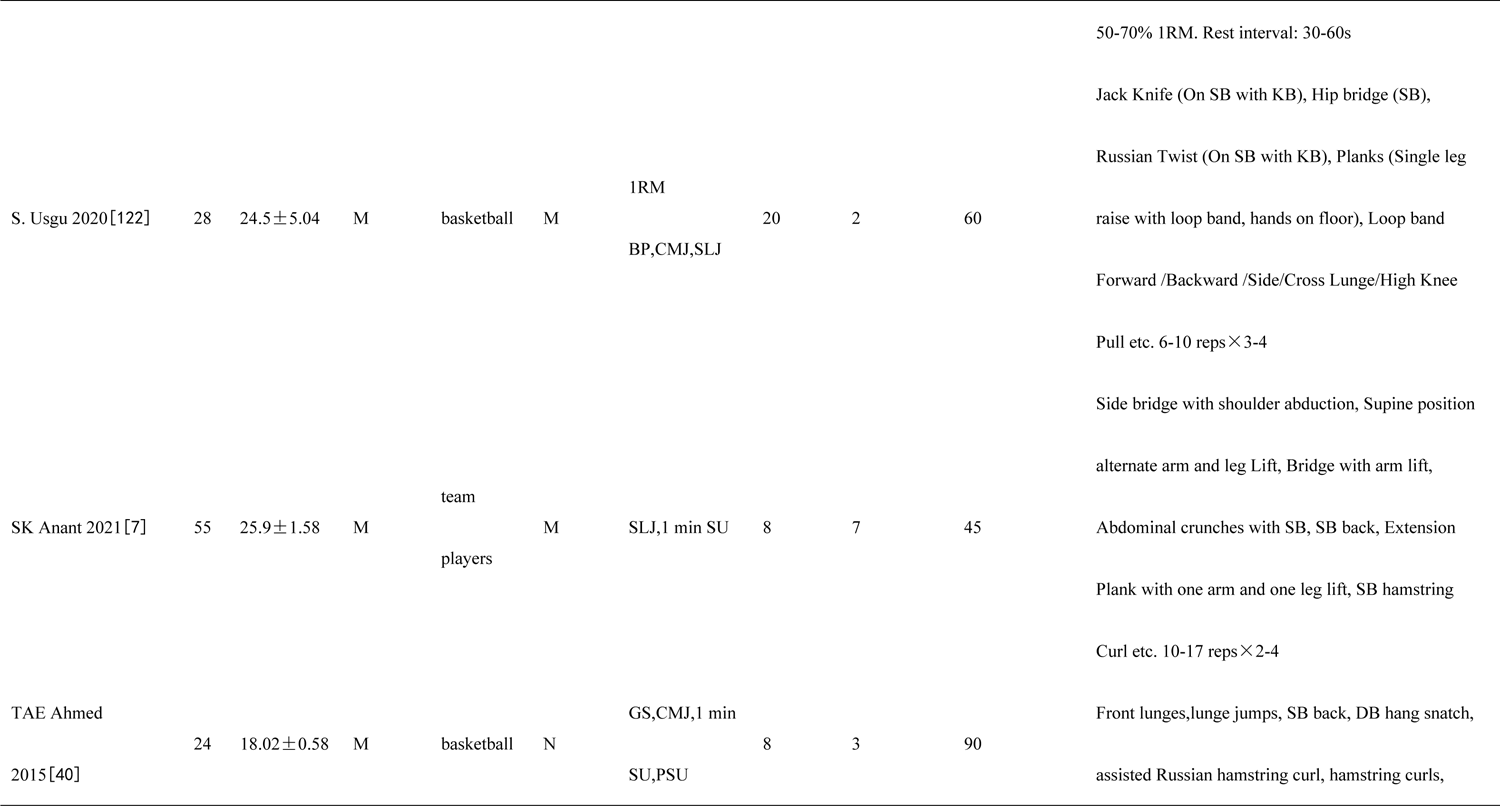
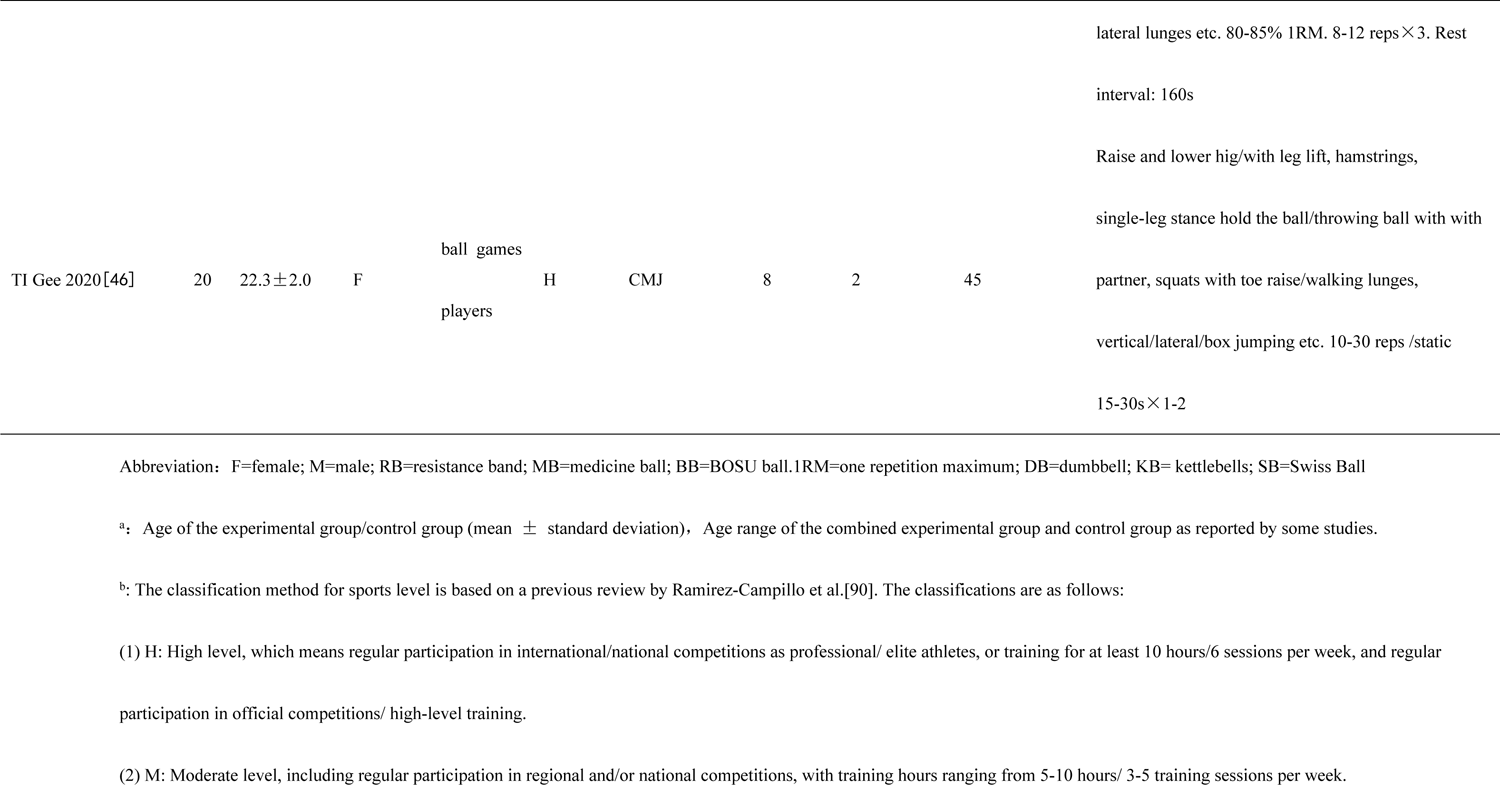
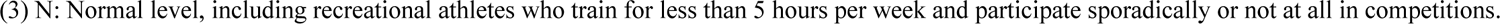
Characteristics of subjects included in the included studies.

### Quality Assessment

The PEDro Scale evaluates the methodological quality of RCT in sports science research[111]. Figure 2 indicates that the reviewed literature generally presents a moderate risk level. In the randomization process, 8 articles were considered low risk, representing 11.9% of the total[6,55,81,121,127,137,138,143]. All articles maintained low risk in areas such as participant identification, intervention adherence, and handling of missing outcome data. For outcome measurement, 19.4% articles was also deemed low risk[6,26,132,135,148,149,151–154,156,157,165]. In terms of result selection, 4.5% articles were classified as low risk[26,59,130]. 77.6% articles employed random allocation methods, with only 2 articles use double-blind methods (2.9%), and 9 are single-blind (13.4%). All studies applied appropriate analysis methods to assess their findings. The attrition rate for all literature is ≤15%, with only 2 articles reported adverse events such as athlete infection with the novel coronavirus[59] and sports injuries[131].

**Fig. 2.**
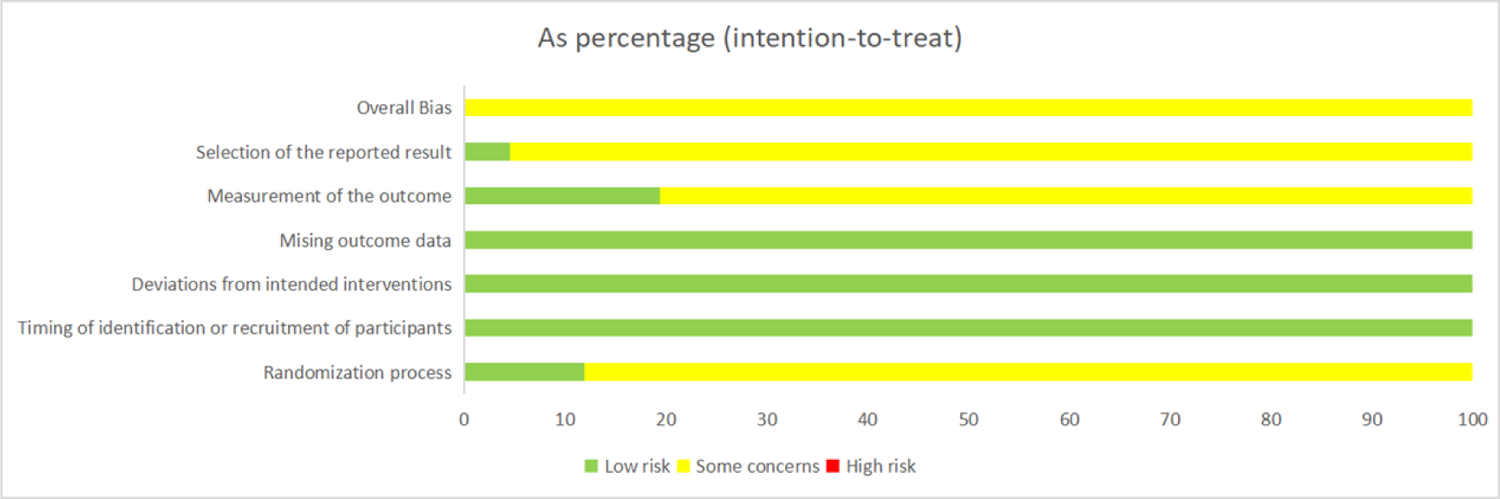
Literature quality evaluation form

### Meta-analysis

The influence of FT on athletes’ strength is detailed in Table 3, complemented by forest plots for each indicator in the supplementary figures. FT significantly enhanced maximal strength in the upper and arm limbs (p<0.05) and had a nearly significant impact on the lower limbs’ maximal strength (p=0.07). The effect sizes for these improvements were large(ES=2.01-3.12; Supplementary Figures 1-3). A notable increase in power was observed with FT (p≤0.001), with effect sizes varying from very small to moderate (ES=0.07-0.68; Supplementary Figures 4-6). Additionally, FT positively affected muscle endurance, as evidenced by significant improvements in the 1-minute SU and PSU (p<0.05), with a nearly significant change in the PLU (p=0.07) and effect sizes ranging from moderate to large (ES=0.74-4.13; Supplementary Figures 7-9). To evaluate publication bias, Egger’s test was employed (Supplementary Figures 10-18), with 8 out of 9 tests indicating no bias. However, for the CMJ analysis, a significant publication bias was identified and addressed using the trim and fill method.

**Table 3.**
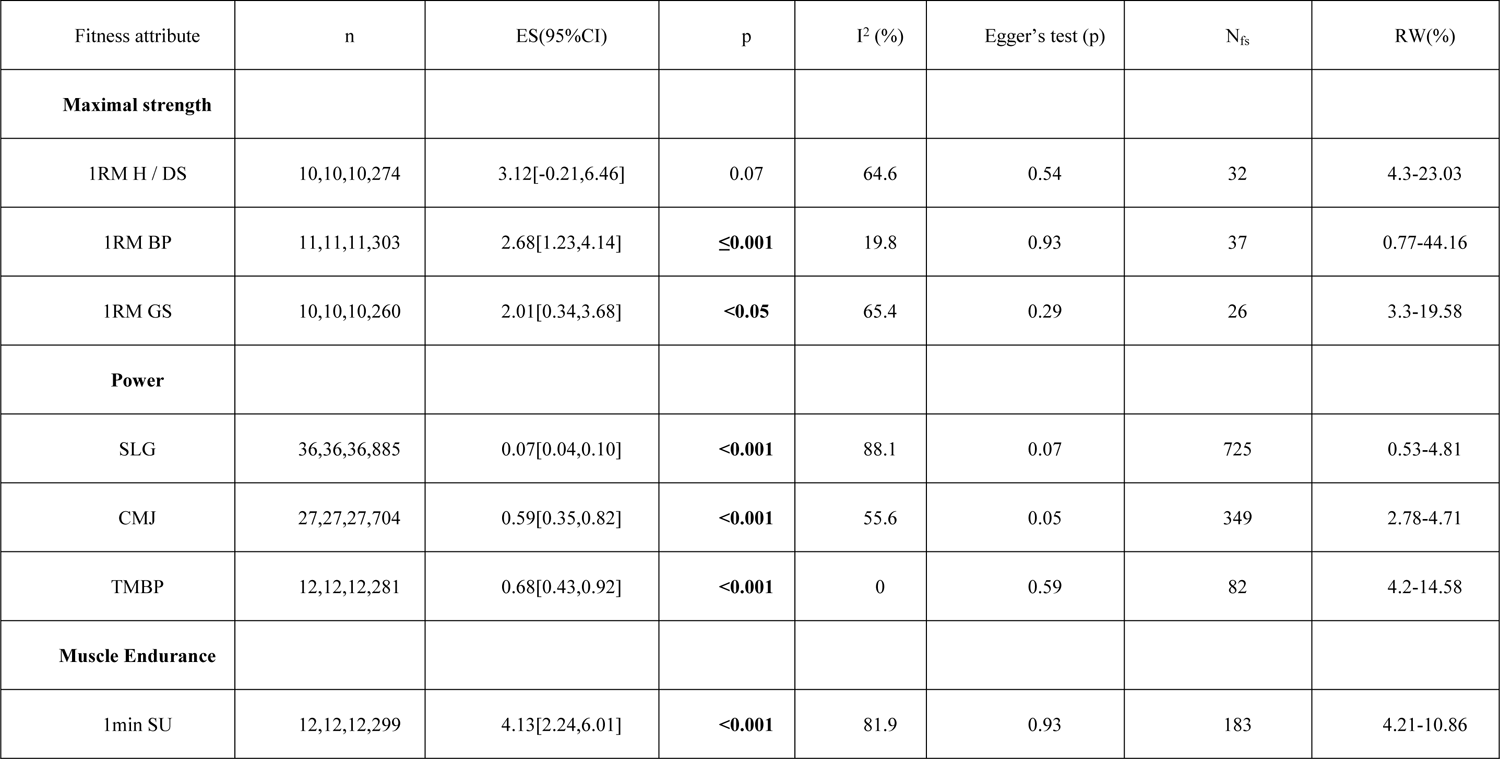
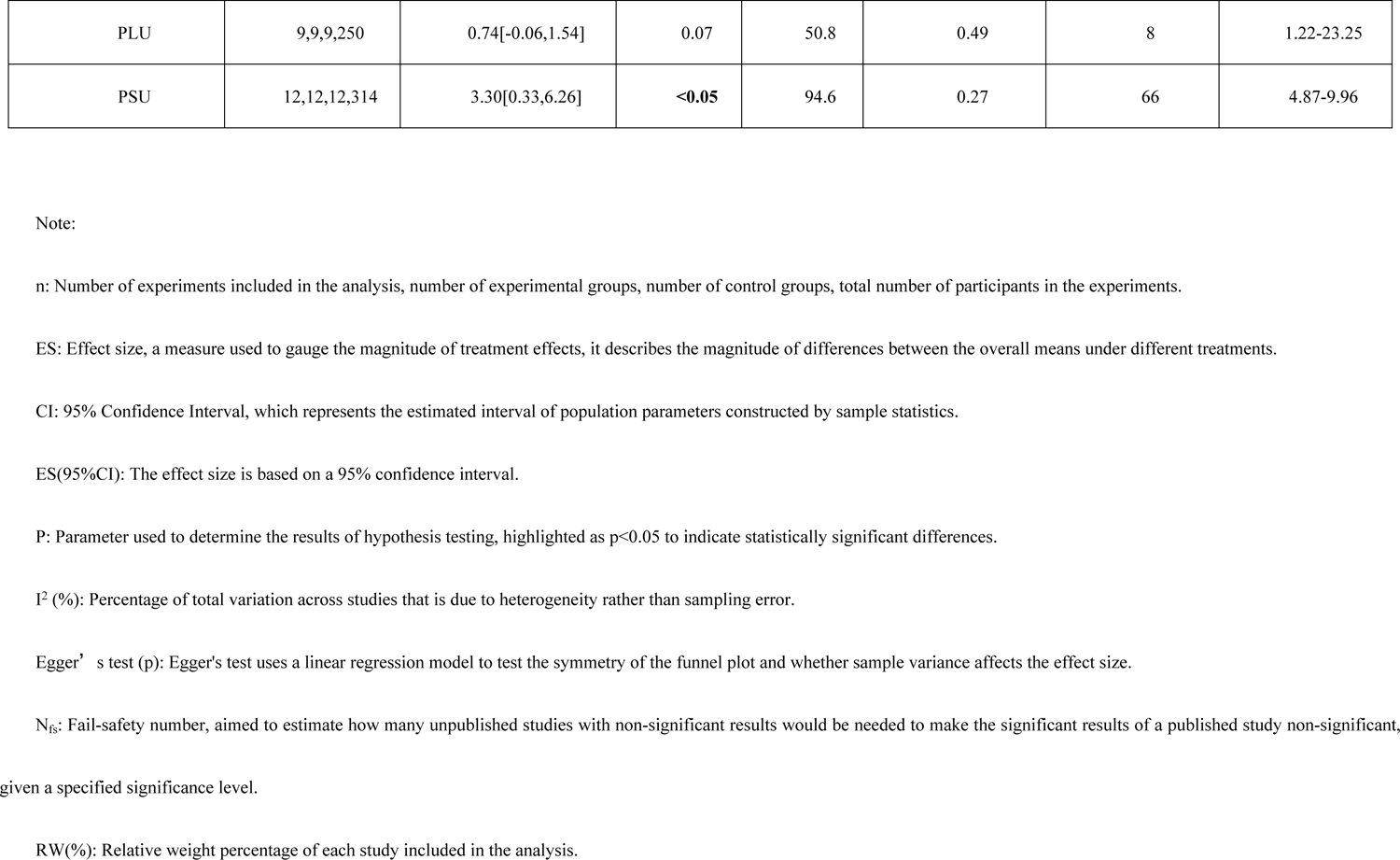
Summary of the effects of FT Vs. TT on muscle strength in athletes.

| Fitness attribute | n | ES(95%CI) | p | I<sup>2</sup> (%) | Egger's test (p) | N<sub>fs</sub> | RW(%) |
| --- | --- | --- | --- | --- | --- | --- | --- |
| **Maximal strength** |  |  |  |  |  |  |  |
| 1RM H / DS | 10,10,10,274 | 3.12[-0.21,6.46] | 0.07 | 64.6 | 0.54 | 32 | 4.3-23.03 |
| 1RM BP | 11,11,11,303 | 2.68[1.23,4.14] | ≤**0.001** | 19.8 | 0.93 | 37 | 0.77-44.16 |
| 1RM GS | 10,10,10,260 | 2.01[0.34,3.68] | **<0.05** | 65.4 | 0.29 | 26 | 3.3-19.58 |
| **Power** |  |  |  |  |  |  |  |
| SLG | 36,36,36,885 | 0.07[0.04,0.10] | **<0.001** | 88.1 | 0.07 | 725 | 0.53-4.81 |
| CMJ | 27,27,27,704 | 0.59[0.35,0.82] | **<0.001** | 55.6 | 0.05 | 349 | 2.78-4.71 |
| TMBP | 12,12,12,281 | 0.68[0.43,0.92] | **<0.001** | 0 | 0.59 | 82 | 4.2-14.58 |
| **Muscle Endurance** |  |  |  |  |  |  |  |
| 1min SU | 12,12,12,299 | 4.13[2.24,6.01] | **<0.001** | 81.9 | 0.93 | 183 | 4.21-10.86 |
| PLU | 9,9,9,250 | 0.74[-0.06,1.54] | 0.07 | 50.8 | 0.49 | 8 | 1.22-23.25 |
| PSU | 12,12,12,314 | 3.30[0.33,6.26] | **<0.05** | 94.6 | 0.27 | 66 | 4.87-9.96 |

Note:
n: Number of experiments included in the analysis, number of experimental groups, number of control groups, total number of participants in the experiments.
ES: Effect size, a measure used to gauge the magnitude of treatment effects, it describes the magnitude of differences between the overall means under different treatments.
CI: 95% Confidence Interval, which represents the estimated interval of population parameters constructed by sample statistics.
ES(95%CI): The effect size is based on a 95% confidence interval.
P: Parameter used to determine the results of hypothesis testing, highlighted as $p < 0.05$ to indicate statistically significant differences.
$I^2$ (%): Percentage of total variation across studies that is due to heterogeneity rather than sampling error.
Egger's test (p): Egger's test uses a linear regression model to test the symmetry of the funnel plot and whether sample variance affects the effect size.
$N_g$ : Fail-safety number, aimed to estimate how many unpublished studies with non-significant results would be needed to make the significant results of a published study non-significant, given a specified significance level.
RW(%): Relative weight percentage of each study included in the analysis.

### Sensitivity Analysis

Sensitivity analysis targeted studies with p<0.1 and I^2^>=50% to assess the stability of the findings (Supplementary Figures 19-27). Exclusion of individual studies did not alter the significant differences, and effect sizes stayed within the 95% confidence interval, demonstrating the robustness of the overall results.

### Subgroup Analysis

Subgroup analyses, each comprising more than 3 studies, were conducted to assess the impact of FT on muscle strength, with sports categorized based on significant heterogeneity levels (I^2^ > 50% and p < 0.1). Ball sport athletes in our study include individuals participating in sports such as soccer, basketball, tennis, baseball, and volleyball. Each of these sports, while involving a ball, demands different physical skills and movements. For example, soccer primarily involves kicking and running, basketball includes jumping and rapid directional changes, and baseball focuses on throwing and batting. These differences highlight the need for tailored training regimens. Our subgroup analysis revealed that ball sport athletes experienced notably greater improvements in muscle strength due to FT compared to athletes from other events(Supplementary Figures 28-34). Specifically, ball sport athletes showed significant advantages in maximal strength (ES=1.94, p≤0.001) over combat sport athletes (p > 0.05). In the domain of power, ball sport athletes (ES=0.09, p < 0.001) outperformed both water sport athletes (p > 0.05) and combat sport athletes (p > 0.05). For muscle endurance, ball sport athletes exhibited more pronounced improvements (ES=0.28, p < 0.05) compared to water sport athletes (p > 0.05). Further analysis (Supplementary Figures 35) highlighted that tennis, soccer, and basketball players saw more substantial enhancements in power following FT compared to athletes in badminton, baseball, and table tennis. The respective effect sizes and p-values were: tennis(ES=0.14, p < 0.05), soccer (ES=0.06, p < 0.05), basketball (ES=0.1, p < 0.05), and badminton, baseball, table tennis (p > 0.05). By defining ball sport athletes and providing examples, we acknowledge the substantial differences in the tasks and physical demands of each sport. This specificity allows for a more accurate interpretation of the impact of FT across diverse athletic activities. *Single (Training) Factor Analysis* The analysis of training factors (Supplementary Figures 36-60) indicates that FT tends to be more effective in younger athletes than in their older counterparts. Our data revealed that extended training sessions, defined as those lasting more than 60 minutes per session, are associated with more significant improvements in maximal strength and muscular endurance. Conversely, shorter sessions, typically under 60 minutes, are more beneficial for power enhancement. The duration of training periods in our study ranged 4-24 weeks. The mean duration across all studies was 12 weeks. Training frequencies varied between 2-5 sessions per week, with the mean frequency being 3 sessions per week. Each session lasted between 30-120 minutes, with a mean session duration of 60 minutes.

#### 4.5.3 Meta-regression results

Regression analysis was conducted on 8 performance indicators (Table 4; Supplementary Figure 61-68). The results revealed that the duration of training sessions had a partial influence on athletes’ power development (p<0.1), with a negative effect size (Z =−1.9). This implies that as the duration of each training session increased, the improvement in power performance was less pronounced.

**Table 4.** Meta-regression results, A multivariate random-effects meta-regression analysis was conducted to predict the impact of FT on athletes’ muscle strength.

| Covariate | Coefficient | 95%CI | Z | p |
| --- | --- | --- | --- | --- |
| **1RM squat/half squat (n=10)** |  |  |  |  |
| Intercept | 9.04 | -102.48 to 120.56 | 0.21 | 0.84 |
| Each session duration | 0.03 | -0.38 to 0.45 | 0.20 | 0.85 |
| Total weeks | -2.25 | -12.78 to 8.27 | -0.55 | 0.61 |
| frequency | -7.53 | -52.59 to 37.53 | -0.43 | 0.69 |
| Total sessions | 1.25 | -2.98 to 5.48 | 0.76 | 0.48 |
| **1RM bench press (n=11)** |  |  |  |  |
| Intercept | -18.79 | -78.76 to 41.18 | -0.77 | 0.47 |
| Each session duration | 0.003 | -0.14 to 0.15 | 0.05 | 0.96 |
| Total weeks | 1.68 | -3.46 to 6.83 | 0.80 | 0.45 |
| frequency | 5.91 | -13.85 to 25.67 | 0.73 | 0.49 |
| Total sessions | -0.46 | -2.22 to 1.30 | -0.64 | 0.55 |
| **1RM grip strength (n=10)** |  |  |  |  |
| Intercept | 0.63 | -15.77 to 17.03 | 0.09 | 0.93 |
| Each session duration | 0.02 | -0.09 to 0.12 | 0.37 | 0.73 |
| Total weeks | 0.55 | -2.56 to 3.65 | 0.43 | 0.68 |
| Total sessions | -0.20 | -0.99 to 0.59 | -0.63 | 0.55 |
| **Standing long jump (n=36)** |  |  |  |  |
| Intercept | 0.38 | -0.01 to 0.75 | 2.07 | **0.05** |
| Each session duration | -0.001 | -0.002 to 0 | -1.90 | **0.07** |
| Total weeks | -0.03 | -0.07 to 0.01 | -1.32 | 0.20 |
| frequency | -0.10 | -0.26 to 0.05 | -1.40 | 0.17 |
| Total sessions | 0.01 | -0.01 to 0.03 | 1.40 | 0.17 |
| **Countermovement jump (n=27)** |  |  |  |  |
| Intercept | 3.83 | 0.64 to 7.03 | 2.49 | **0.02** |
| Each session duration | -0.001 | -0.01 to 0.01 | -0.32 | 0.75 |
| Total weeks | -0.24 | -0.53 to 0.06 | -1.67 | 0.11 |
| frequency | -1.05 | -2.39 to 0.29 | -1.62 | 0.12 |
| Total sessions | 0.08 | -0.04 to 0.20 | 1.32 | 0.20 |
| **Throw medicine ball (n=12)** |  |  |  |  |
| Intercept | 9.26 | -0.19 to 18.71 | 2.32 | **0.05** |
| Each session duration | -0.004 | -0.02 to 0.01 | -0.78 | 0.46 |
| Total weeks | -0.71 | -1.56 to 0.14 | -1.97 | 0.09 |
| frequency | -2.69 | -5.88 to 0.50 | -1.99 | 0.09 |
| Total sessions | 0.23 | -0.06 to 0.51 | 1.88 | 0.10 |
| **1 min sit-ups (n=12)** |  |  |  |  |
| Intercept | 8.17 | -17.86 to 34.20 | 0.74 | 0.48 |
| Each session duration | 0.08 | -0.07 to 0.23 | 1.24 | 0.25 |
| Total weeks | -1.33 | -4.61 to 1.96 | -0.95 | 0.37 |
| frequency | -4.90 | -16.22 to 6.43 | -1.02 | 0.34 |
| Total sessions | 0.62 | -0.82 to 2.07 | 1.02 | 0.34 |
| **Push-ups (n=12)** |  |  |  |  |
| Intercept | 41.69 | -91.64 to 175.02 | 0.74 | 0.48 |
| Each session duration | -0.02 | -0.18 to 0.14 | -0.31 | 0.77 |
| Total weeks | -2.52 | -15.73 to 10.69 | -0.45 | 0.67 |
| frequency | -11.15 | -57.66 to 35.35 | -0.57 | 0.59 |
| Total sessions | 0.73 | -3.84 to 5.30 | 0.38 | 0.72 |

## 5 Discussion

Prior reviews highlight the efficacy of FT and TRT on muscle strength, physical functioning, and daily activities. FT improves movement skills, balance, agility, and neuromuscular efficiency, making it valuable for athletic training. Optimal FT involves at least 8 weeks of sessions over 60 minutes, 3 times per week. These findings support FT as a complement to TRT for enhancing muscle strength and athletic performance.

### Maximal Strength

Integrating FT into sports programs has gained attention for its potential to boost athletic performance. Evidence shows FT can lead to significant improvements, especially when well-structured. For example, a study with 31 elite male high school soccer players found that a 16-week FT program significantly enhanced lower limb strength and balance through varied exercises (e.g., squat variations, unilateral deadlifts, core exercises)[107]. Conversely, an 8-week FT intervention with table tennis players reported less pronounced effects, possibly due to fewer and less intense exercises per session[165].

This indicates that FT’s effectiveness depends on exercise diversity and volume. Compared to TRT, FT significantly enhances upper limb and grip maximum strength (ES = 1.02-2.68; p ≤ 0.001) and nearly significantly improves lower limb strength (ES = 3.12, p = 0.07). A study with young male soccer players found FT significantly influences peak force, with TRT showing slower effects in training adaptation[134]. FT addresses issues like young athletes’ common lack of trunk balance and strength by incorporating unstable environments and multi-joint exercises, enhancing core muscle engagement and overall strength training efficacy[53]. Another study with skilled basketball athletes showed that the FT group had superior improvements in maximal strength, musculoskeletal health, and basketball performance (WMD = 4.5)[63]. Despite positive outcomes, FT’s role in exercise science is debated due to its lower muscle load compared to TRT[63,125].Critics argue FT may not enhance muscle strength as effectively as TRT[11,105]. For instance, a study with 12 professional baseball athletes found grip strength increased 85% less after 12 weeks of FT compared to TRT(WMD = −3.82)[148]. However, many studies confirm FT can effectively increase muscle strength, comparable to TRT[11,28,35,52]. For example, in older women, FT significantly improved upper limb strength and balance/agility compared to stretching exercises, though no significant changes in body composition were observed[121]. While TRT is more effective for large muscle groups, FT stimulates both large and small muscle groups, enhancing overall strength, functional strength, and neuromuscular proficiency[64,108]. The combined approach of FT followed by TRT (FT->TRT) or vice versa (TRT->FT) demonstrated significant improvements in balance, agility, and upper limb strength, suggesting FT may provide faster and greater magnitude adaptations[66]. Our analysis confirms FT significantly enhances muscle strength, especially in older adults[31,68]. For optimal results, longer durations (>60 minutes), higher frequencies (≥3 times/week), increased practice variations (≥5/session), and extended total weeks and sessions are recommended. This aligns with the muscle development overload principle, suggesting more intense stimulation leads to faster muscle growth[114].

### Power

FT significantly enhances power in athletes, with effect sizes ranging from small to large (ES = 0.07-0.68) and consistent statistical significance (p < 0.001) compared to control groups. For instance, a 12-week FT program for 57 amateur soccer players significantly increased power[92]. FT improves muscle strength by enhancing neuromuscular coordination and efficiency, aligning with research on high-velocity power muscle contractions for high power output [14,15,81,99]. In contrast, TRT, focusing on slow and high-repetition muscle hypertrophy, has not shown the same ability to improve power[16]. FT often includes bodyweight exercises and rapid eccentric-concentric transitions, leveraging the stretch-shortening cycle (SSC) to enhance force production and power performance[12,18,45,62,70,72,116,128]. Studies have shown significant improvements in countermovement jump (CMJ) performance after FT, highlighting the benefits of integrating high-speed and explosive movements in training. However, some studies note less progress in specific metrics like the standing long jump (SLJ) when FT is not combined with heavy resistance training[63,148]. FT’s advantage over TRT lies in coordinating the entire kinetic chain, improving neural activation, and precise muscle control, enhancing movement efficiency[19]. While FT significantly improves power, training programs should be carefully tailored to avoid overtraining and maximize benefits[39,44,61,81]. Additionally, research has demonstrated that FT can enhance sprinting, jumping, and functional movement skills, particularly in young athletes engaged in team sports. The systematic review found that FT improved athletes’ performance in 5, 10, 15, 20, 25, and 30-meter sprints, as well as in functional movement skills tests[89,117,129].

### Muscle Endurance

FT effectively enhances muscle endurance, showing moderate to large effect sizes. FT significantly increased 1-minute sit-ups (ES=4.13), PSU (ES=3.3), with near-significant gains in pull-ups (p=0.07) compared to TRT[30]. An 8-week FT program with dynamic and static core exercises (e.g., seated cable pulldowns, leg raises) significantly improved 1-minute sit-ups versus a control group. Despite smaller gains in pull-ups compared to TRT, FT’s focus on multi-joint movements enhances overall muscle endurance (WMD=-2.75)[66,164]. Another randomized controlled trial (RCT) found significant improvements in baseball performance-related physical factors in both FT and TRT groups, suggesting complementary effects[136,138]. FT’s emphasis on core stability through multi-plane and unstable environment exercises enhances proprioception, optimizes movement patterns, and strengthens neuromuscular connections, crucial for endurance[82]. While FT includes diverse core exercises, TRT employs fewer[74]. For older adults, FT has shown to improve muscle endurance, balance, and overall physical fitness, making it a suitable approach to combat age-related decline in physical abilities[45,70,121]. In conclusion, FT effectively enhances muscle endurance through core stability, proprioception, and neuromuscular connections. For rapid muscle endurance improvements, particularly in athletes with high initial strength, TRT may provide more immediate results.

### Various Sports

The sports analyzed in this paper fall into four main categories: ball sports, water sports, combat sports, and artistic sports.

### Ball Sports

The enhancements in power for tennis, soccer, and basketball players following FT are more significant than those for baseball, badminton, and table tennis athletes. This is due to the higher volume of running and jumping in the former sports. Soccer players perform 150-250 high-intensity actions per match, basketball players cover several kilometers with numerous high-speed maneuvers and about 50 power jumps per quarter, and tennis involves repetitive, brief sprints[77,78,96]. In contrast, baseball, badminton, and table tennis require more agility and reaction speed. FT compensates for these differences by boosting power, improving overall performance.[63,85].

### Water Sports

FT significantly enhances physical attributes and performance in water-based sports like swimming, dragon boating, and windsurfing. FT improves swimmers’ shoulder flexibility, upper body coordination, and overall aquatic movement, reducing injury risk[160]. In dragon boating, FT boosts functional movement, core strength, muscle endurance, and rowing speed, aiding consistent and powerful race rhythms[130]. Windsurfers benefit from FT’s emphasis on muscle strength, crucial for sail control and balance. These improvements have been attributed to FT’s focus on core stability, multi-joint movements, and neuromuscular coordination[70,176].

### Comat Sports

Research on combat sports like wrestling, taekwondo, boxing, and sanda shows FT enhances agility, lower limb power, and core strength, while TRT quickly builds muscle strength and size[155,157]. Combining FT and TRT is recommended for optimal results, as TRT is essential for stimulating large muscle groups required in combat sports[156].FT’s ability to improve agility and core strength complements TRT’s muscle-building effects, providing a balanced approach to training in combat sports[63].

### Artistic Sports

FT benefits male gymnasts by improving landing stability, movement functionality, balance control, core strength, muscle endurance, and stability[123].These enhancements translate to better landing movements and may improve power and jumping performance over time[146,158]. FT is an important training modality for artistic sports, though conclusions should be cautiously evaluated due to limited research[26,75]. In artistic sports, FT’s emphasis on balance and core stability can enhance performance by improving movement efficiency and reducing injury risk[70].

This meta-analysis consolidates the understanding that while both FT and TRT have unique advantages, their combination may offer the most comprehensive benefits for athletes across various sports. FT’s focus on functional movement, core stability, and neuromuscular coordination complements TRT’s emphasis on muscle hypertrophy and strength, leading to well-rounded athletic development. The combined approach of FT followed by TRT (FT->TRT) or vice versa (TRT->FT) demonstrated significant improvements in various performance metrics. Future research should continue to explore the optimal integration of FT and TRT to maximize athletic performance across different sports disciplines.

### Limitations and Perspectives

This meta-analysis faces certain limitations, such as incomplete athlete data and unclear reporting of training specifics, which could influence the analysis. Further investigation is needed to understand how age affects the efficacy of FT. Diverse training criteria and inconsistent reporting may lead to biases, undermining the meta-analysis’s reliability. Future studies should aim to standardize assessment techniques to reduce potential errors.

### Practical Applications

FT may improve an athlete’s strength, and it seems that younger athletes might benefit more from it. For coaches, it is recommended that when designing FT sessions to boost power, these should be kept relatively brief. Conversely, sessions that aim to develop maximal strength and endurance can be longer. It is crucial to incorporate a variety of training methods that are both diverse and targeted, while also ensuring that the training load is sufficient. Moreover, more frequent training sessions and a longer overall training period can lead to improved outcomes.

## Acknowledgments

Conceptualization and design: Junyan Liu and Lei Shang Literature search: Junyan Liu and Lei Shang. Analysis: Junyan Liu and Lei Shang. Manuscript writing, first draft: Junyan Liu Manuscript writing, revision: Junyan Liu. Supervision: Shang Lei and Hongjun Yu. All authors read and approved the final manuscript. Availability of data: Full data coded of the included studies can be shared upon reasonable request from the corresponding author. The authors have no conflicts of interest to disclose. The results of this study do not constitute endorsement of the product by the authors or the National Strength and Conditioning Association.

## Supplementary Figure

**Figure 1.**
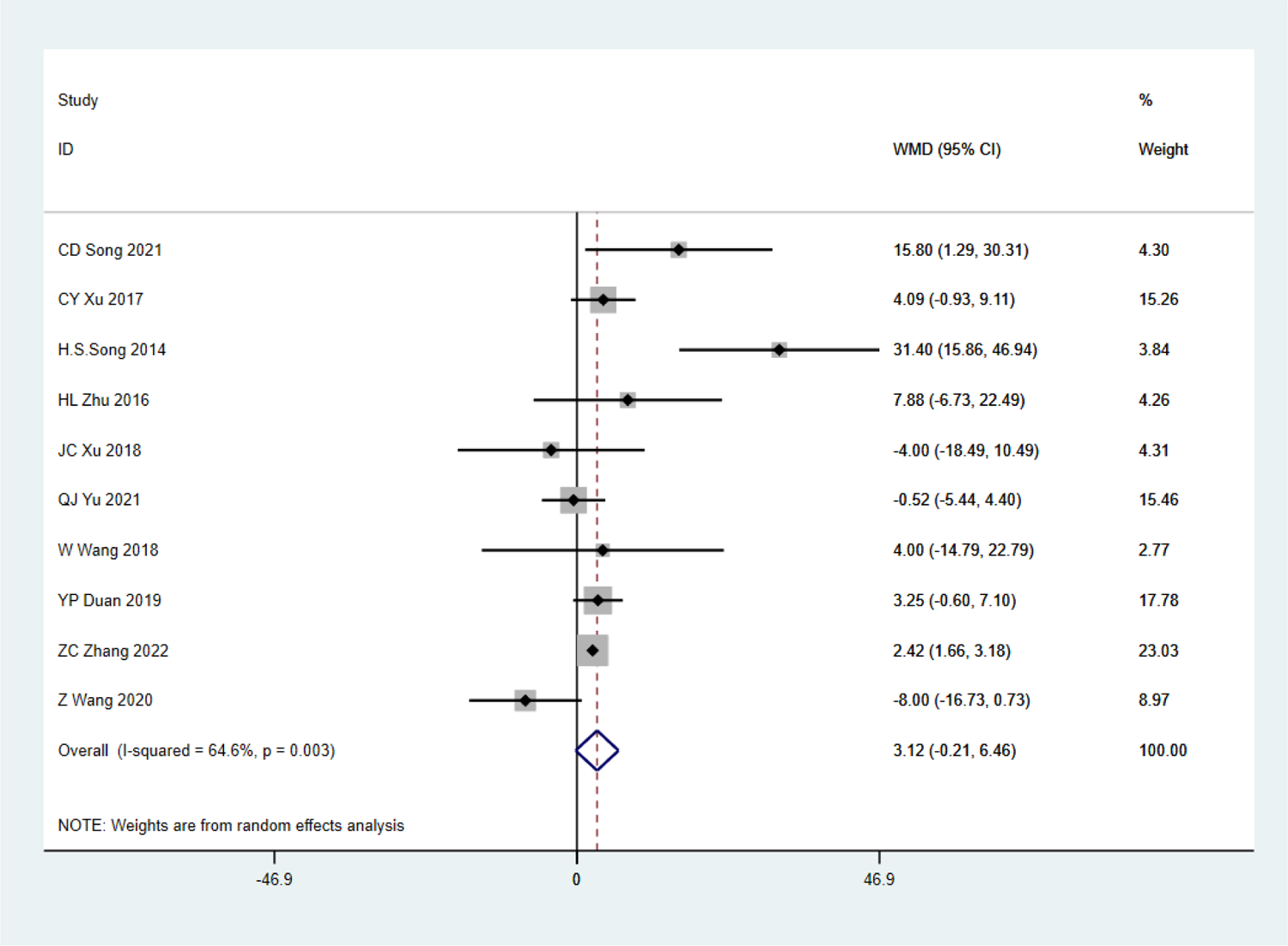
Forest plot - 1RM H/DS

**Figure 2.**
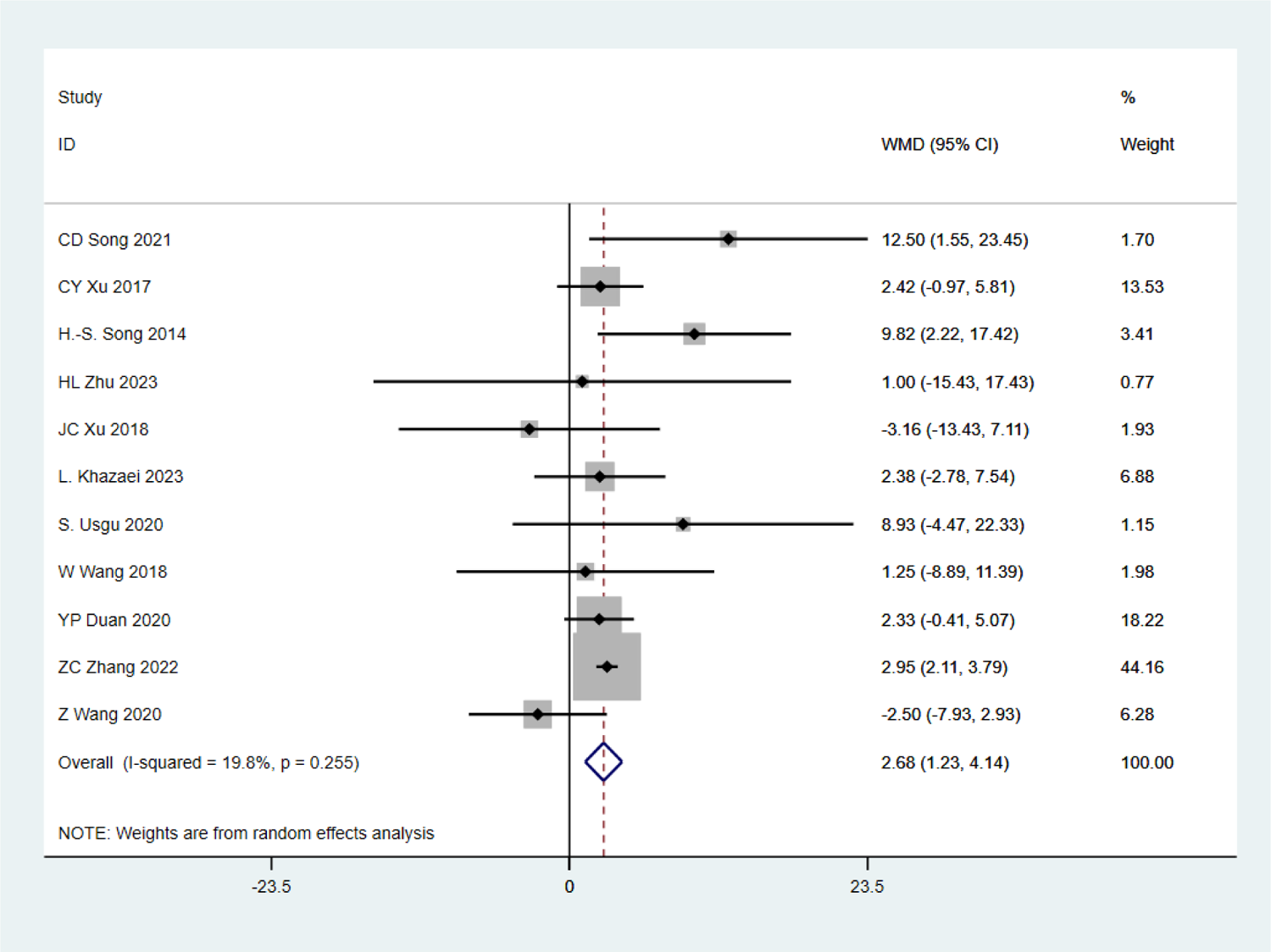
Forest plot - 1RM BP

**Figure 3.**
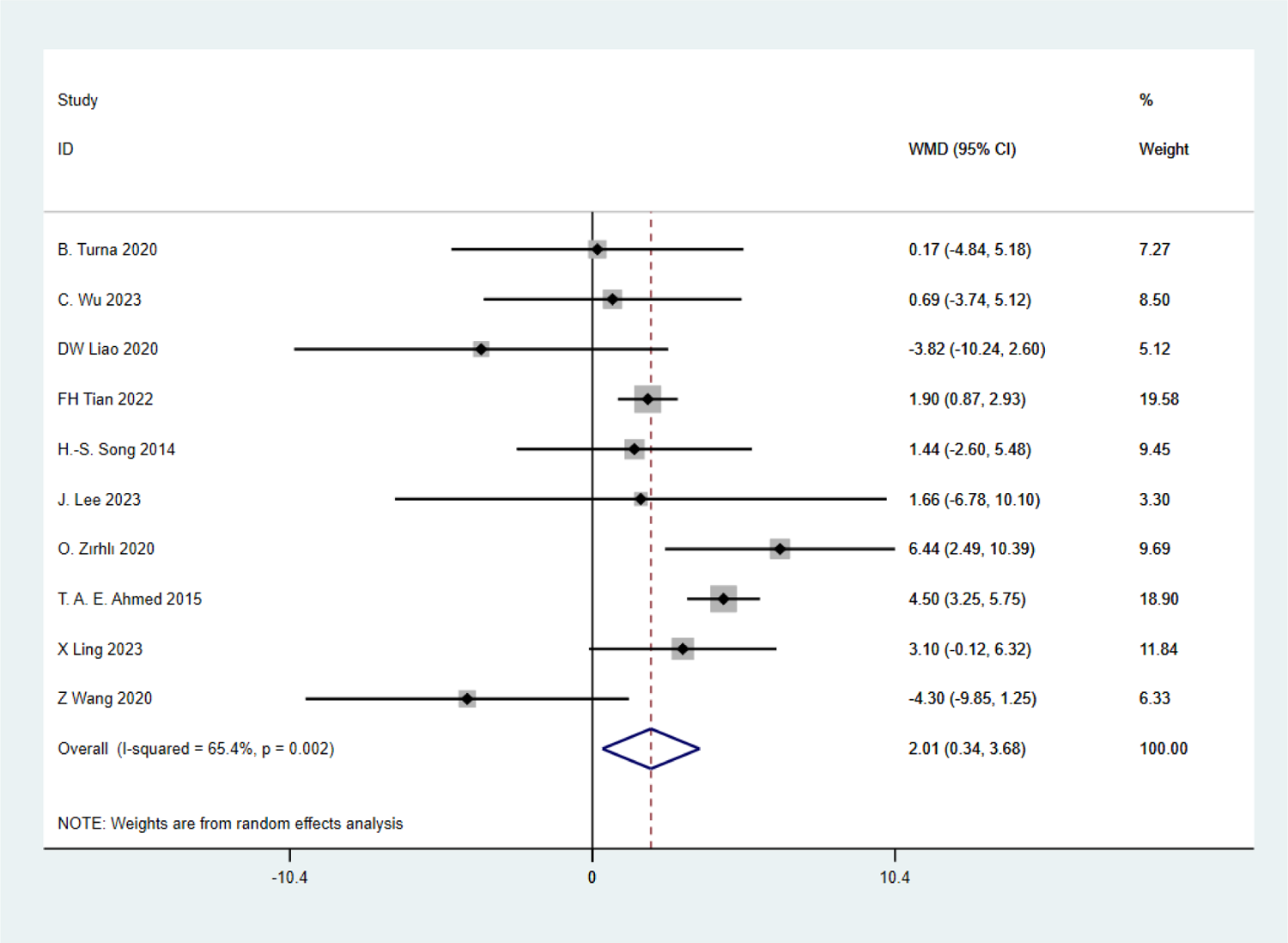
Forest plot 1RM GS.

**Figure 4.**
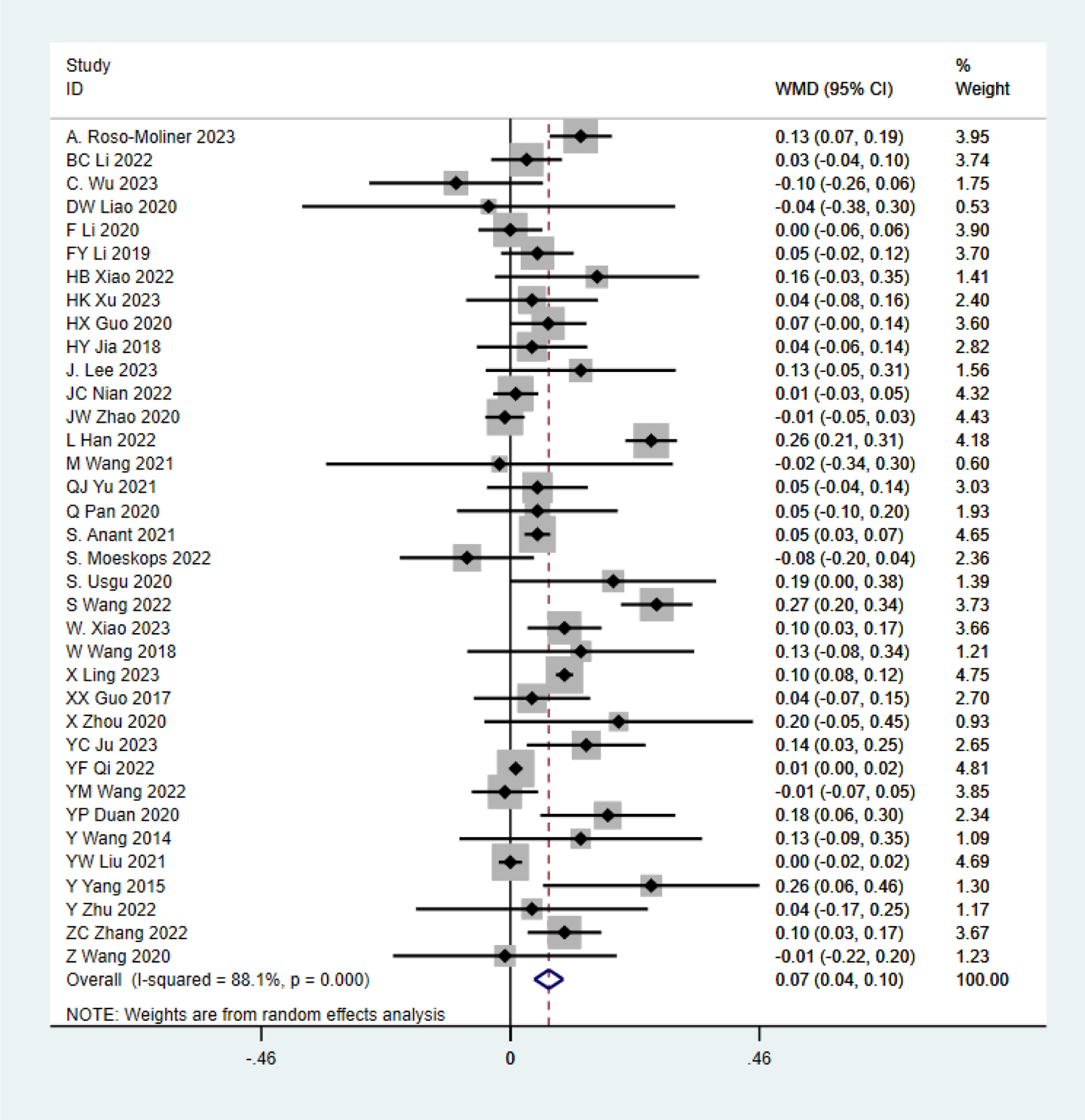
Forest plot - SLJ.

**Figure 5.**
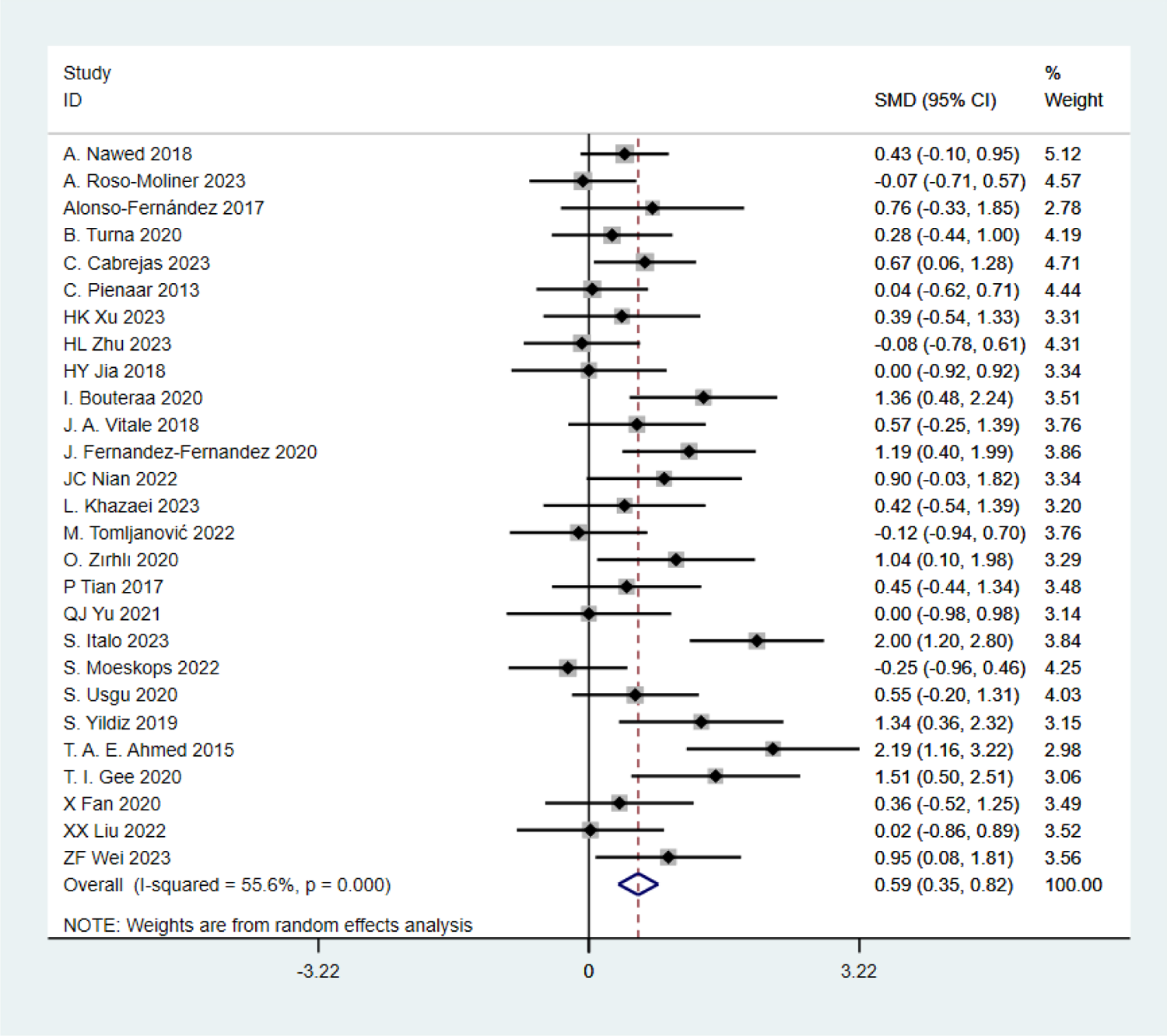
Forest plot - CMJ.

**Figure 6.**
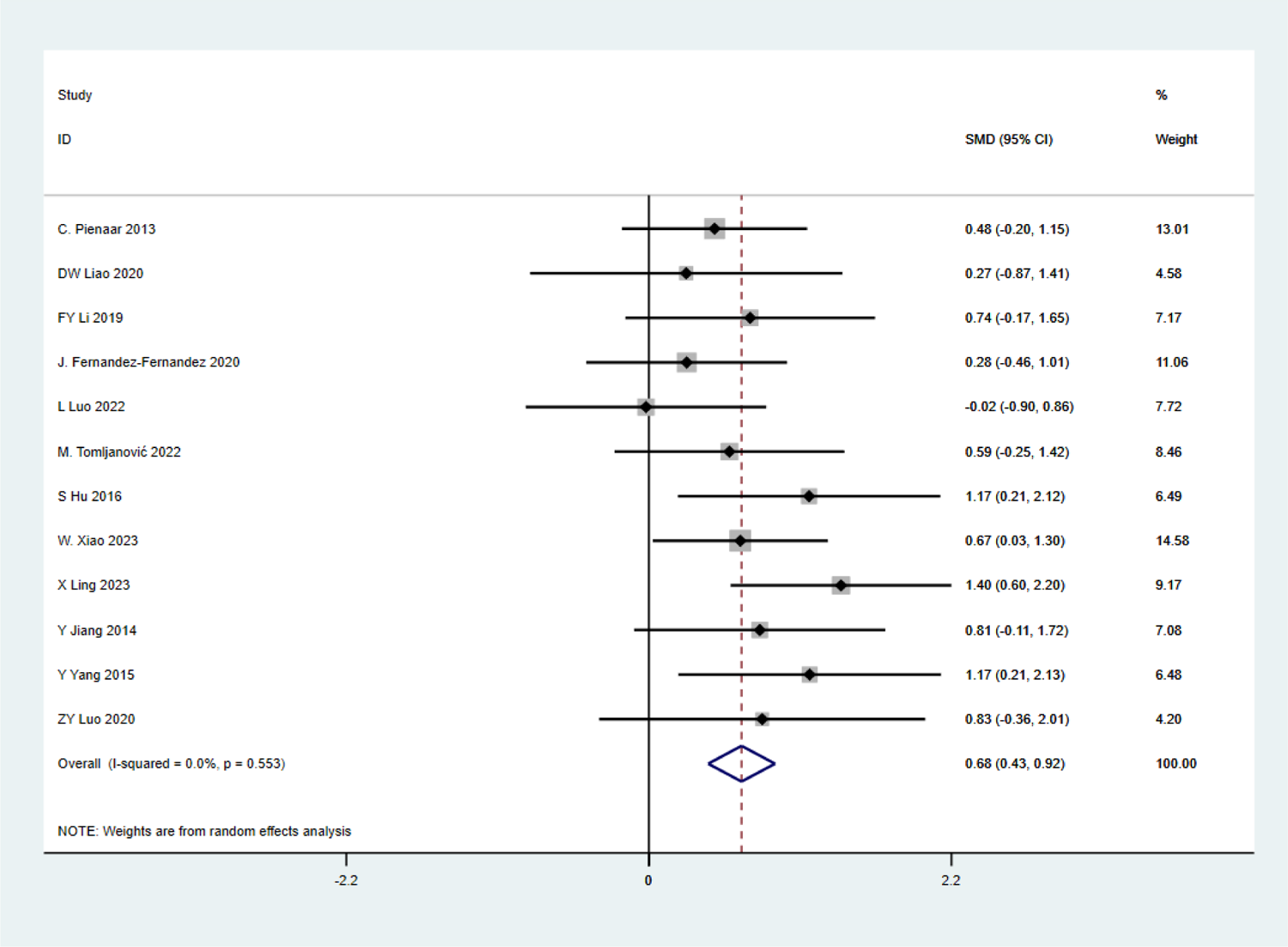
Forest plot - TMBP.

**Figure 7.**
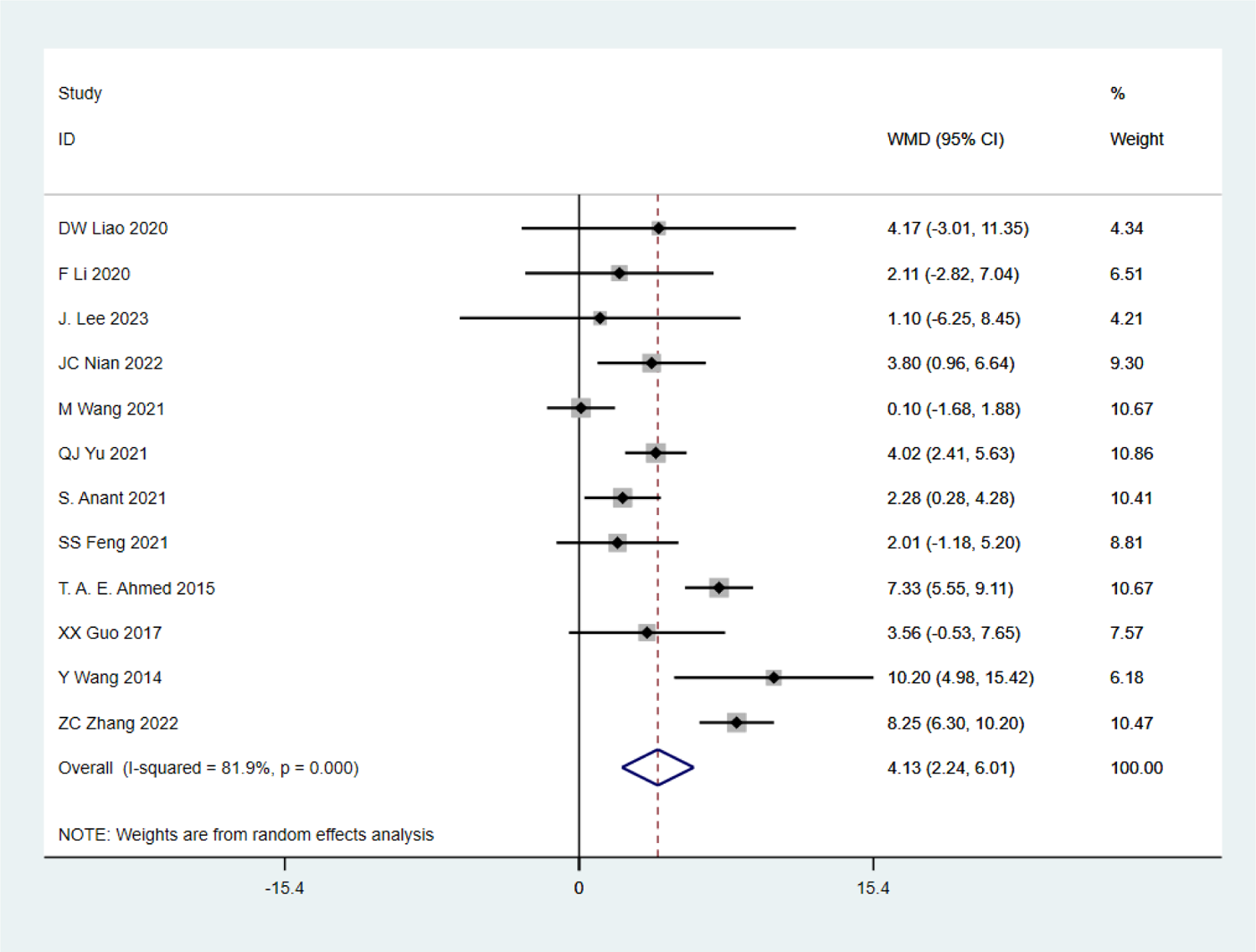
Forest plot - 1min SU.

**Figure 8.**
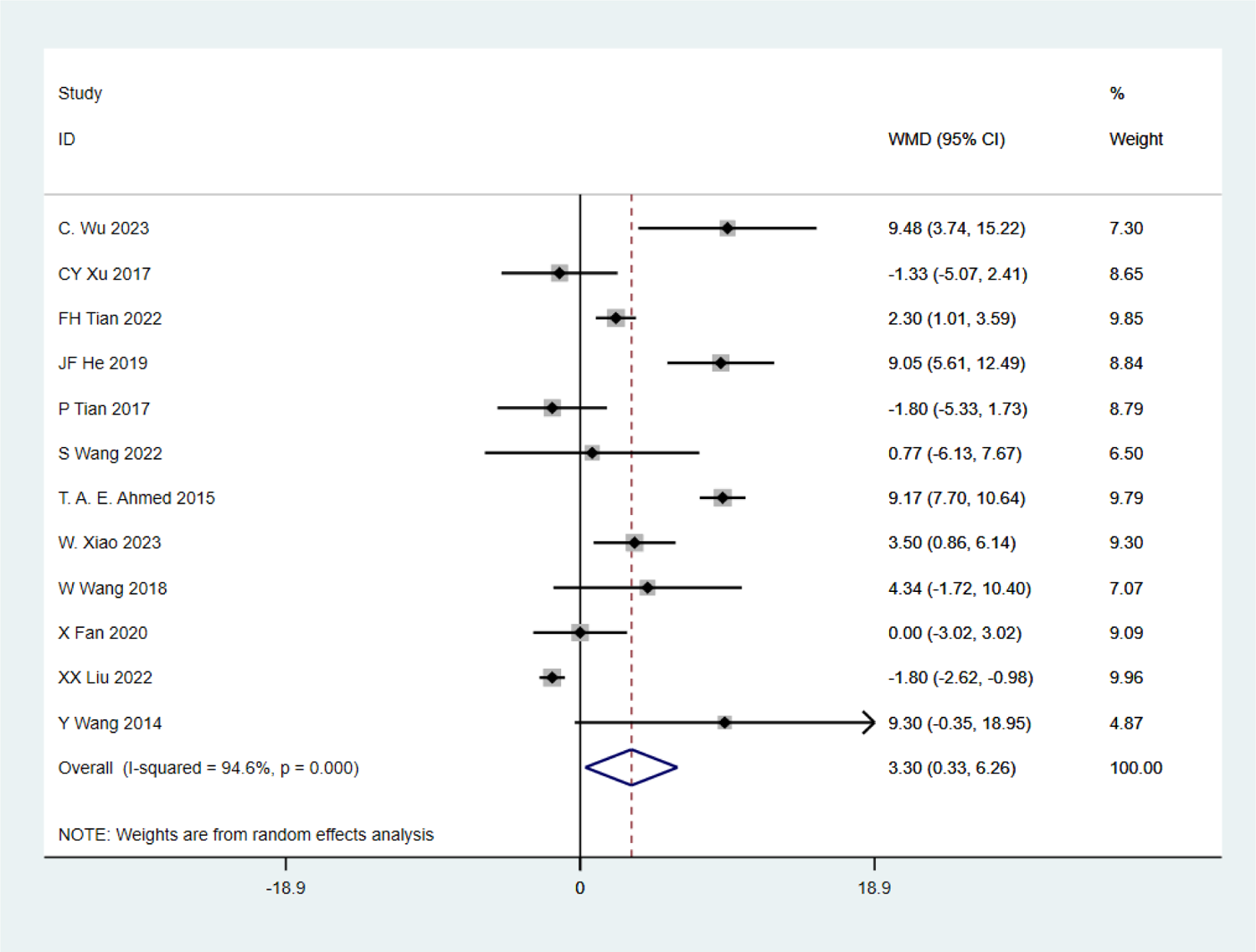
Forest plot - PSU.

**Figure 9.**
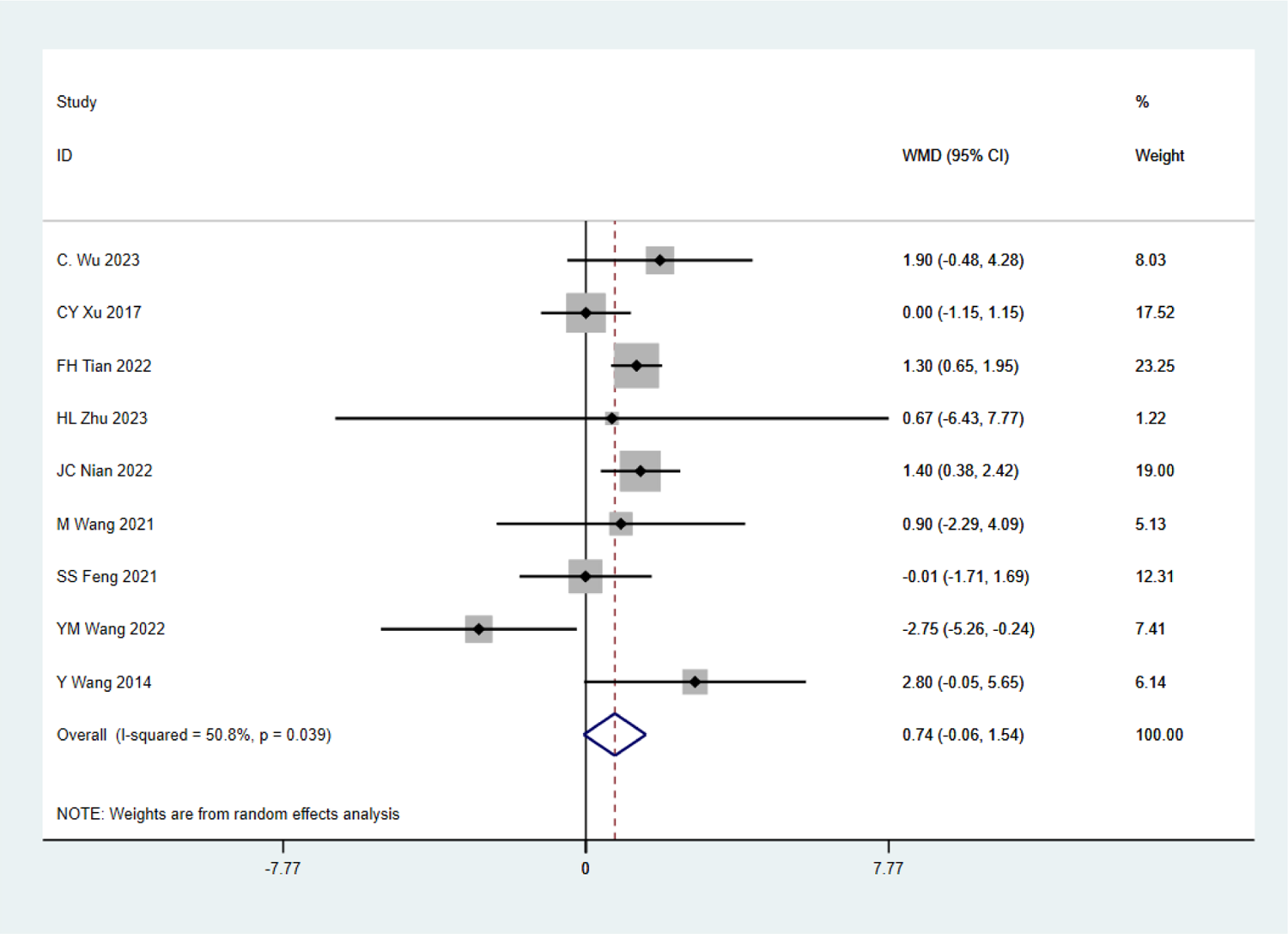
Forest plot - PLU.

**Figure 10:**
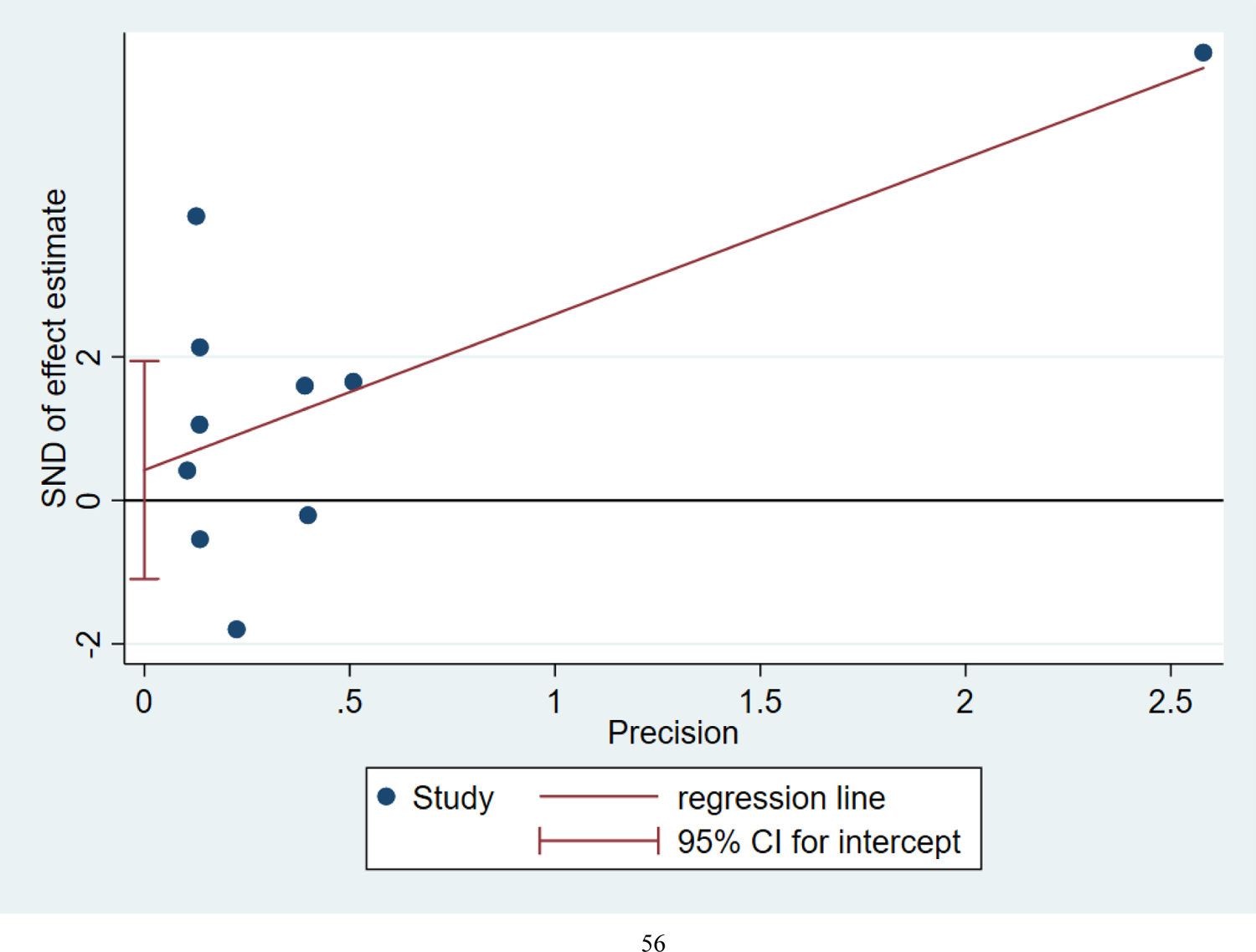
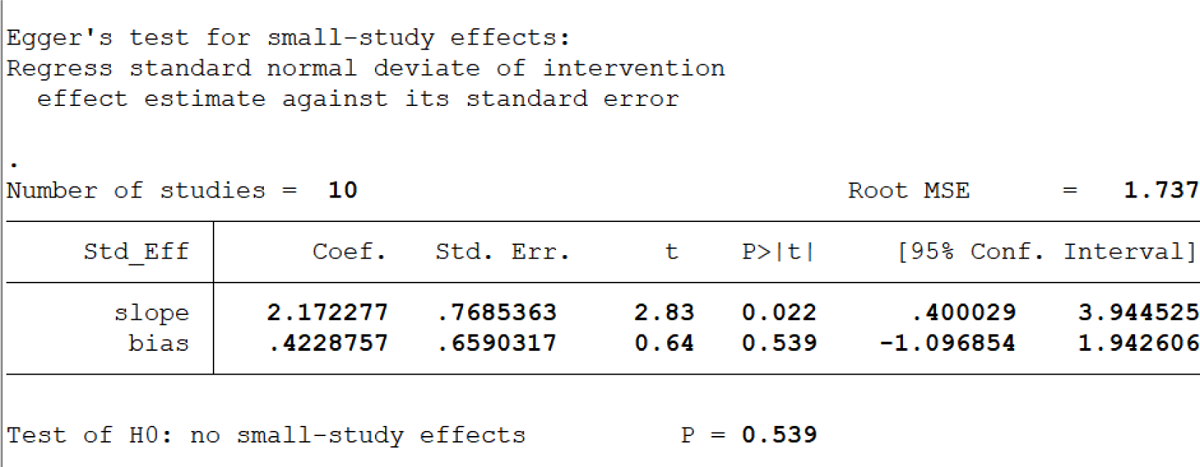
Egger’s test - 1RM H/DS

**Figure 11:**
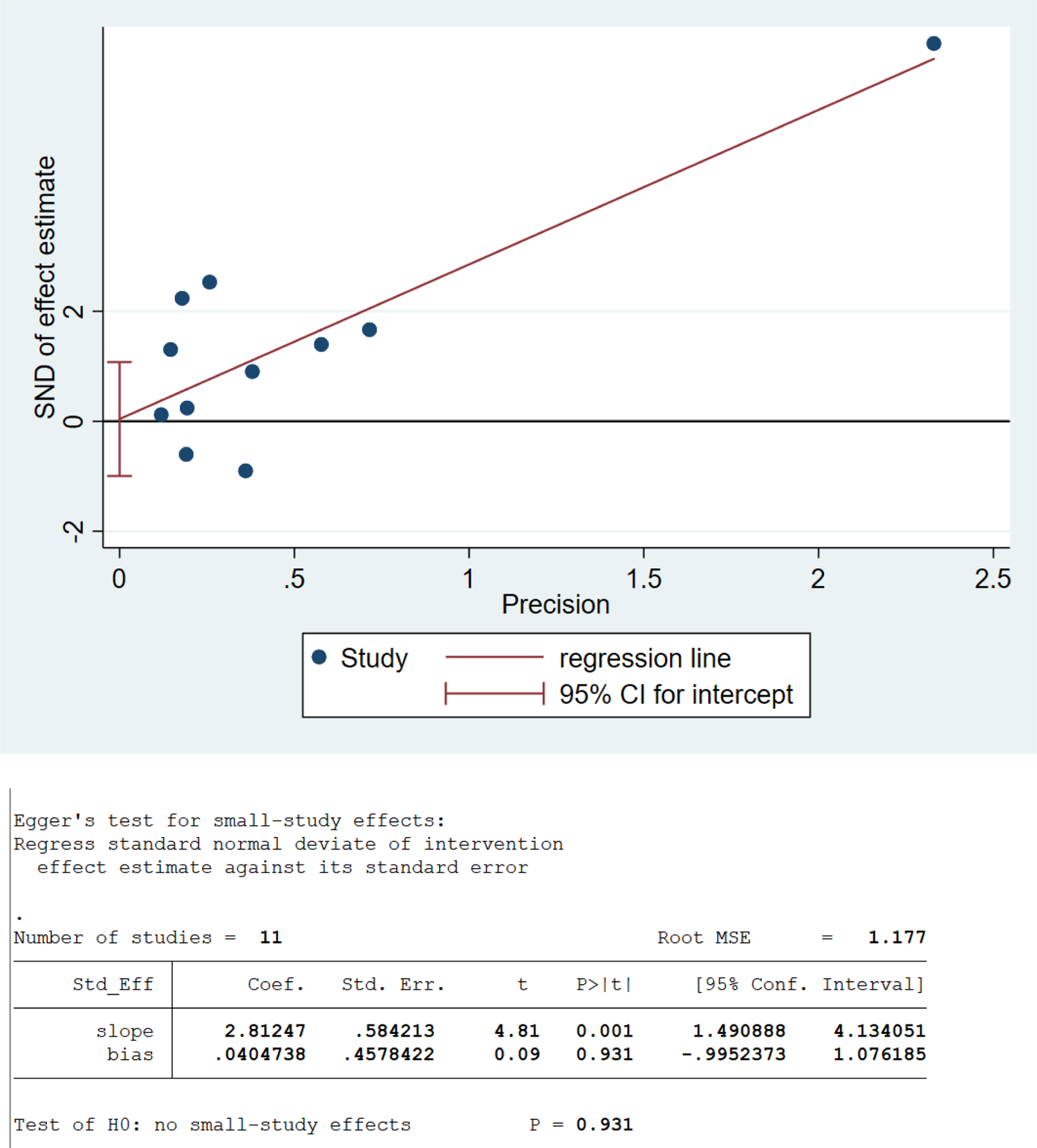
Egger’s test - 1RM BP

**Figure 12:**
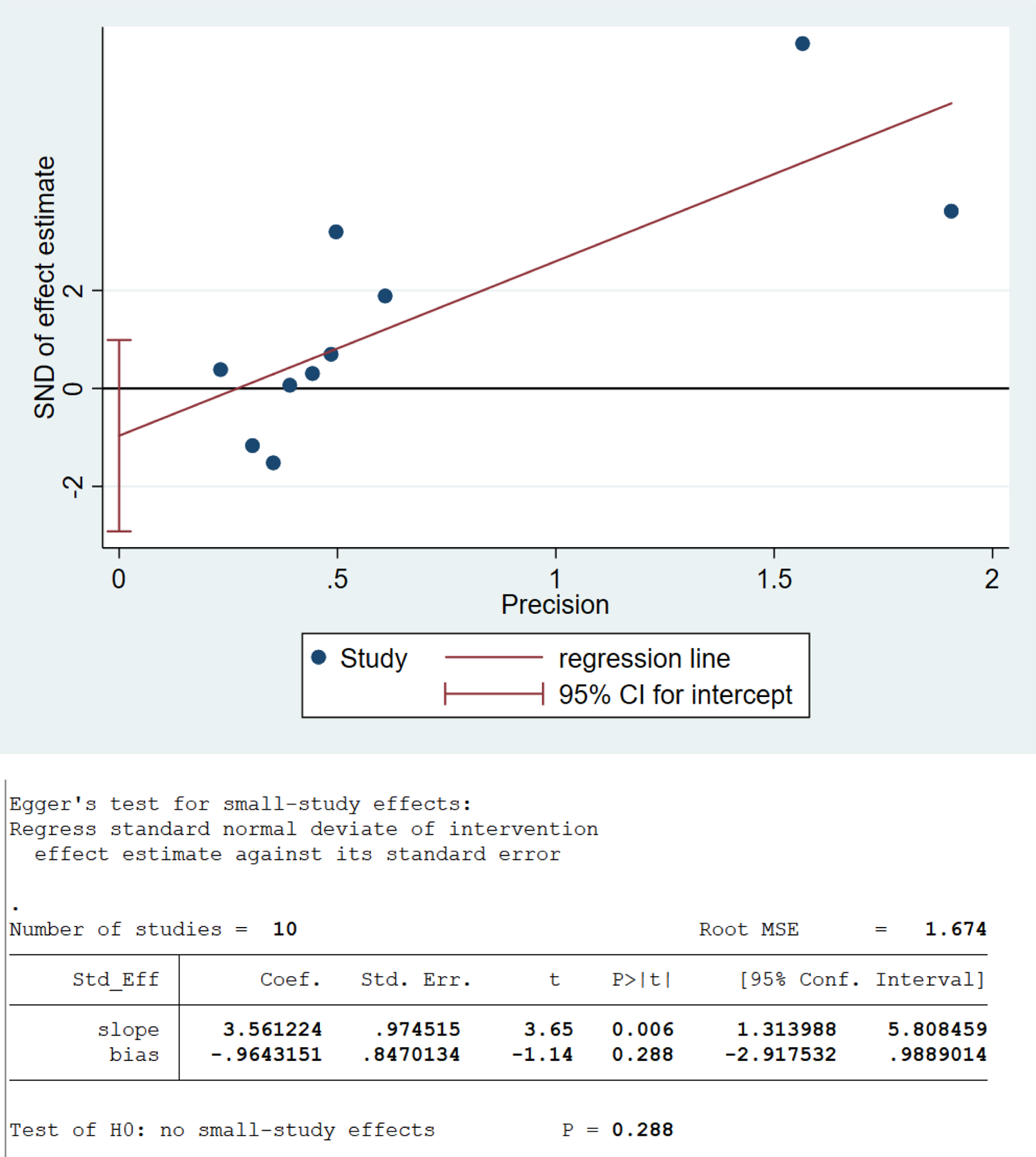
Egger’s test - 1RM GS

**Figure 13:**
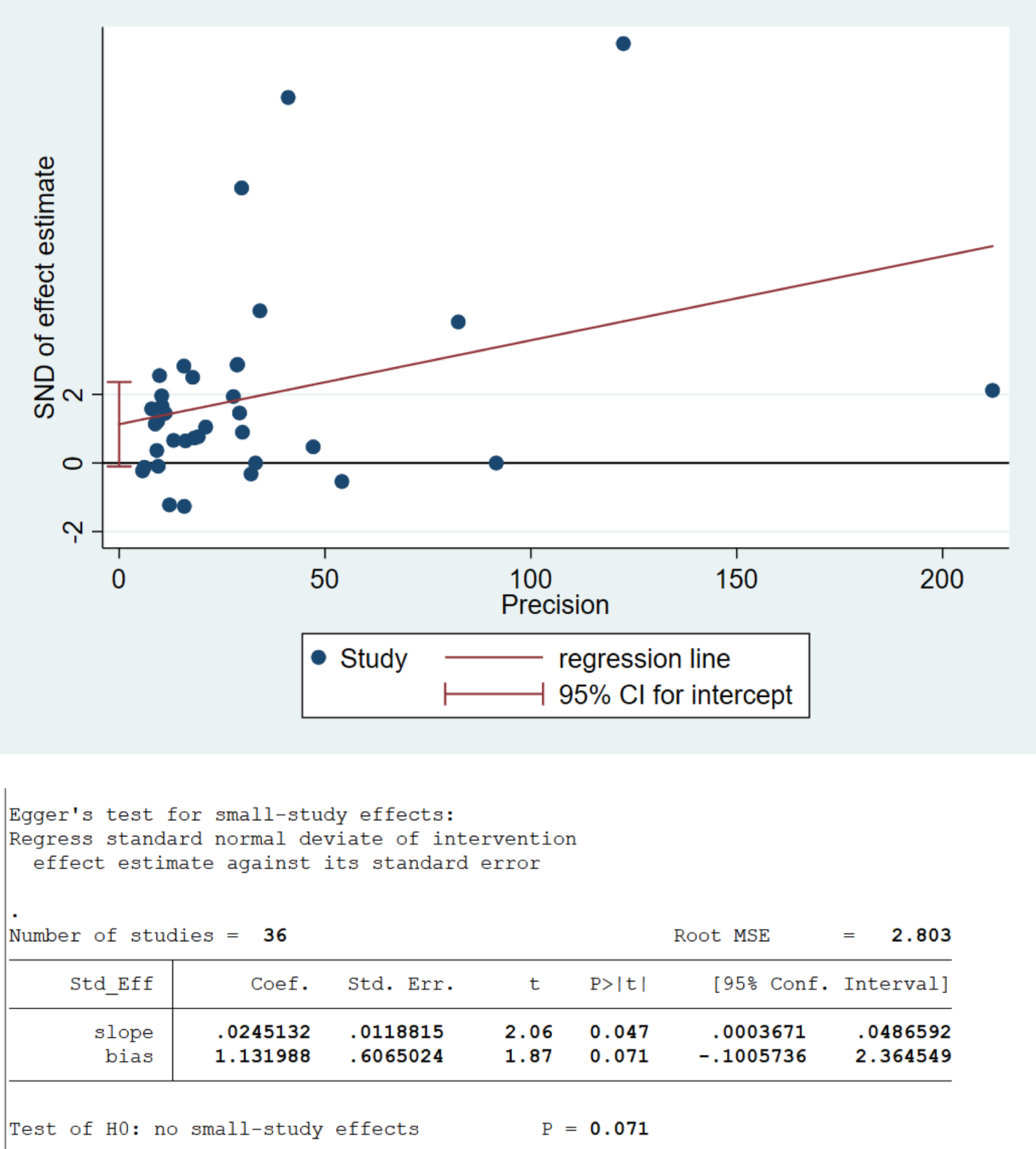
Egger’s test - SLJ

**Figure 14:**
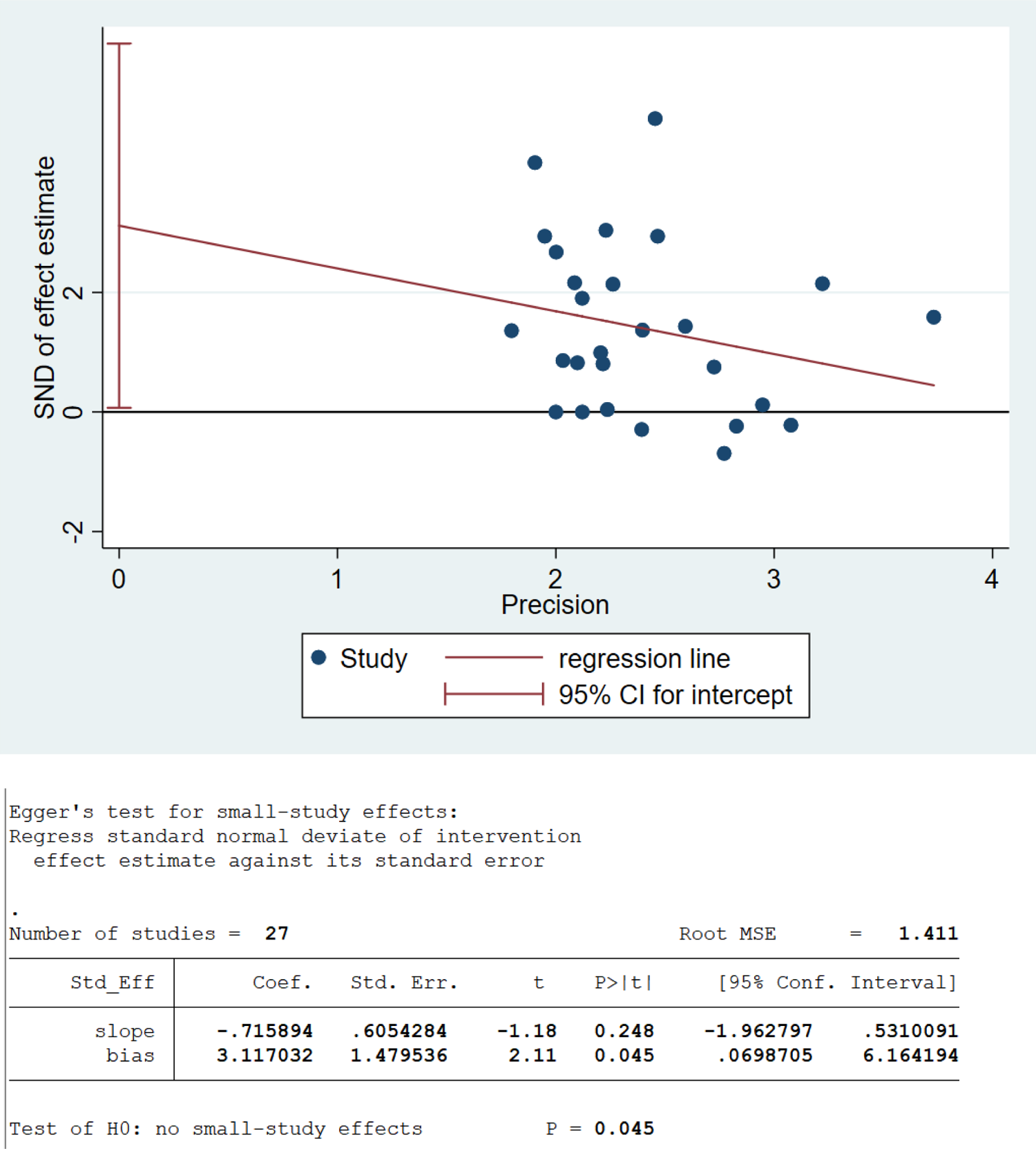
Egger’s test - CMJ

**Figure 15:**
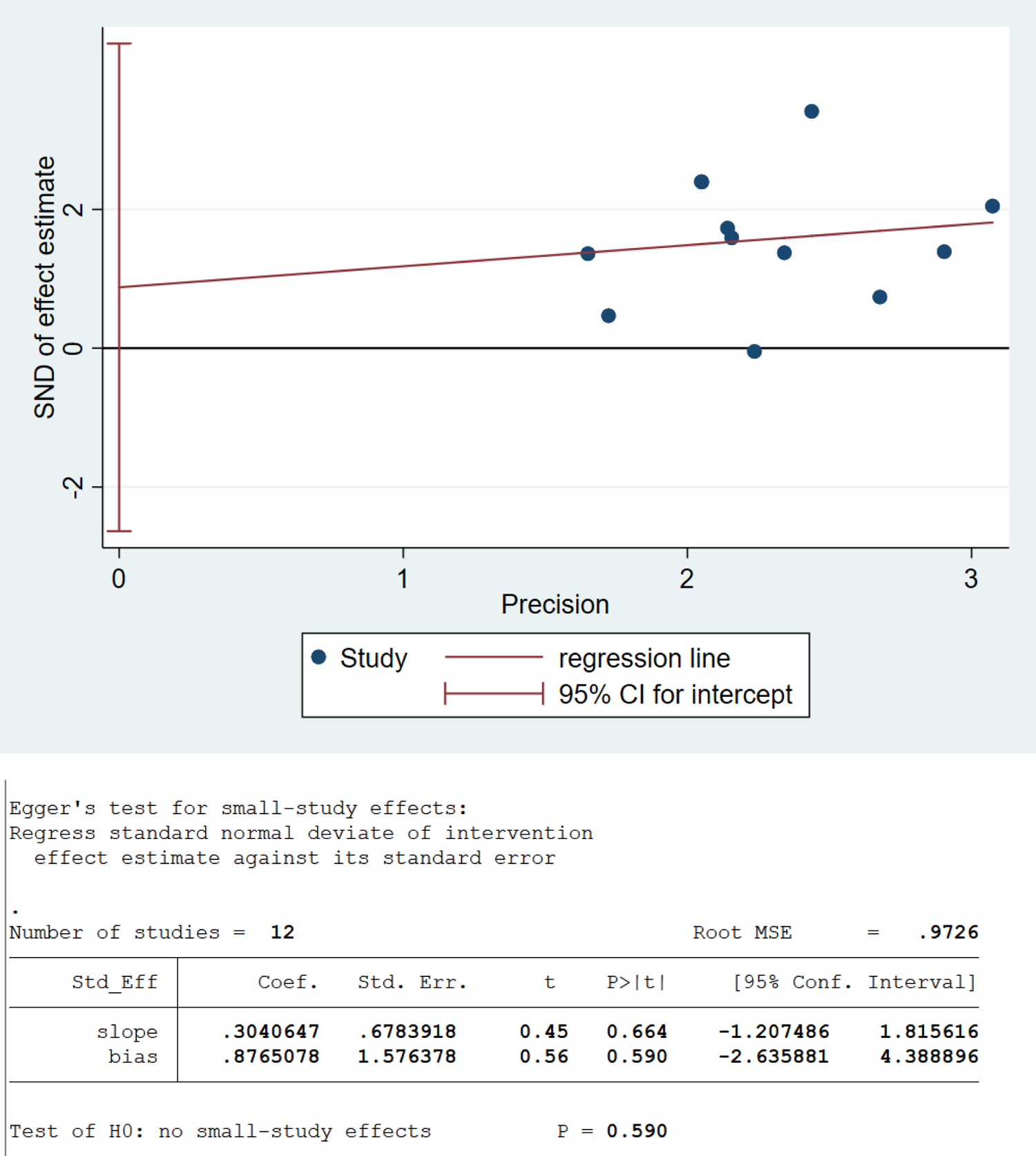
Egger’s test - TMBP

**Figure 16:**
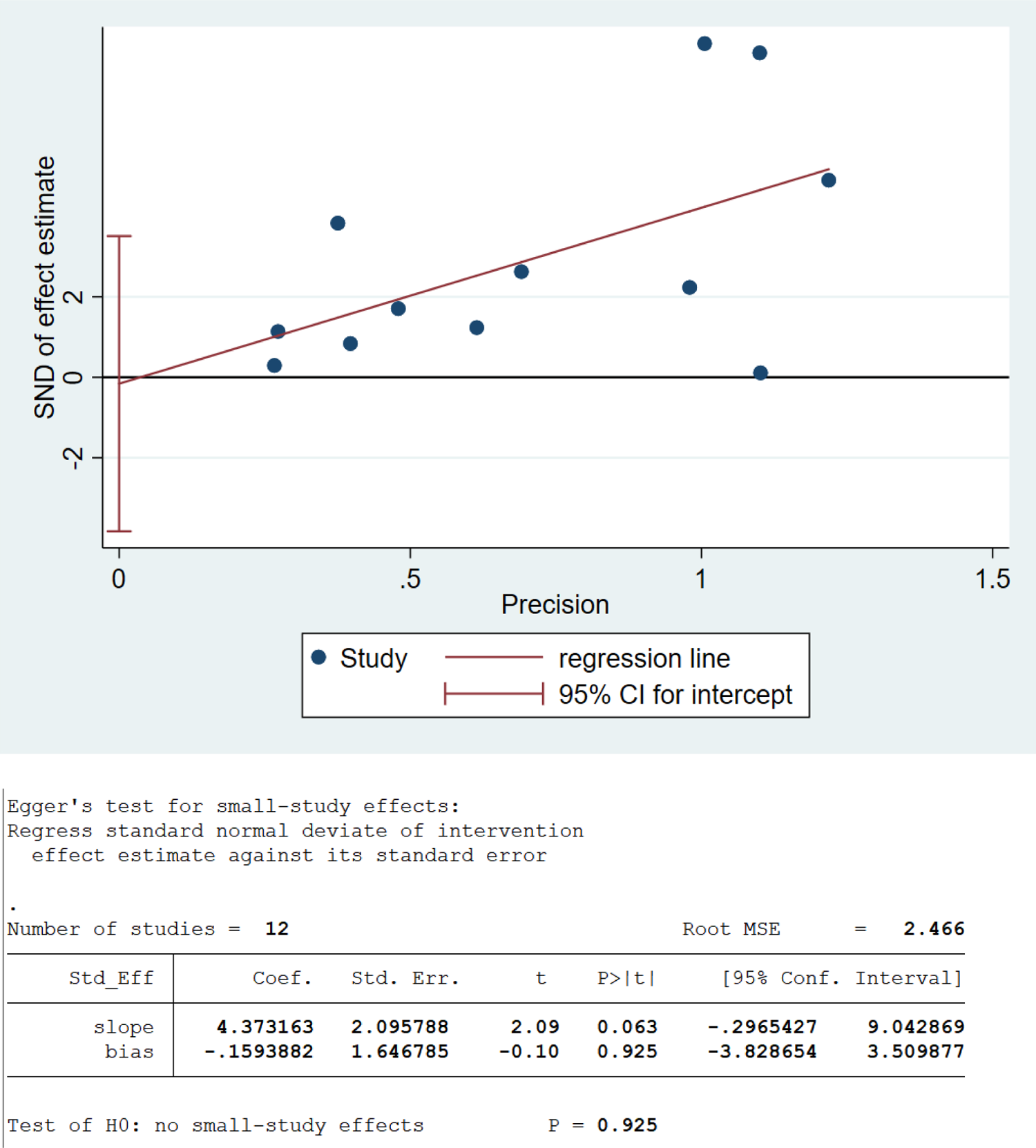
Egger’s test - 1min SU

**Figure 17:**
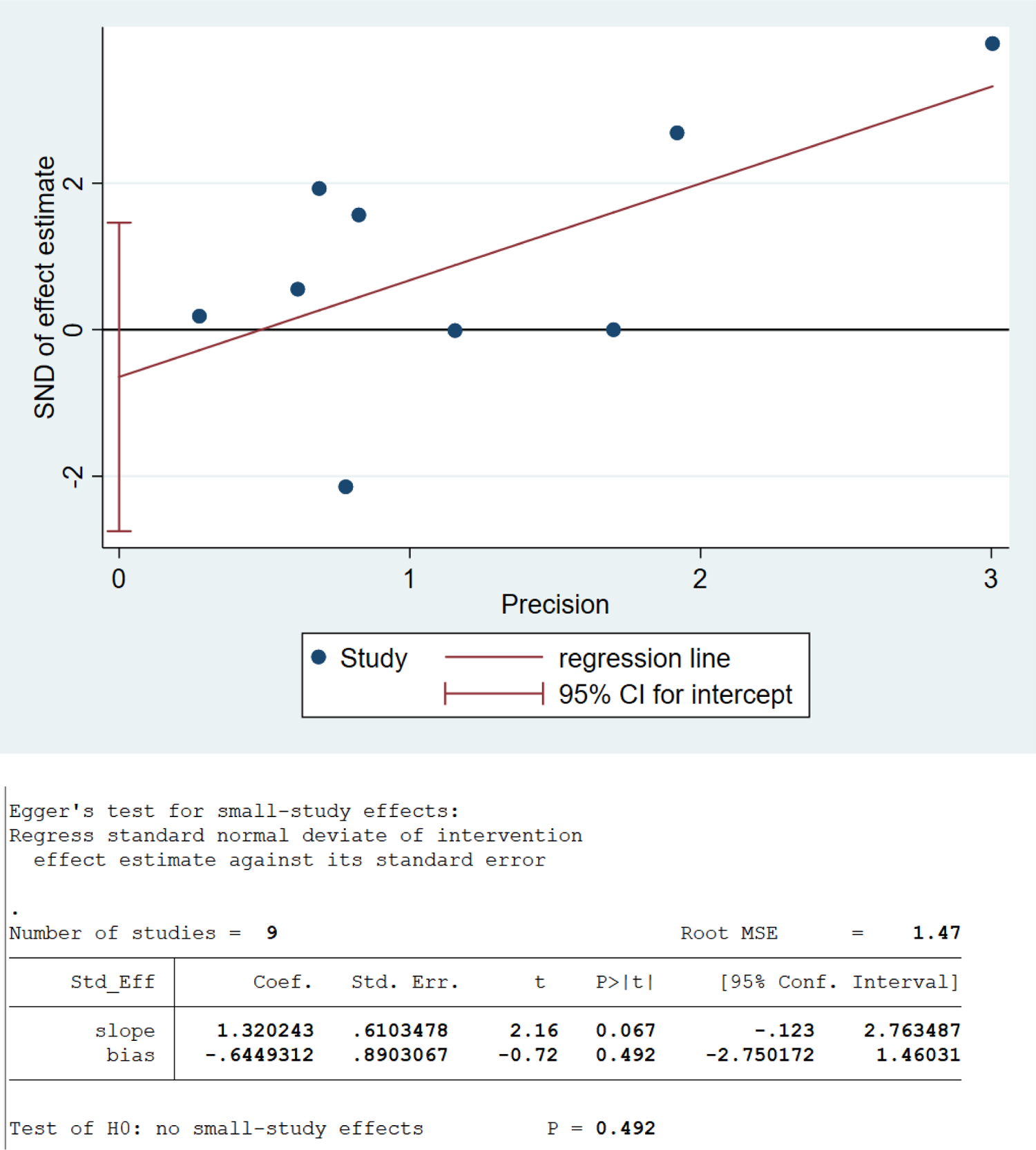
Egger’s test - PLU

**Figure 18:**
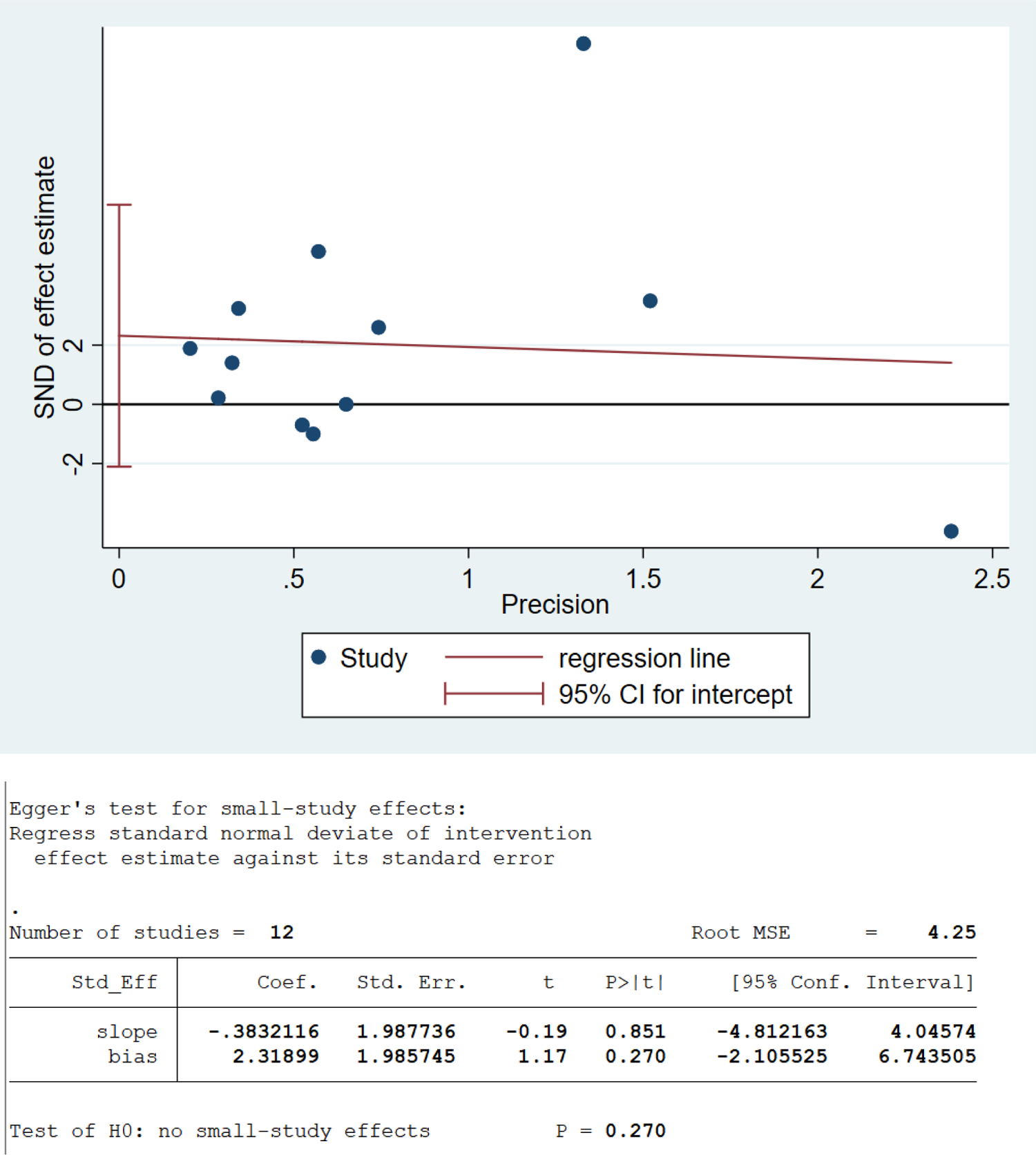
Egger’s test - PSU

**Figure 19:**
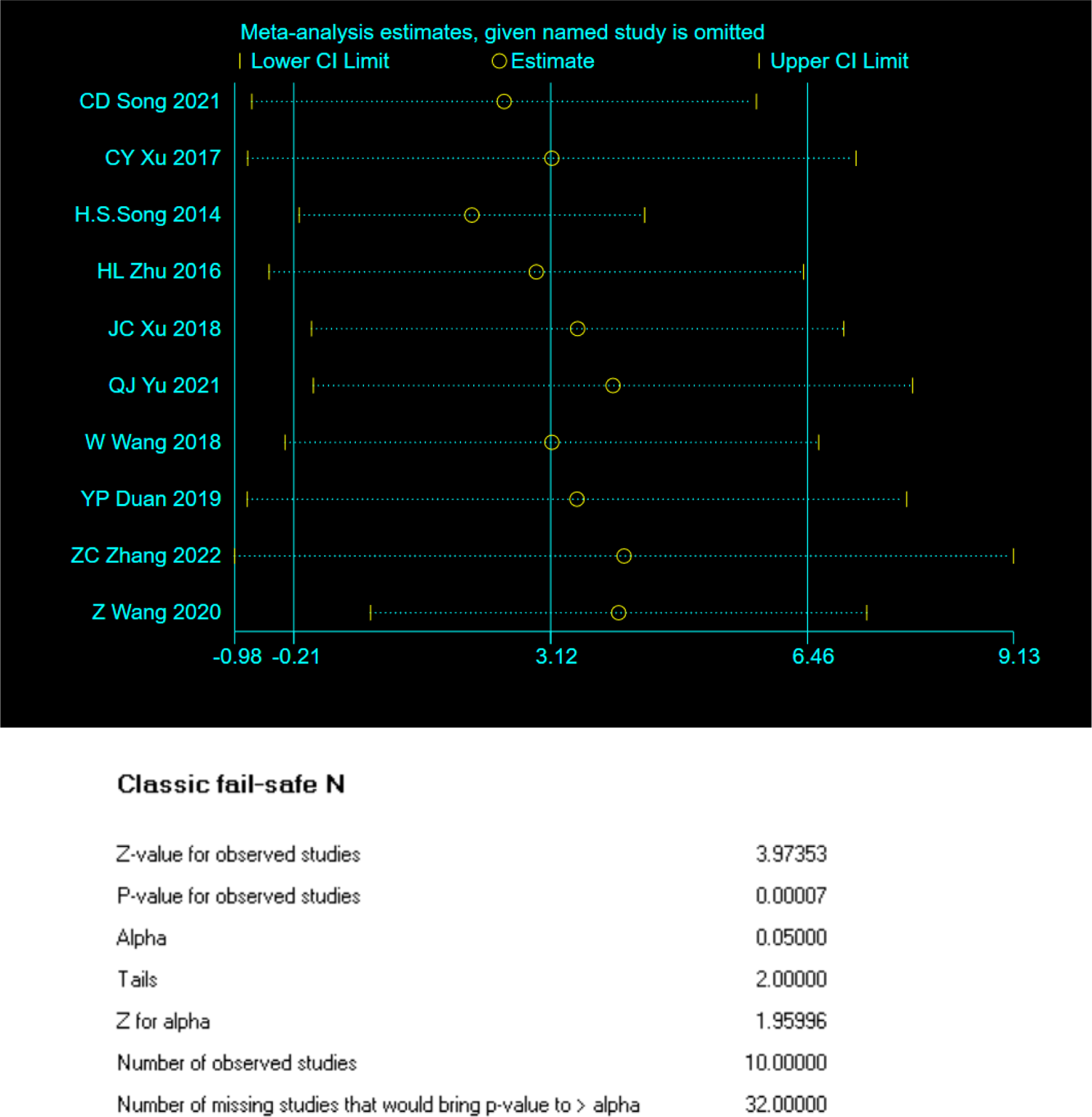
Sensitivity analysis - 1RM H/DS.

**Figure 20:**
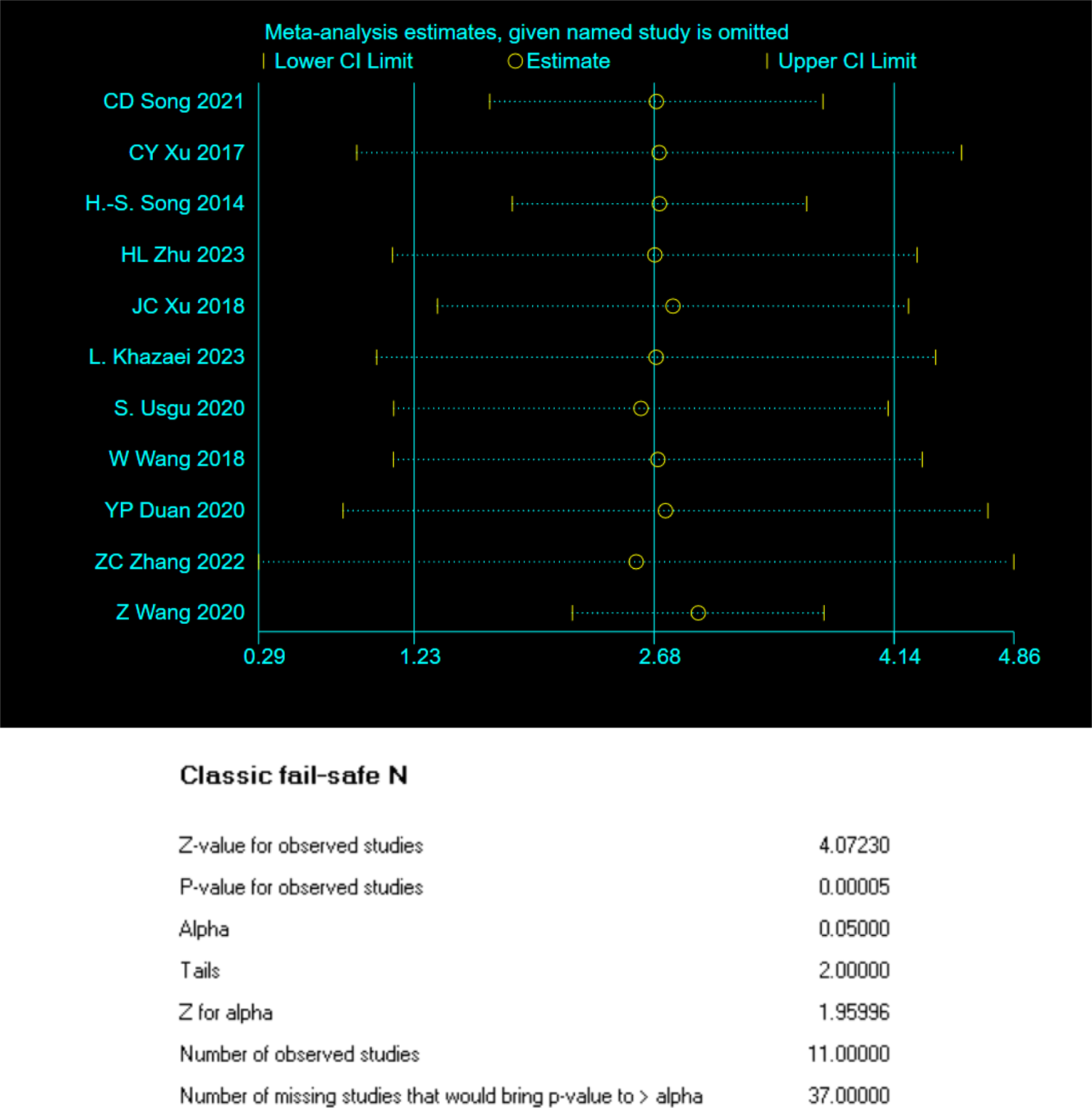
Sensitivity analysis - 1RM BP.

**Figure 21:**
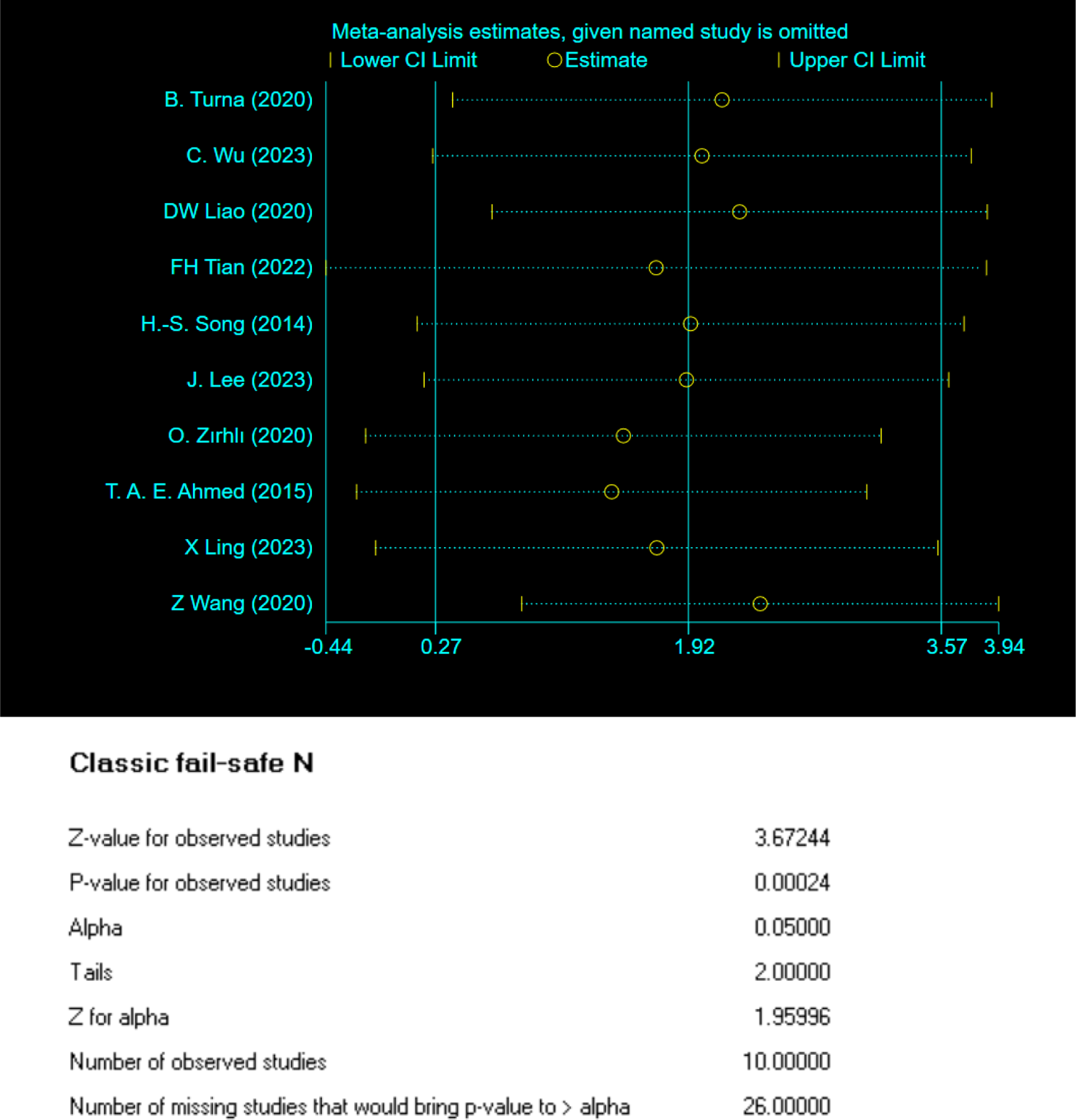
Sensitivity analysis - 1RM GS.

**Figure 22:**
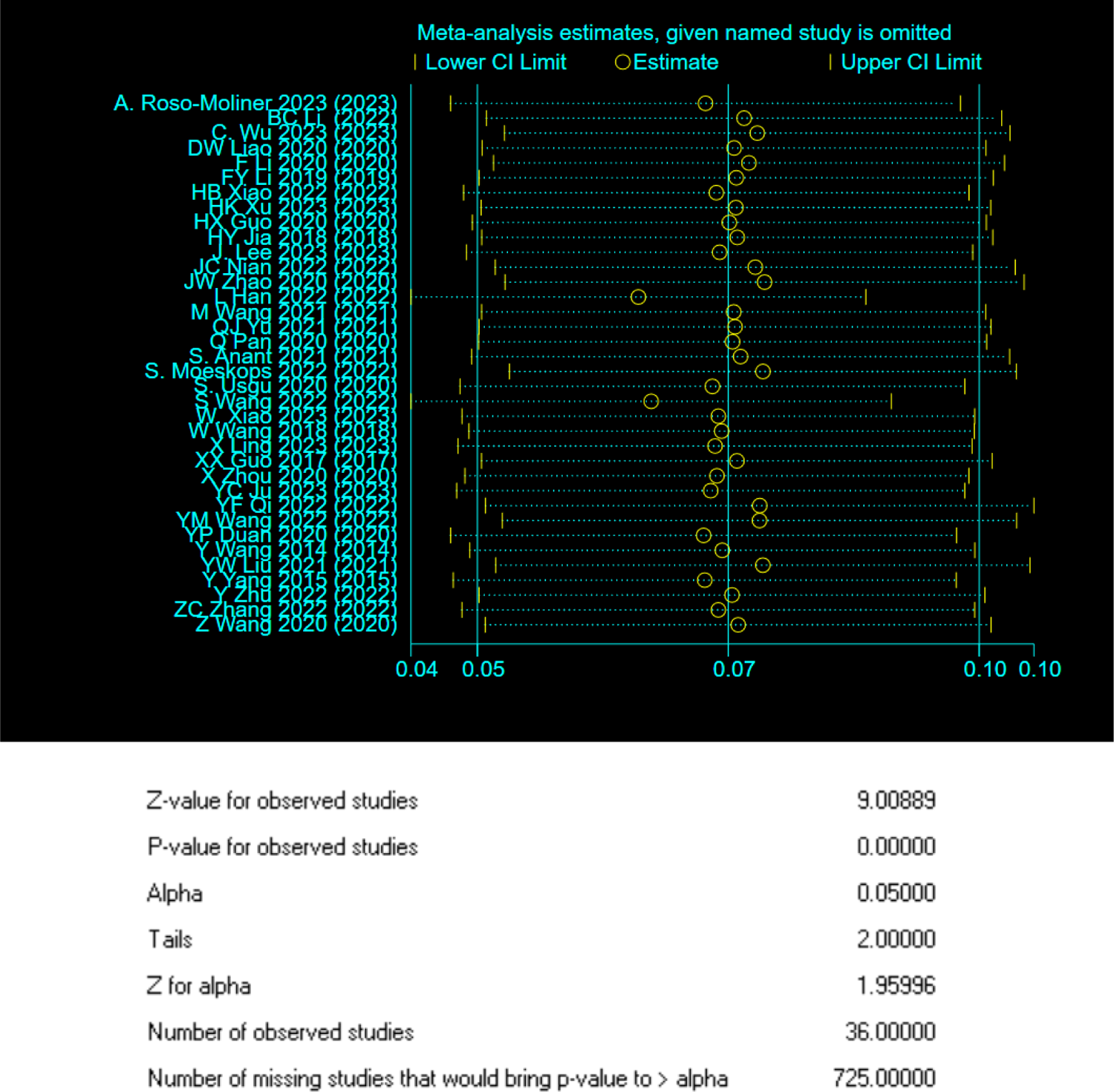
Sensitivity analysis - SLJ.

**Figure 23:**
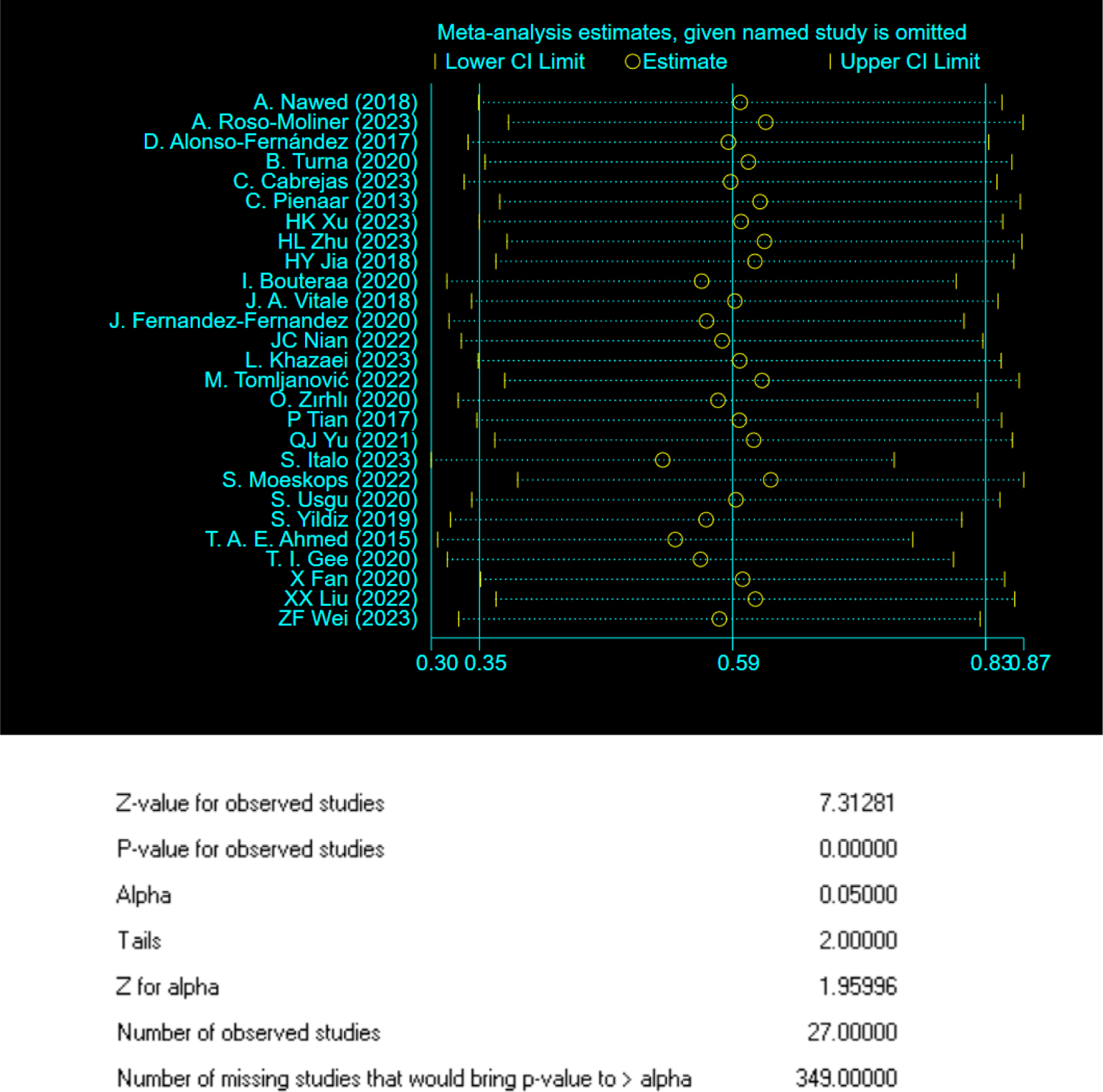
Sensitivity analysis - CMJ.

**Figure 24:**
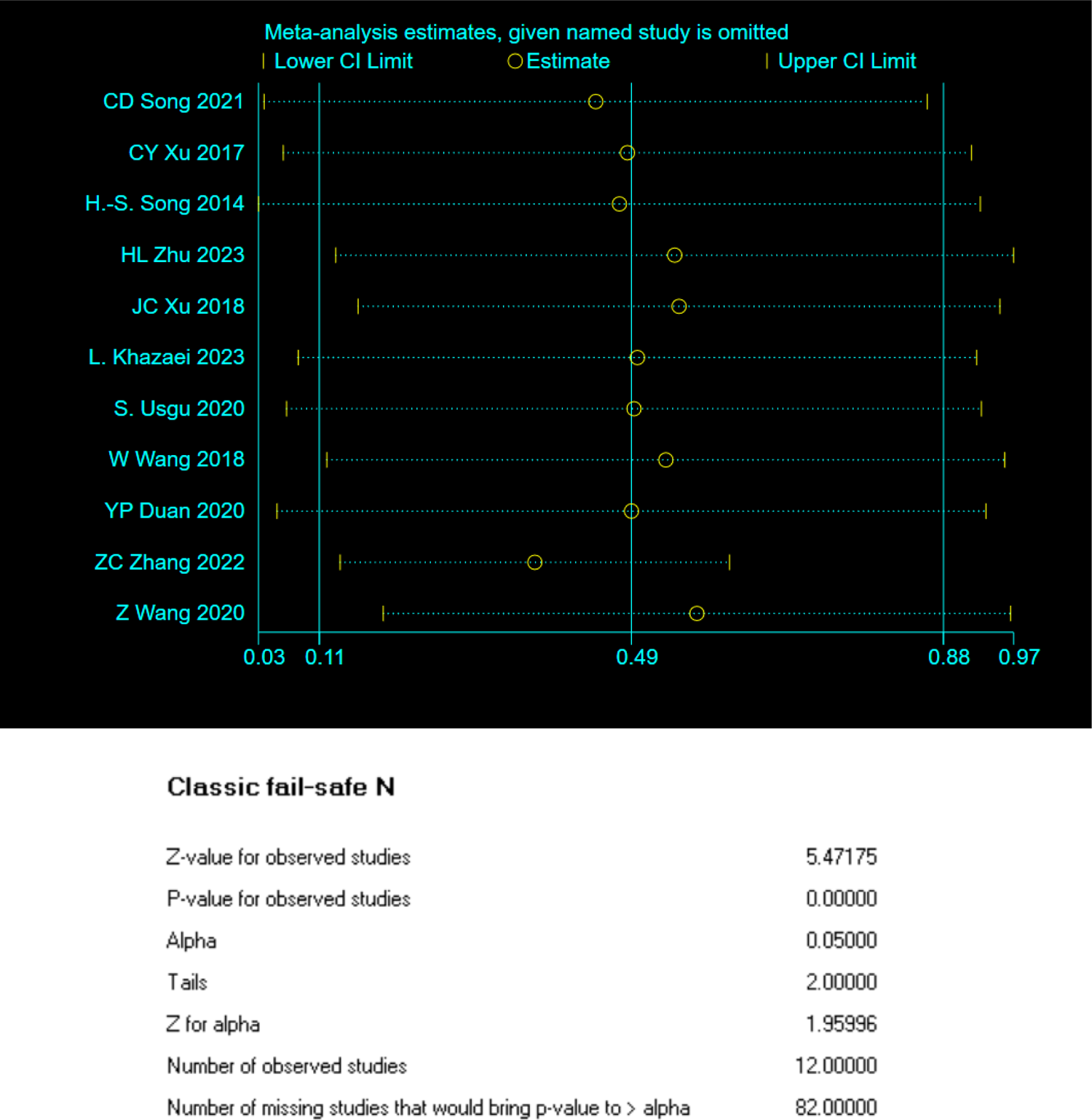
Sensitivity analysis 5 - TMBP.

**Figure 25:**
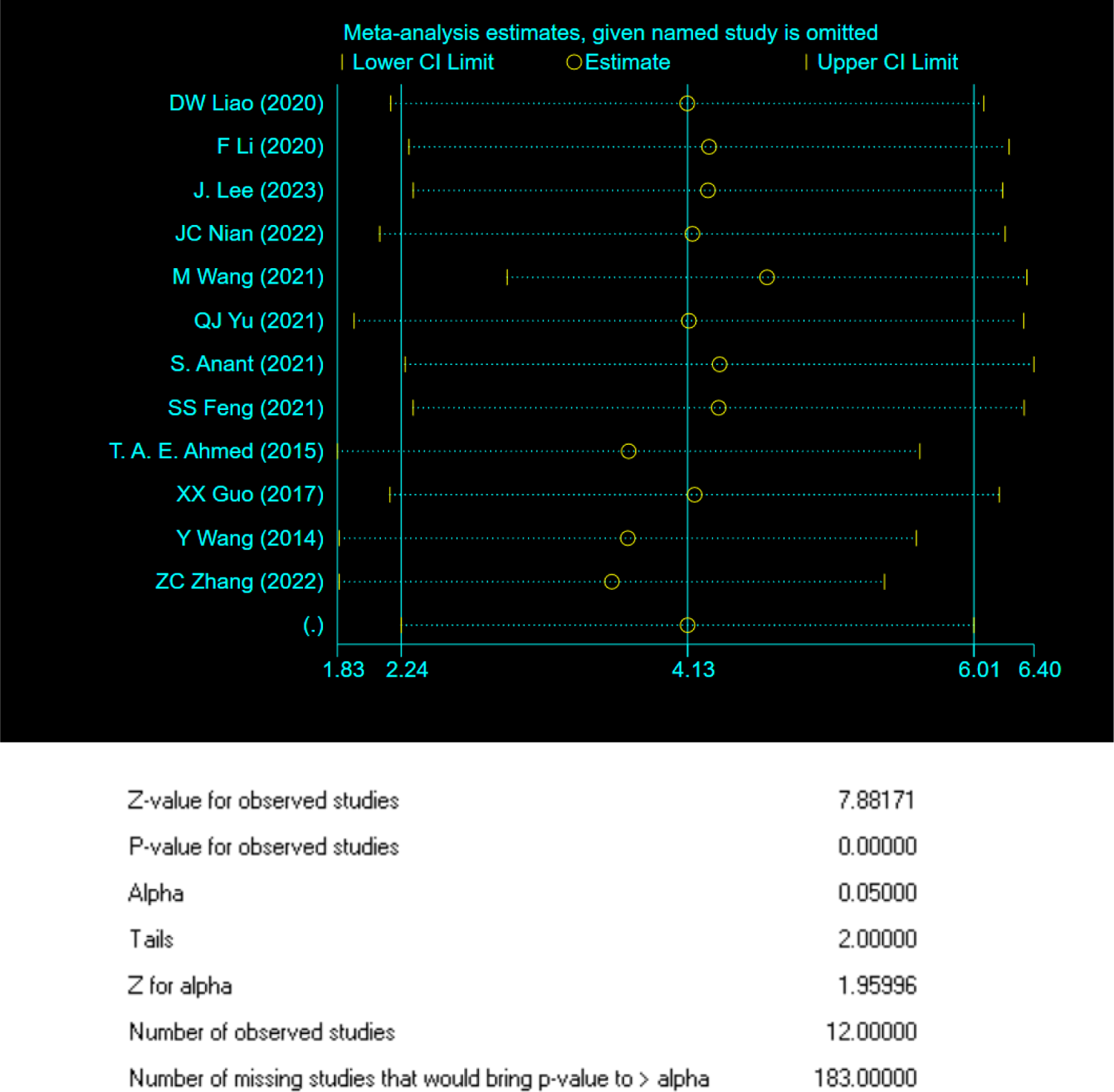
Sensitivity analysis 5 - 1min SU.

**Figure 26:**
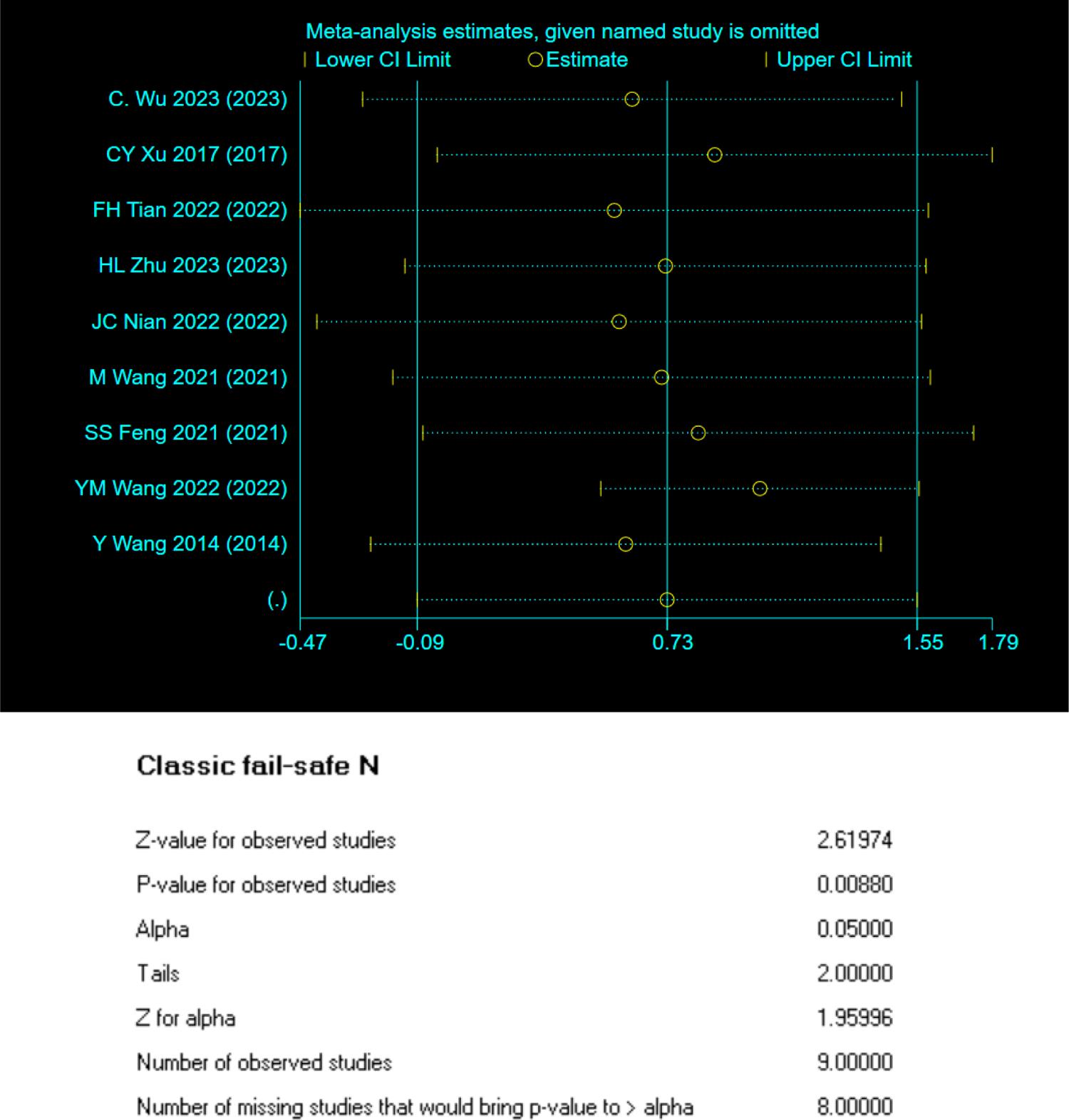
Sensitivity analysis - PLU.

**Figure 27:**
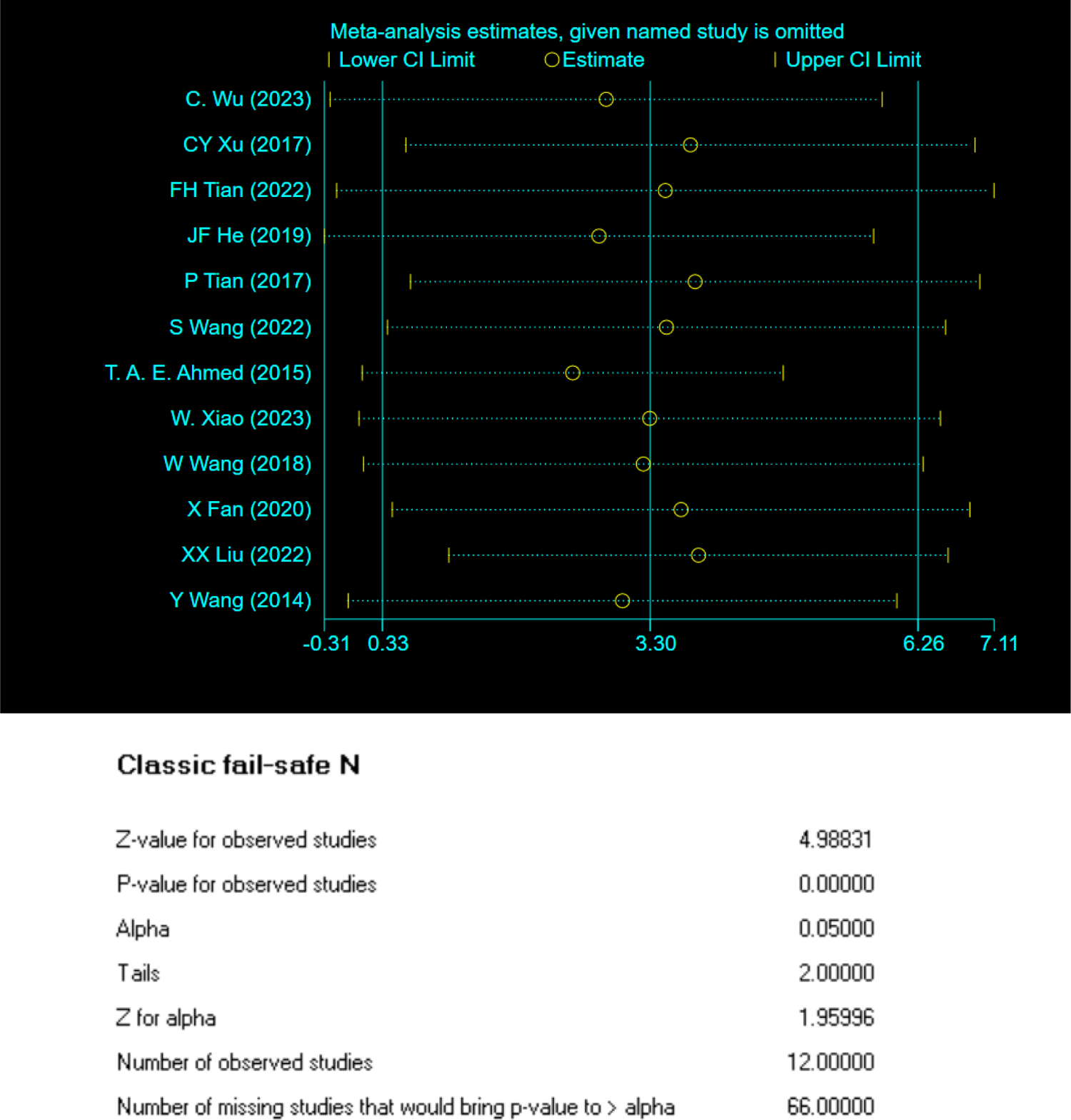
Sensitivity analysis - PSU.

**Figure 28:**
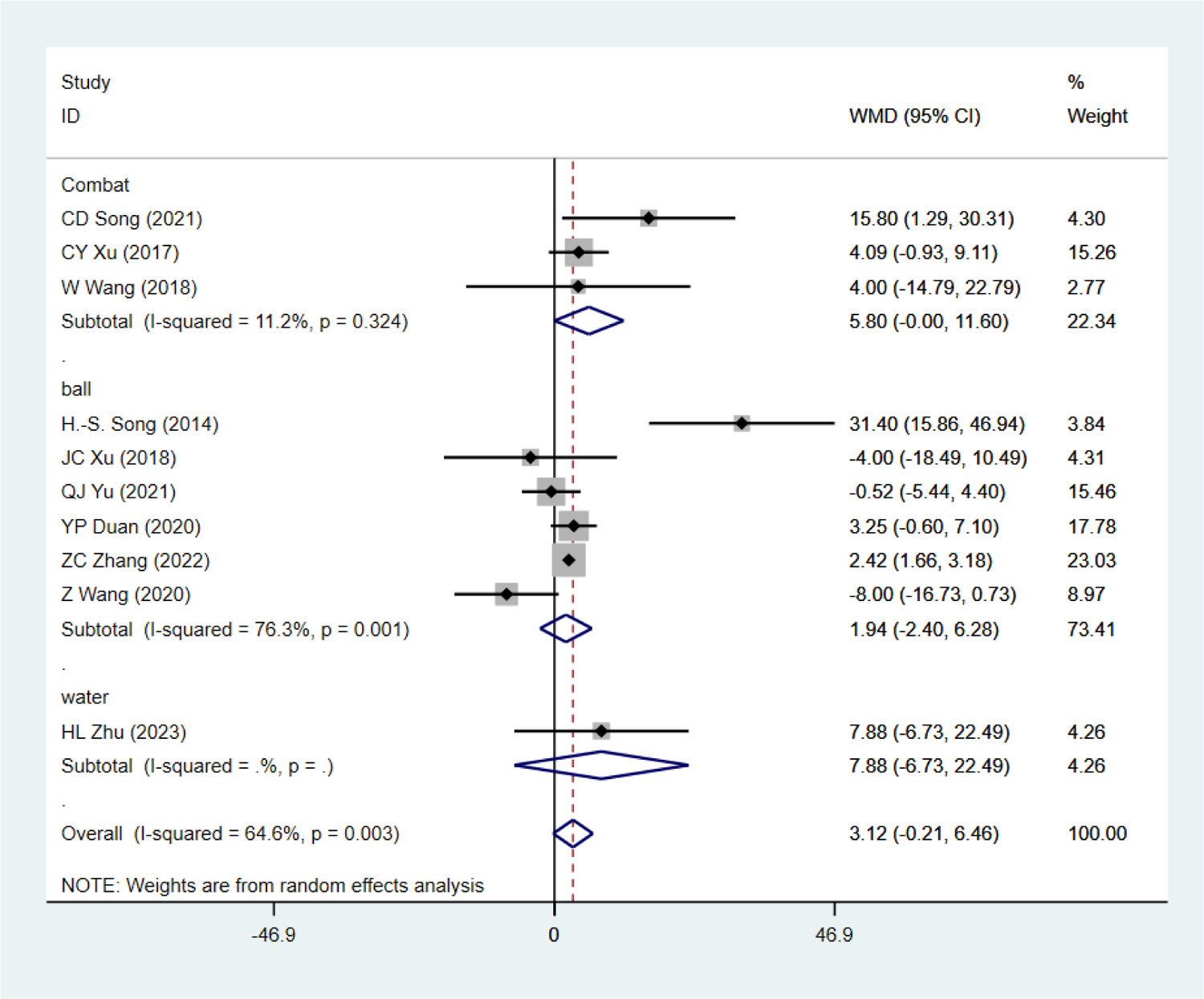
Subgroup Analysis - 1RM D/HS)

**Figure 29:**
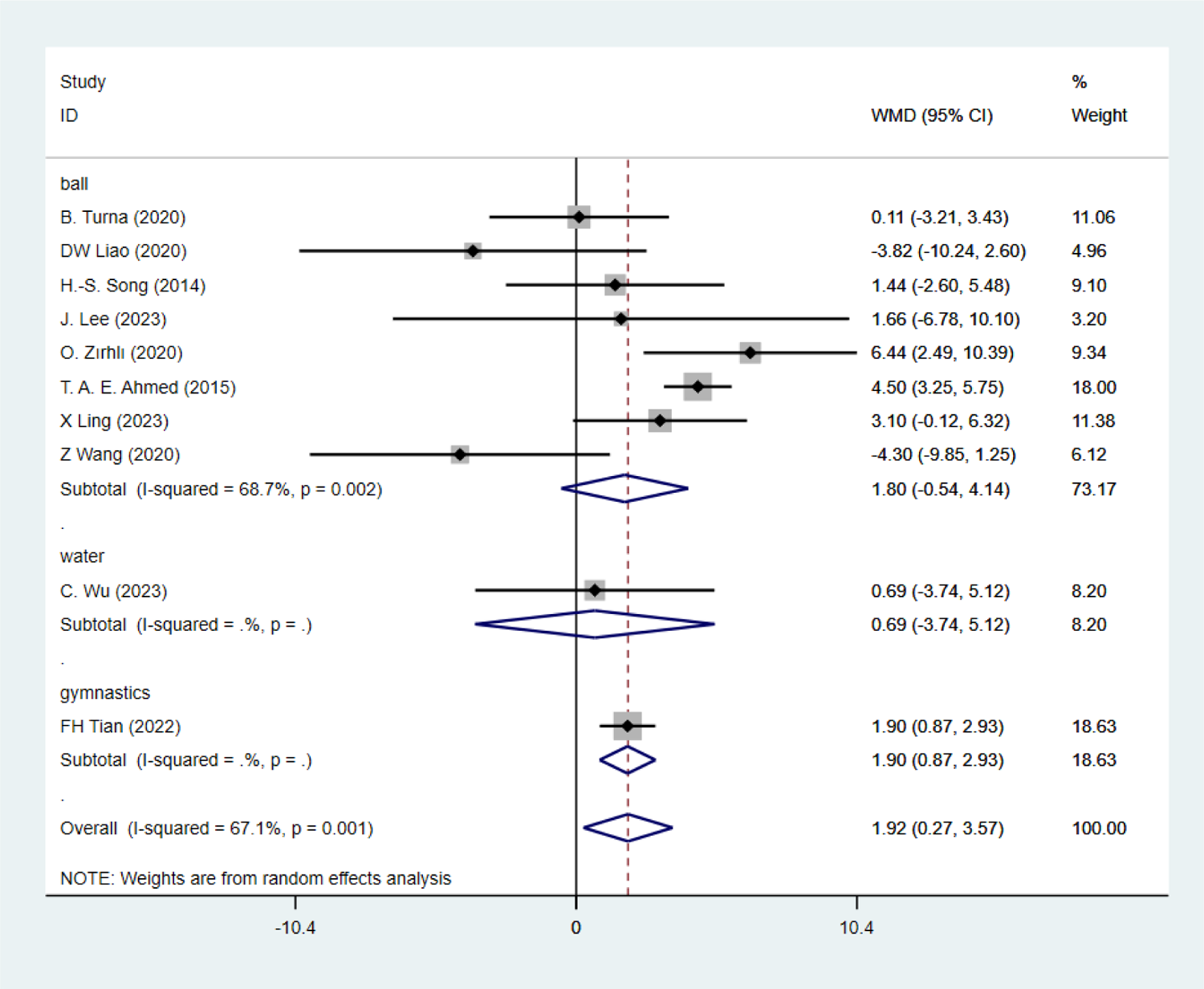
Subgroup Analysis - 1RM GS

**Figure 30:**
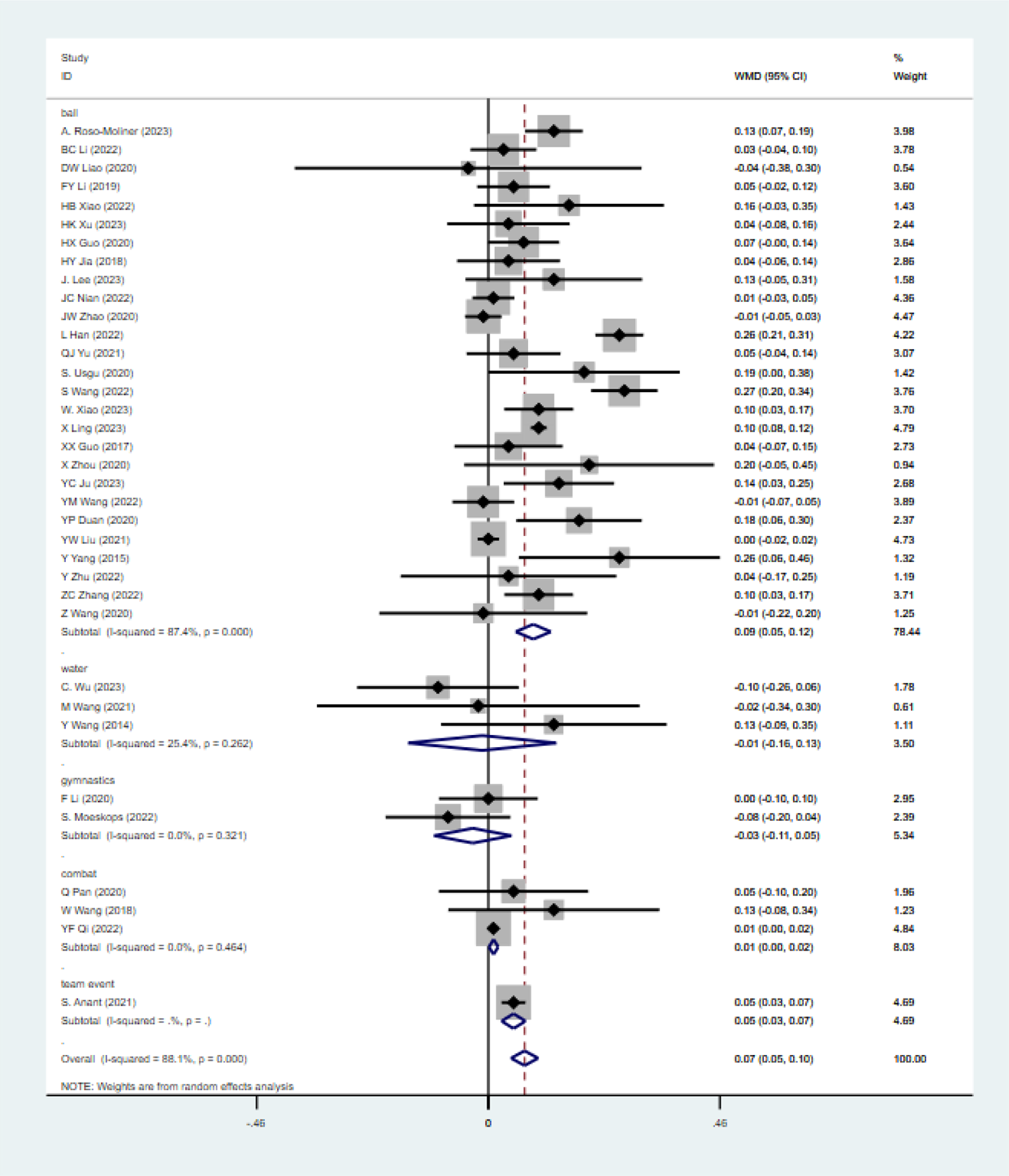
Subgroup Analysis - SLJ

**Figure 31:**
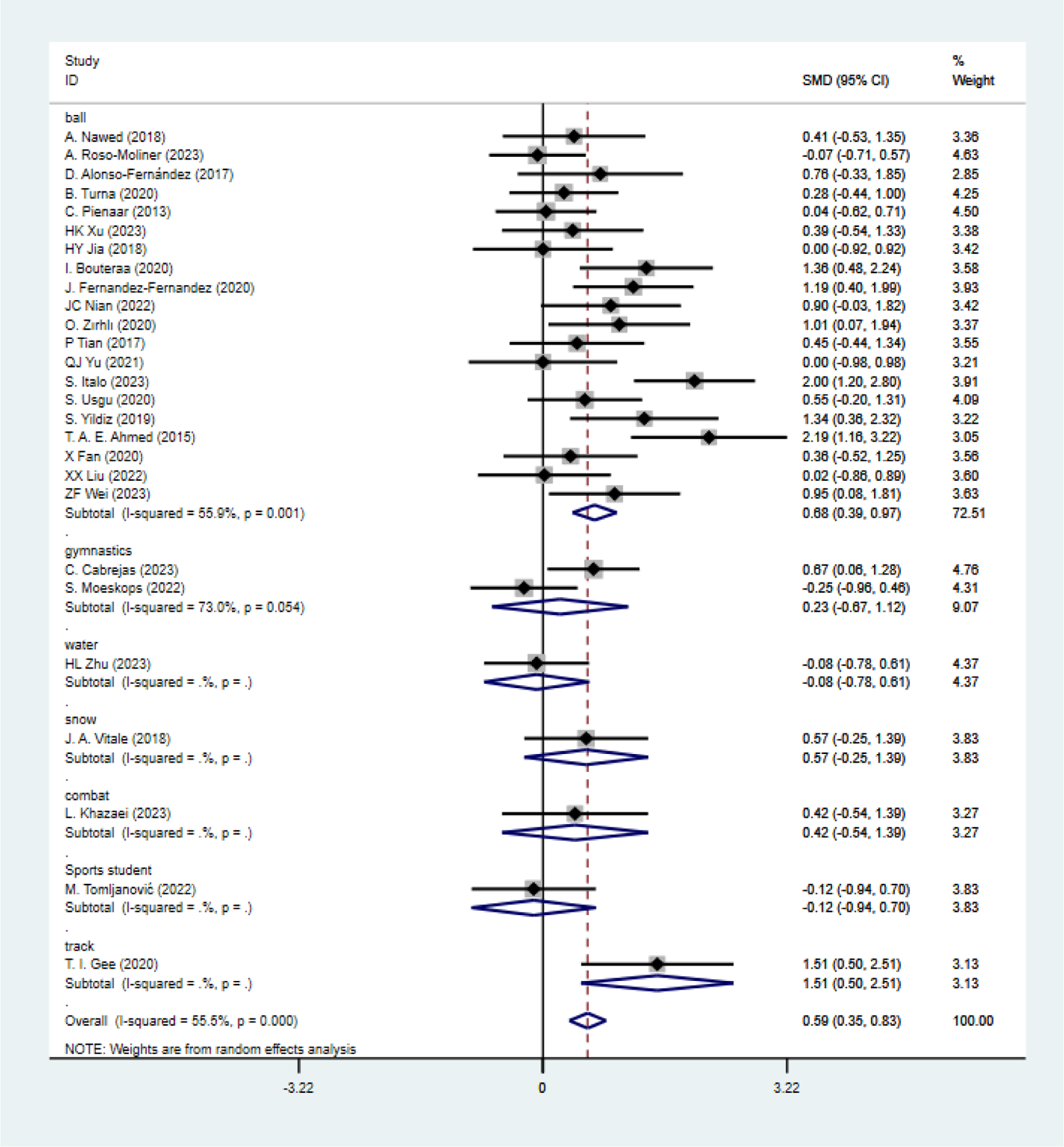
Subgroup Analysis - CMJ

**Figure 32:**
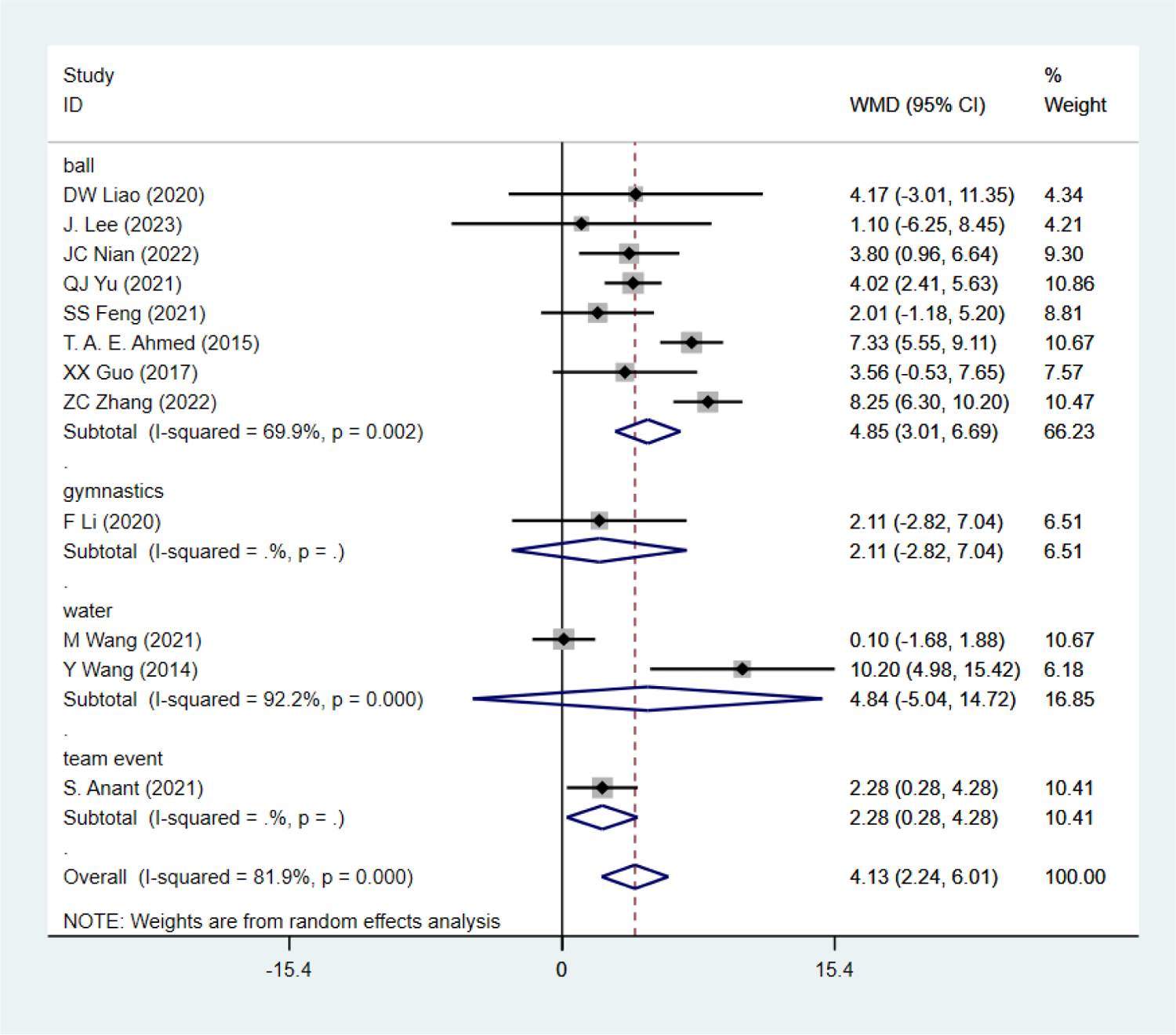
Subgroup Analysis - 1 min SU

**Figure 33:**
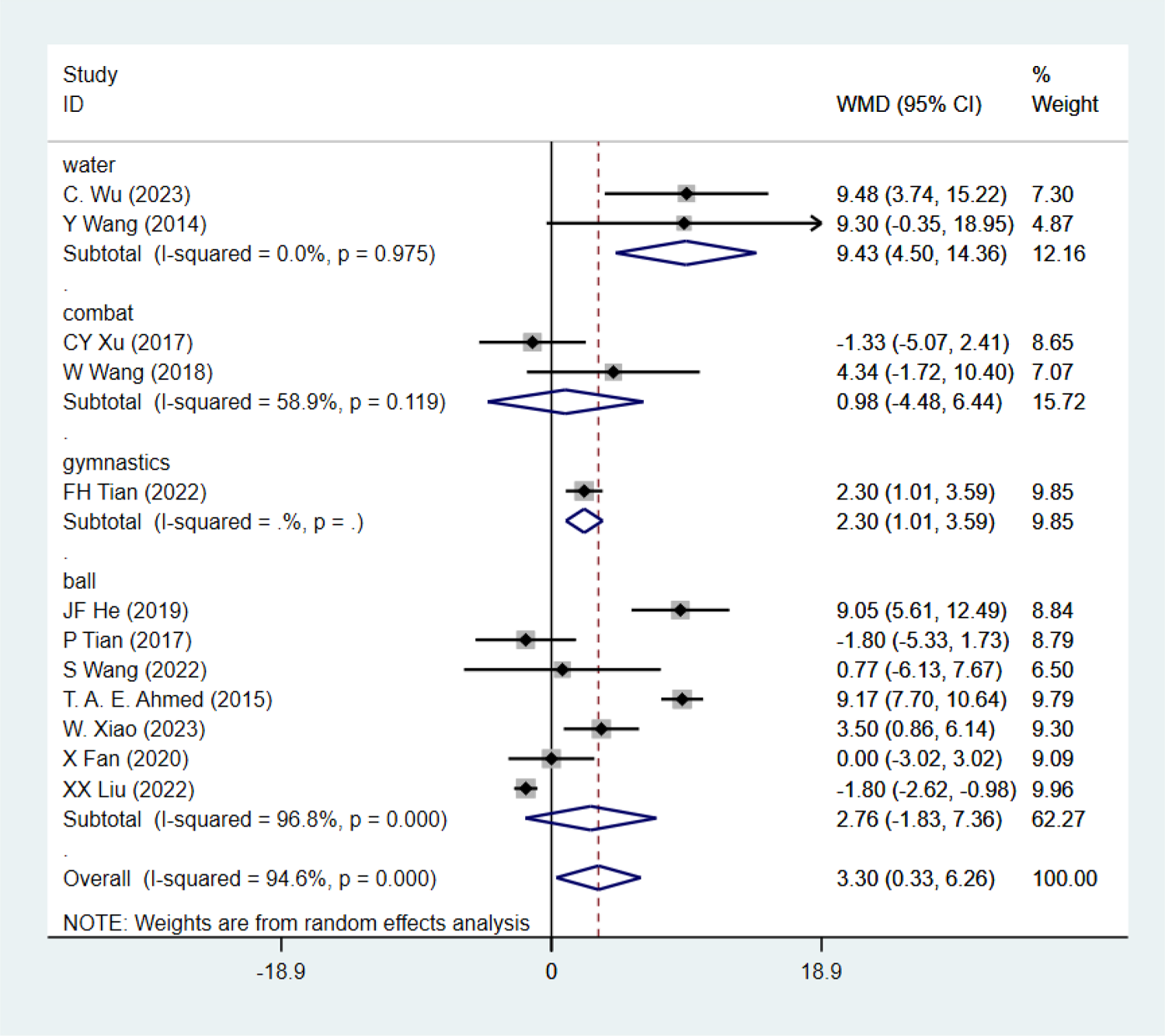
Subgroup Analysis - PSU

**Figure 34:**
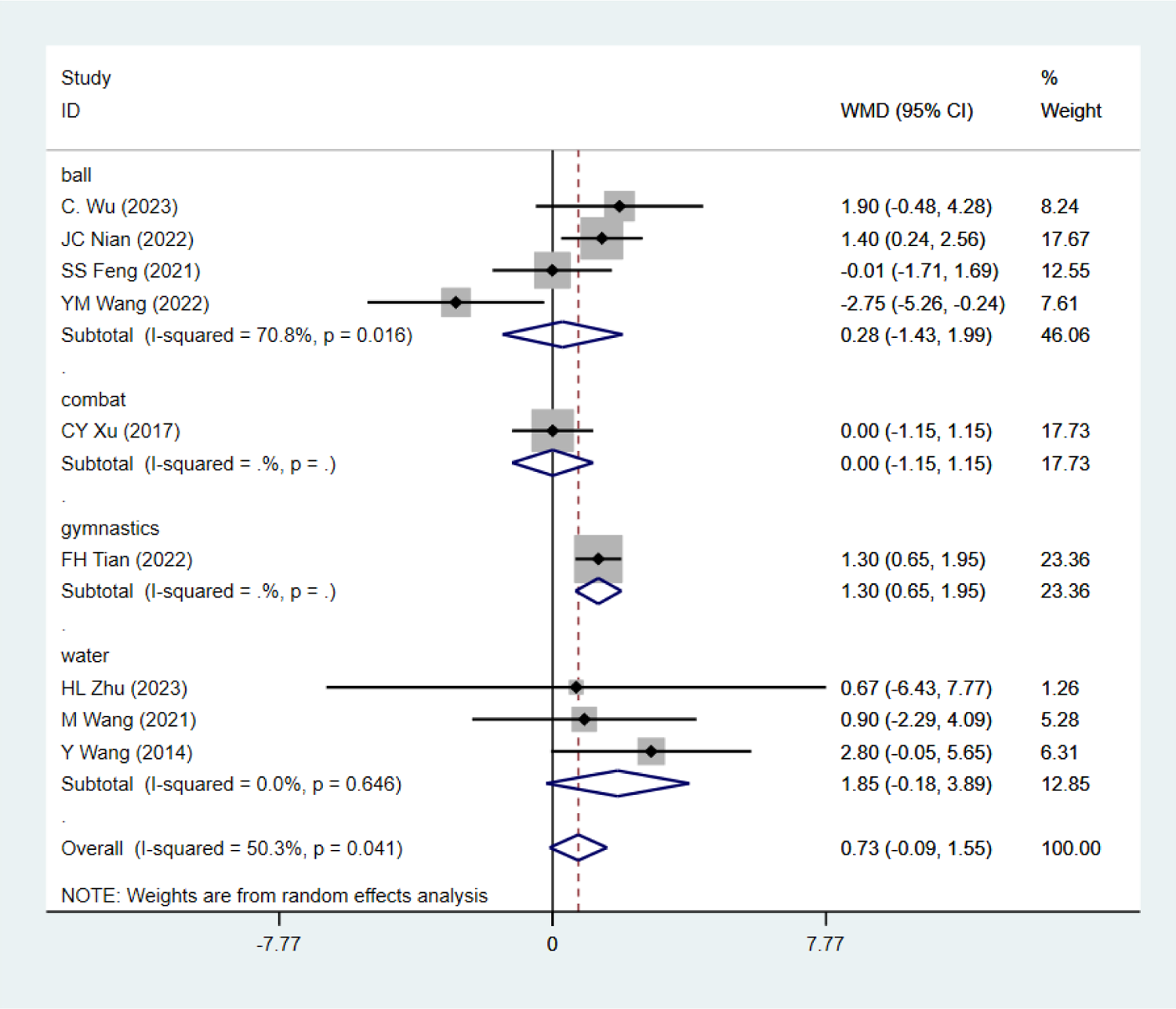
Subgroup Analysis - PLU

**Figure 35:**
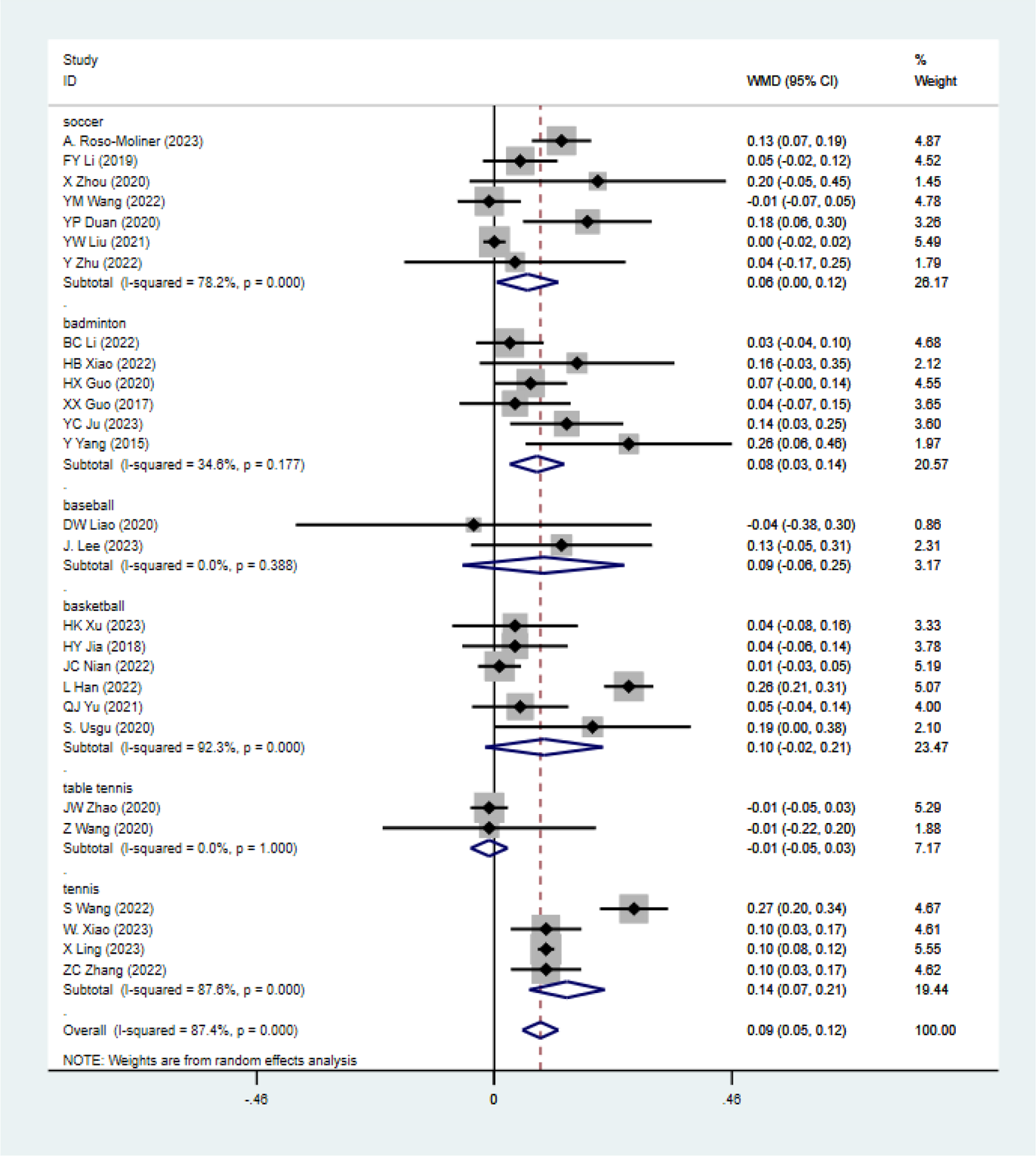
Subgroup Analysis - Impact of FT vs. TRT on Power (SLJ) in Ball Sport Athletes

**Figure 36:**
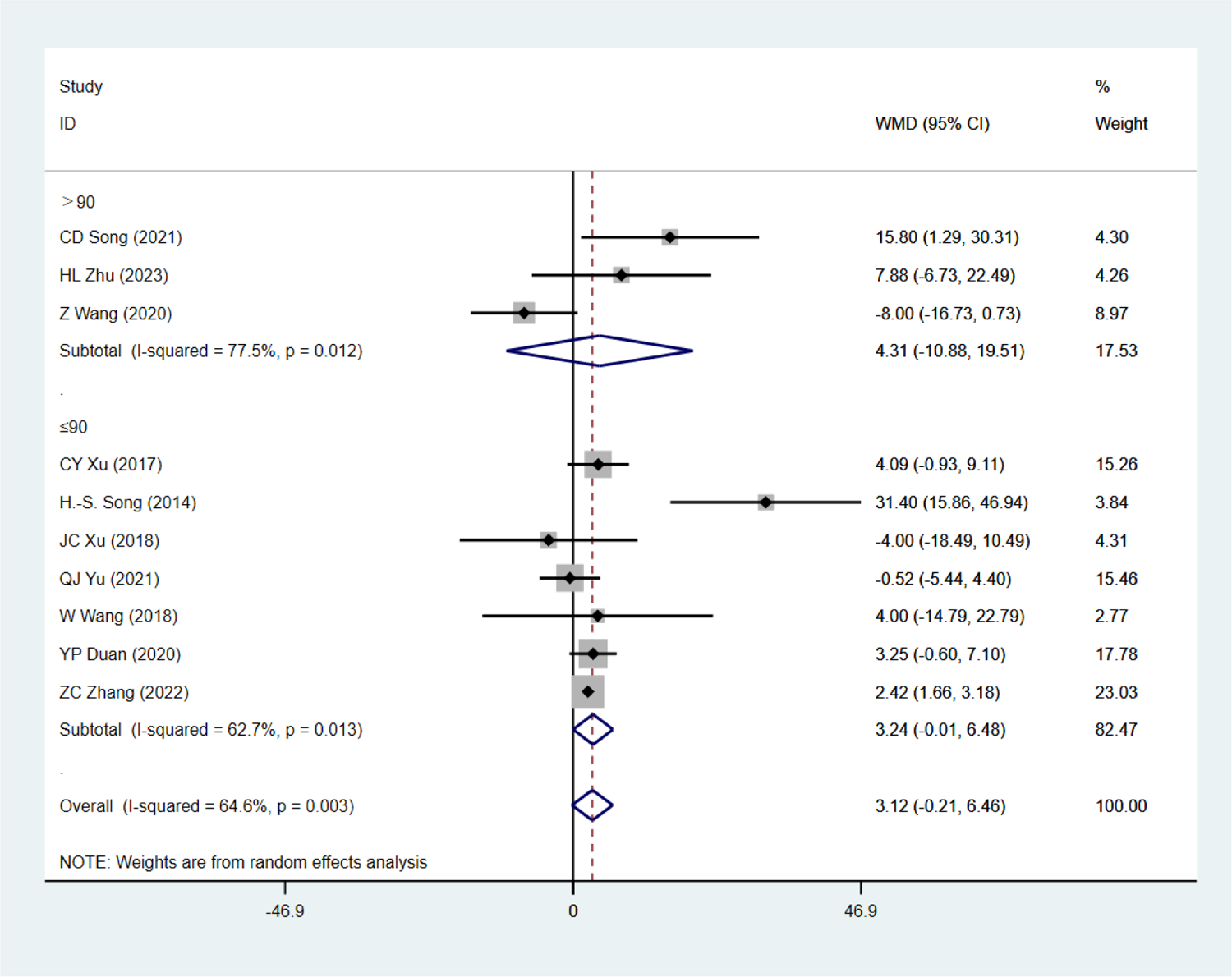
Single factor analysis on 1RM D/HS - Each session duration.

**Figure 37:**
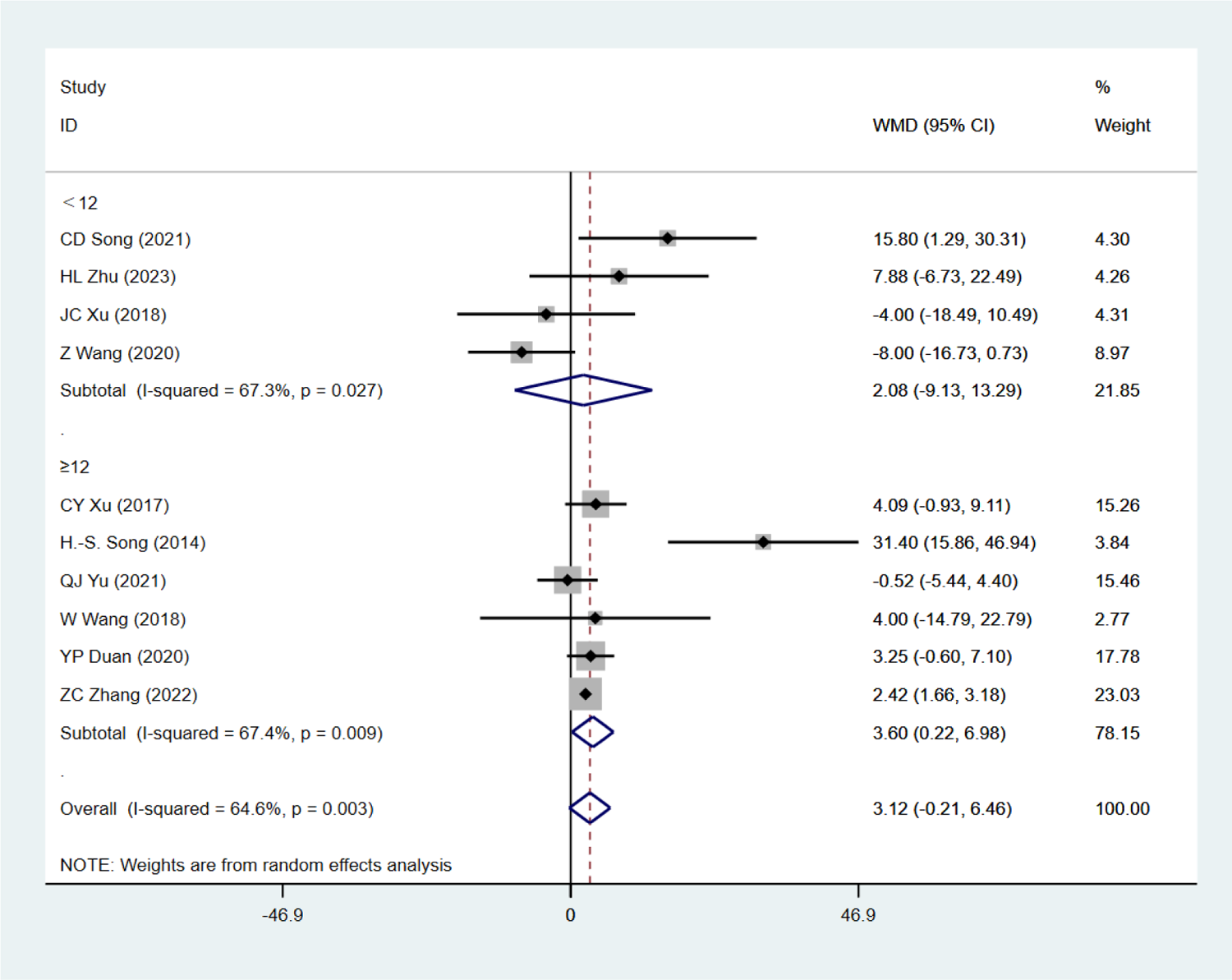
Single factor analysis on 1RM D/HS - Total weeks

**Figure 38:**
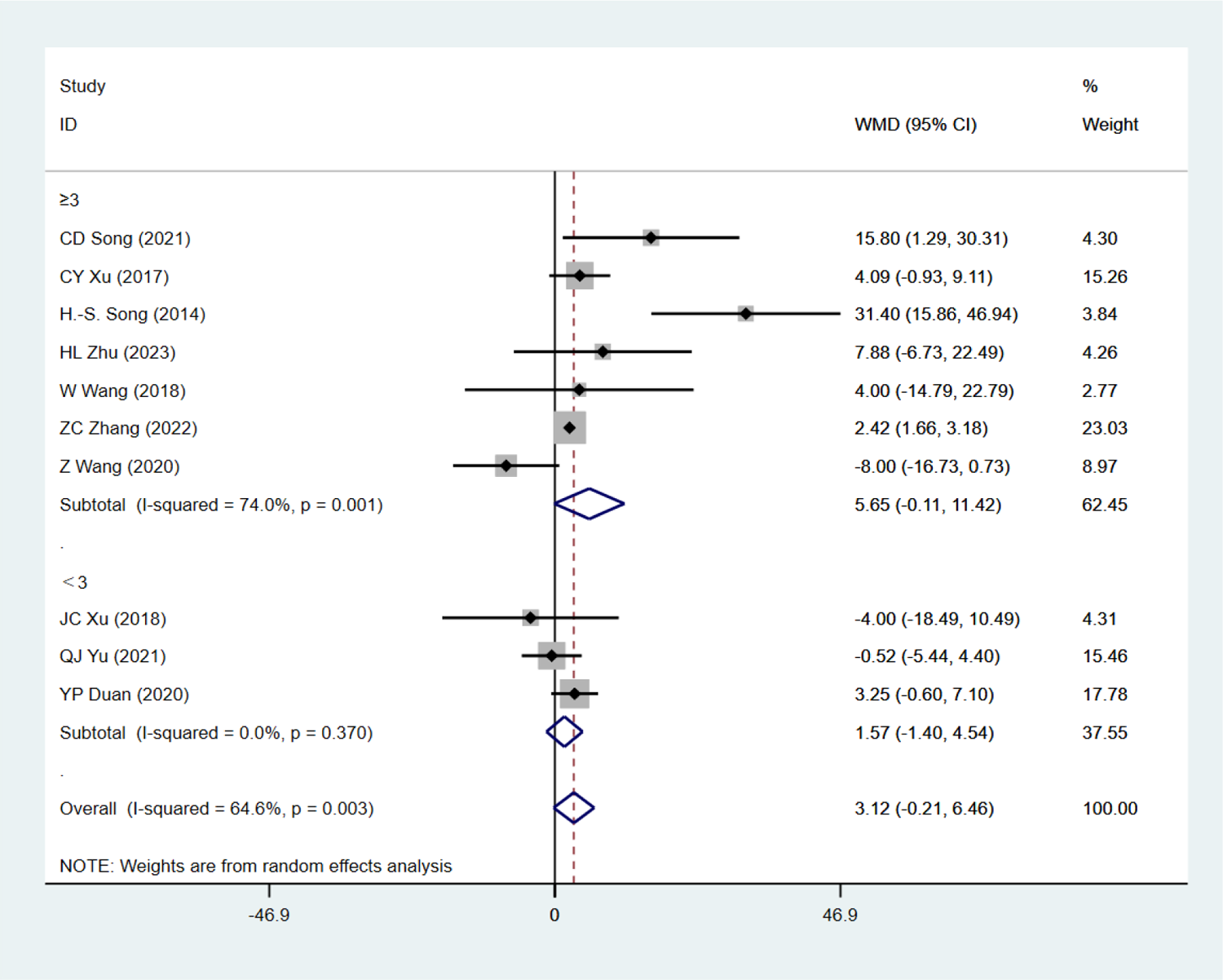
Single factor analysis on 1RM D/HS -Frequency

**Figure 39:**
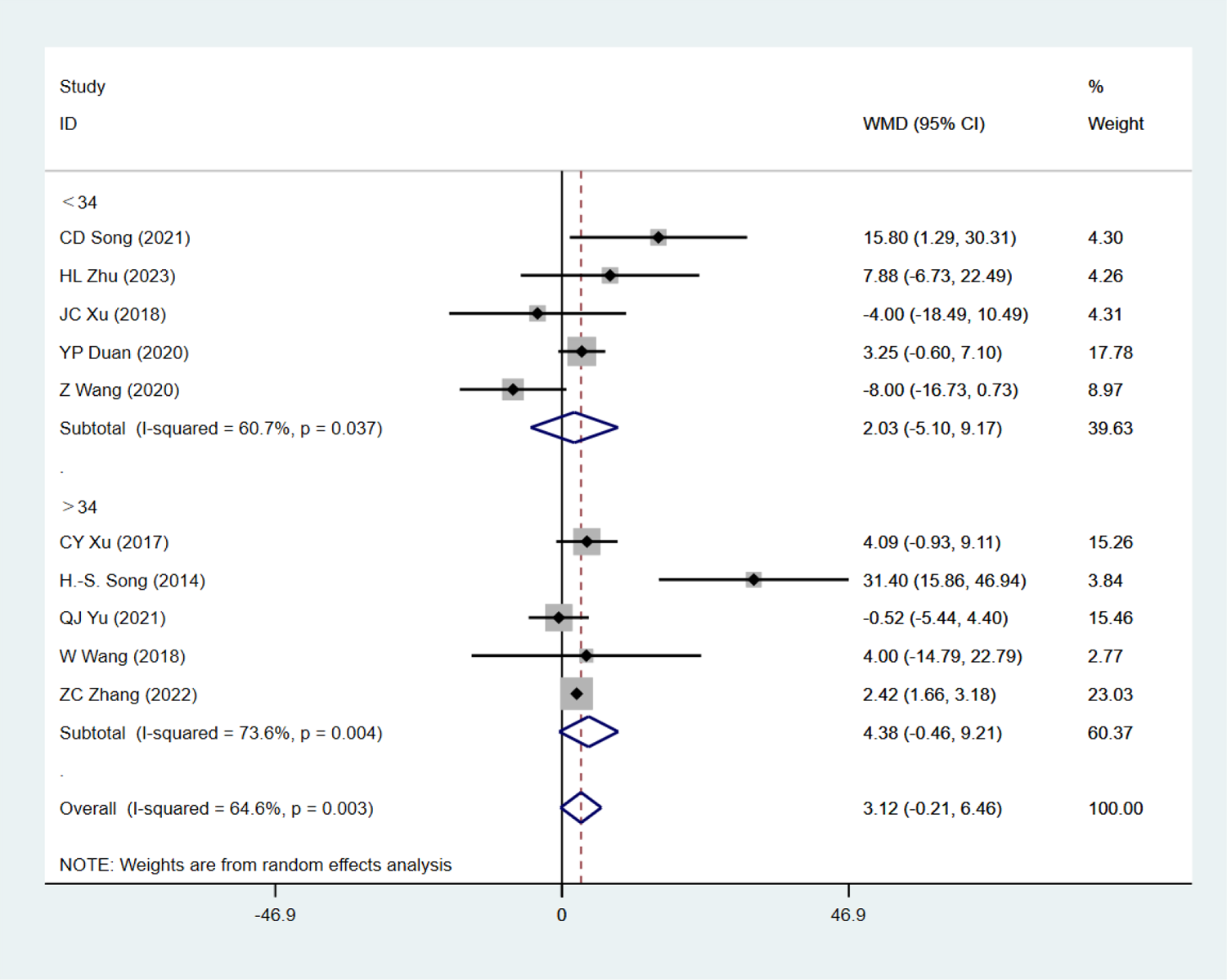
Single factor analysis on 1RM D/HS - Total sessions

**Figure 40:**
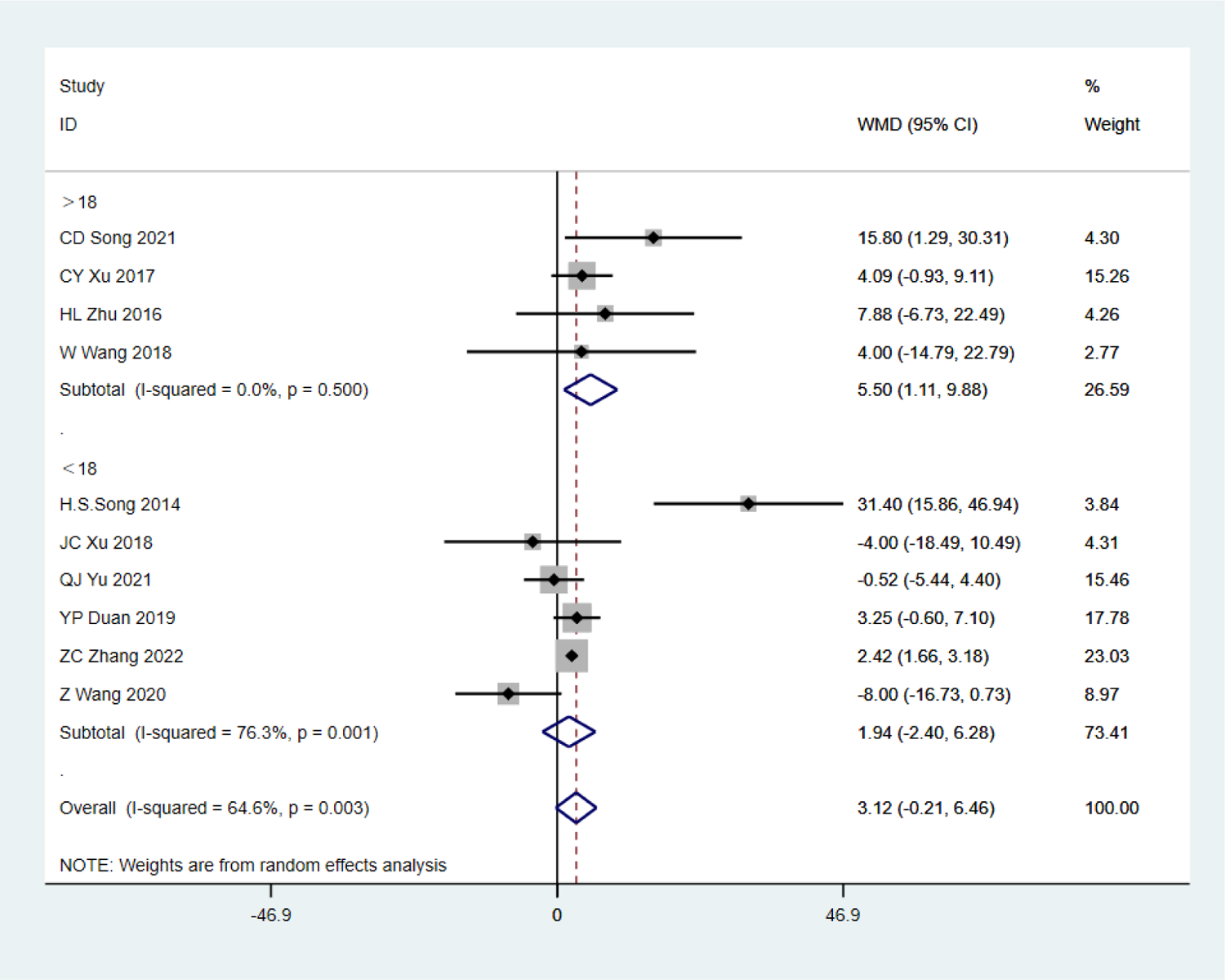
Single factor analysis on 1RM D/HS - Age.

**Figure 41:**
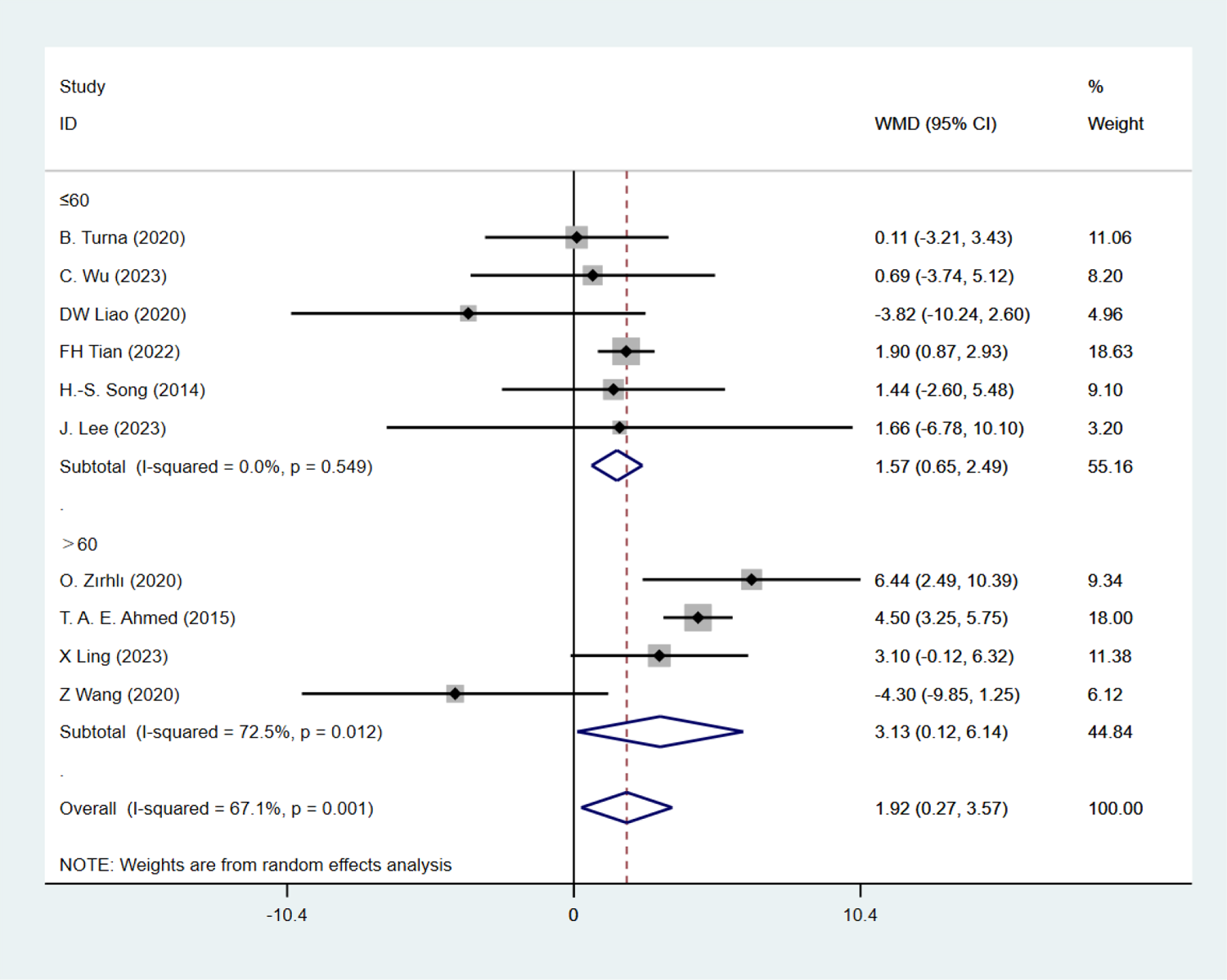
Single factor analysis on 1RM GS - Each session duration.

**Figure 42:**
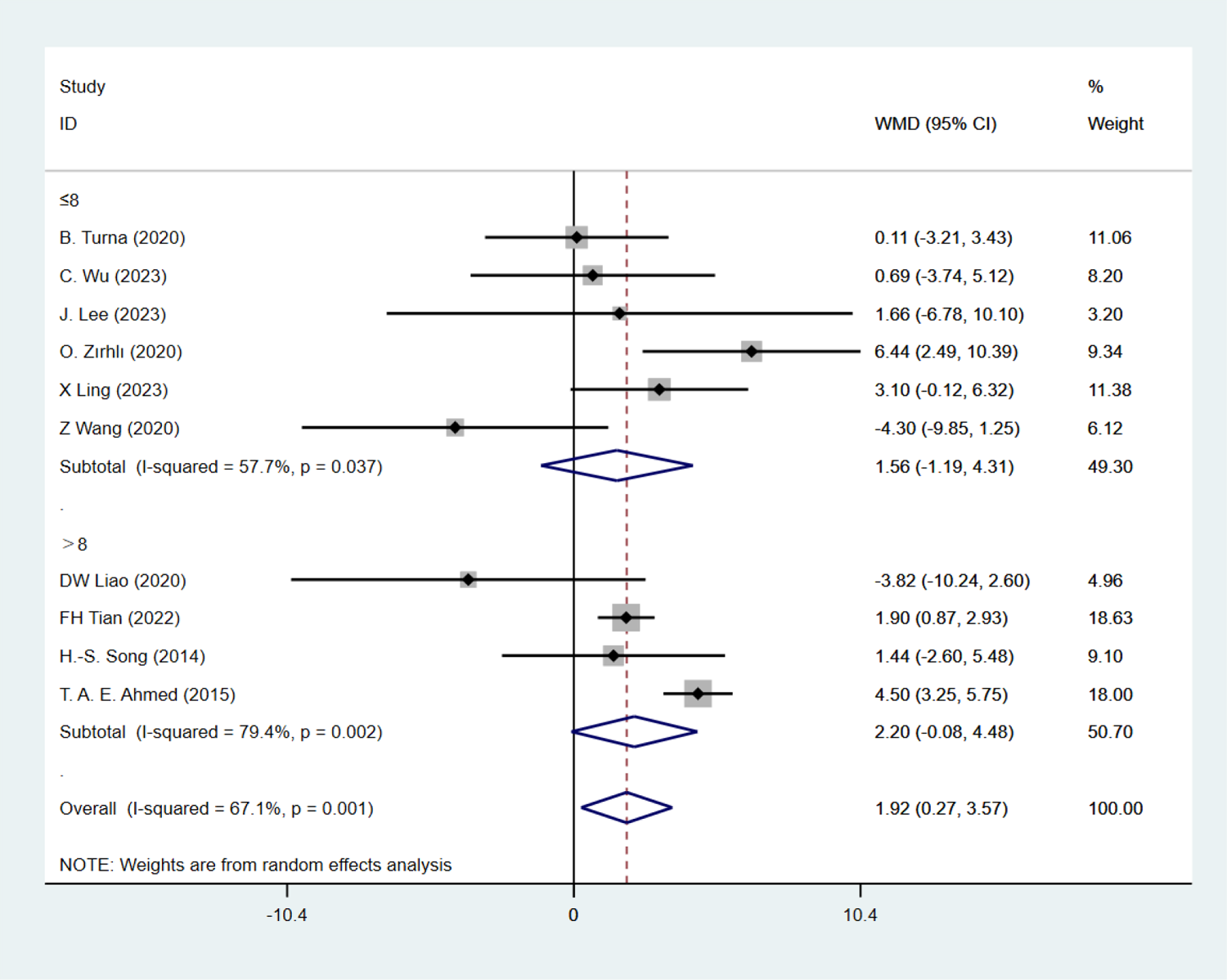
Single factor analysis on 1RM GS - Total weeks

**Figure 43:**
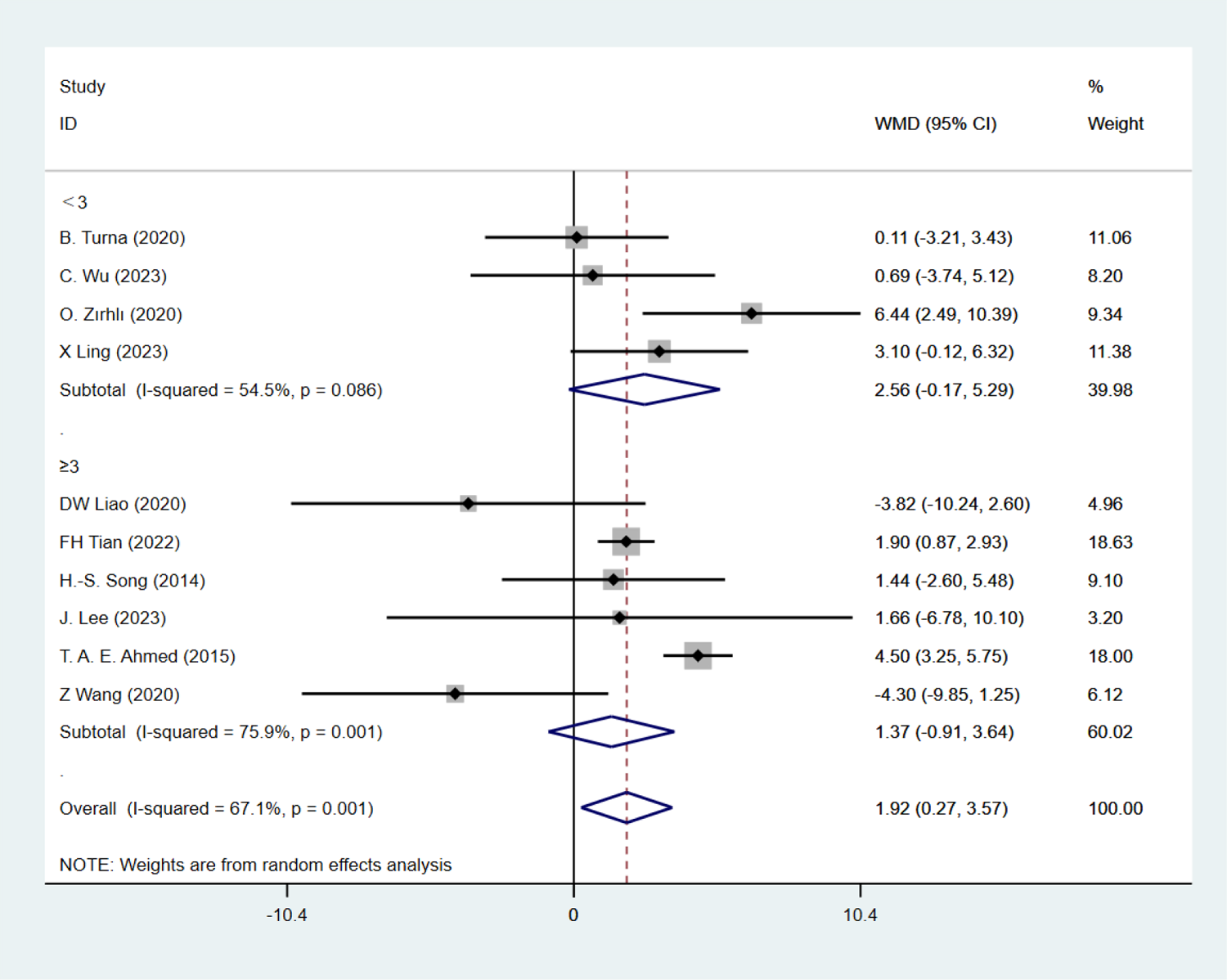
Single factor analysis on 1RM GS - Frequency

**Figure 44:**
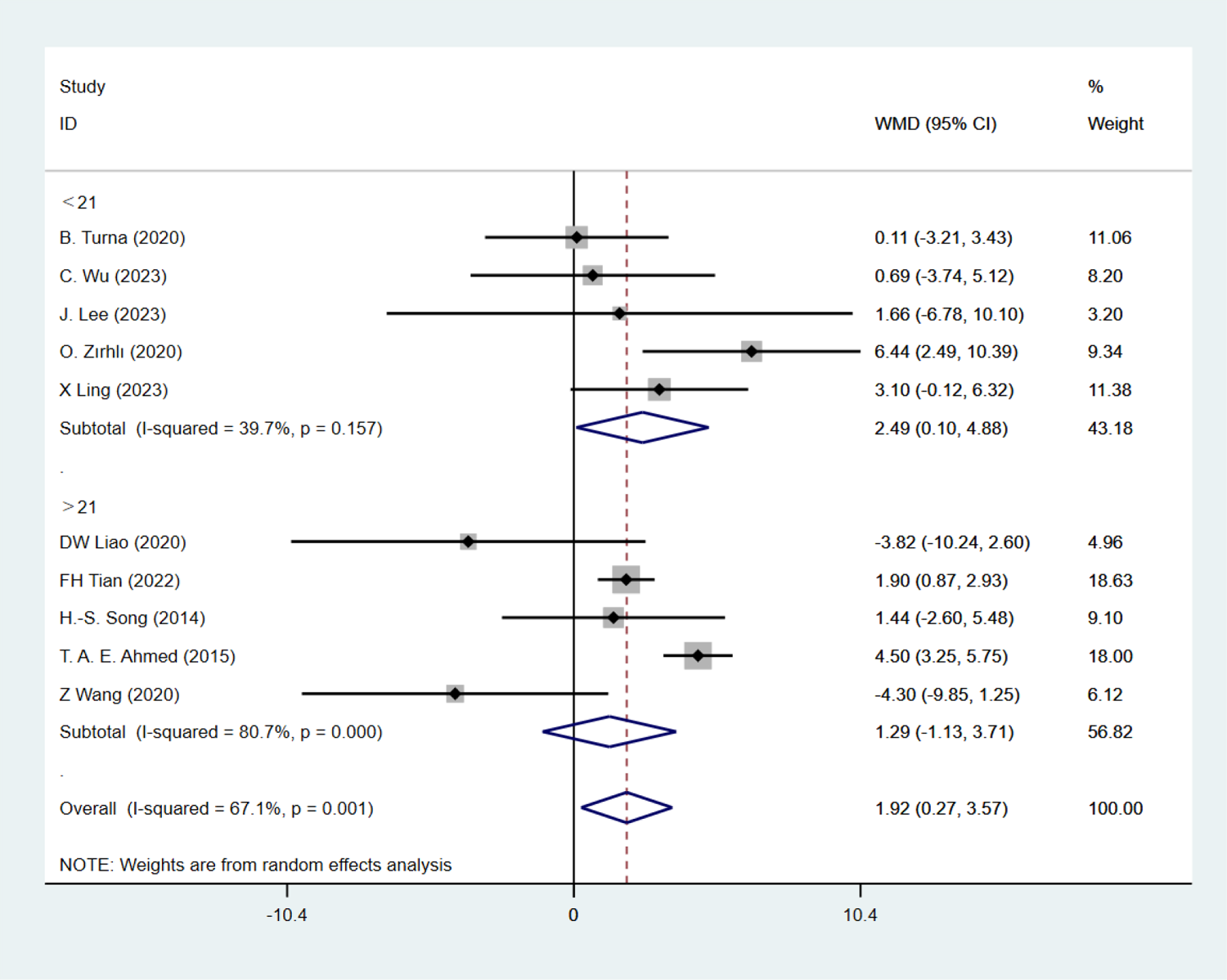
Single factor analysis on 1RM GS - Total sessions

**Figure 45:**
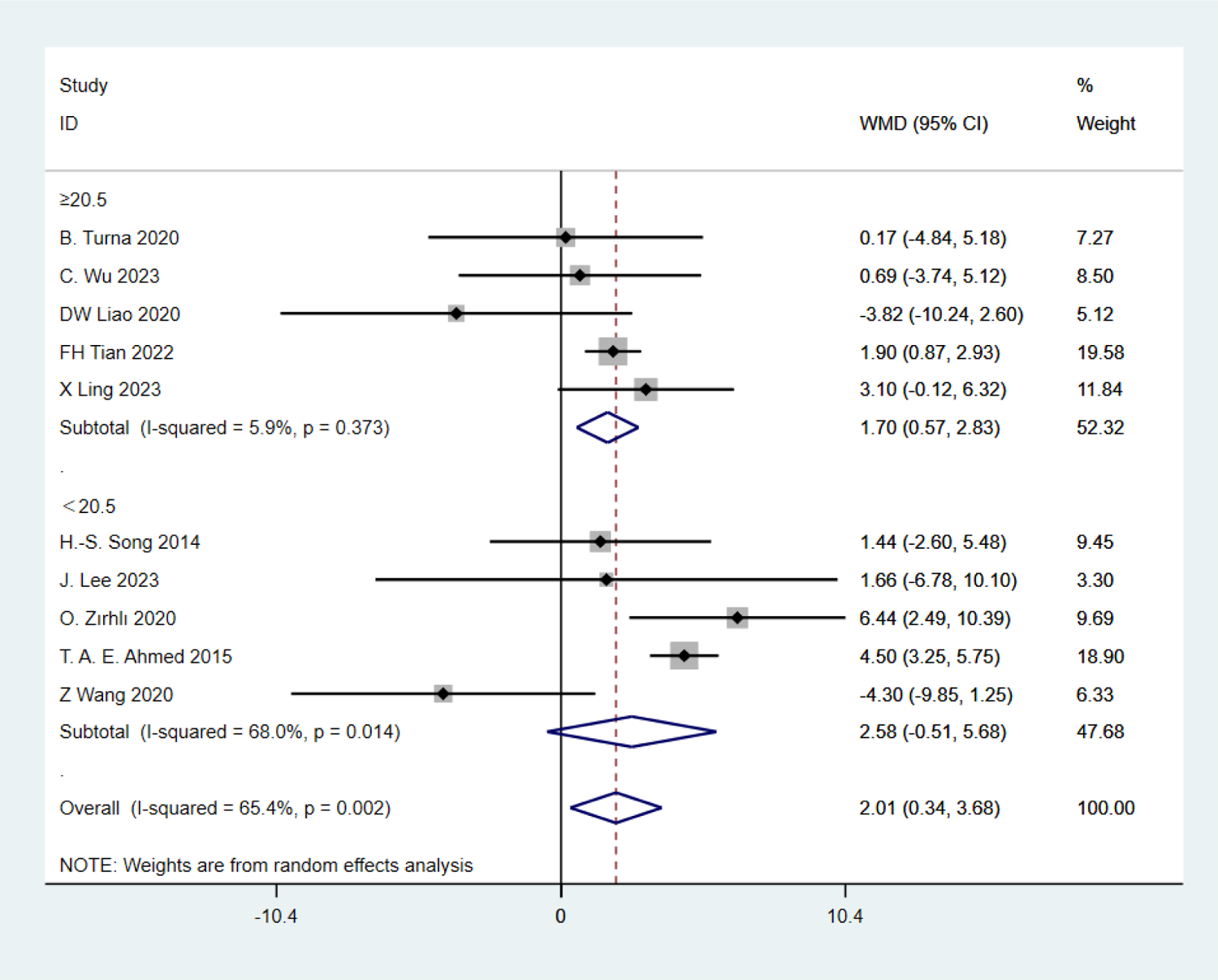
Single factor analysis on 1RM GS - Age

**Figure 46:**
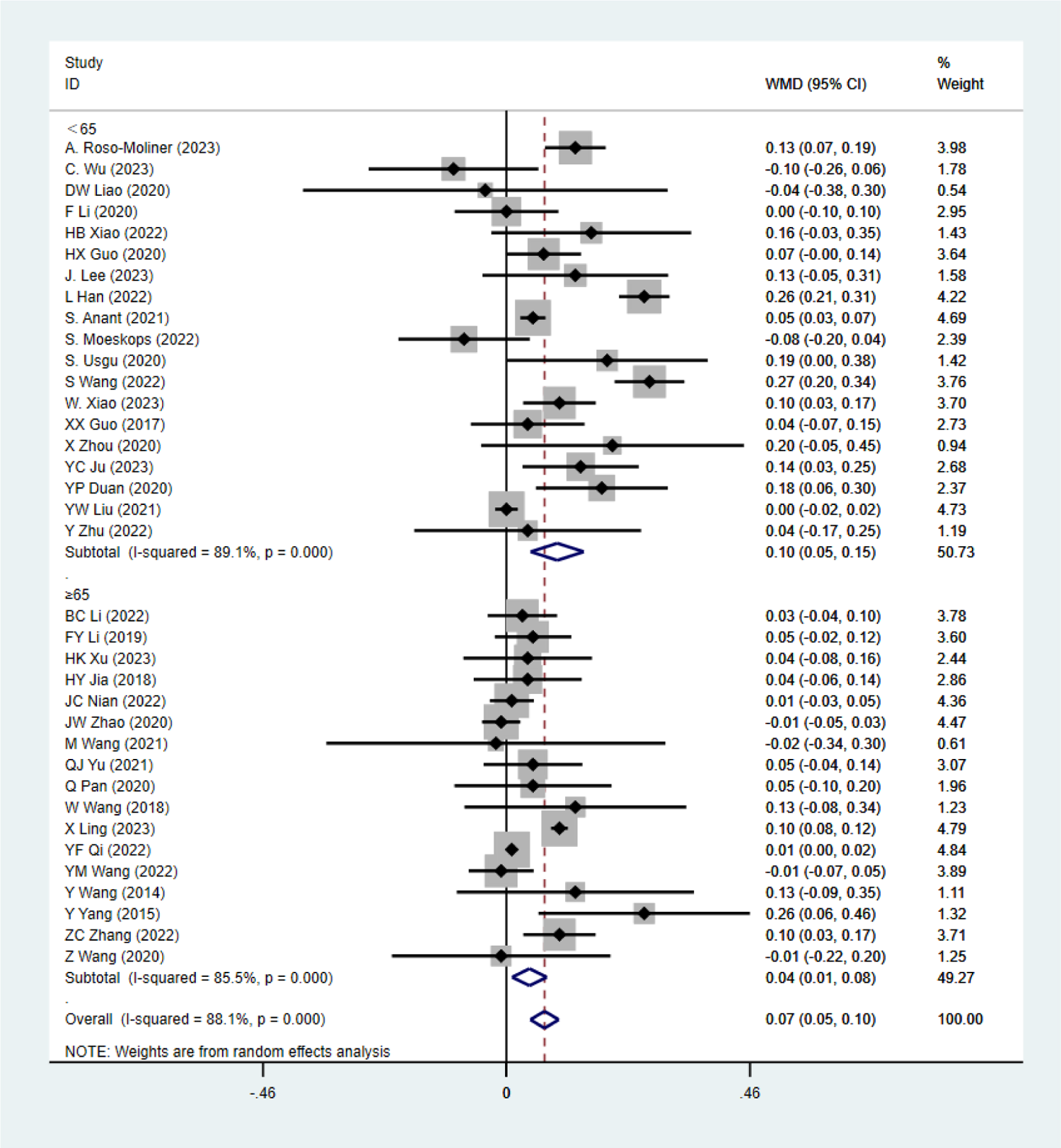
Single factor analysis on SLJ - Each session duration.

**Figure 47:**
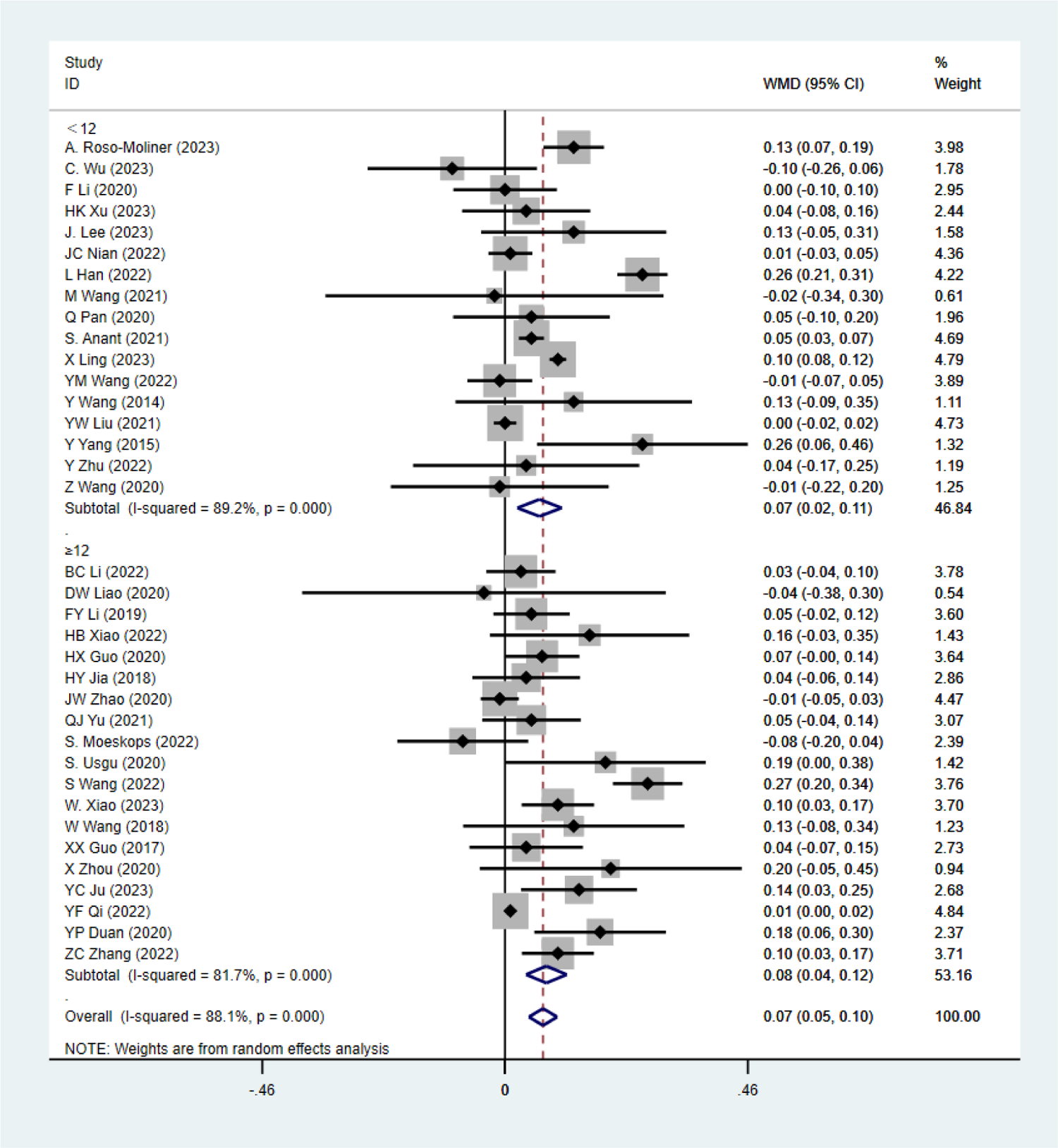
Single factor analysis on SLJ - Total weeks.

**Figure 48:**
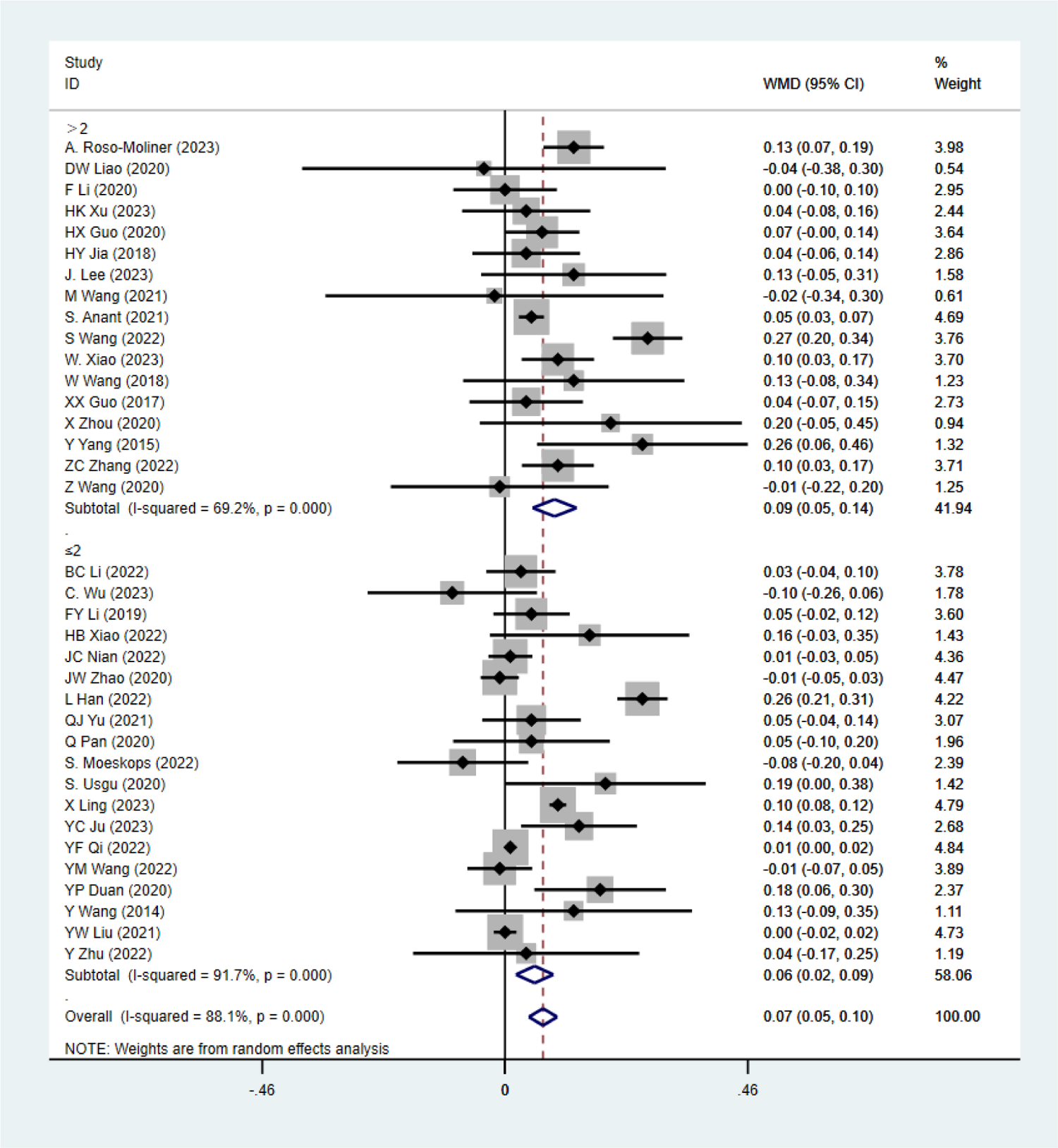
Single factor analysis on SLJ - Frequency

**Figure 49:**
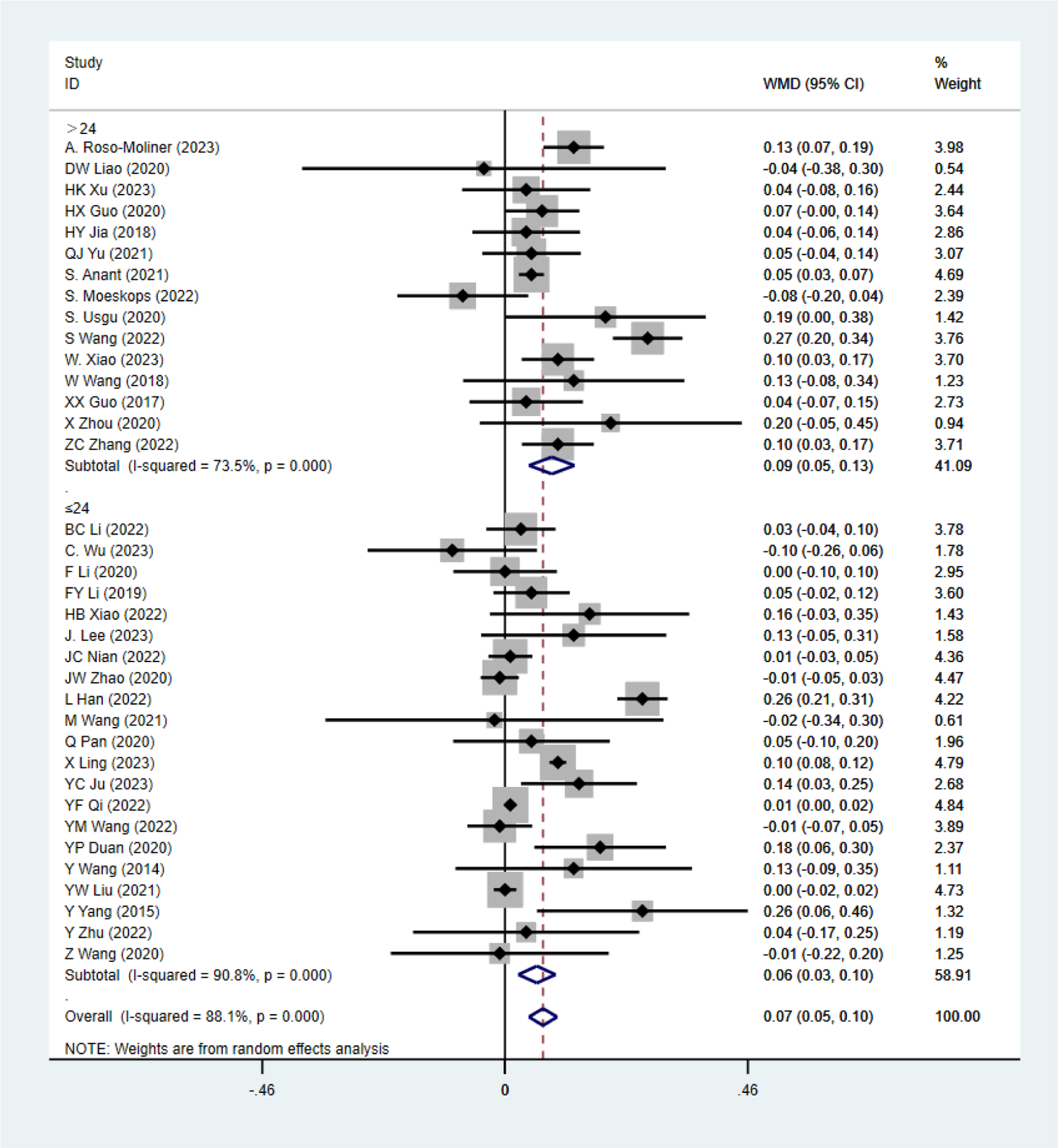
Single factor analysis on SLJ - Total sessions

**Figure 50:**
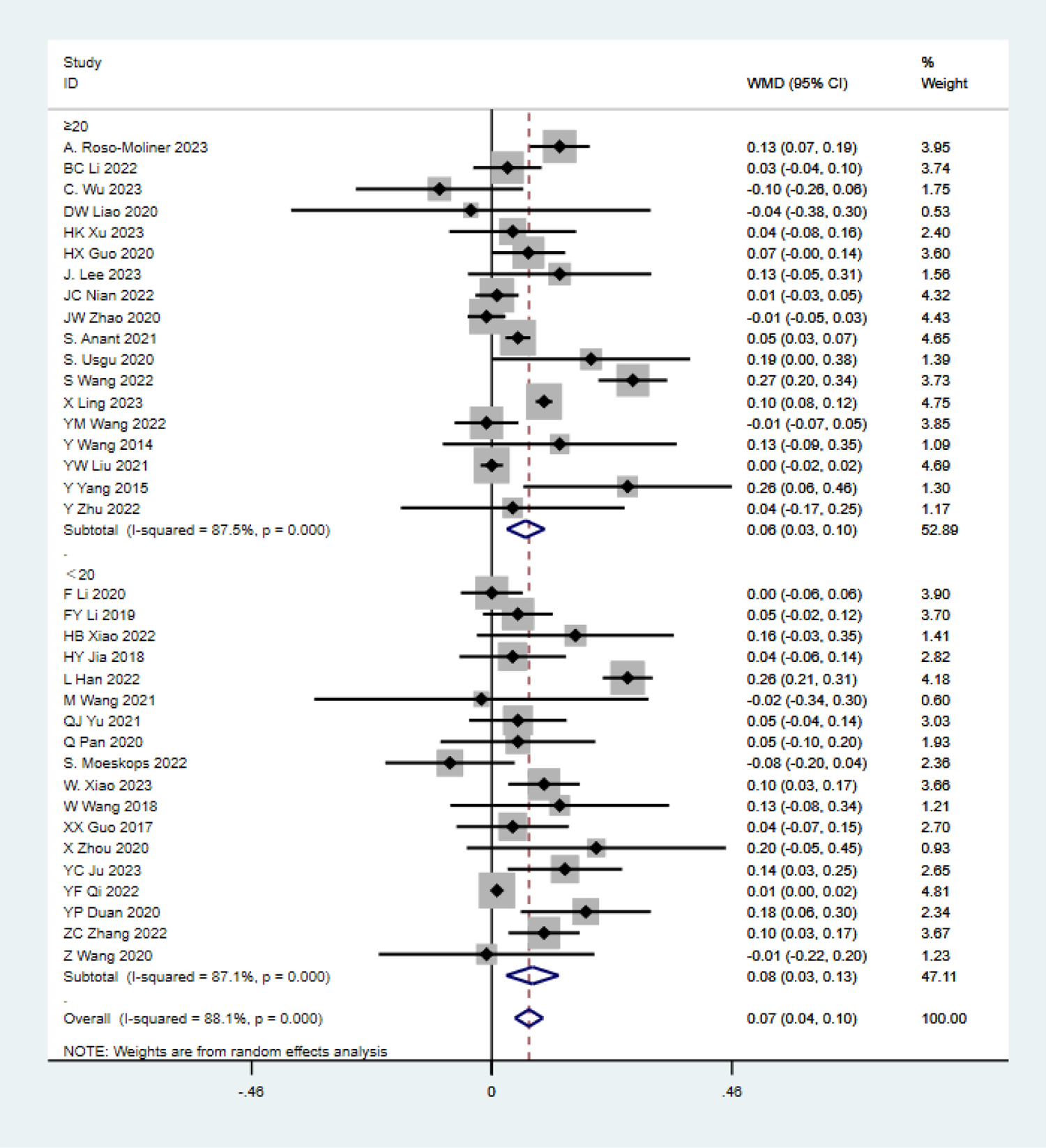
Single factor analysis on SLJ - Age.

**Figure 51:**
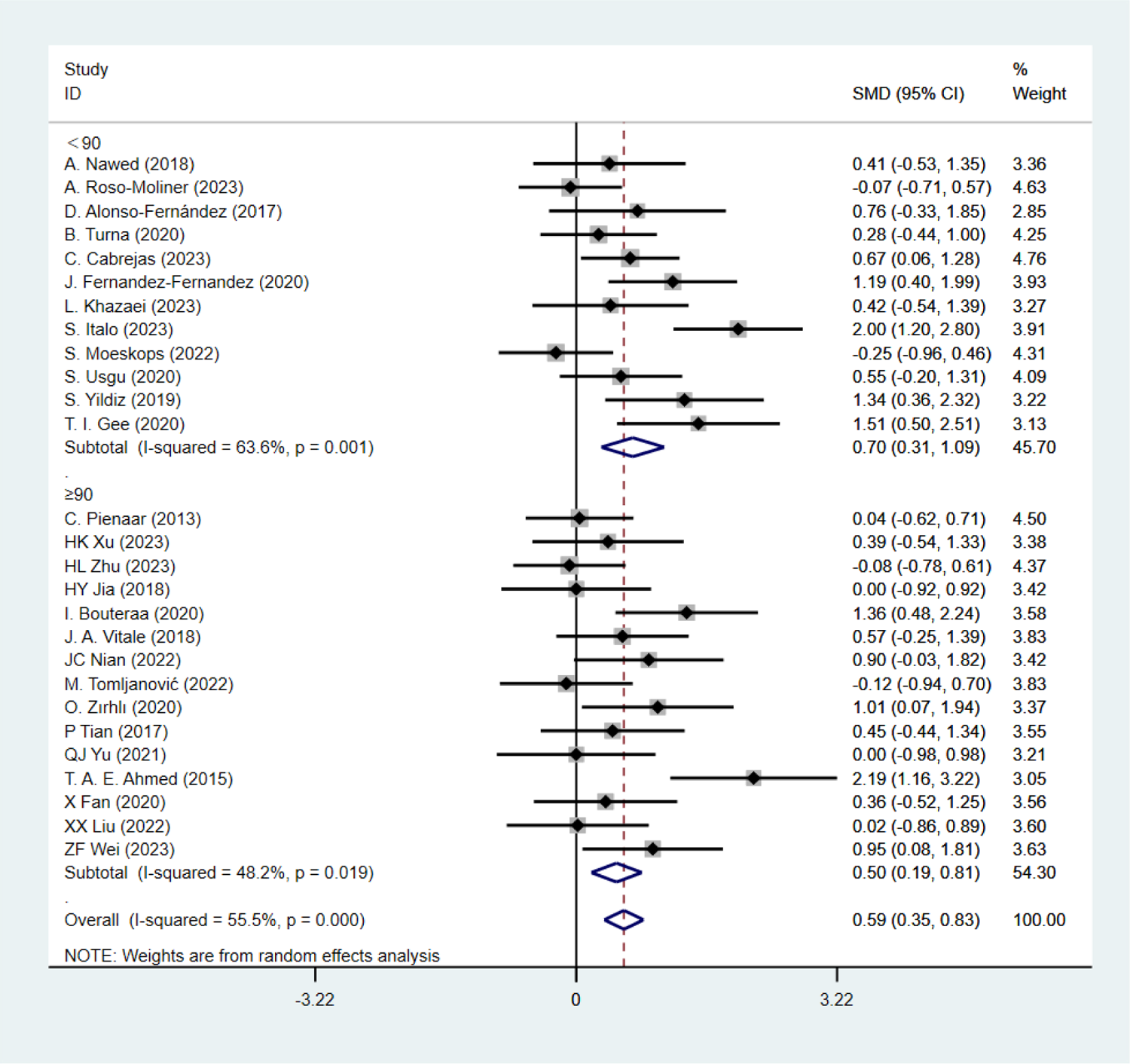
Single factor analysis on CMJ - Each session duration.

**Figure 52:**
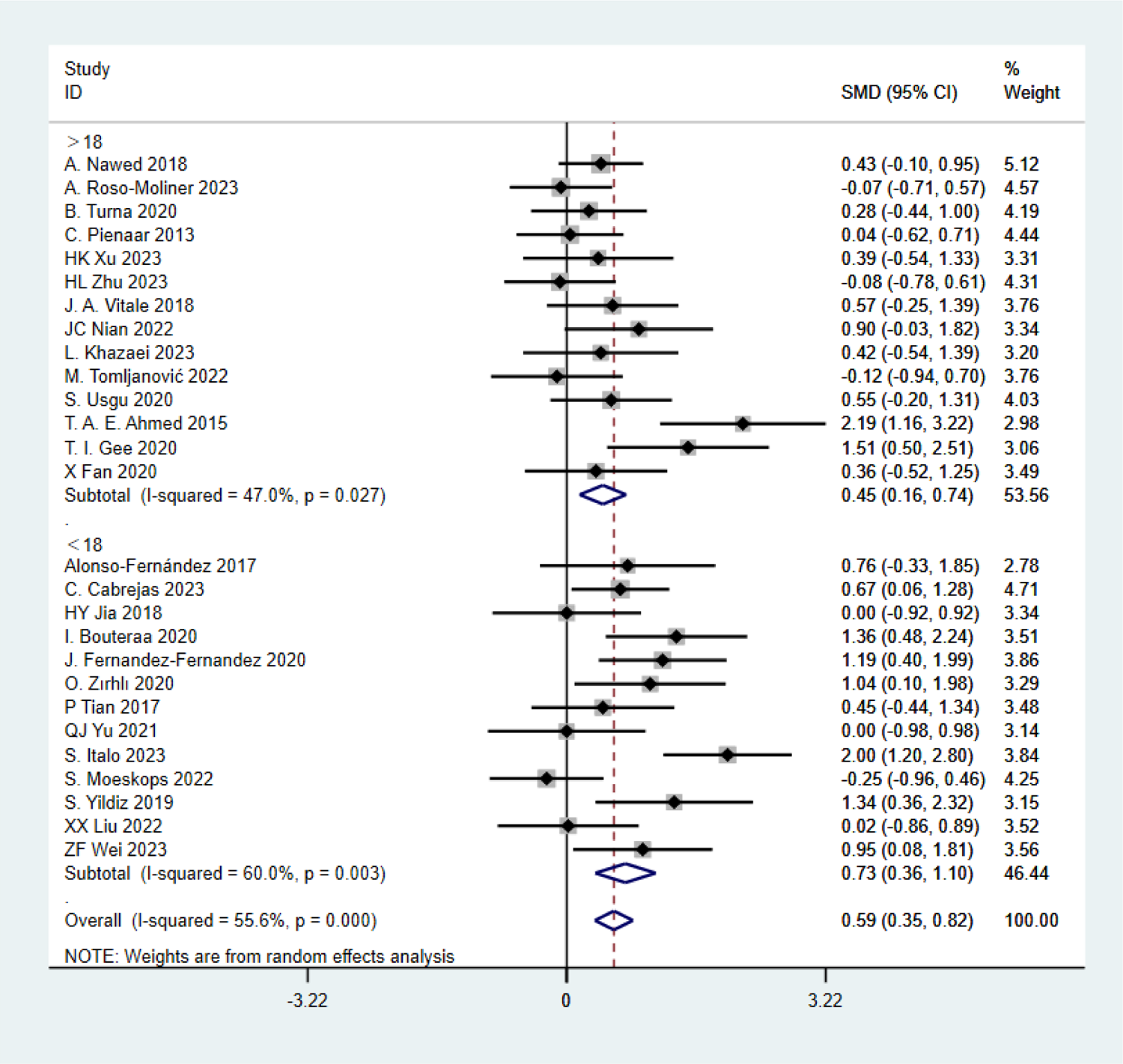
Single factor analysis on CMJ - Age.

**Figure 53:**
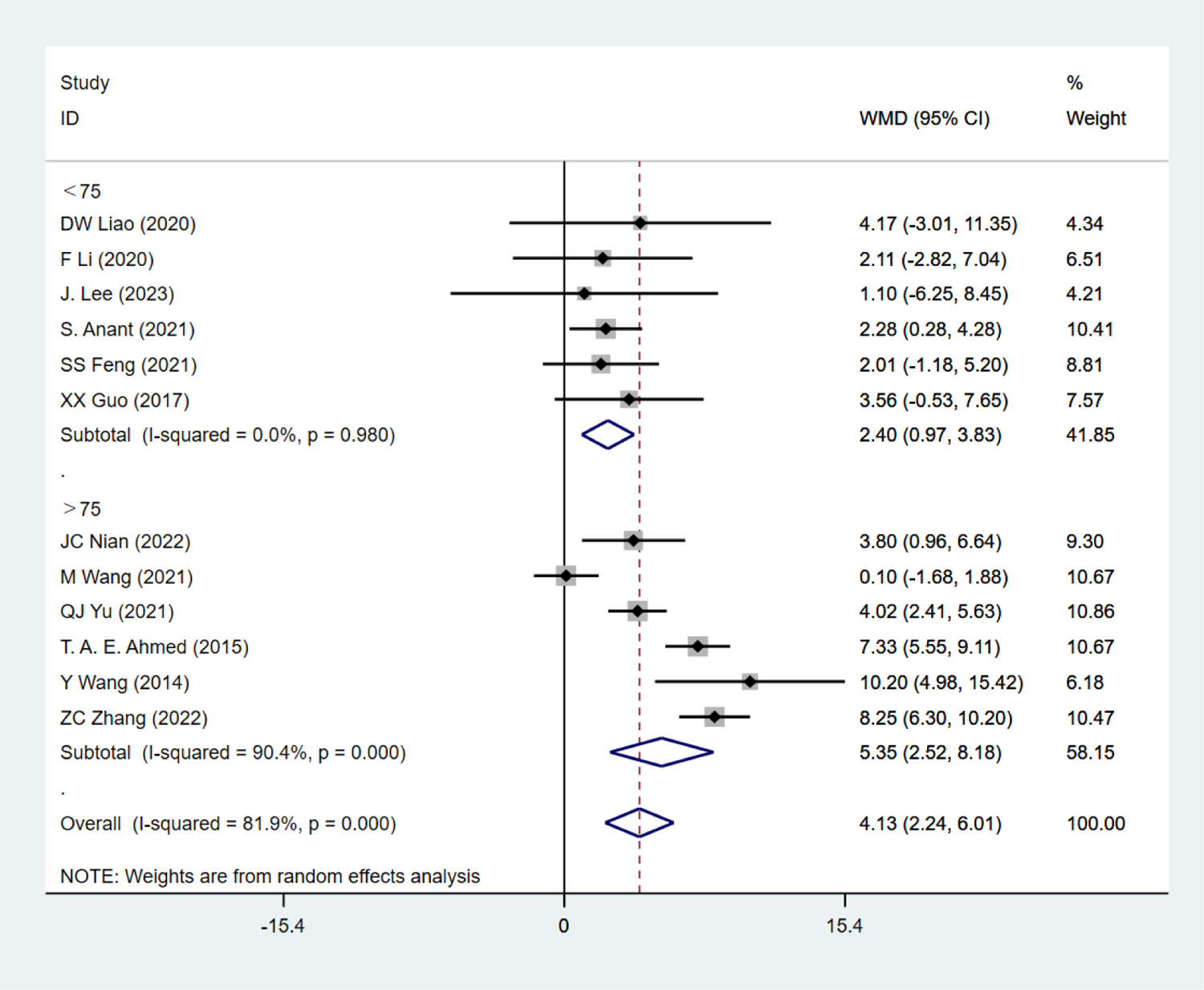
Single factor analysis on 1min SU - Each session duration.

**Figure 54:**
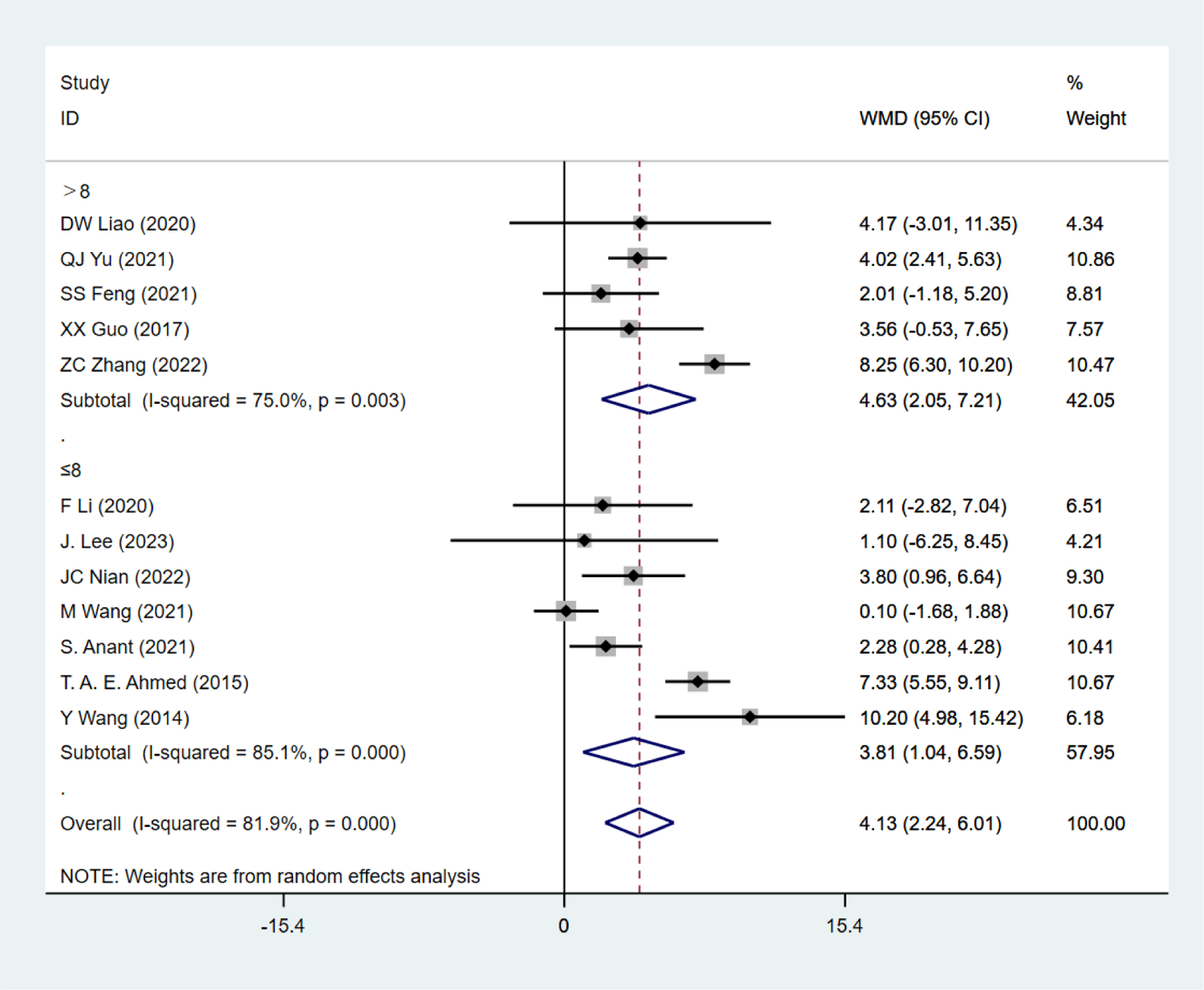
Single factor analysis on 1min SU - Total weeks

**Figure 55:**
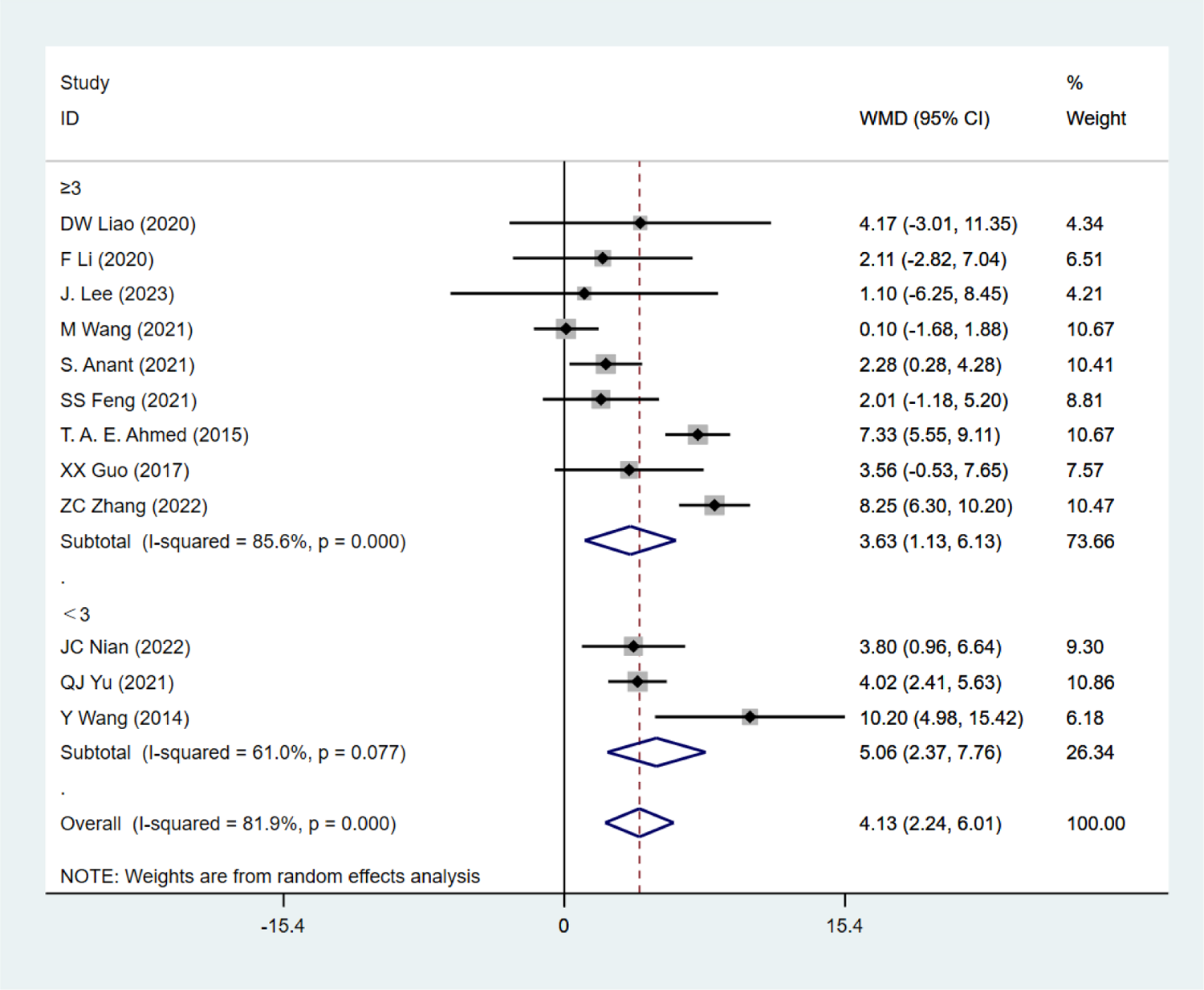
Single factor analysis on 1min SU - Frequency.

**Figure 56:**
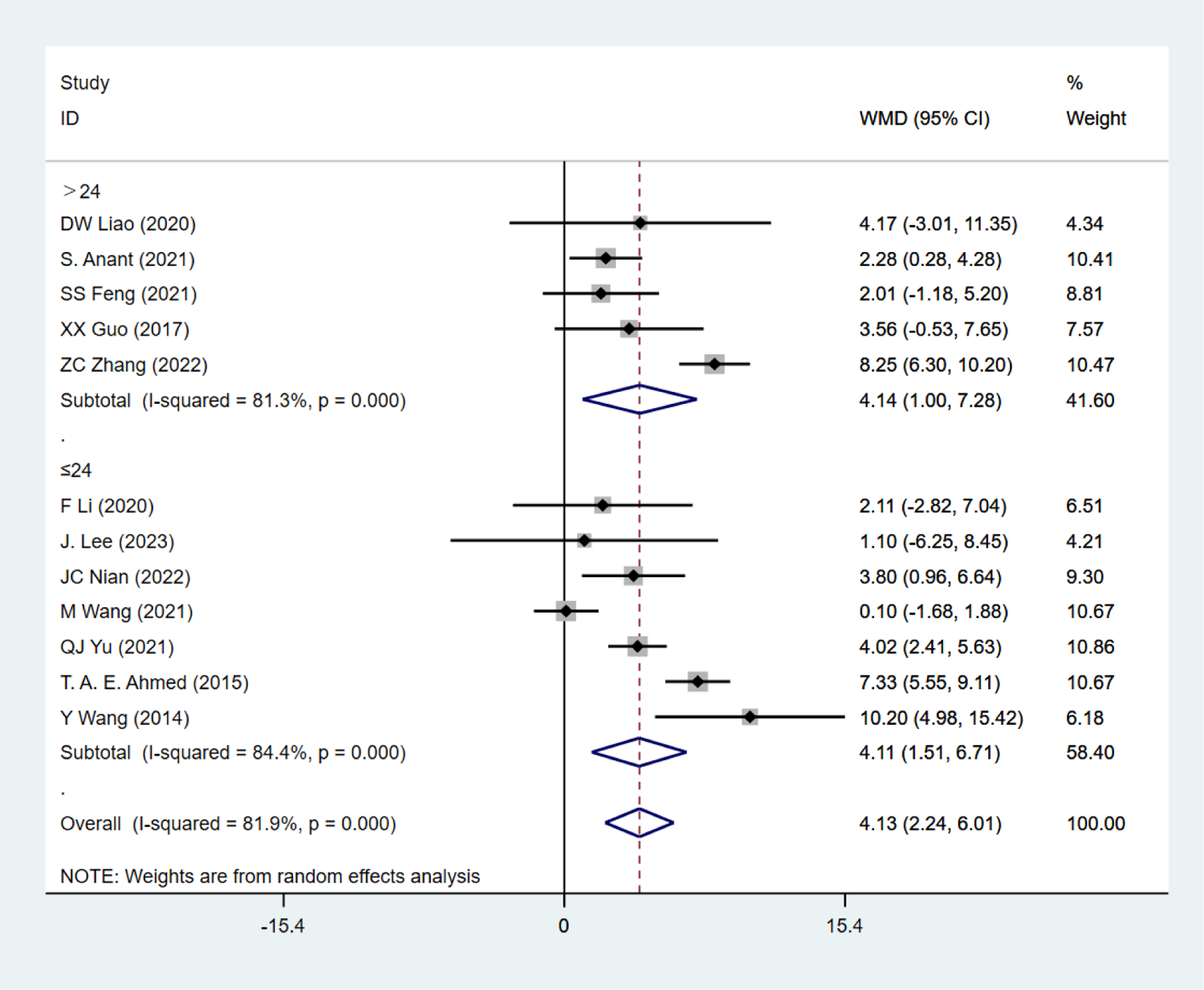
Single factor analysis on 1min SU - Total sessions.

**Figure 57:**
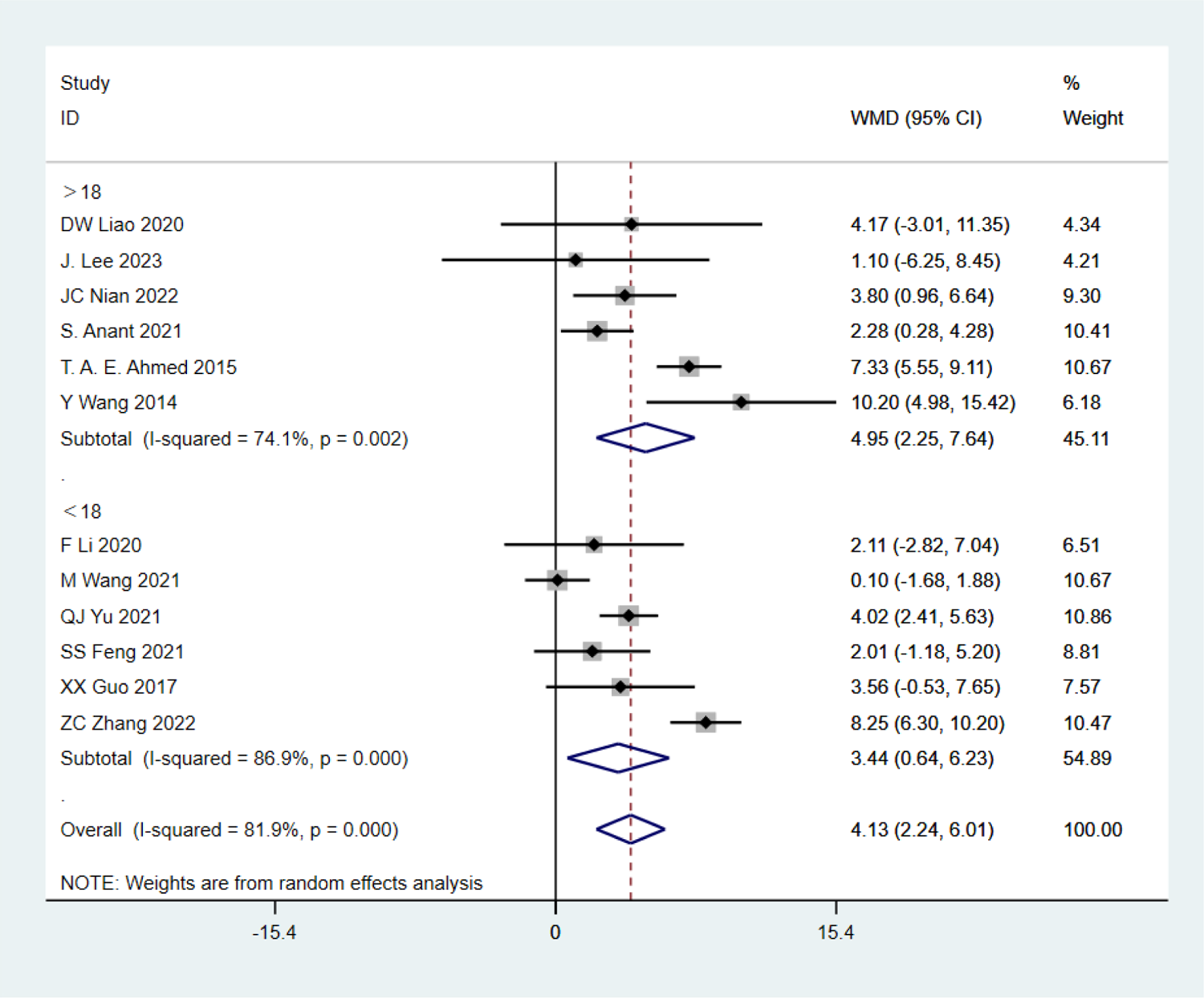
Single factor analysis on 1min SU - Age.

**Figure 58:**
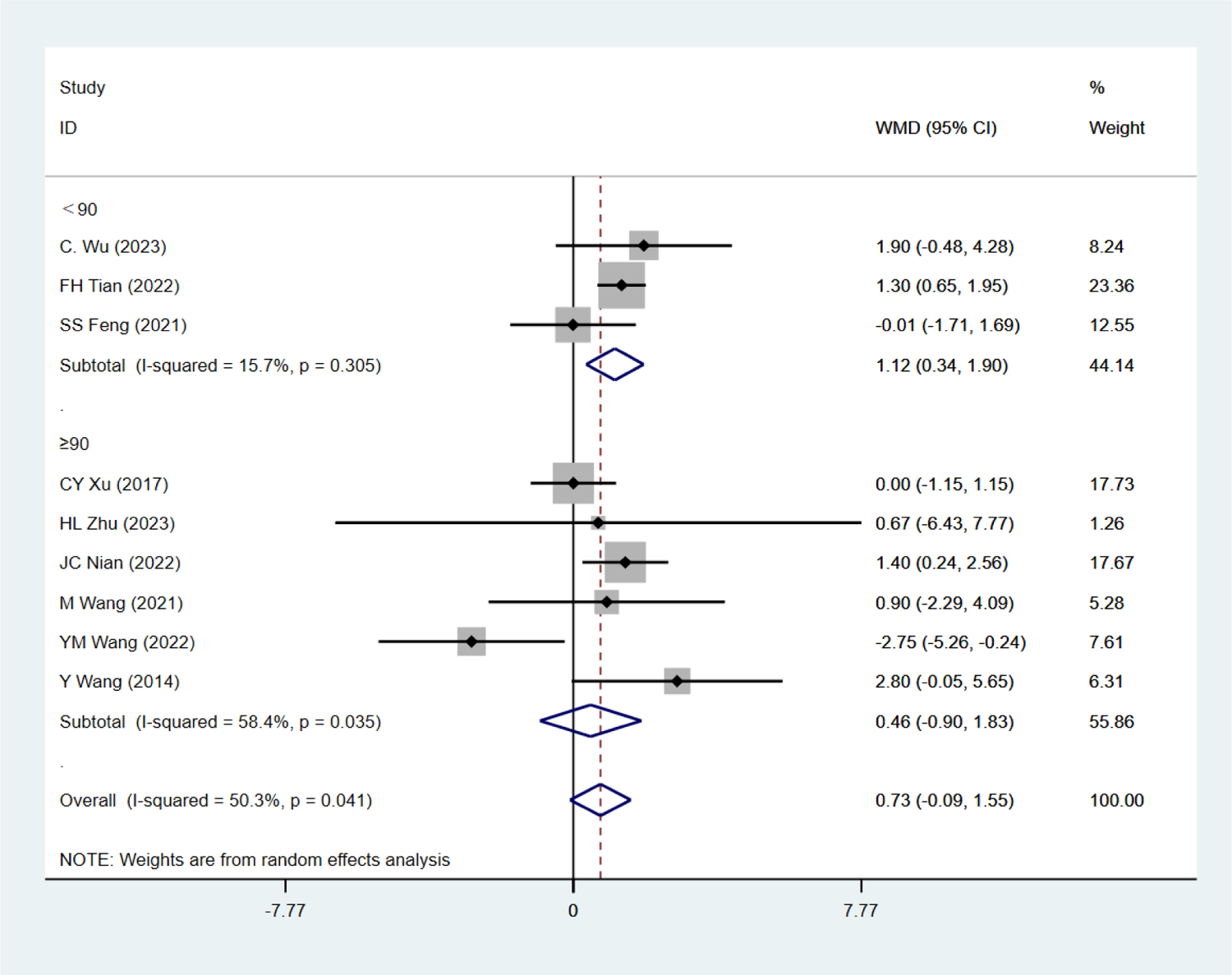
Single factor analysis on PLU - Each session duration.

**Figure 59:**
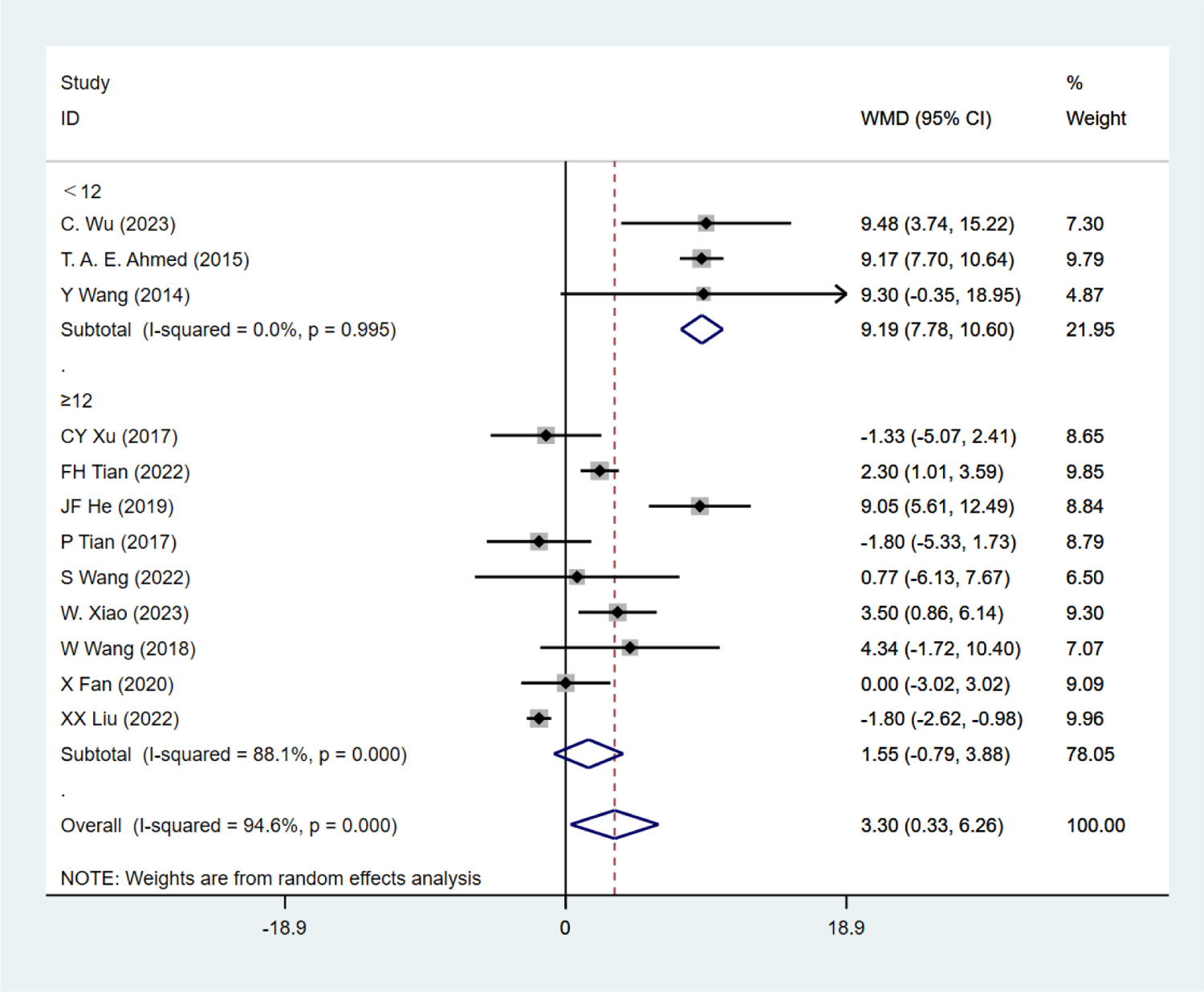
Single factor analysis on PSU - Total weeks.

**Figure 60:**
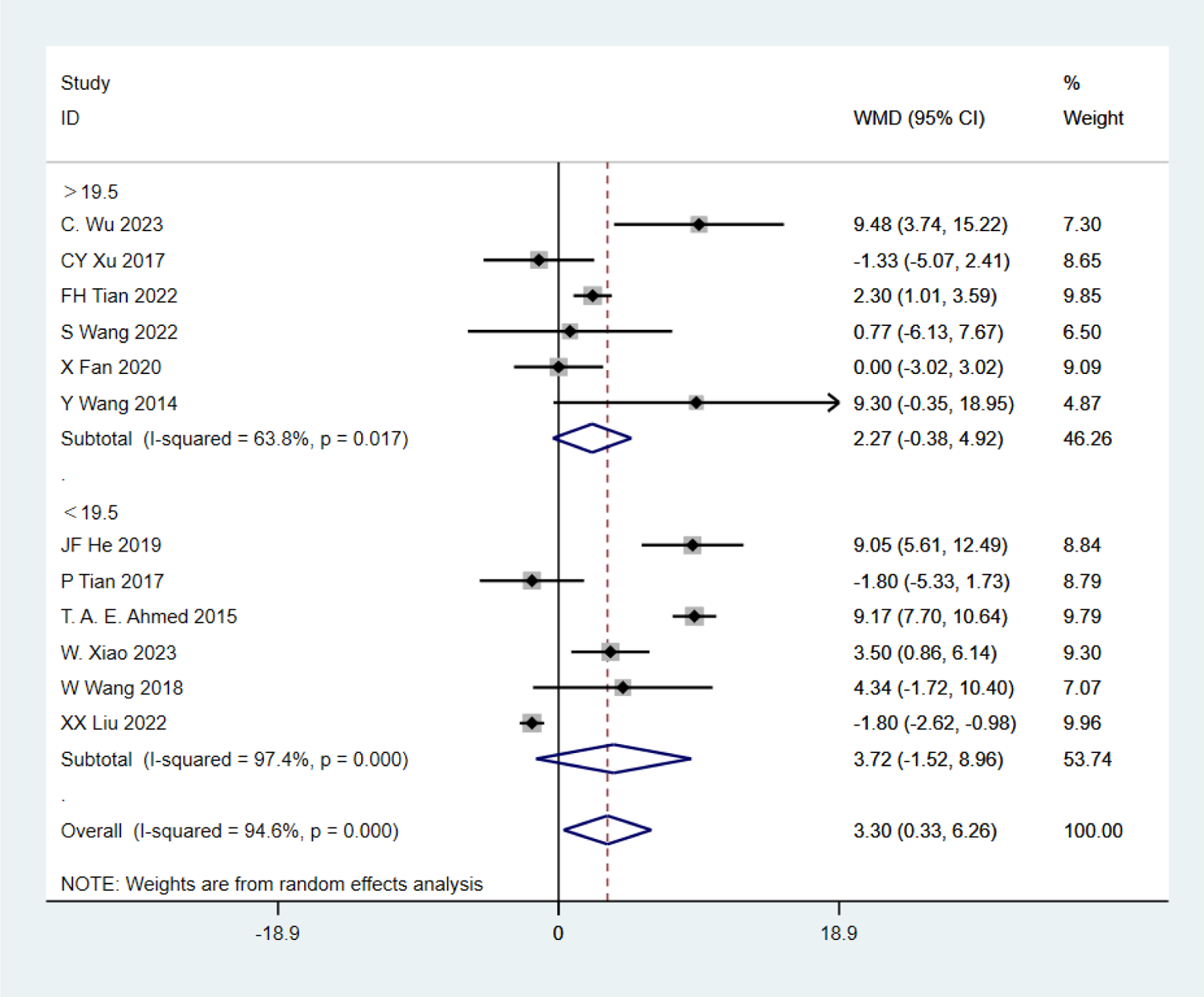
Single factor analysis on PSU - age.

**Figure 61:**
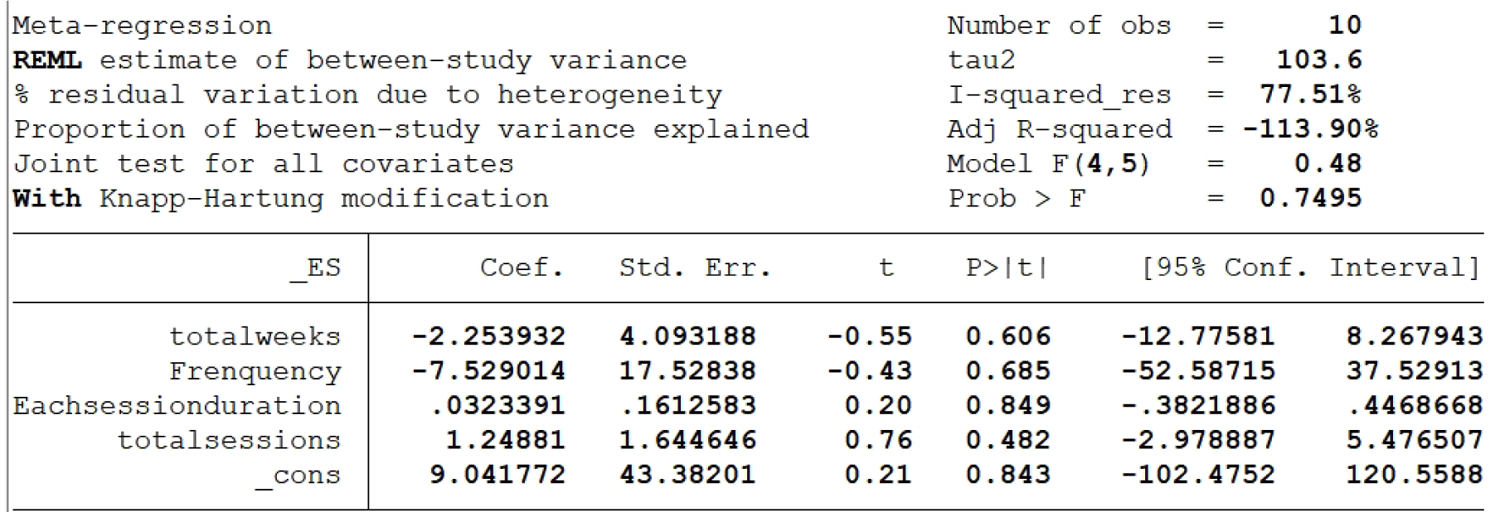
Meta Regression - 1RM HS/DS.

**Figure 62:**
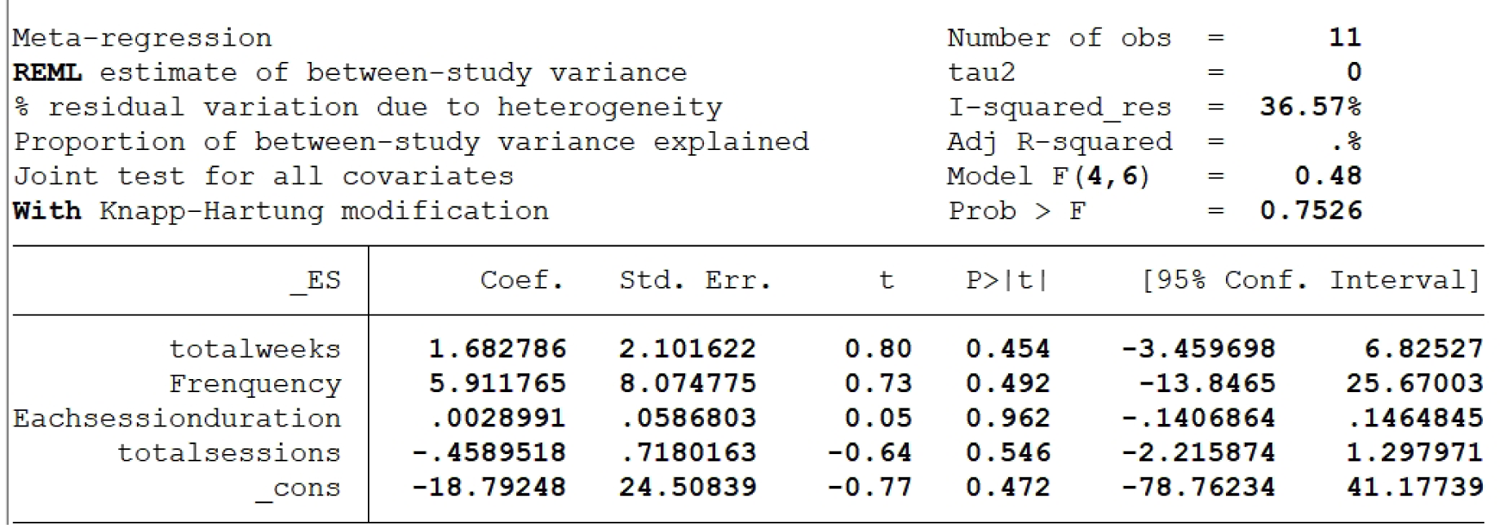
Meta Regression - 1RM BP.

**Figure 63:**
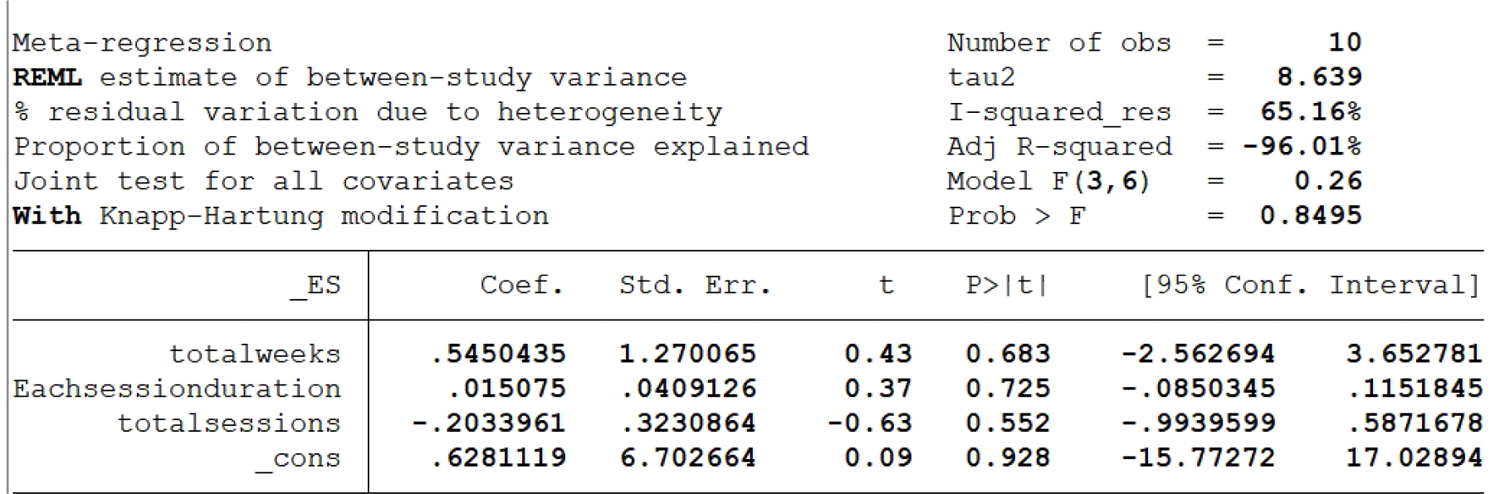
Meta Regression - 1RM GS.

**Figure 64:**
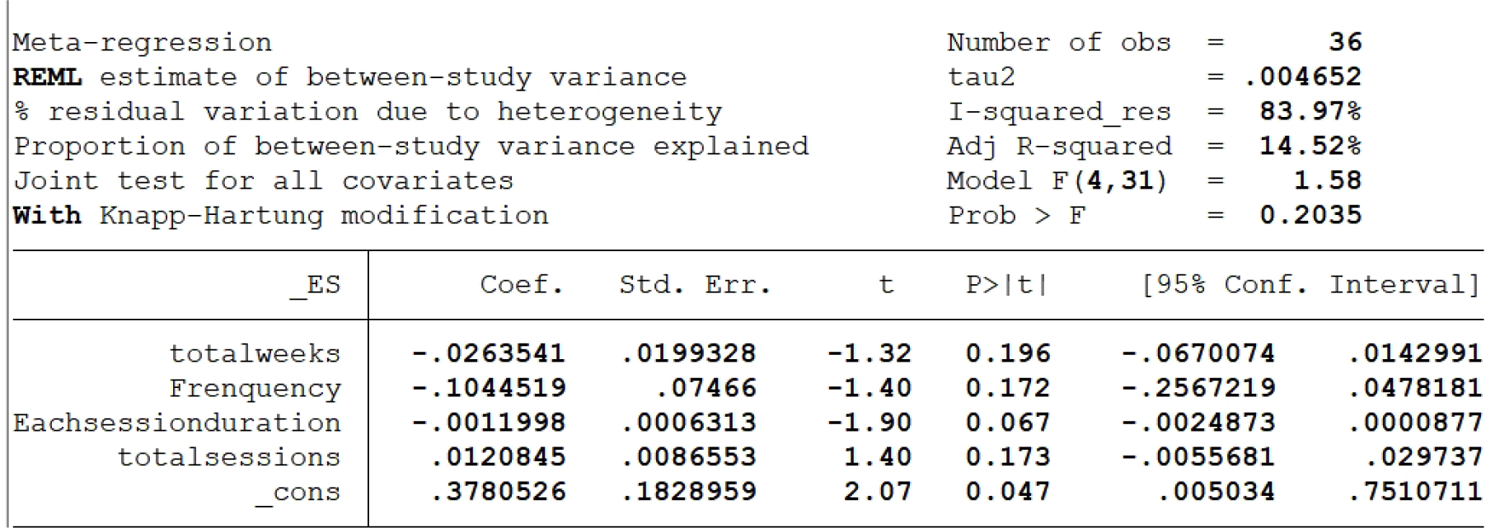
Meta Regression - SLJ.

**Figure 65:**
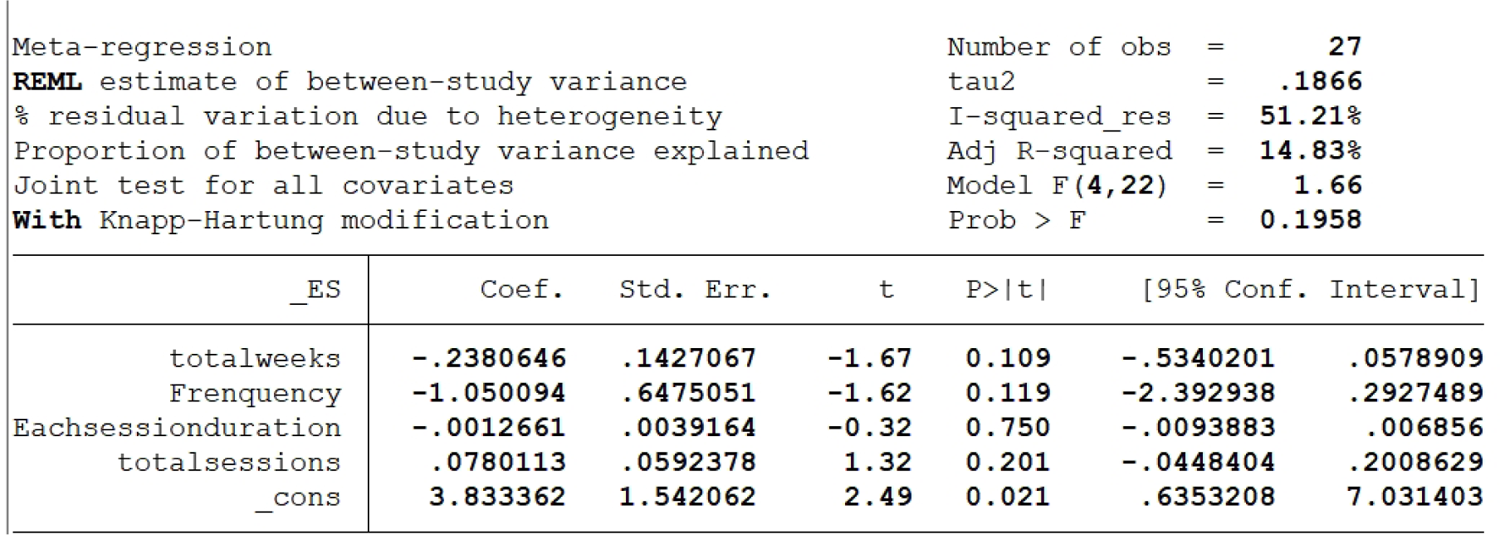
Meta Regression - CMJ.

**Figure 66:**
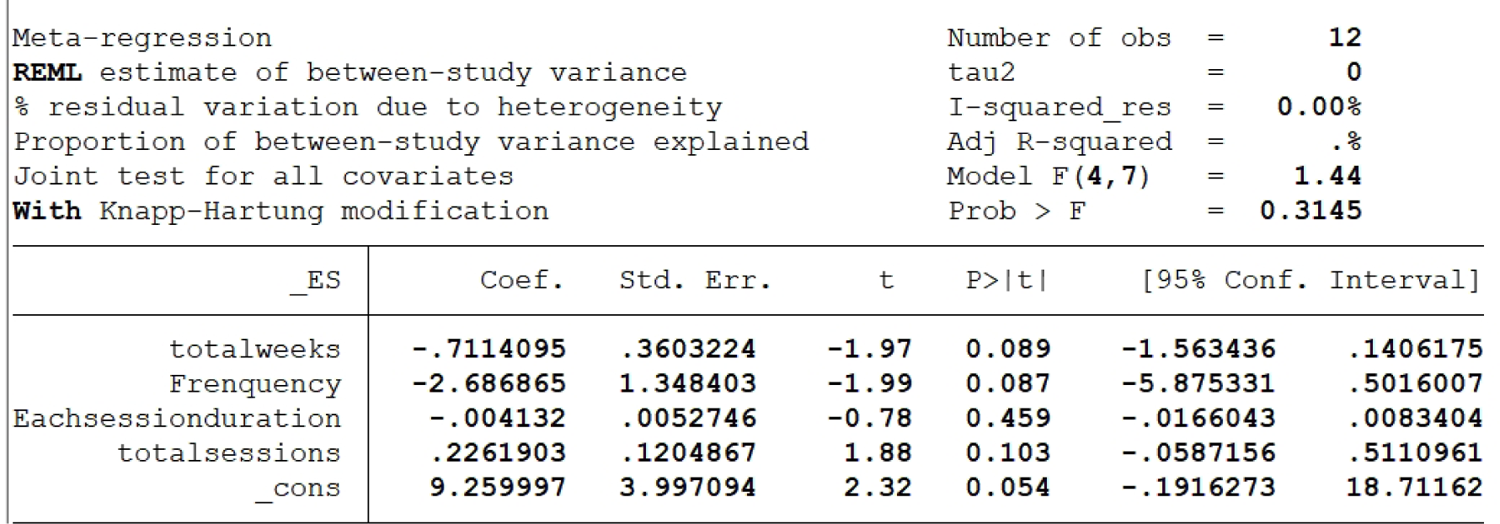
Meta Regression - TMBP.

**Figure 67:**
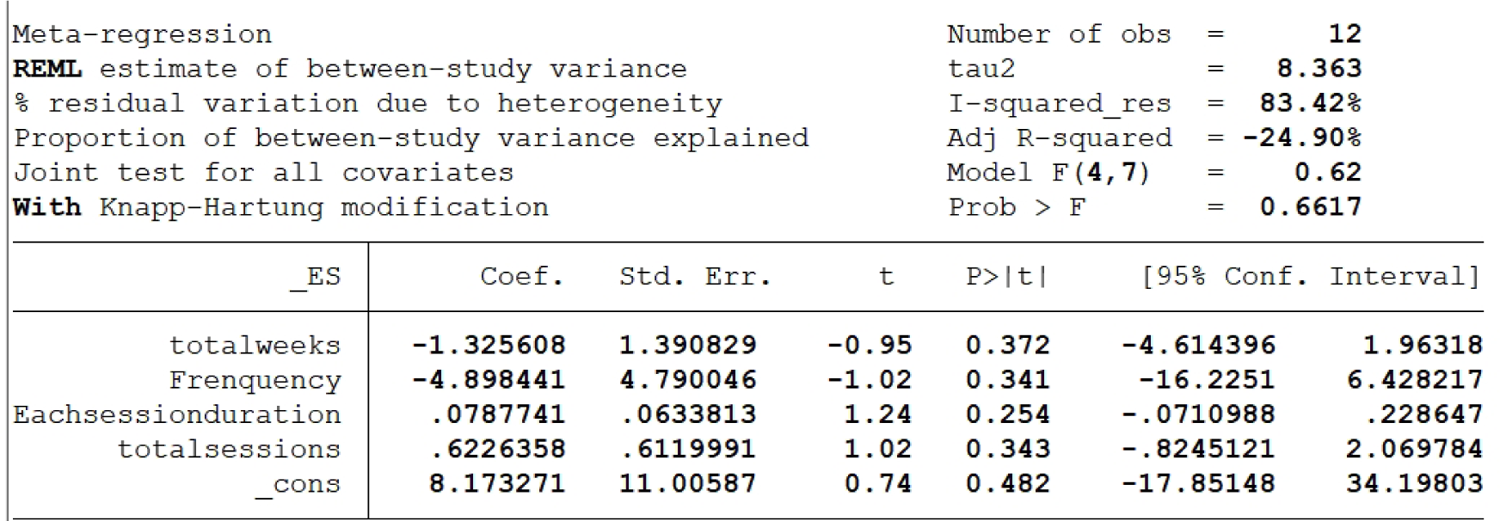
Meta Regression - 1min SU.

**Figure 68:**
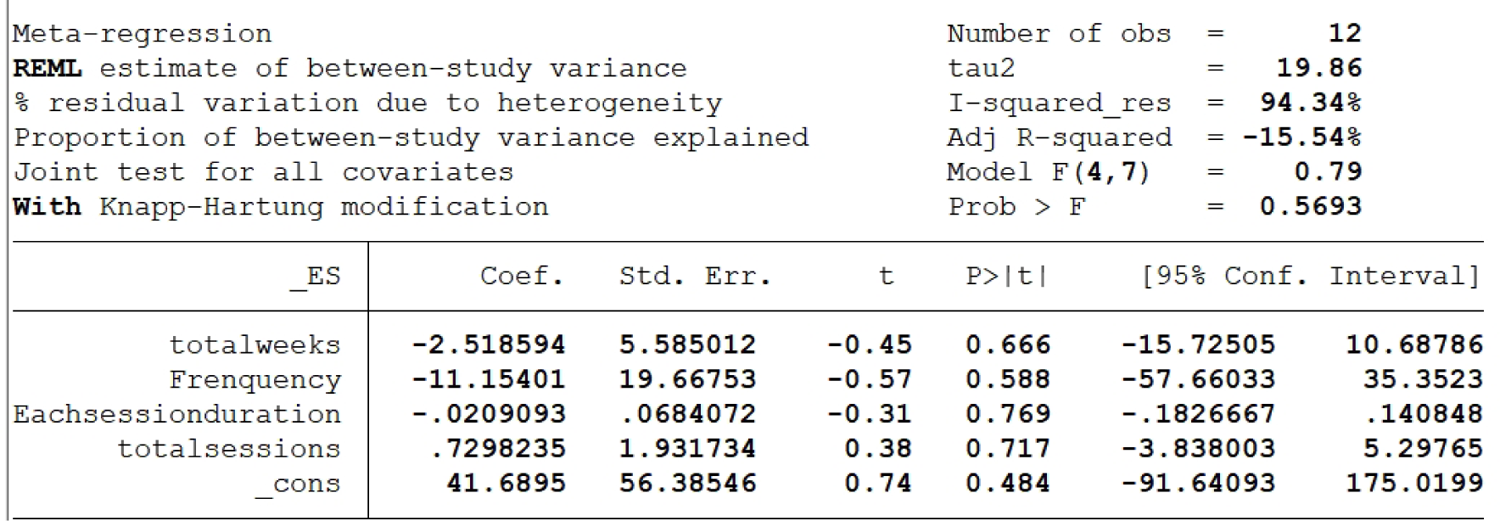
Meta Regression - PSU.

## Supplementary Figure

**Figure 1.**
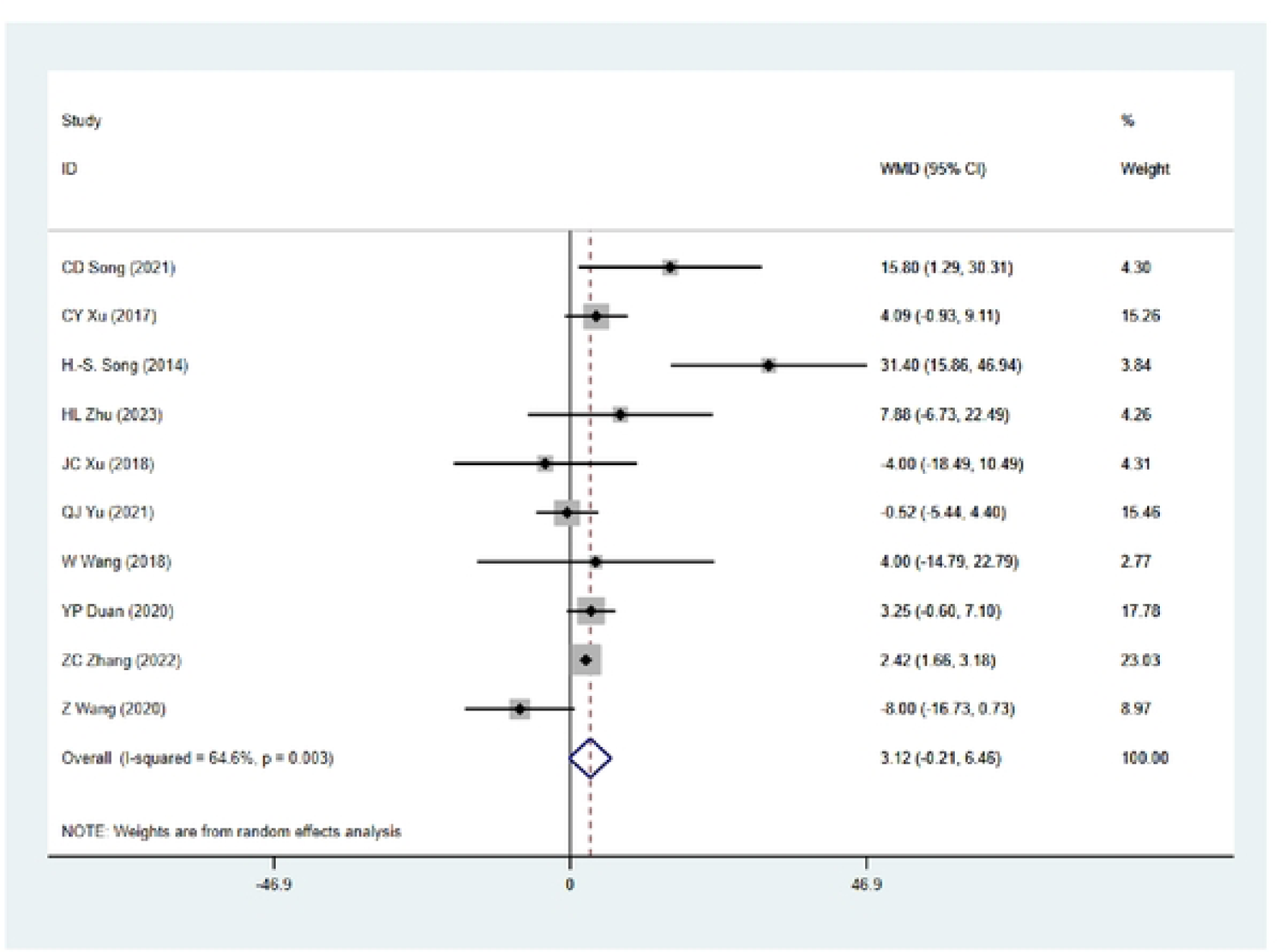
Forest plot 2 The effects of FT vs. TRT on lower limb maximal strength (1RM squat/half squat).

**Figure 2.**
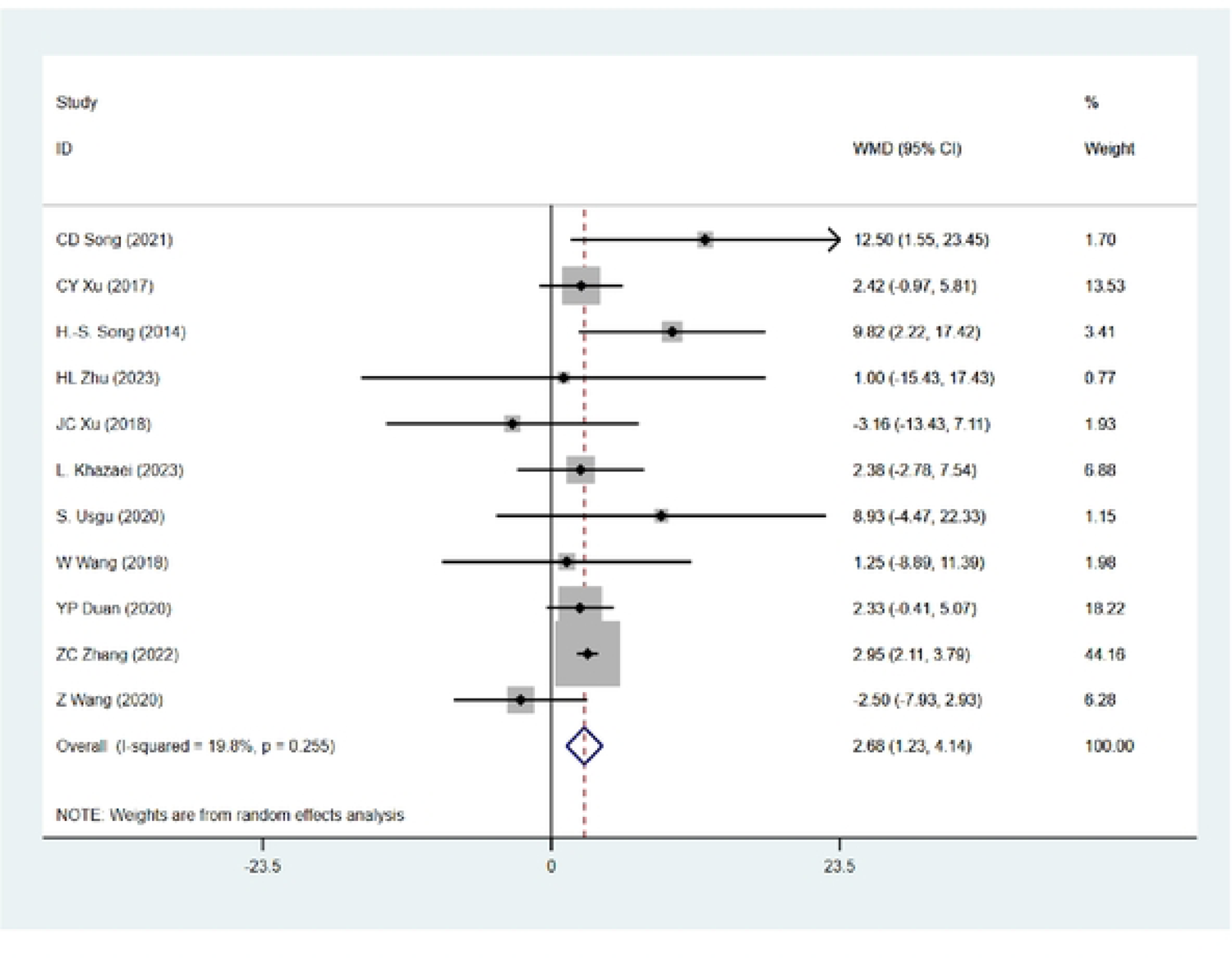
Forest plot 2 The effects of FT vs. TRT on upper limb maximal strength (1RM bench press).

**Figure 3.**
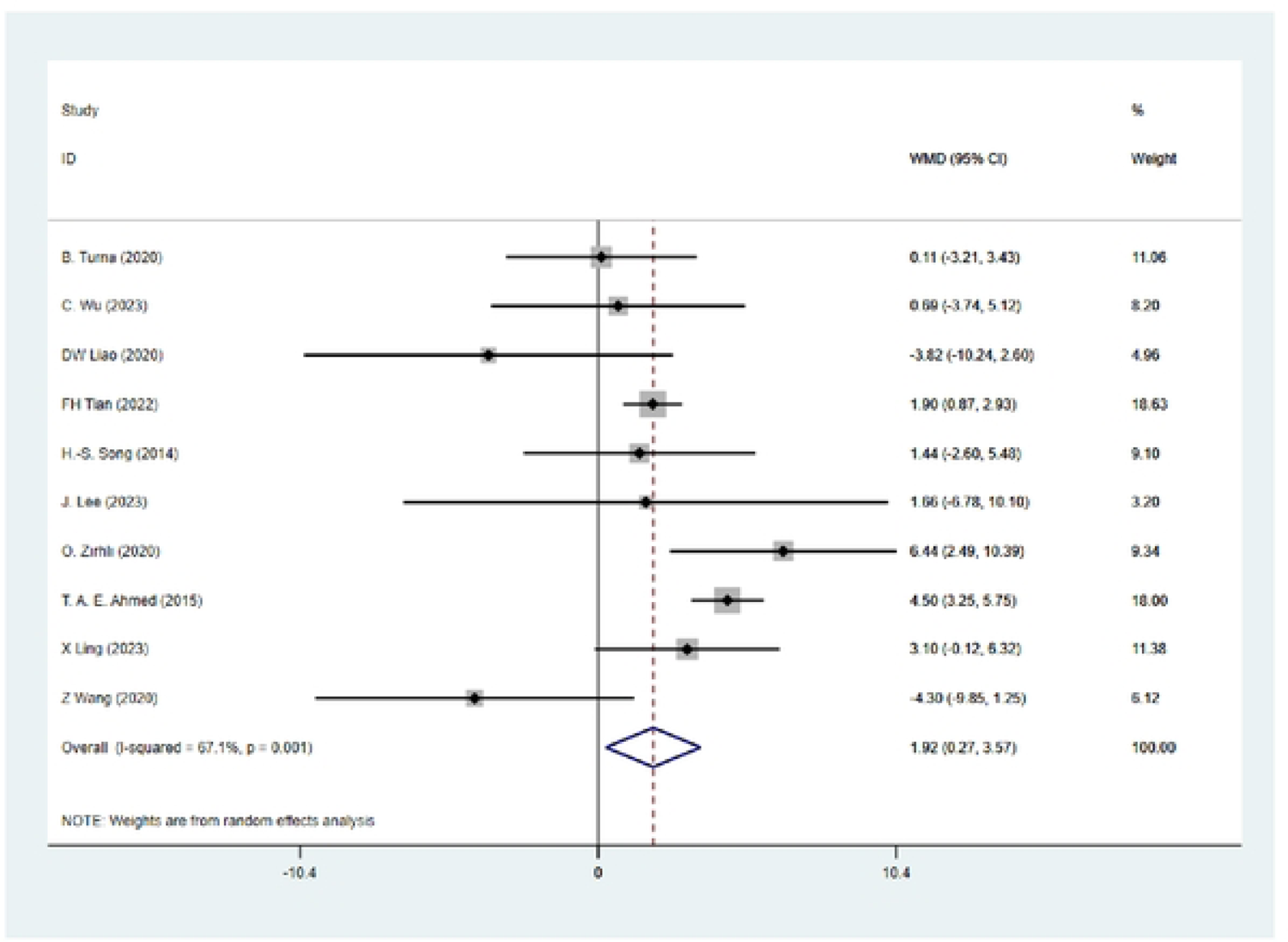
Forest plot 3 The effects of FT vs. TRT on grip maximal strength (1RM grip strength).

**Figure 4.**
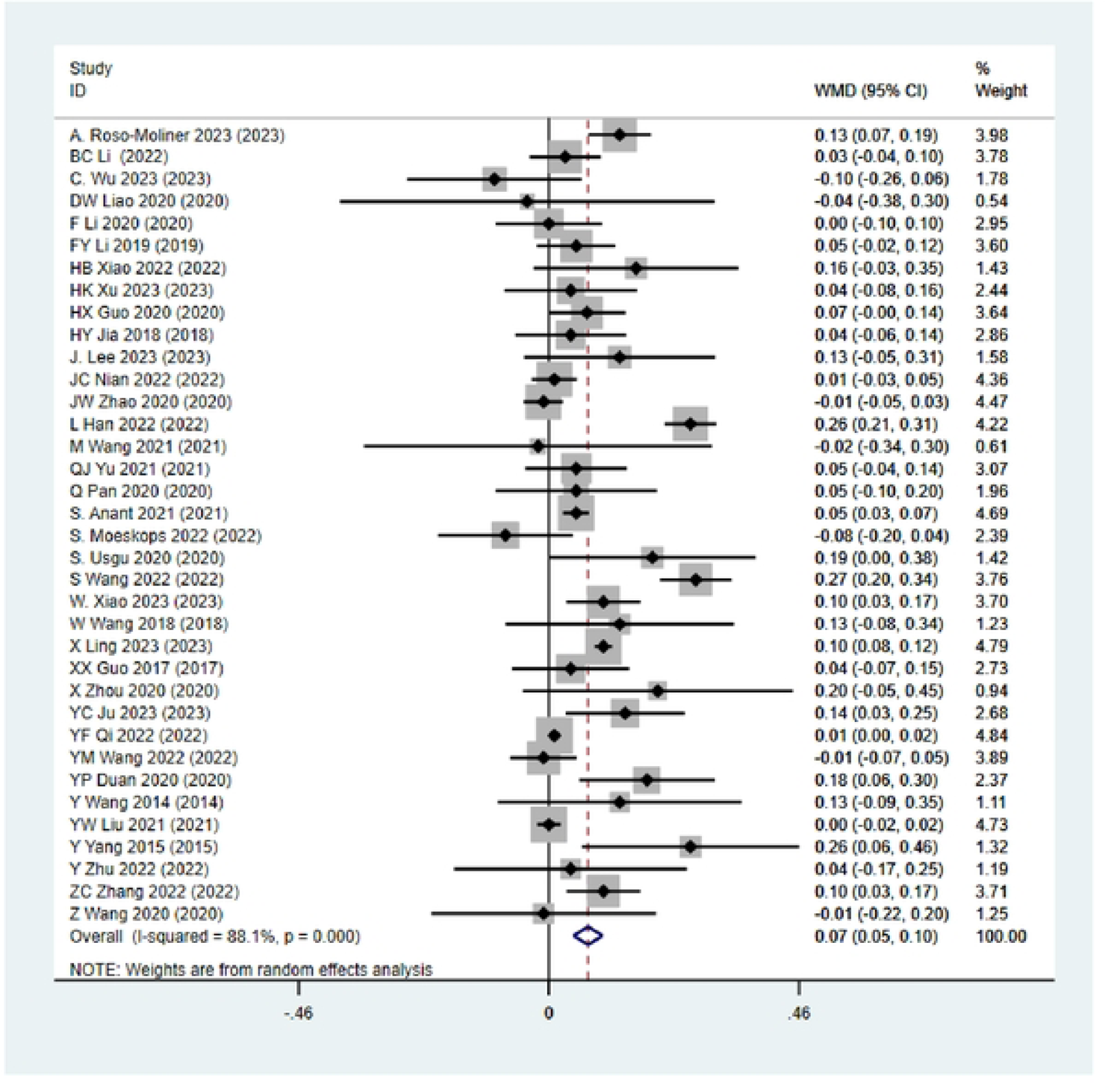
Forest plot 4 The effects of FT vs. TRT on power (standing long jump).

**Figure 5.**
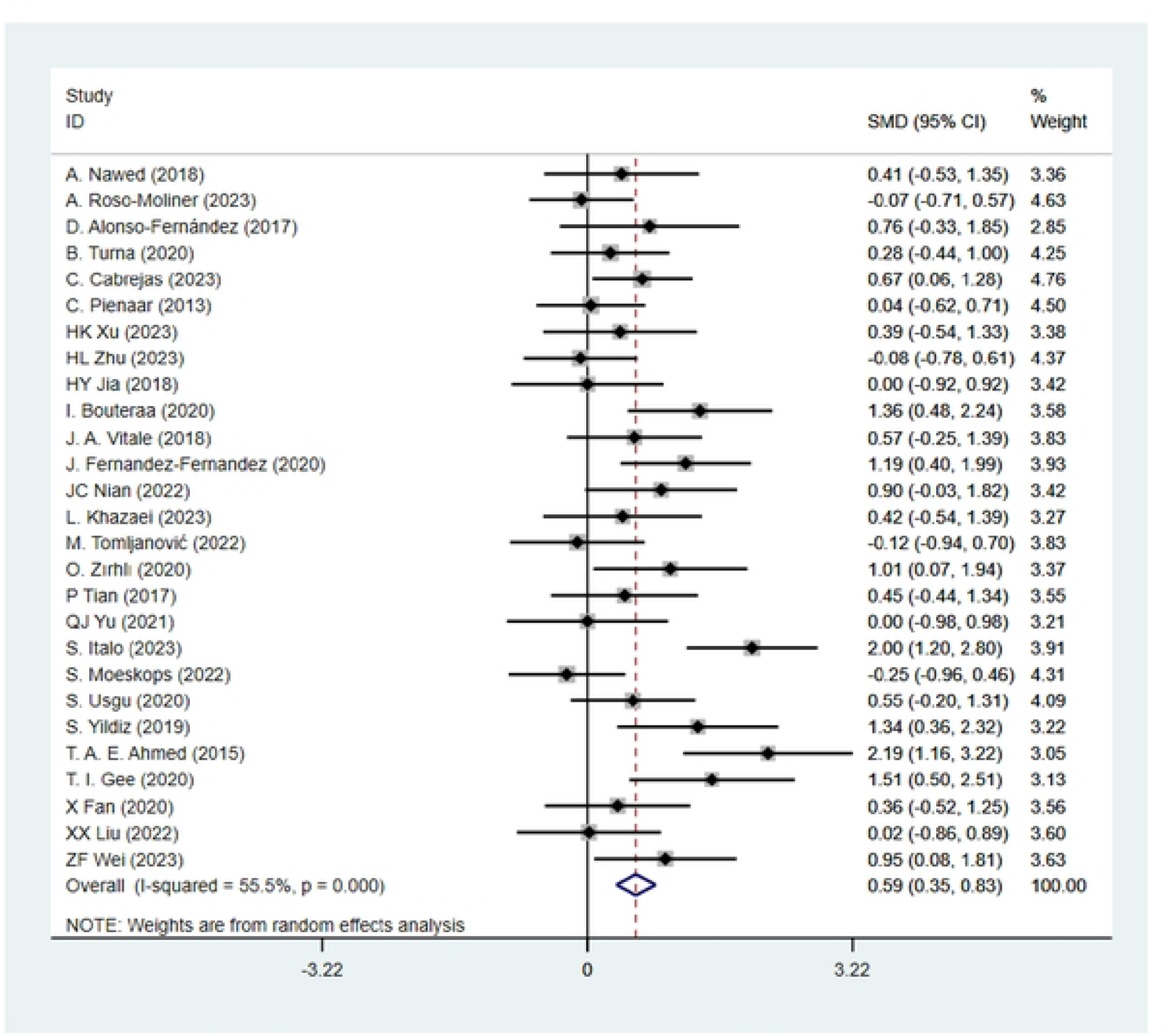
Forest plot5 The effects of FT vs. TRT on power (countermovement jump).

**Figure 6.**
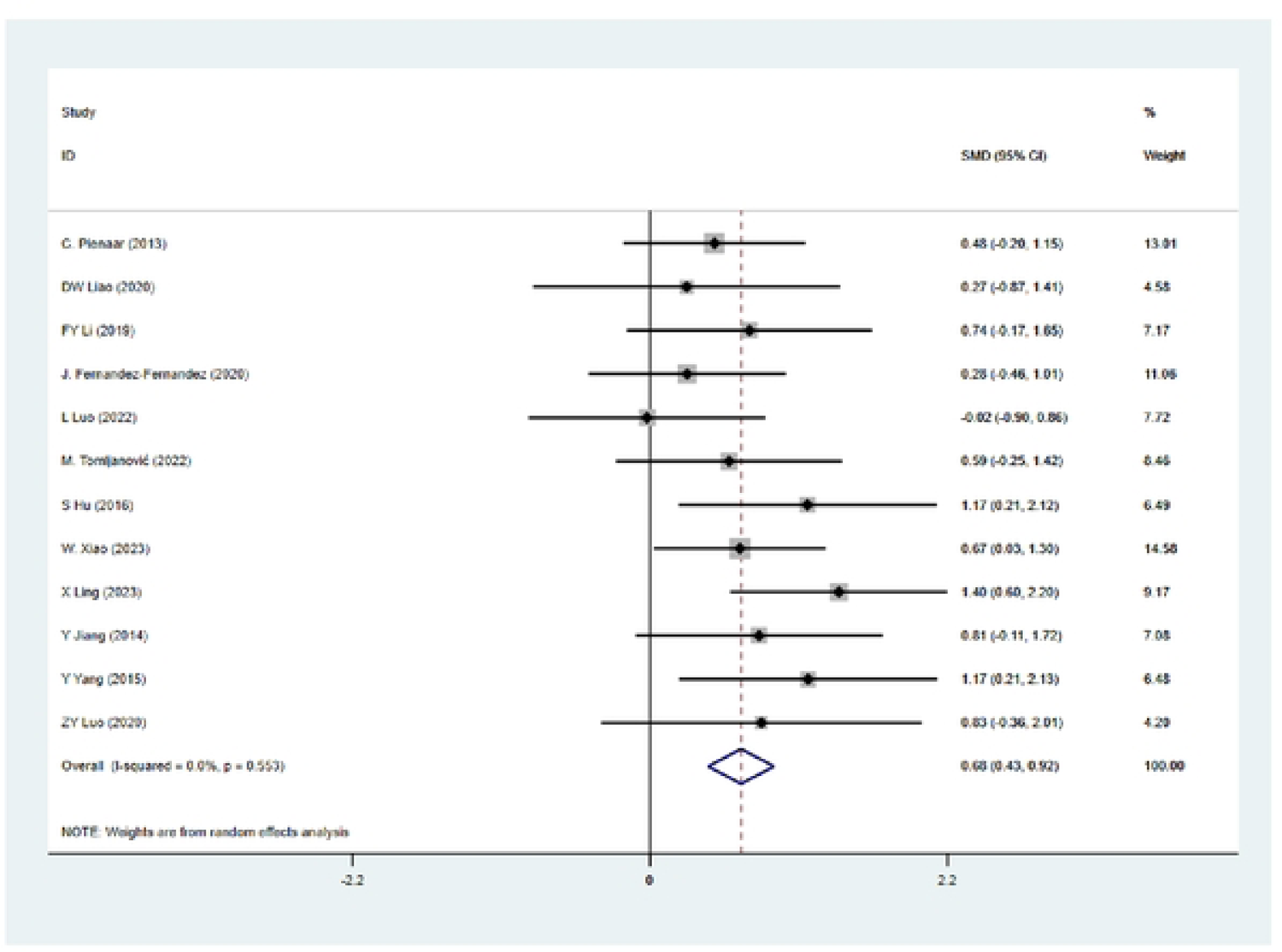
Forest plot 6 The effects of FT vs. TRT on power (throw medicine ball in place).

**Figure 7.**
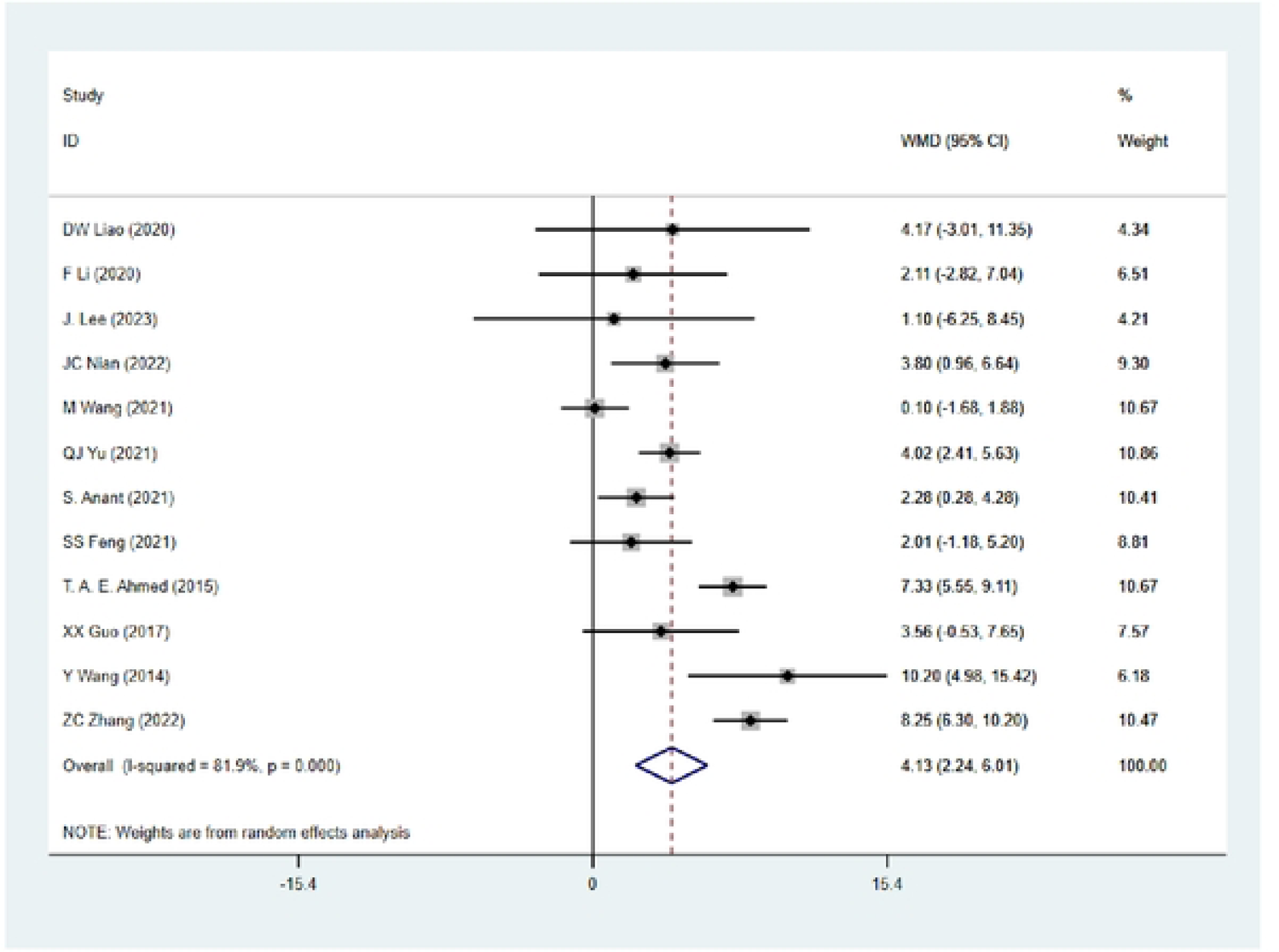
Forest plot 7 The effects of FT vs. TRT on muscle endurance (limit-ups).

**Figure 8.**
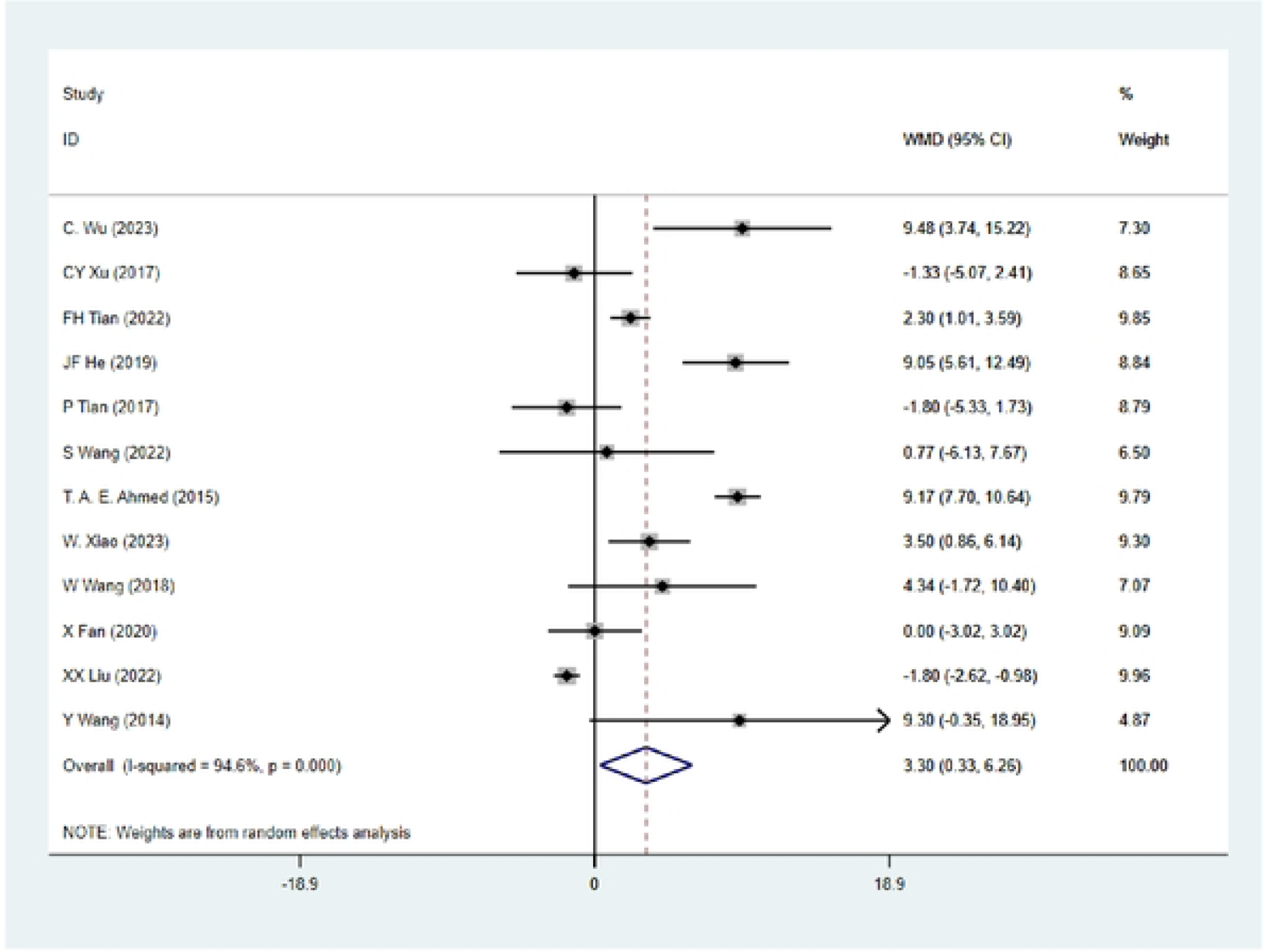
Forest plot 8 The effects of FT vs. TRT on muscle endurance (push-ups).

**Figure 9.**
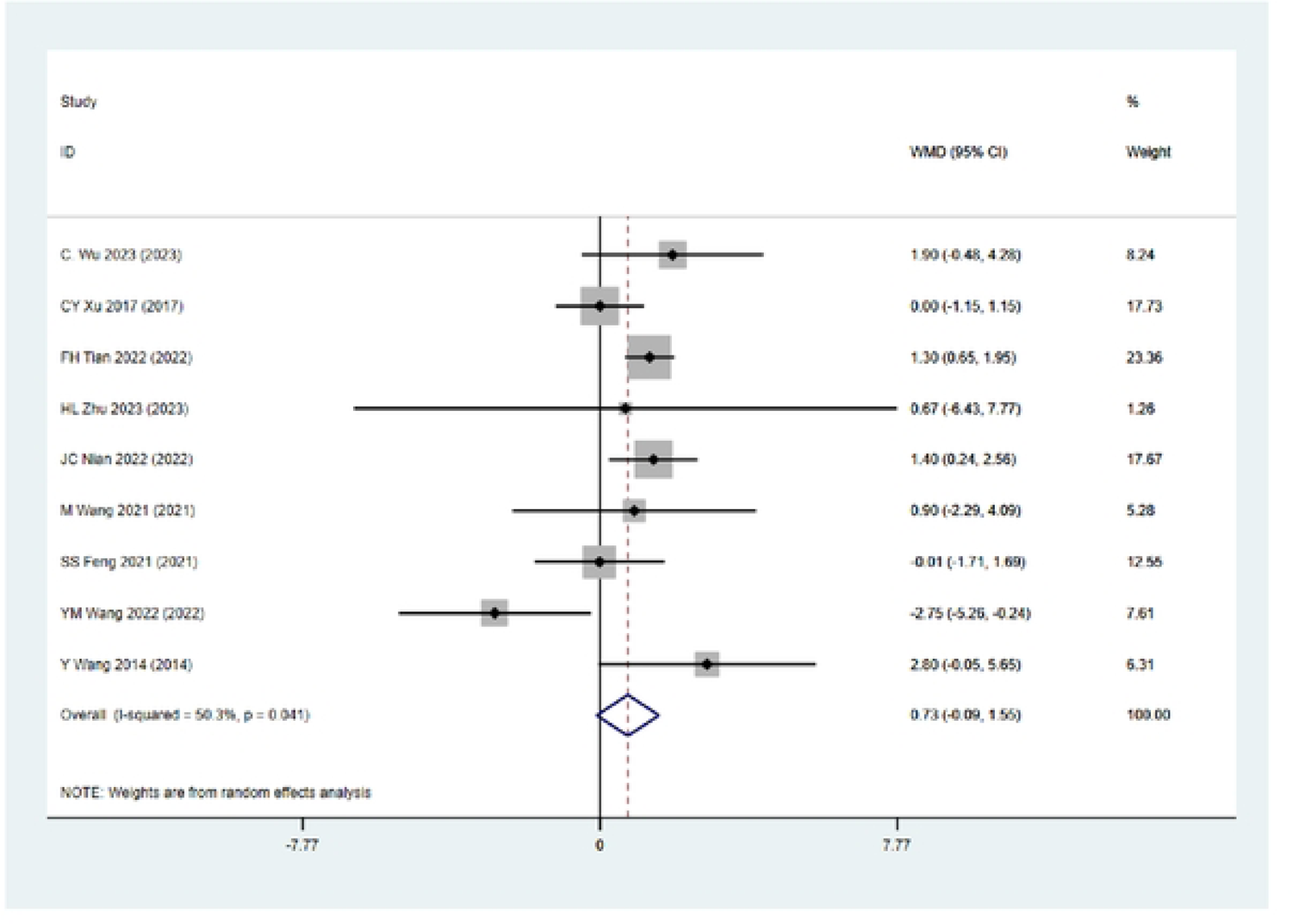
Forest plot 9 The effects of FT vs. TRT on muscle endurance (pull-ups).

**Figure 10:**
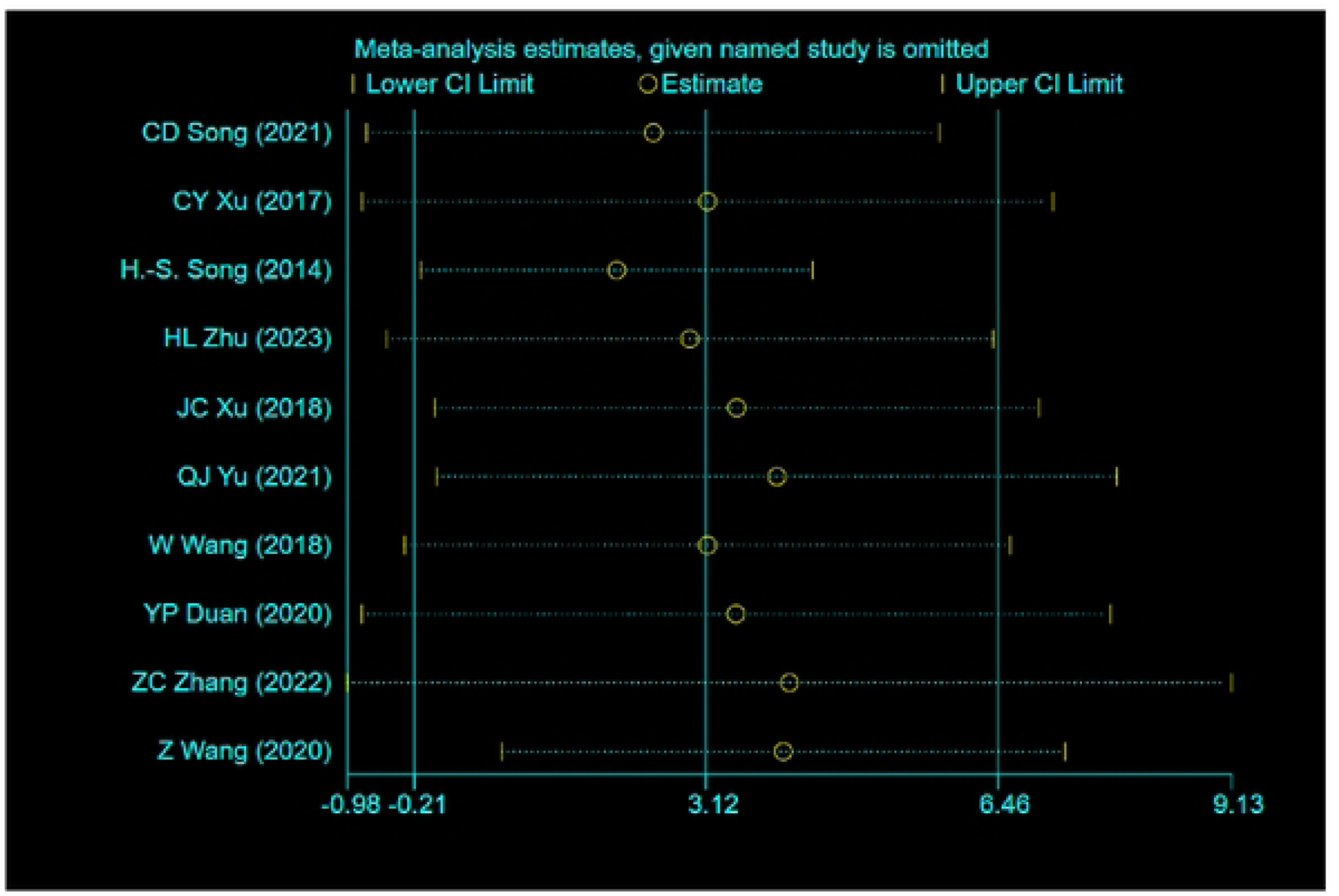
Sensitivity analysis 1 The effects of FT vs. TRT on lower limb maximal strength (1RM squat/half squat).

**Figure 11:**
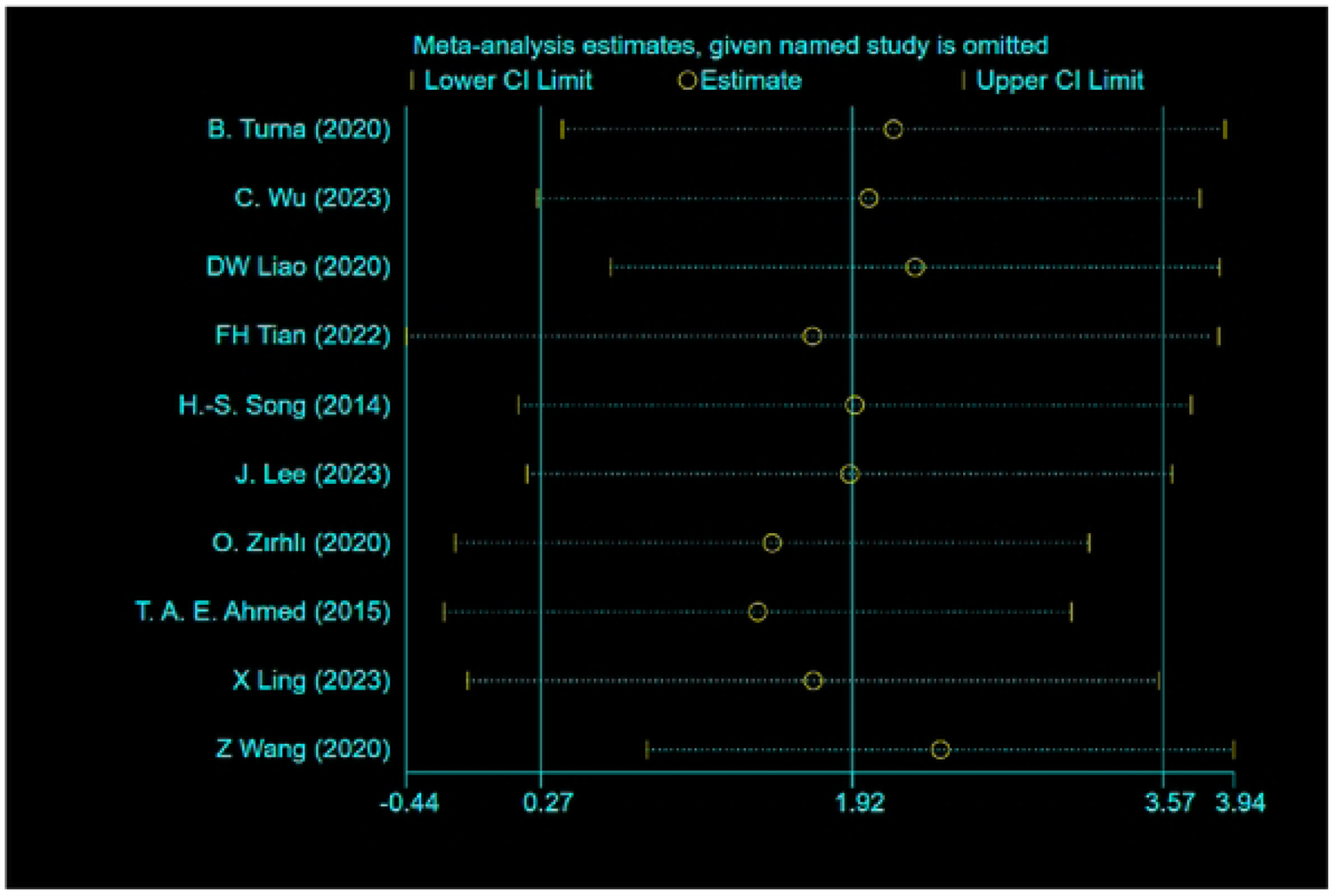
Sensitivity analysis 2 The effects of FT vs. TRT on grip strength (1RM grip strength).

**Figure 12:**
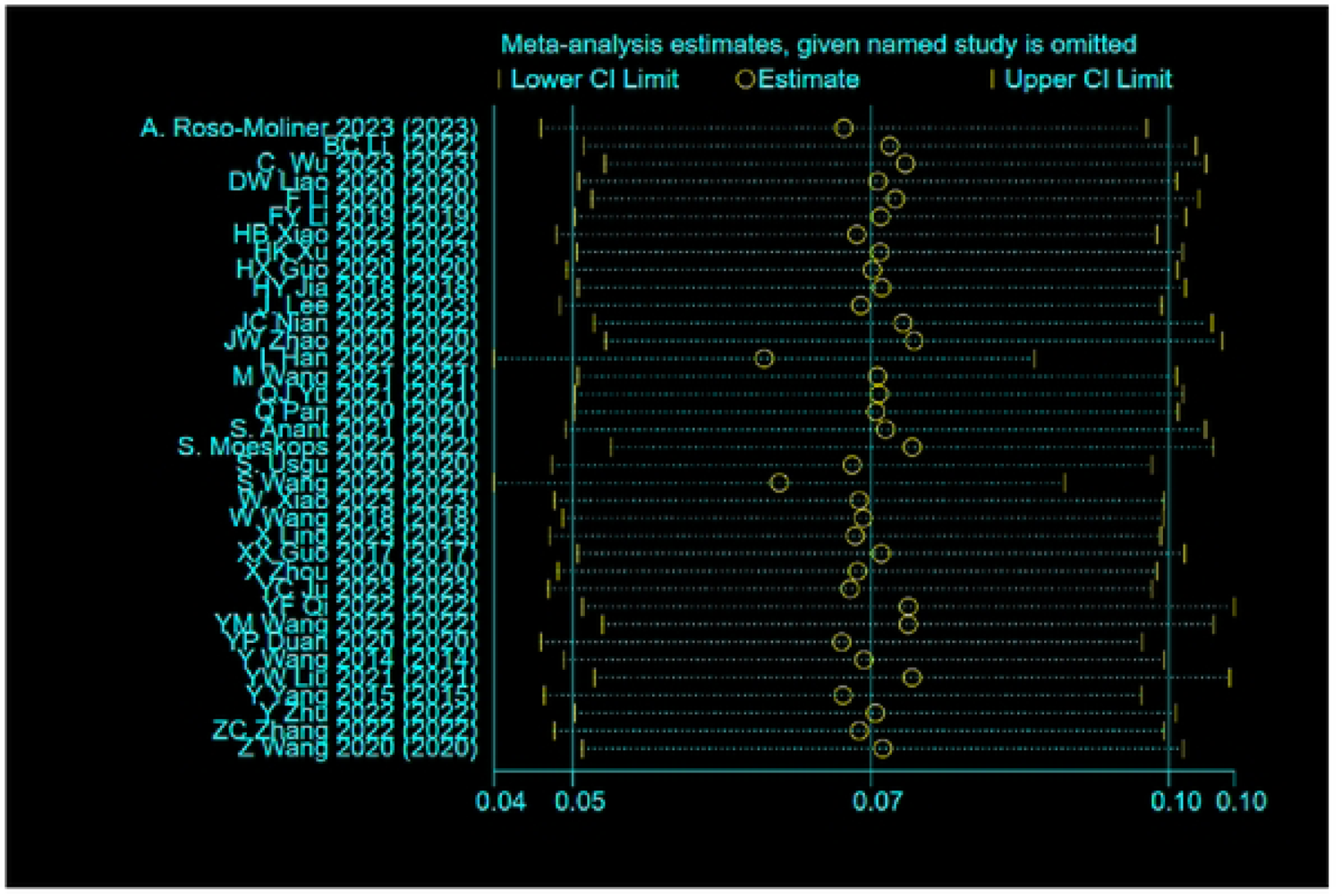
Sensitivity analysis 3 The effects of FT vs. TRT on power (standing long jump).

**Figure 13:**
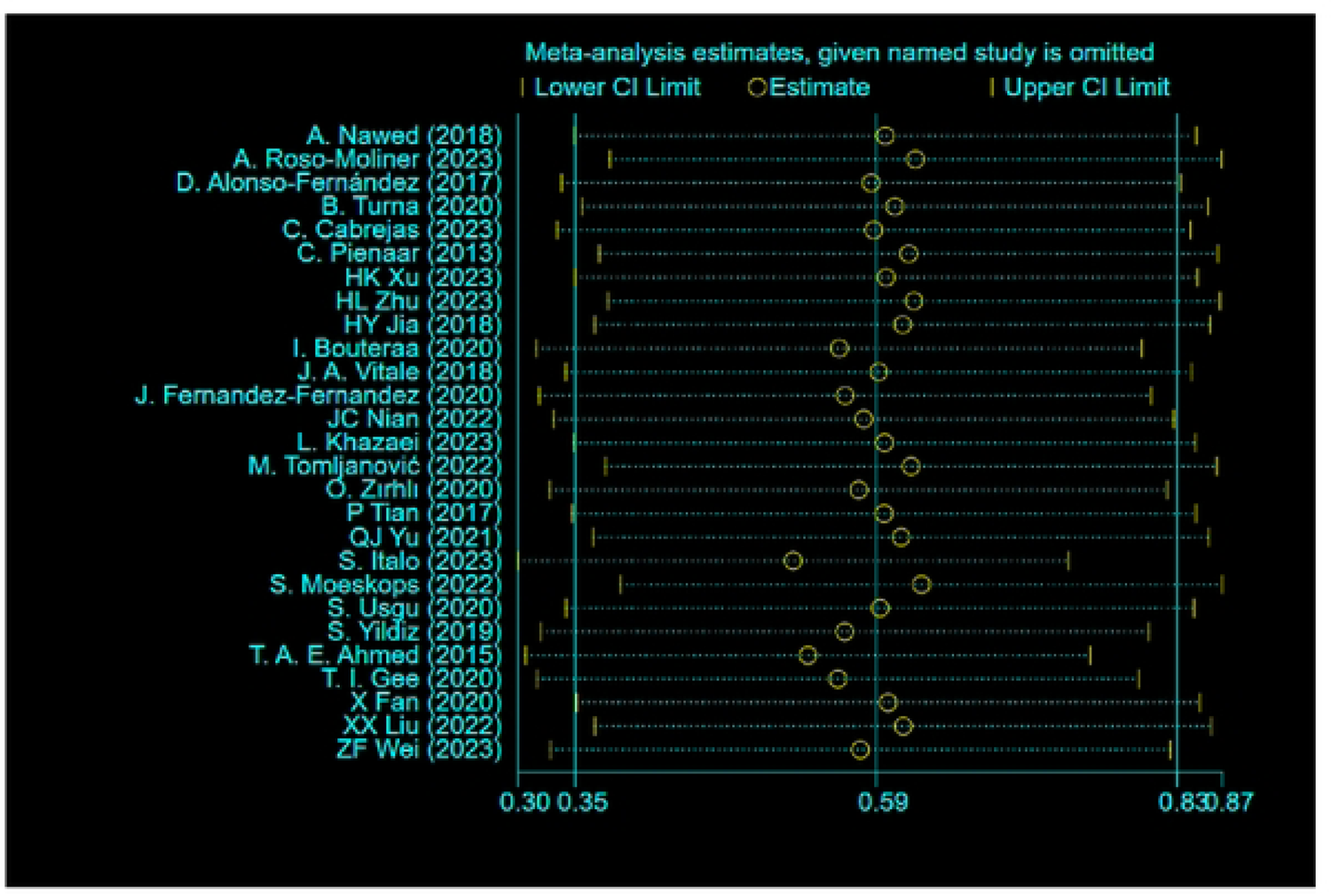
Sensitivity analysis 4 The effects of FT vs. TRT on power (countermovement jump).

**Figure 14:**
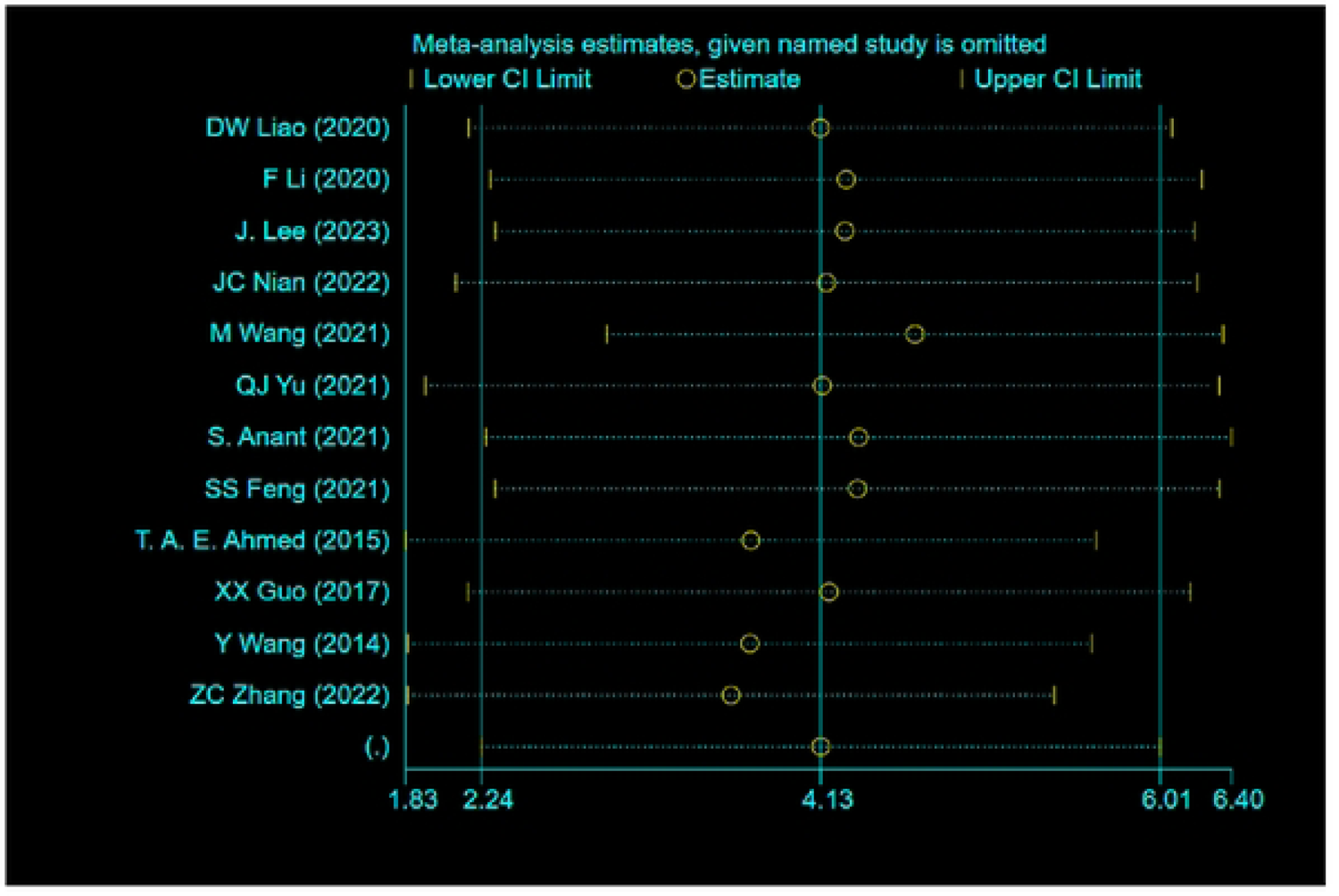
Sensitivity analysis 5 The effects of FT vs. TRT on muscle endurance (lmin sit-ups).

**Figure 15:**
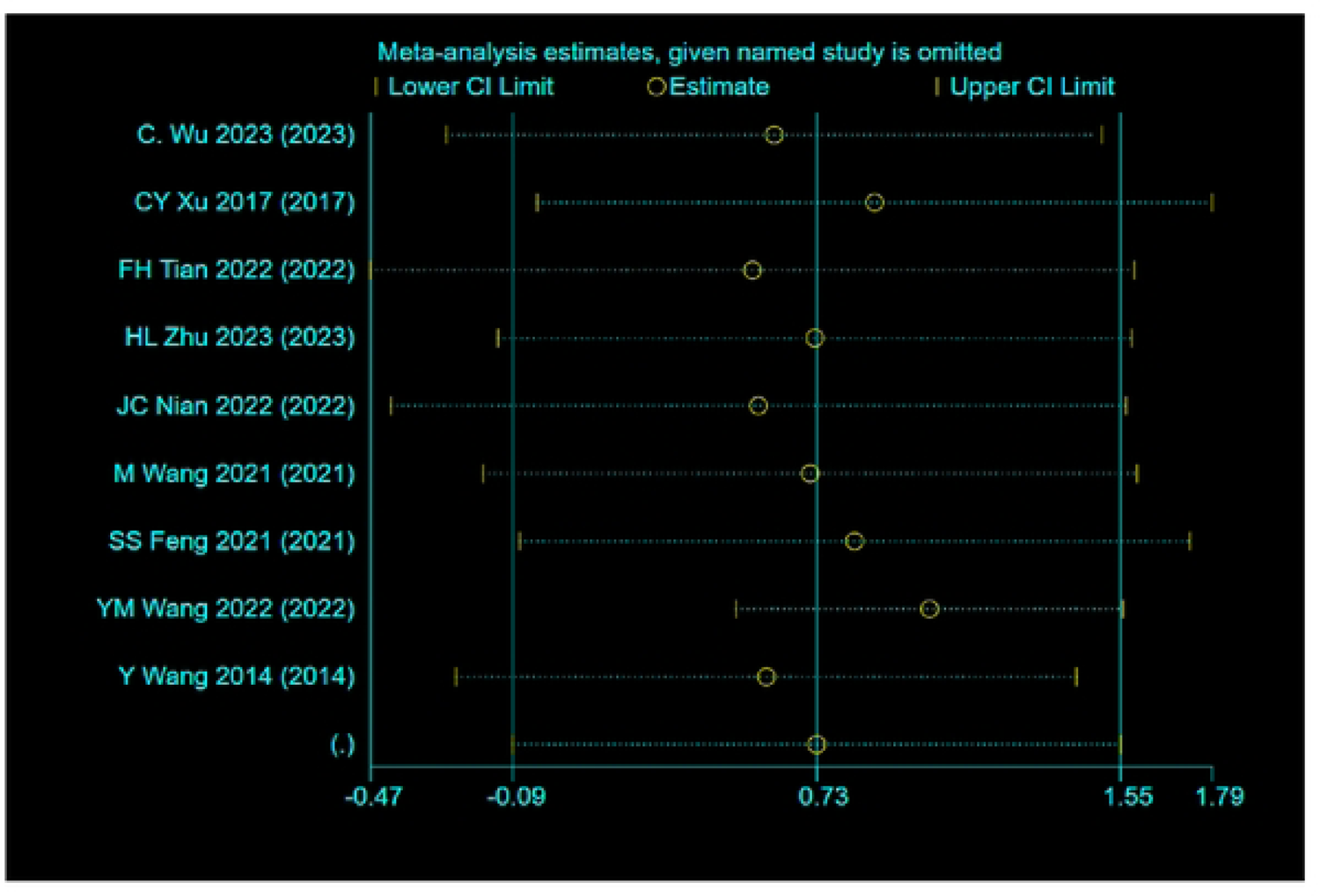
Sensitivity analysis 6 The effects of FT vs. TRT on muscle endurance (pull-ups).

**Figure 16:**
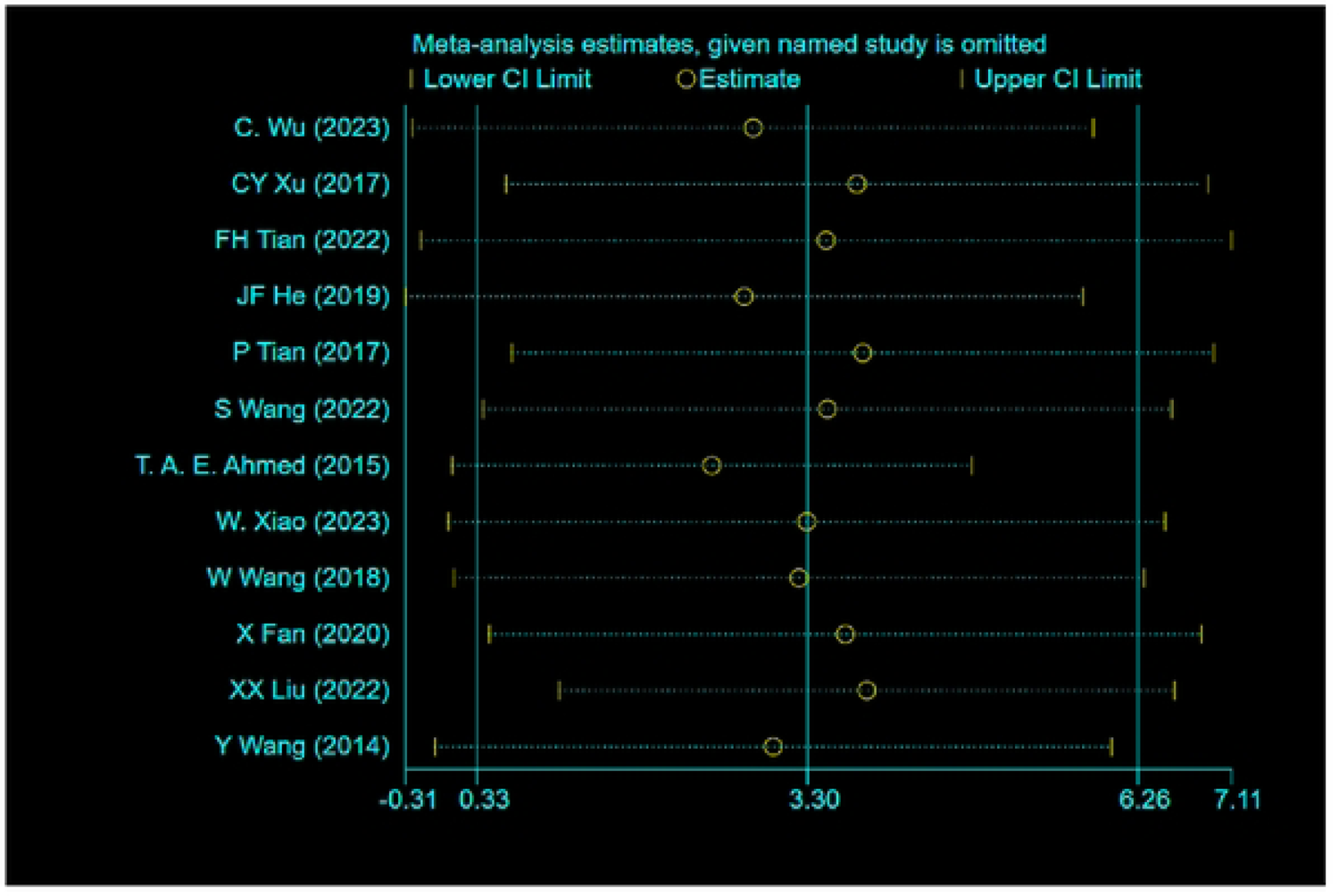
Sensitivity analysis 7 The effects of FT vs. TRT on muscle endurance (push-ups).

**Figure 17:**
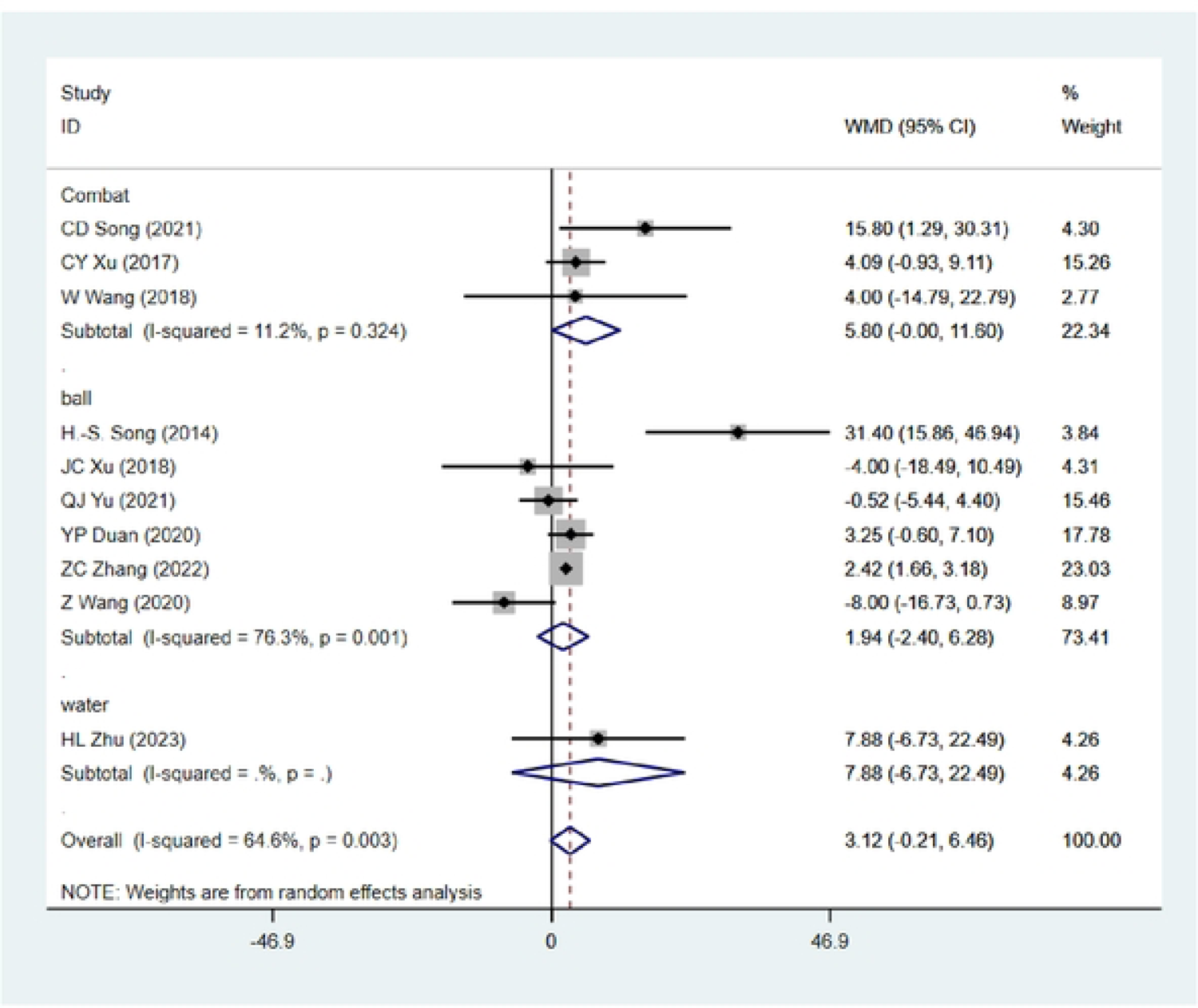
Subgroup Analysis-FT vs. TRT on Lower Limb Maximum Strength (DS/HS)

**Figure 18:**
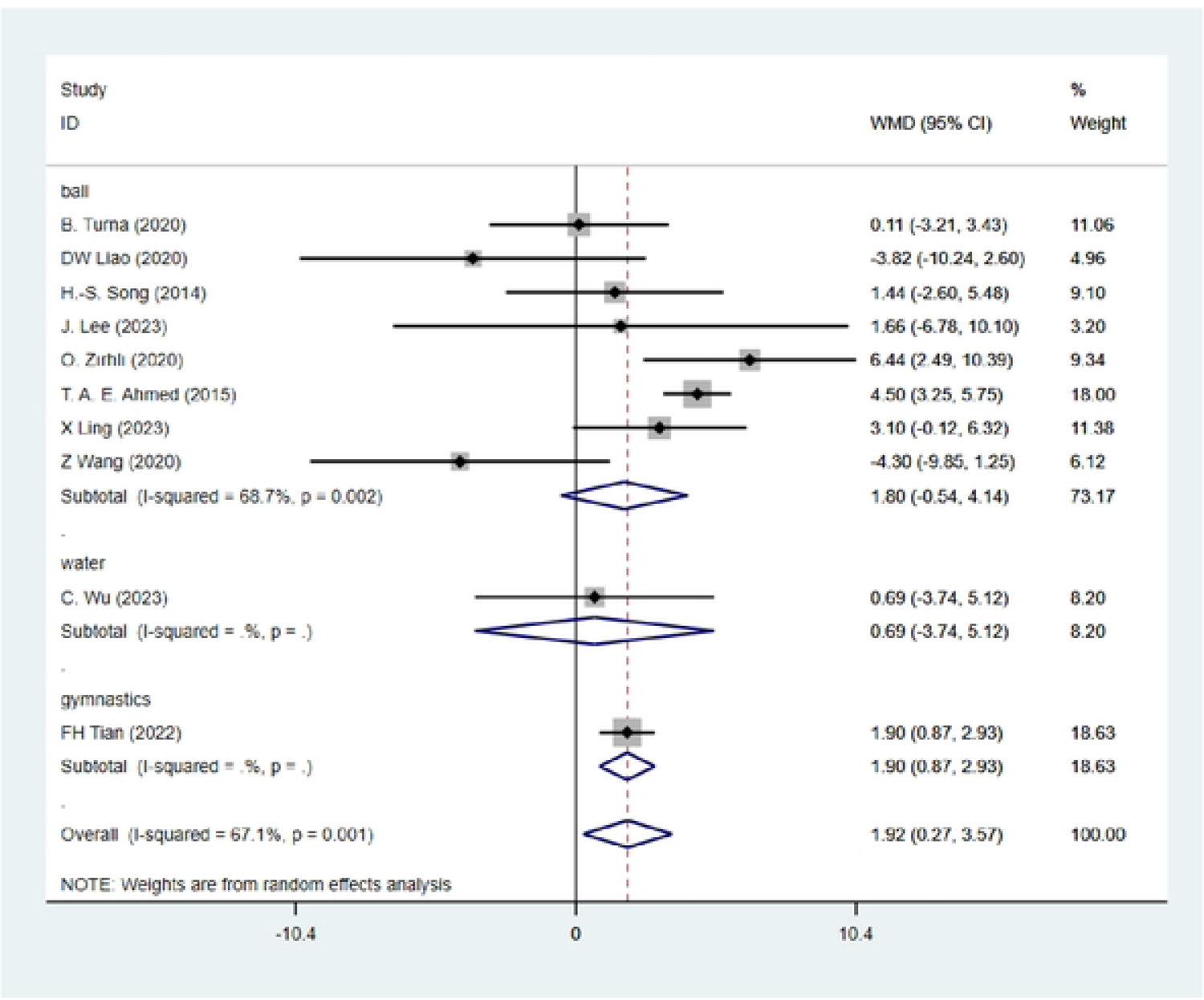
Subgroup Analysis-FT vs. TRT on Grip Strength (GS)

**Figure 19:**
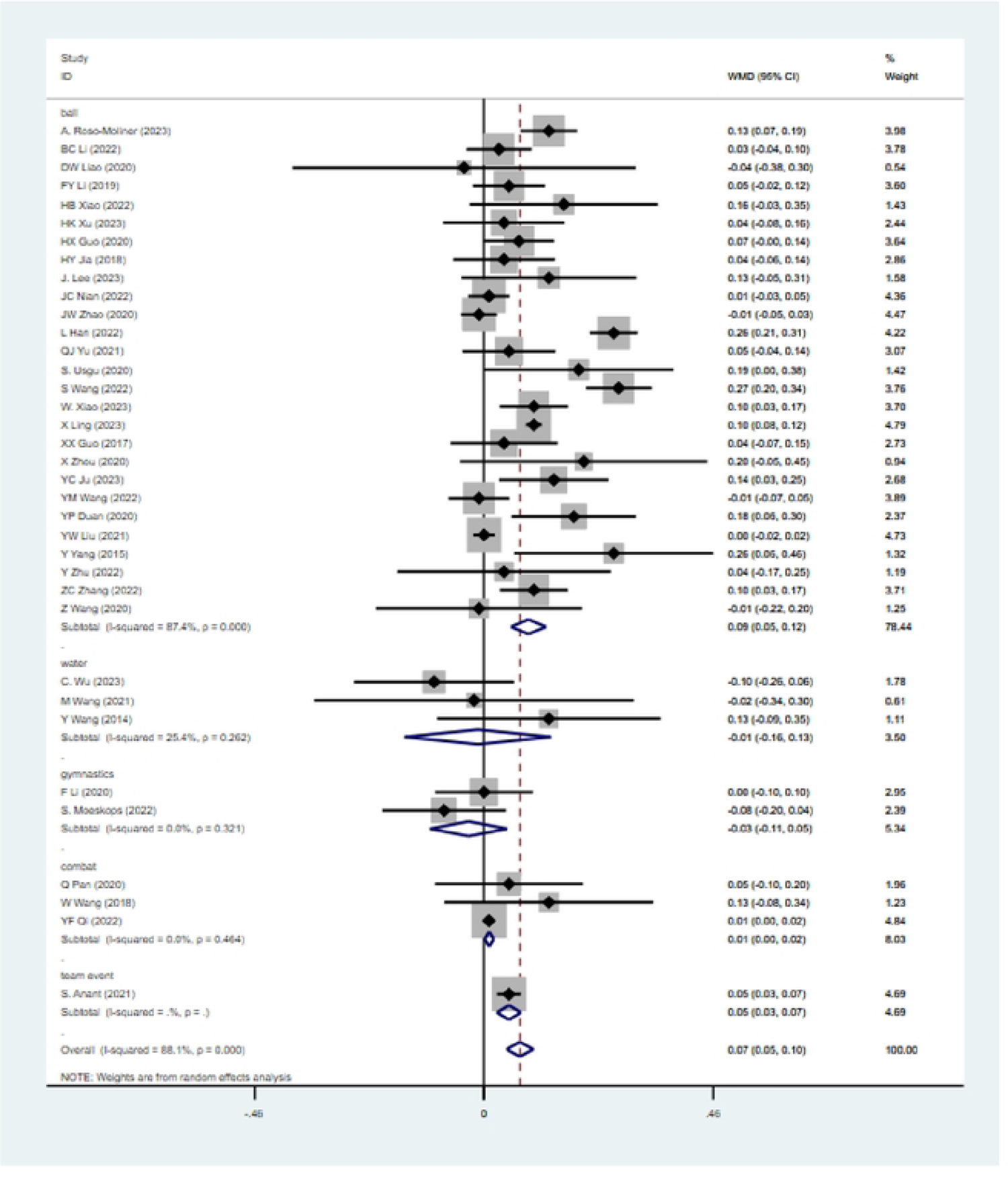
Subgroup Analysis-FT vs. TRT on Power (SLJ)

**Figure 20:**
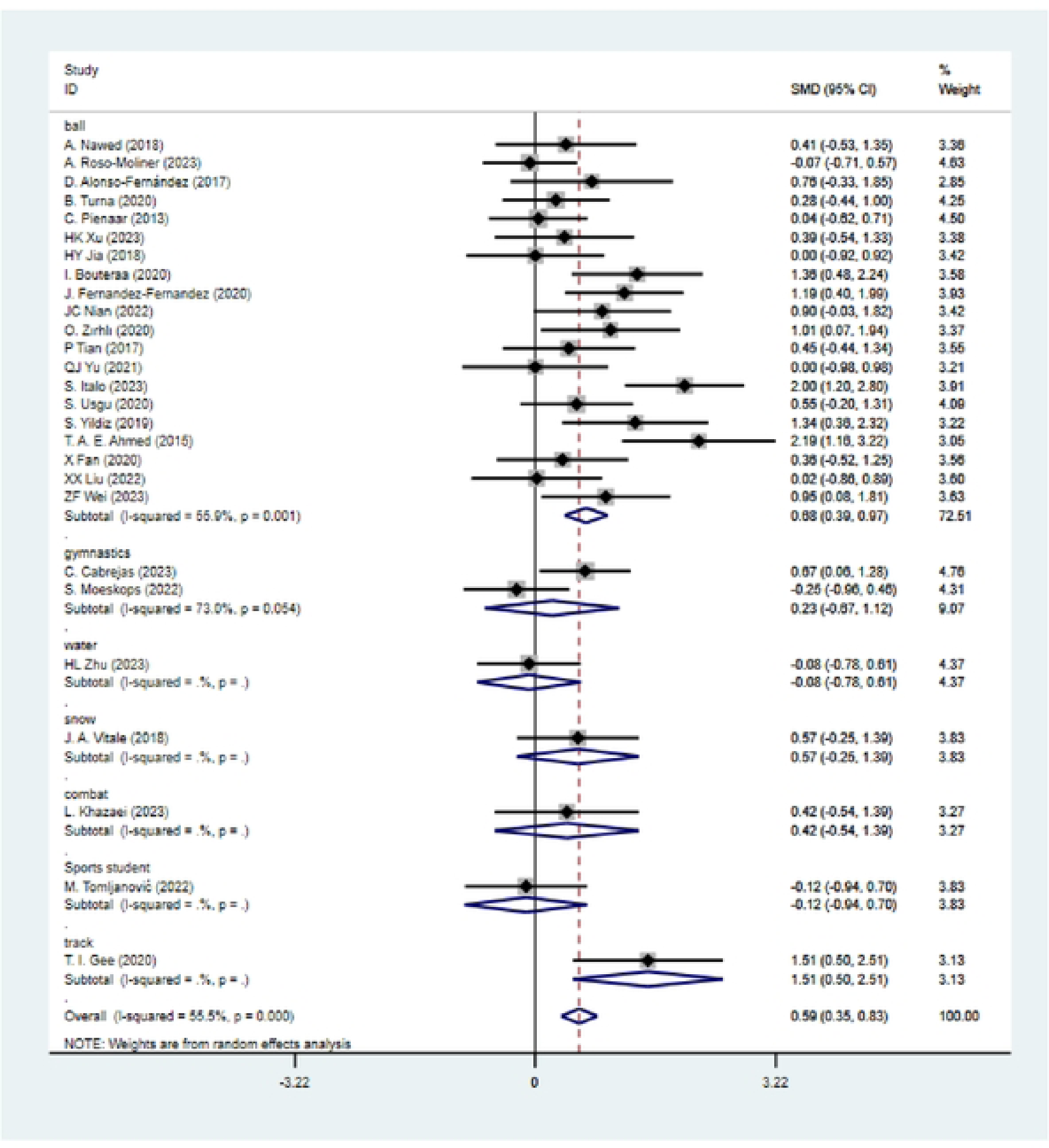
Subgroup Analysis-FT vs. TRT on Power (CMJ)

**Figure 21:**
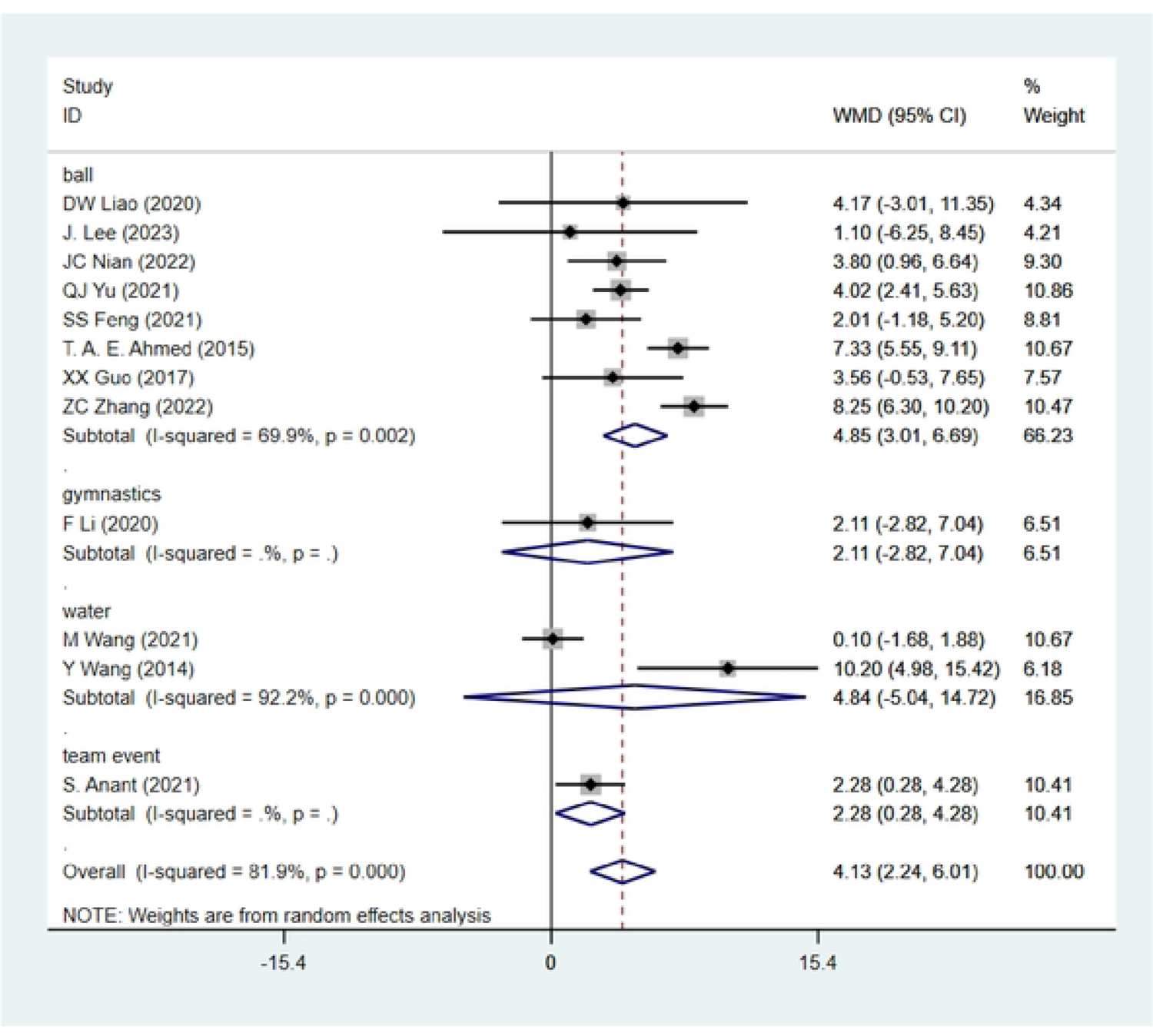
Subgroup Analysis-FT vs. TRT on Muscle Endurance (1 min SU)

**Figure 22:**
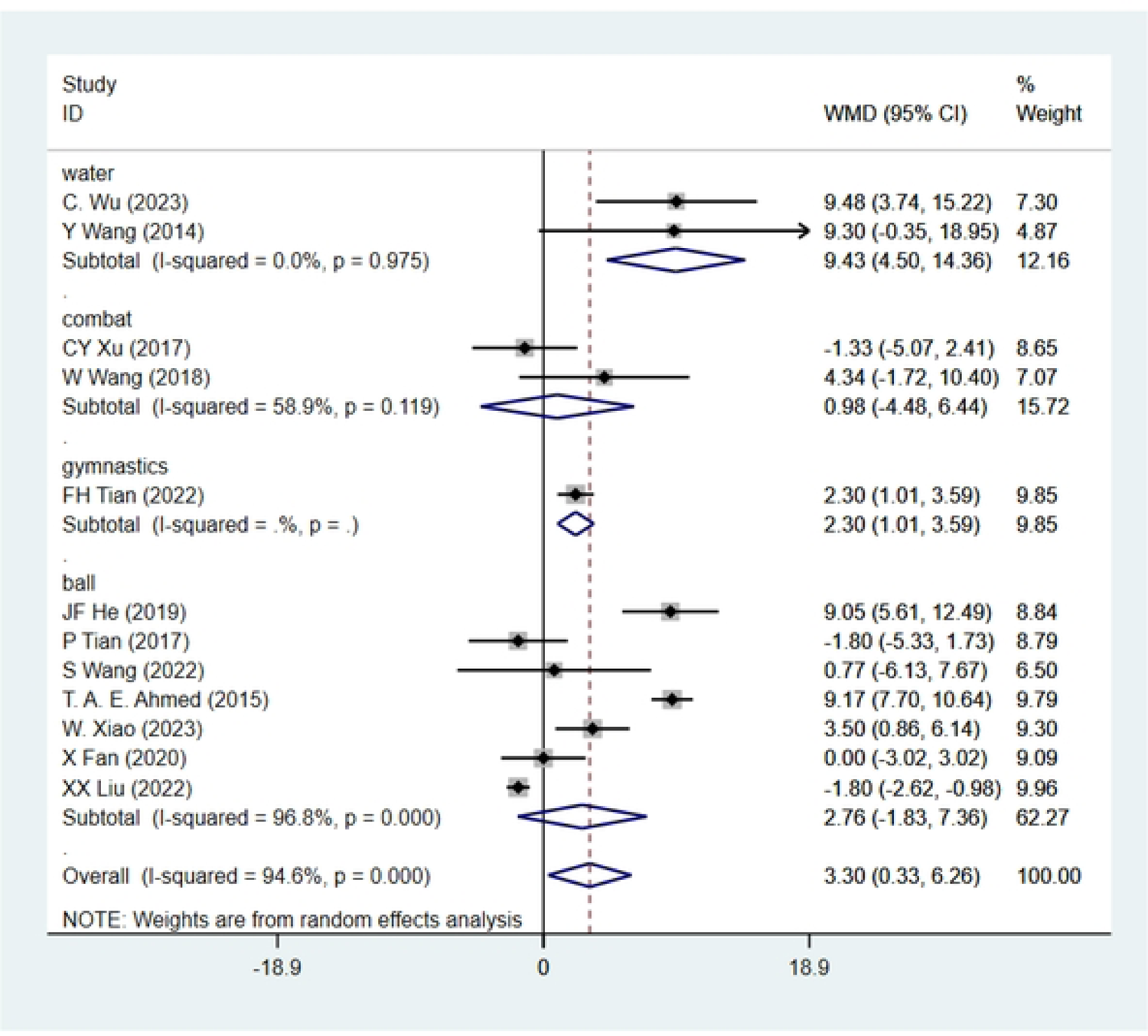
Subgroup Analysis-FT vs. TRT on Muscle Endurance (PSU)

**Figure 23:**
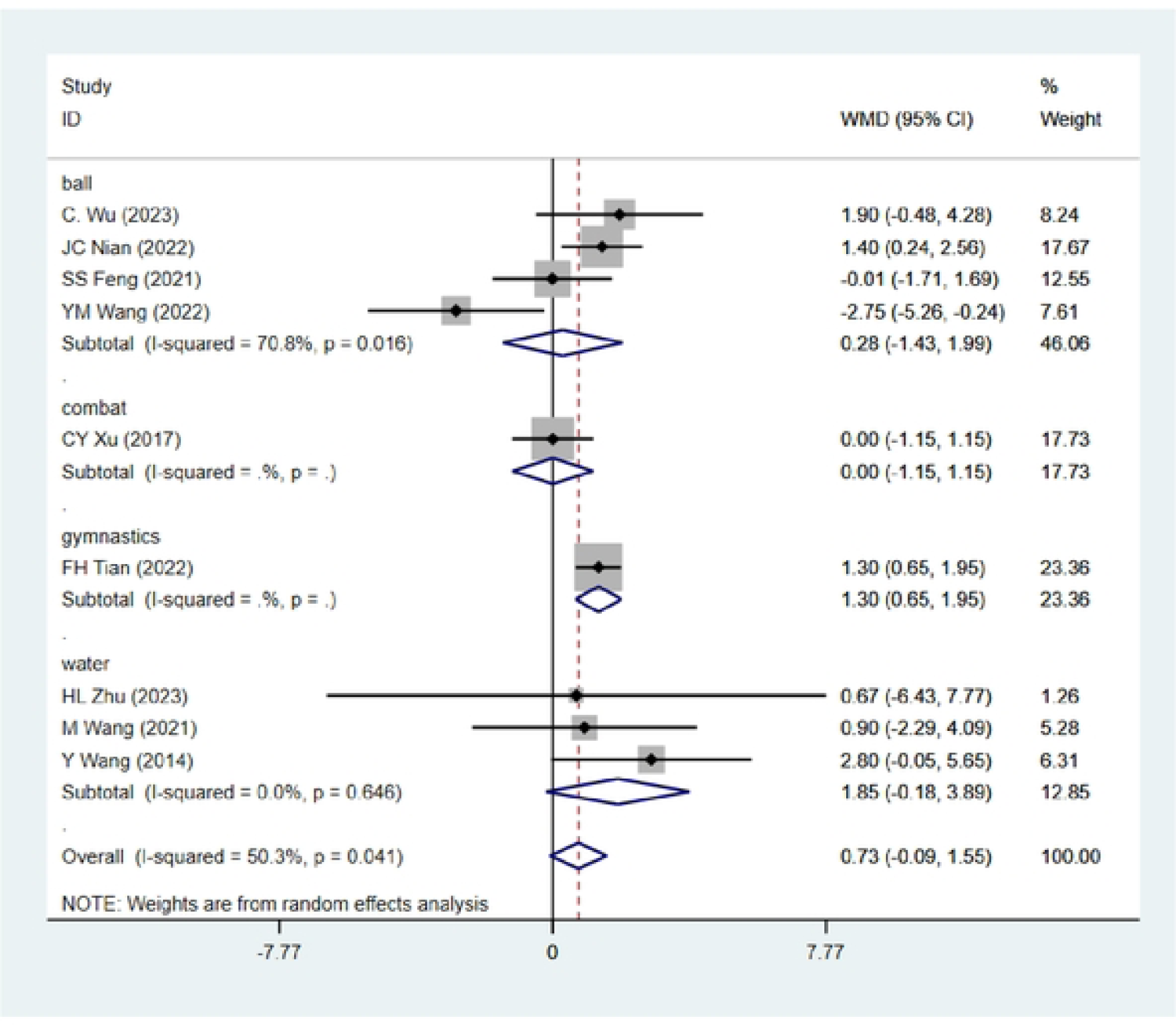
Subgroup Analysis-FT vs. TRT on Muscle Endurance (PLU)

**Figure 24:**
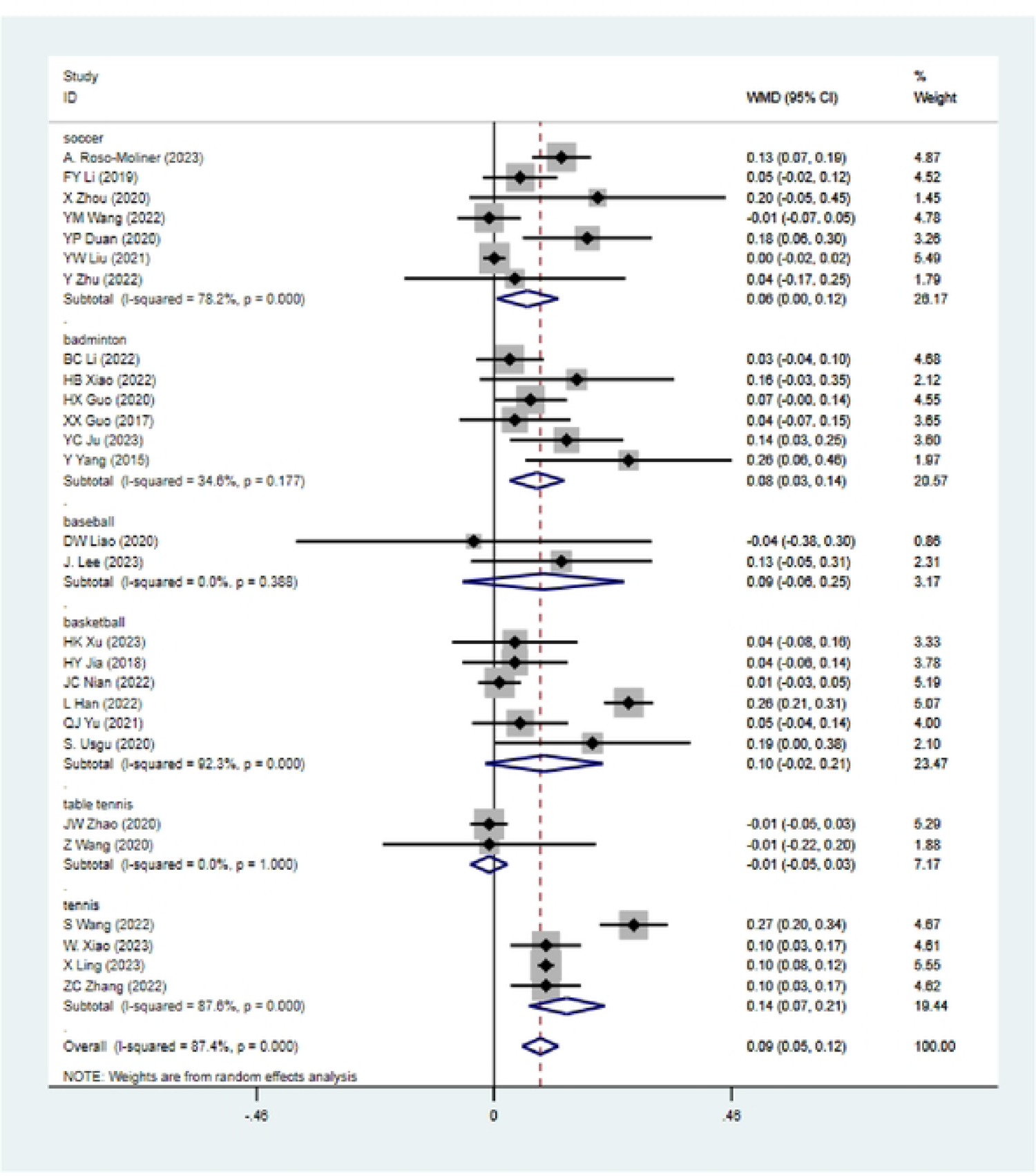
Subgroup Analysis-Impact of FT vs. TRT on Power (SLJ) in Ball Sport Athletes

**Figure 25:**
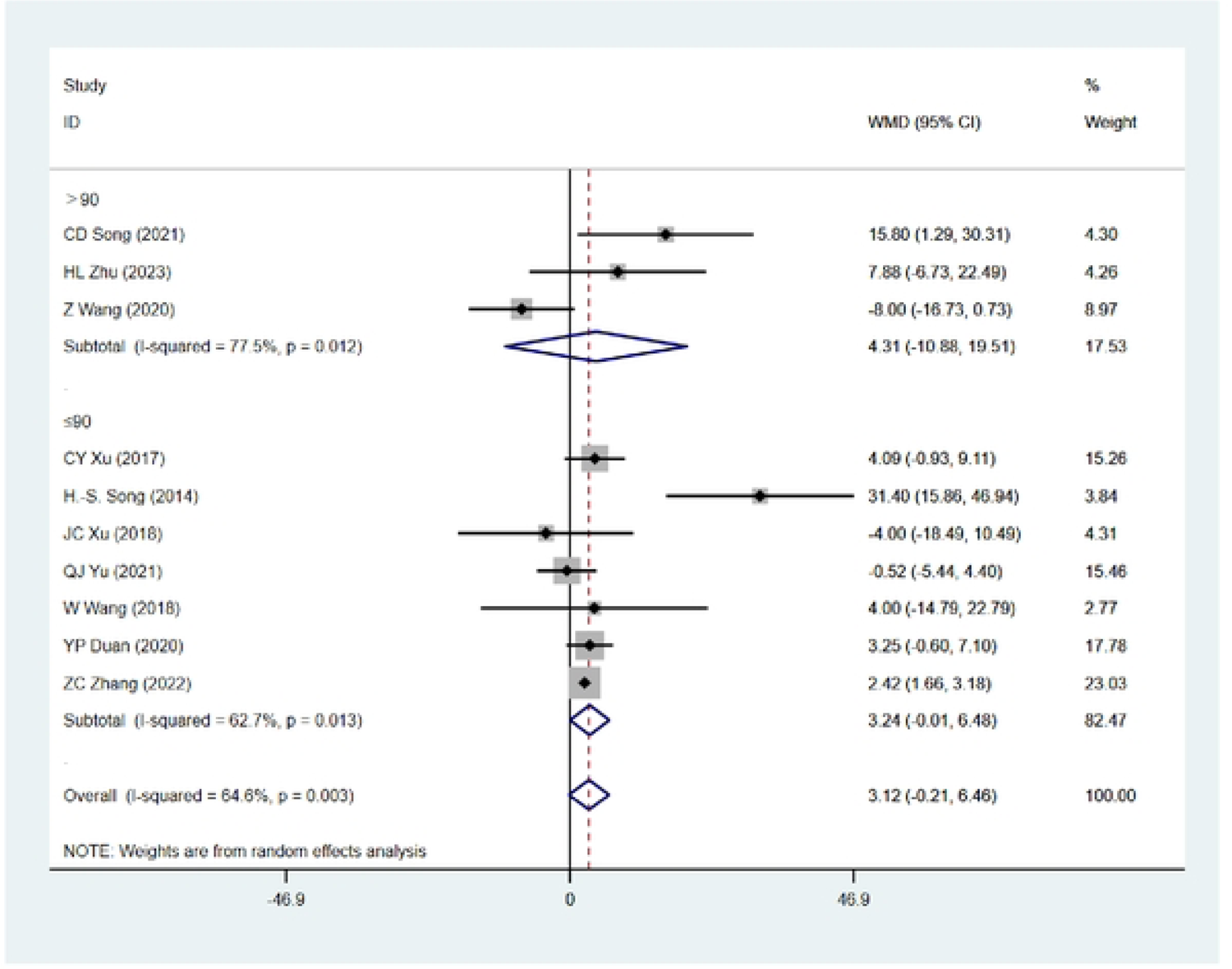
The effects of FT vs. TRT on lower limb maximal strength (DS/HS)-Each session duration.

**Figure 26:**
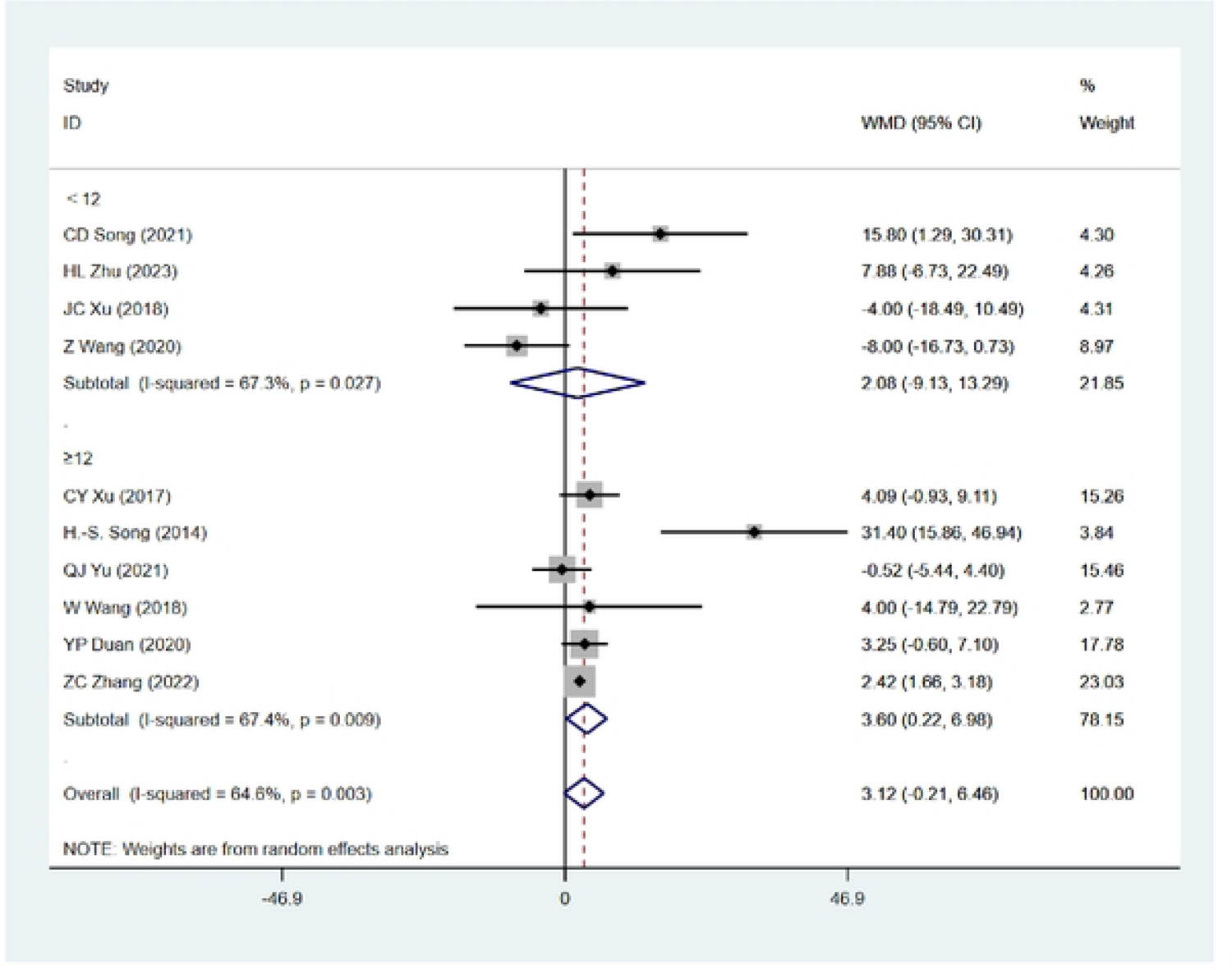
The effects of FT vs. TRT on lower limb maximal strength (DS/HS)-Tolal weeks

**Figure 27:**
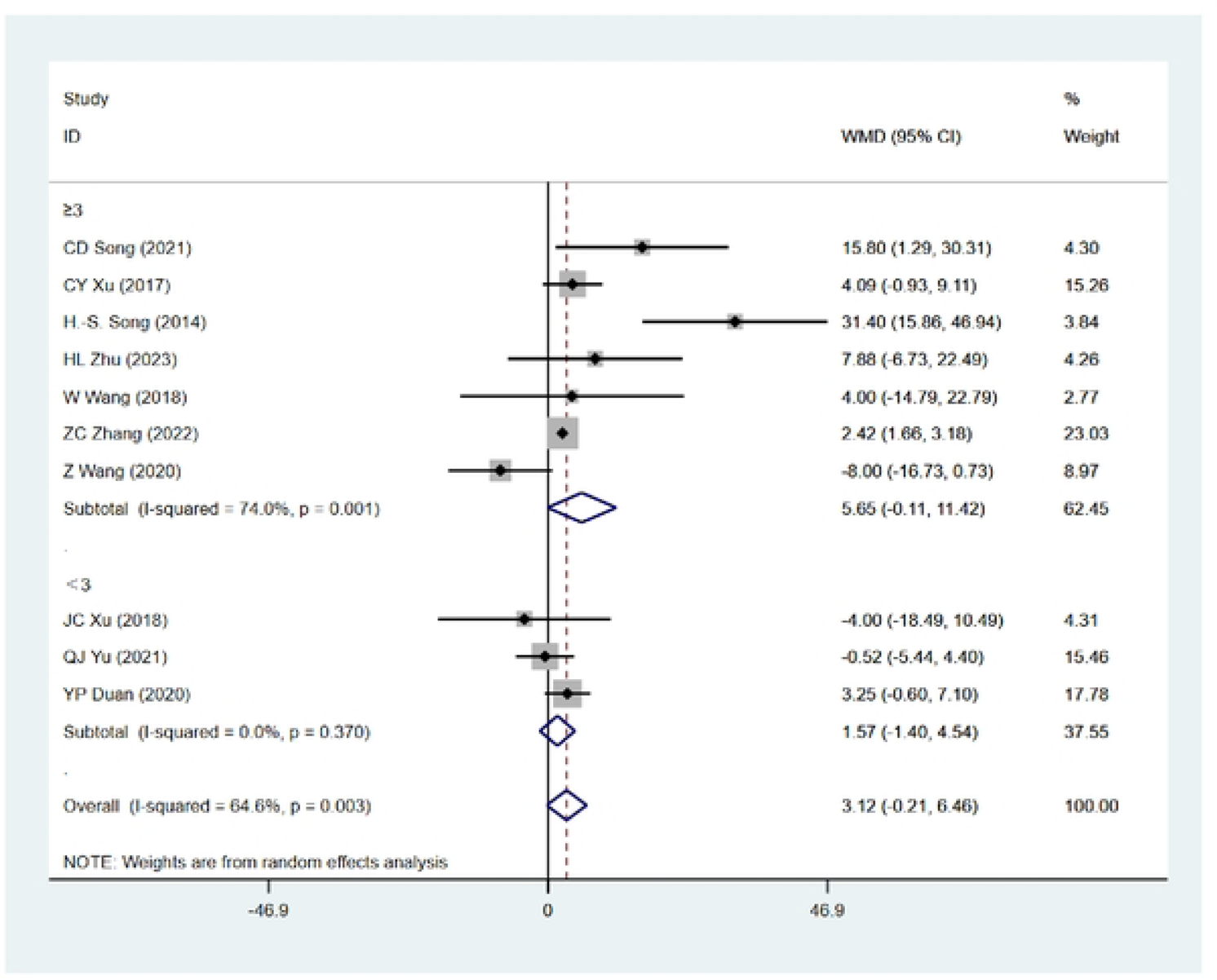
The effects of FT vs. TRT on lower limb maximal strength (DS/HS)-Frequency

**Figure 28:**
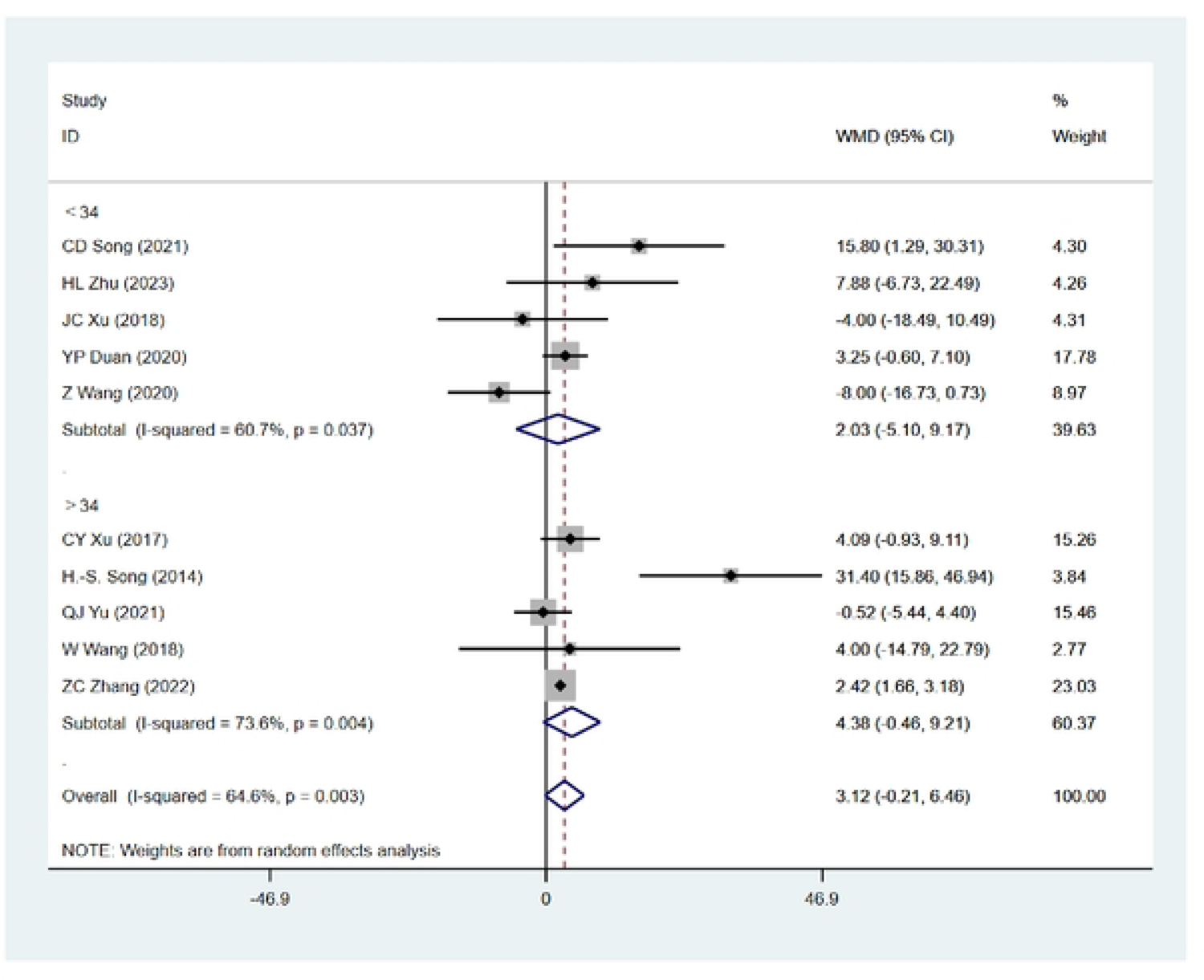
The effects of FT vs. TRT on lower limb maximal strength (DSZHS)­Total sessions

**Figure 29:**
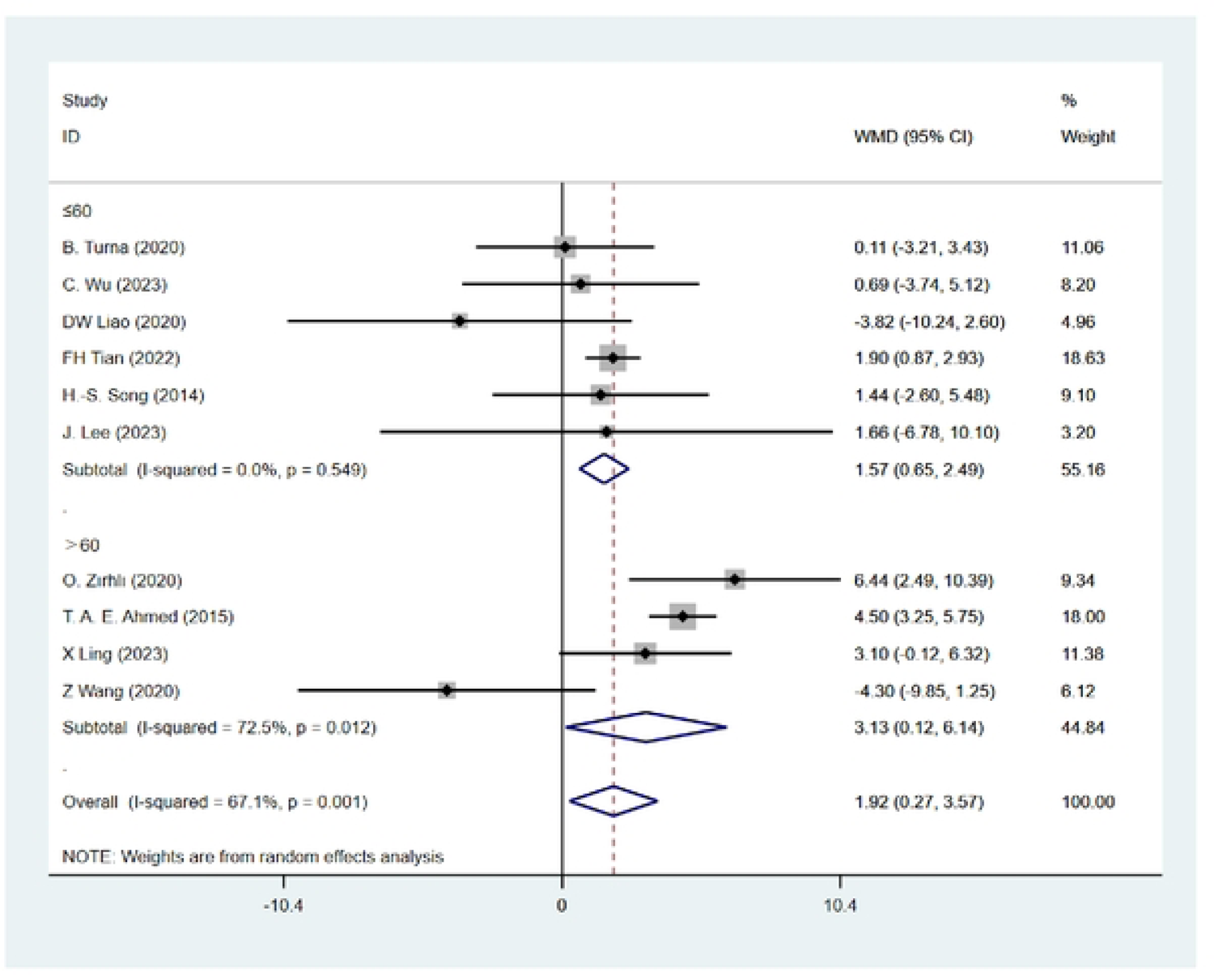
The effects of FT vs. TRT on grip strength (GS)­Each session duration.

**Figure 30:**
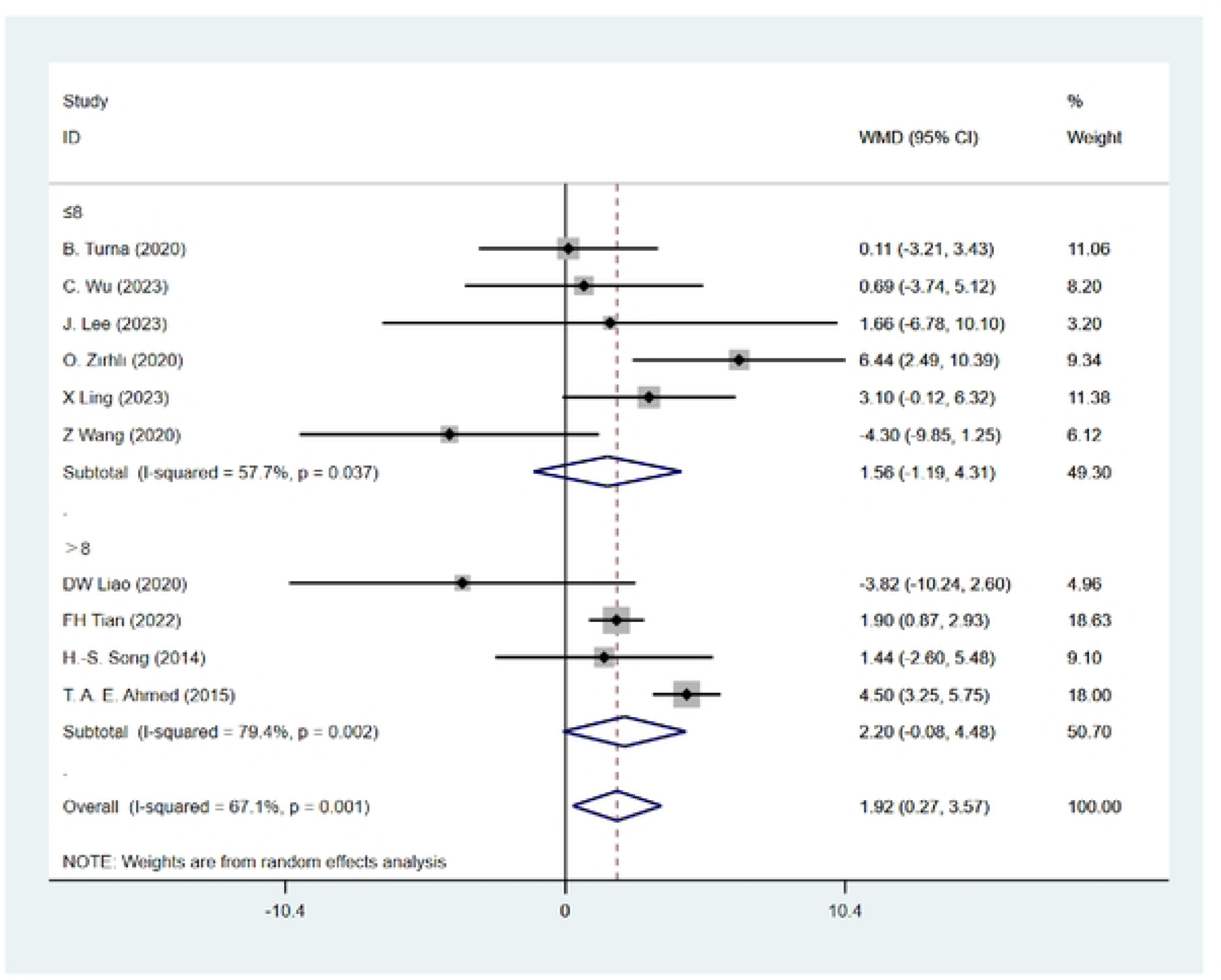
The effects of FT vs. TRT on grip strength (GS) - Total weeks

**Figure 31:**
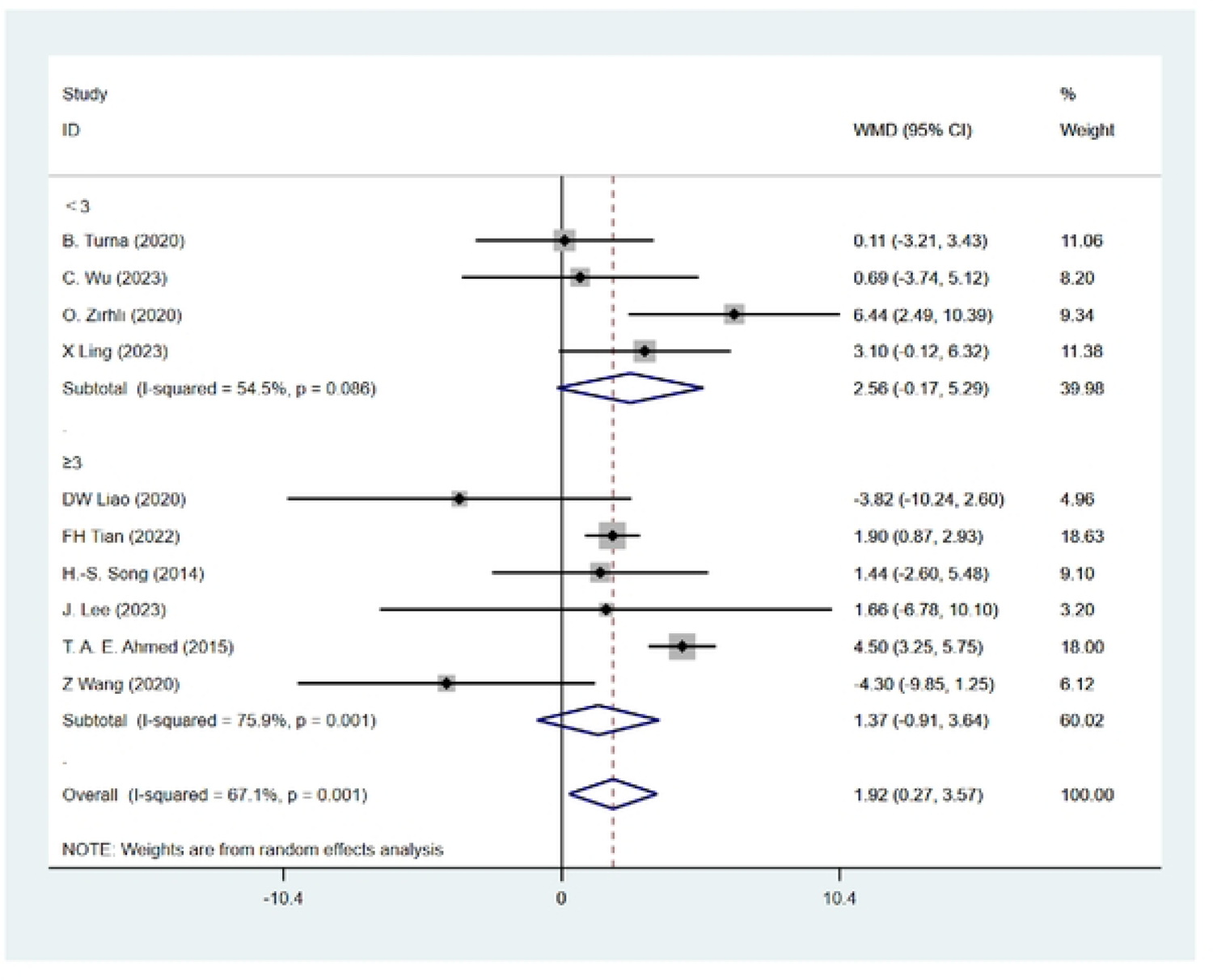
The effects of FT vs. TRT on grip strength (GS) - Frequency

**Figure 32:**
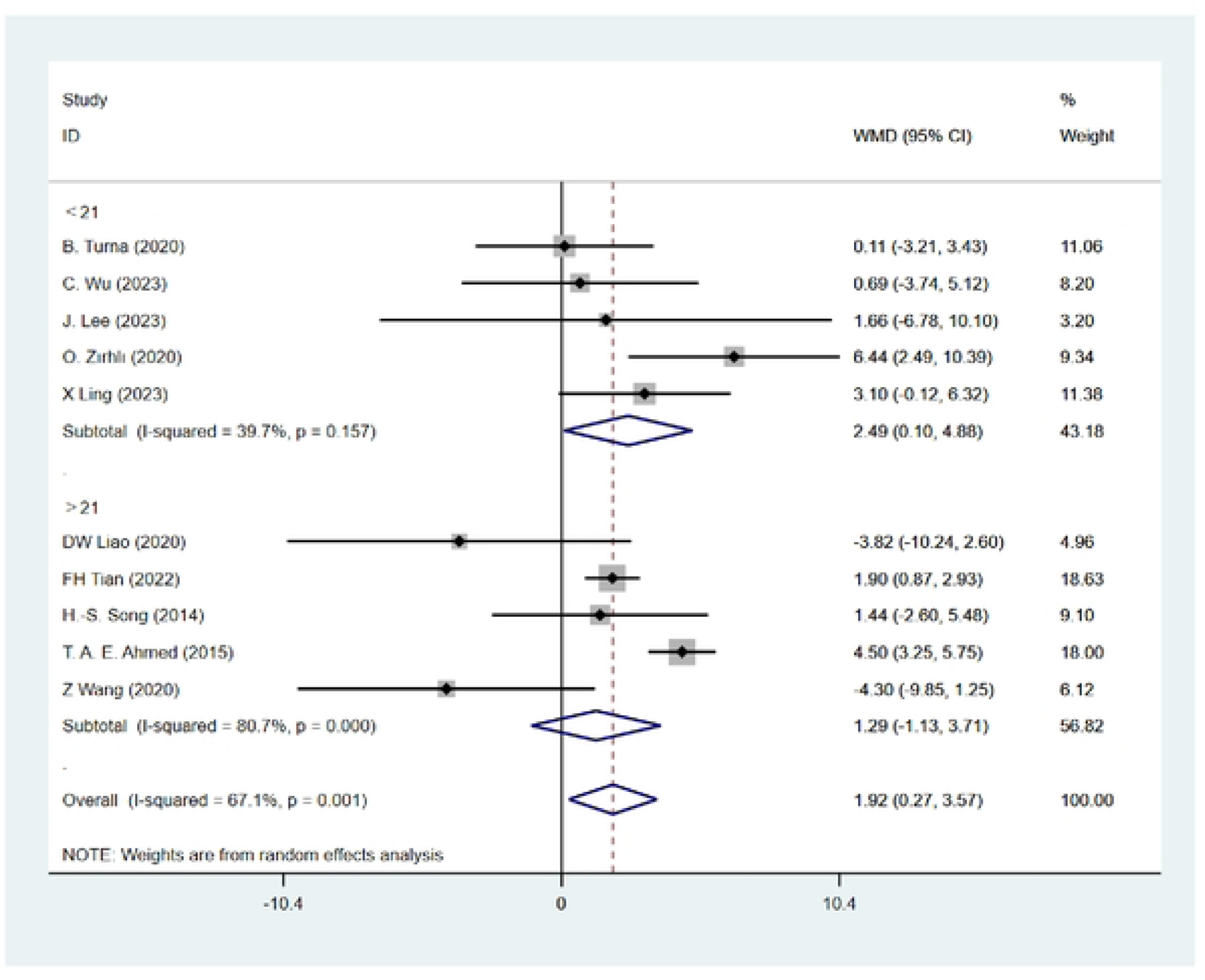
The effects of FT vs. TRT on grip strength (GS) - Total sessions

**Figure 33:**
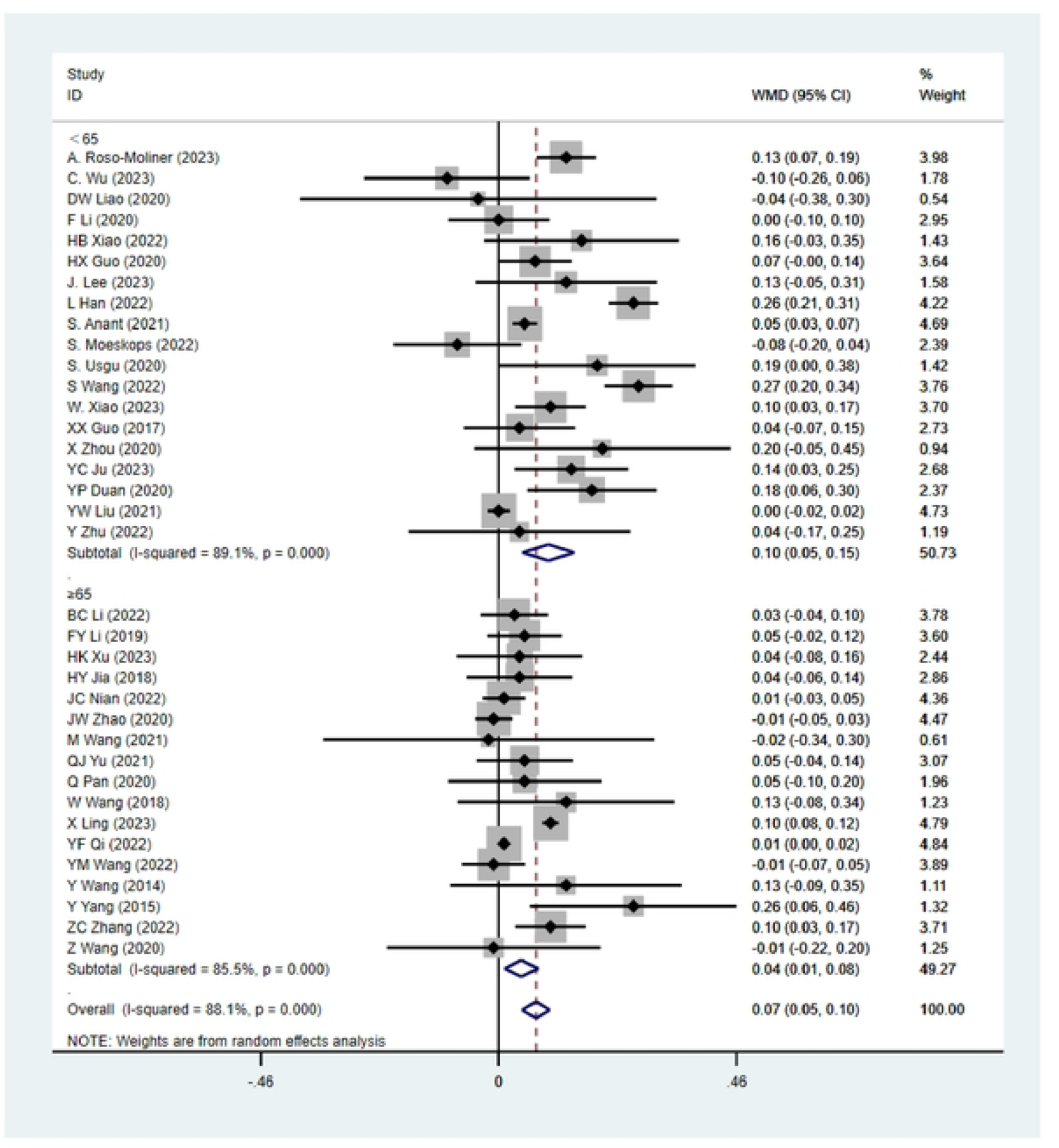
The effects of FT vs. TRT on power (SLJ) - Each session duration.

**Figure 34:**
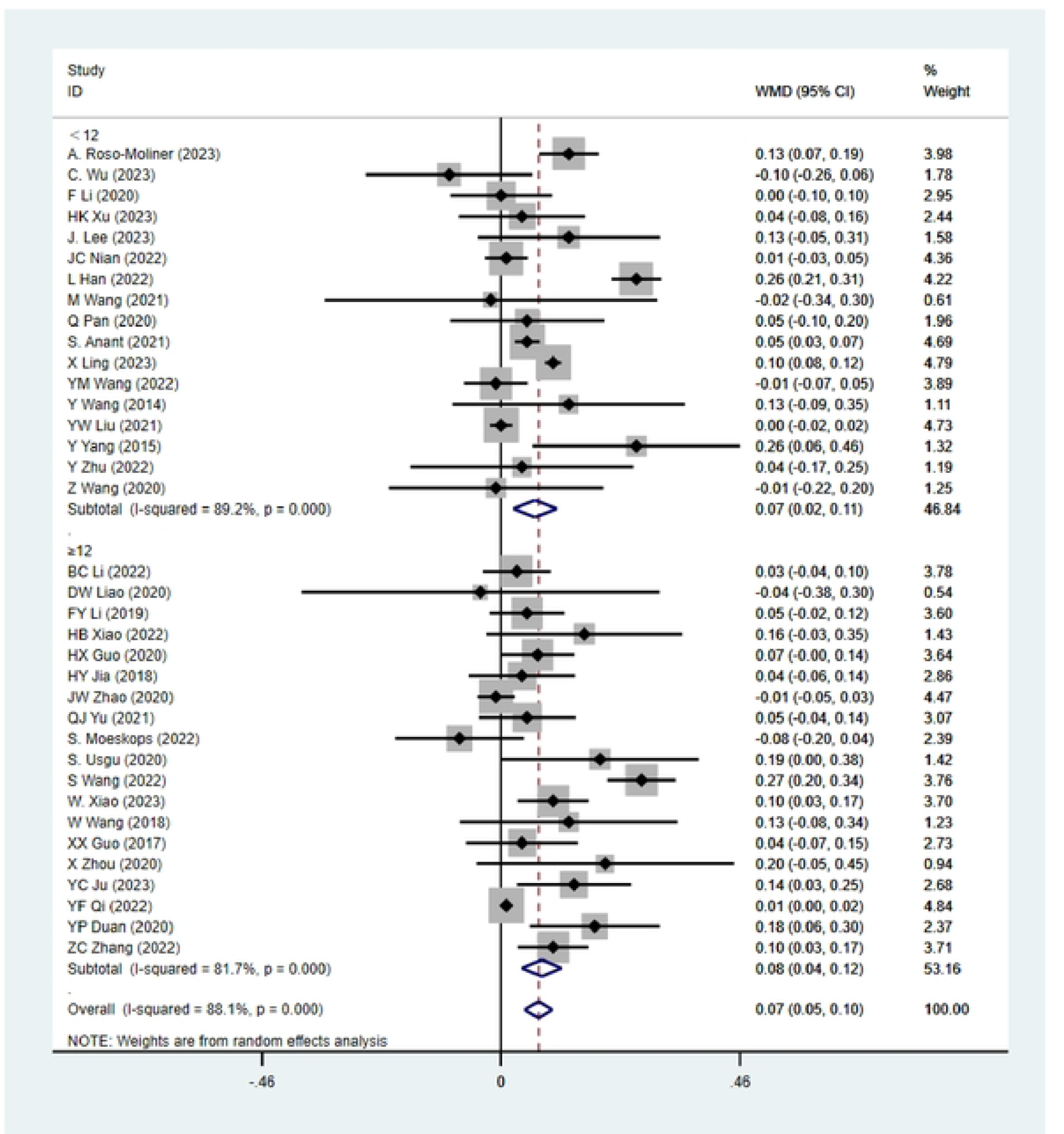
The effects of FT vs. TRT on power (SLJ) - Total weeks.

**Figure 35:**
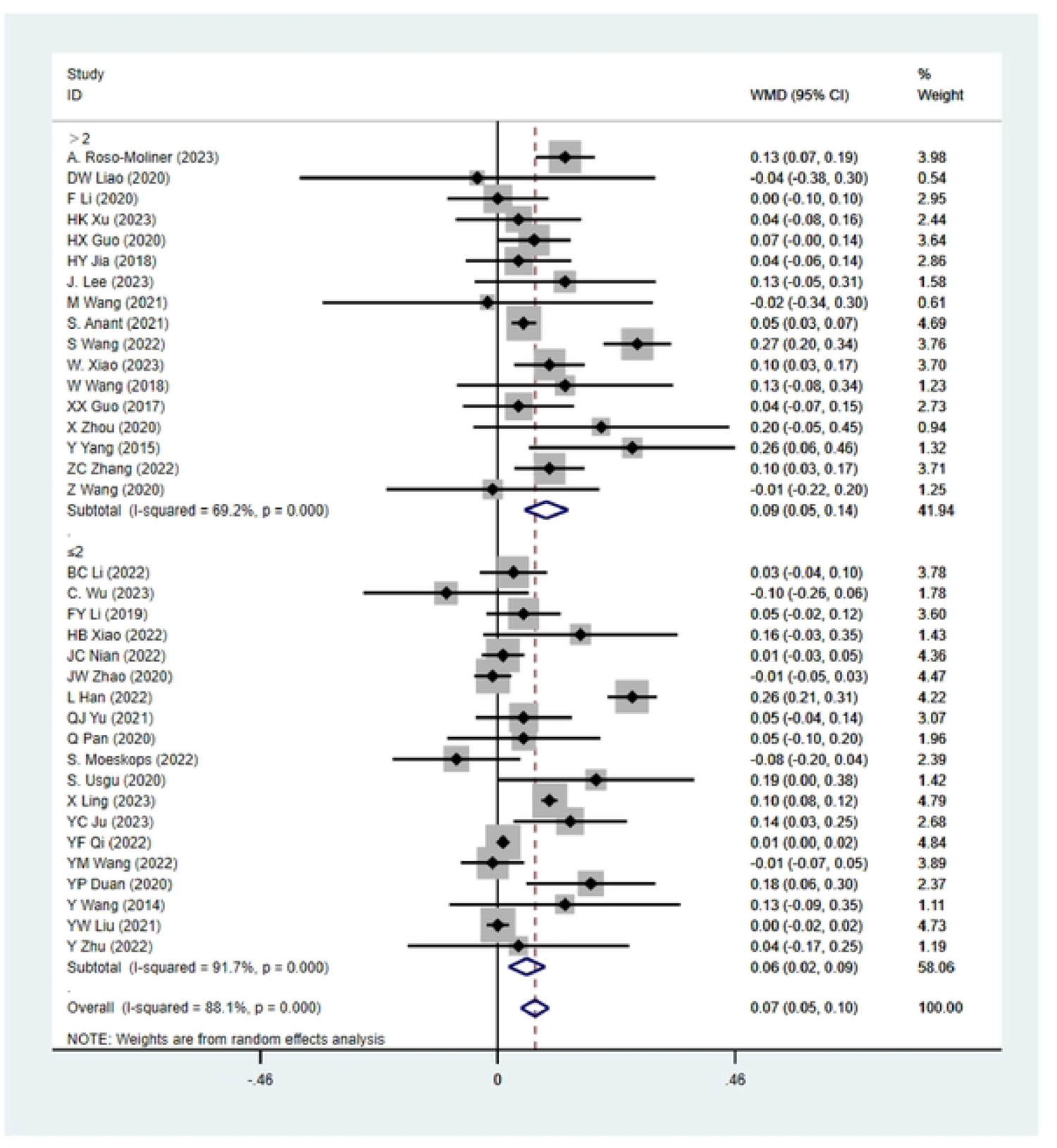
The effects of FT vs. TRT on power (SLJ) - Frequency

**Figure 36:**
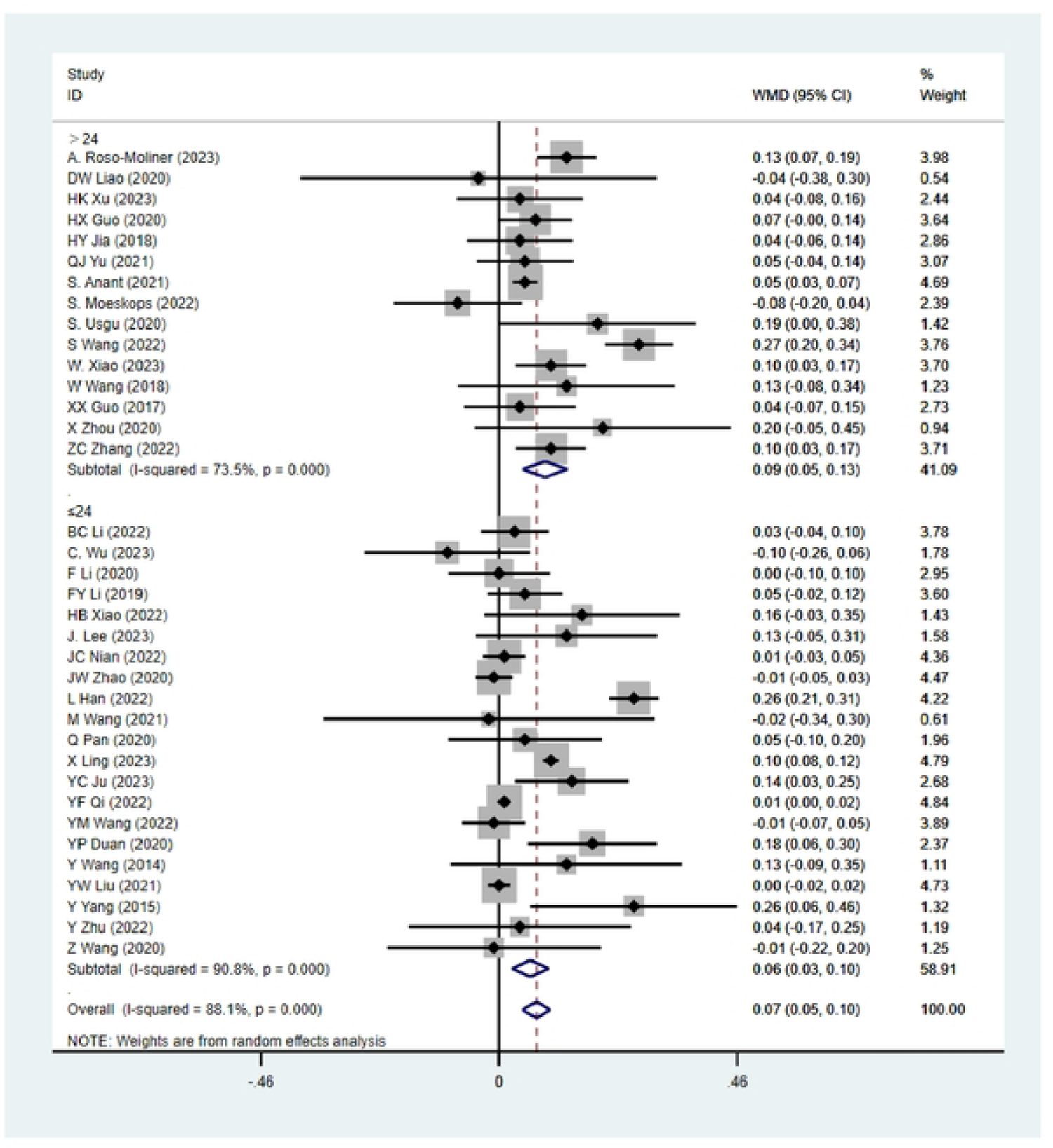
The effects of FT vs. TRT on power (SLJ) - Total sessions

**Figure 37:**
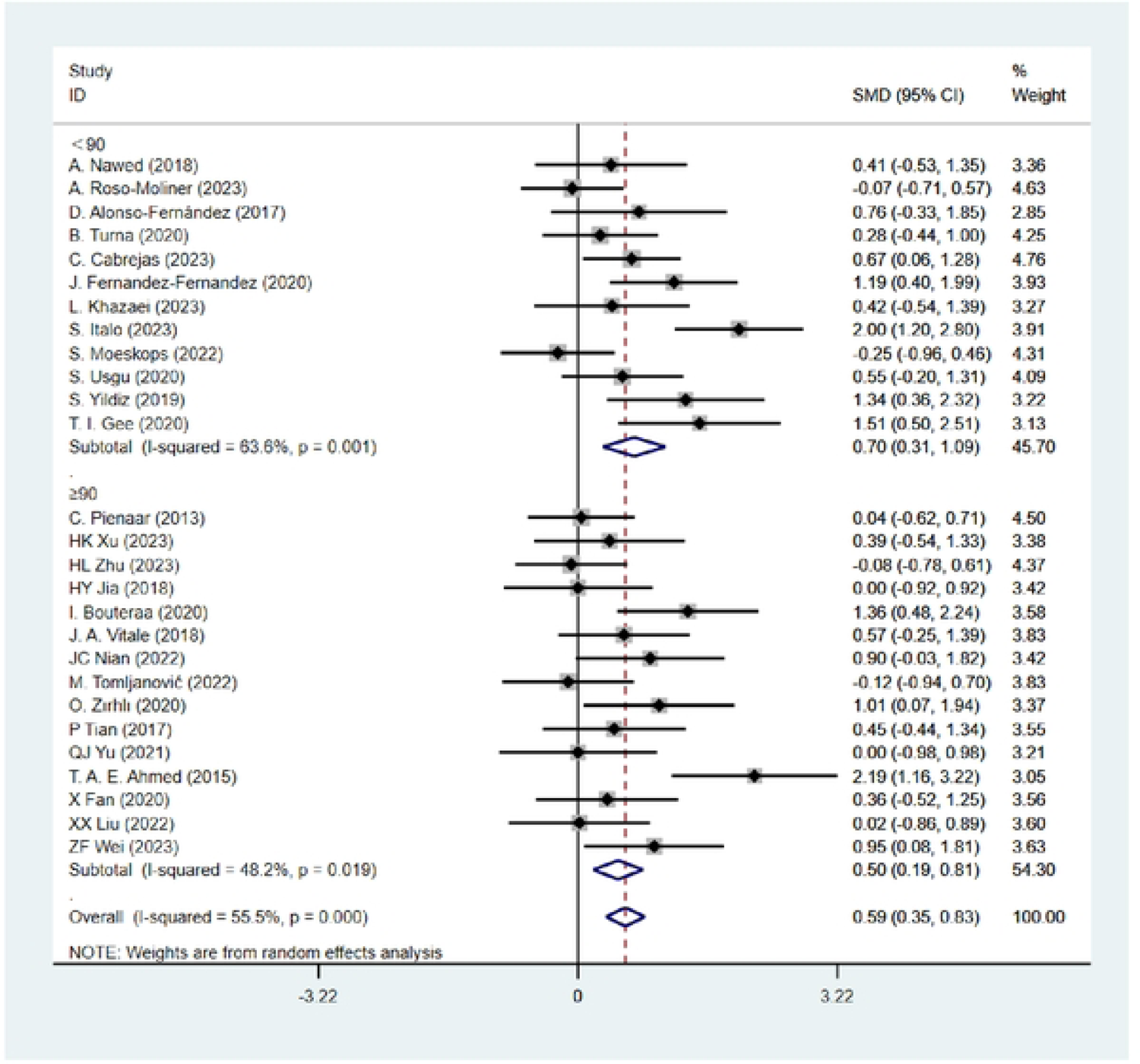
The effects of FT vs. TRT on power (CMJ) - Each session duration.

**Figure 38:**
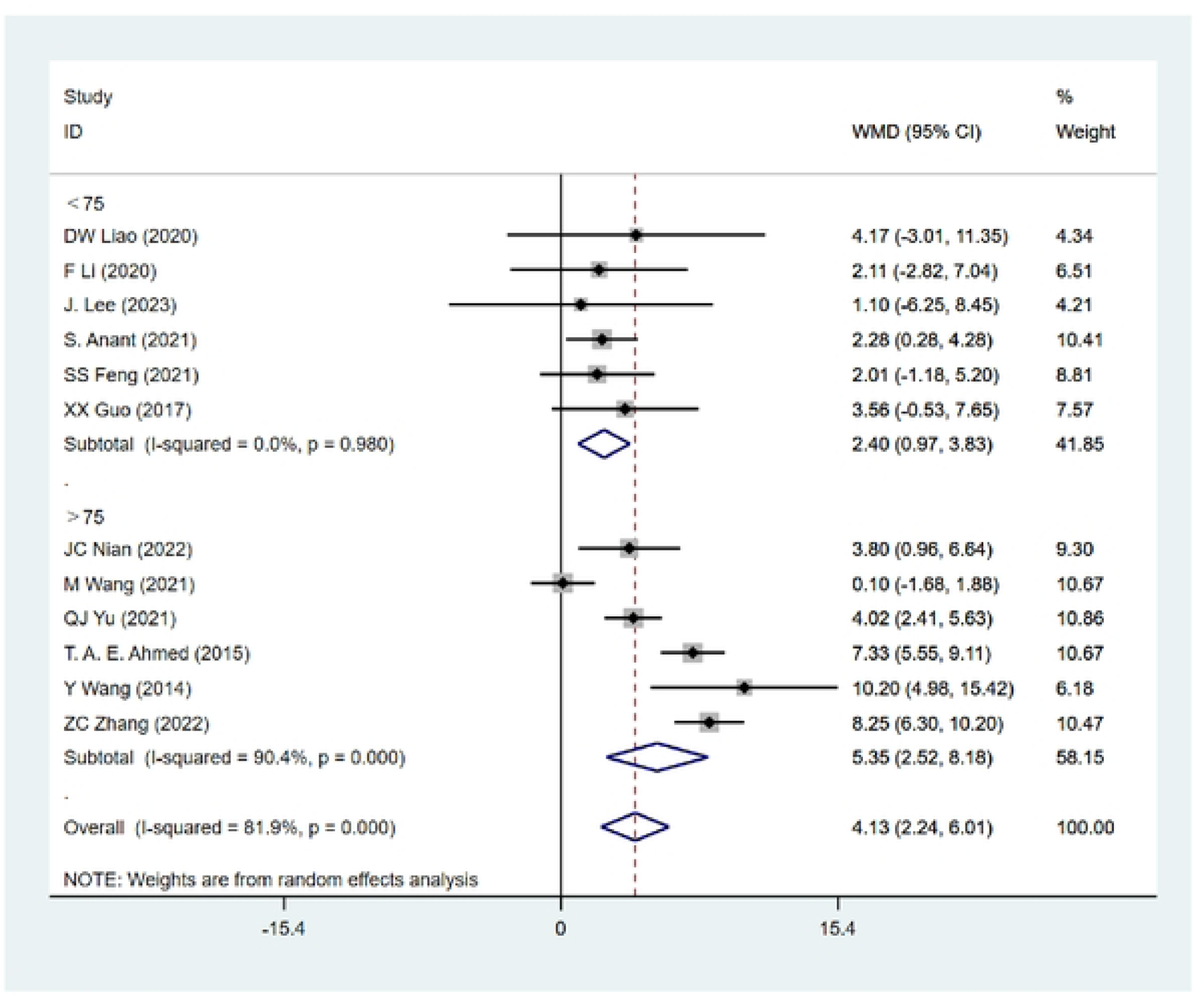
The effects of FT vs. TRT on muscle endurance (1 min SU) - Each session duration.

**Figure 39:**
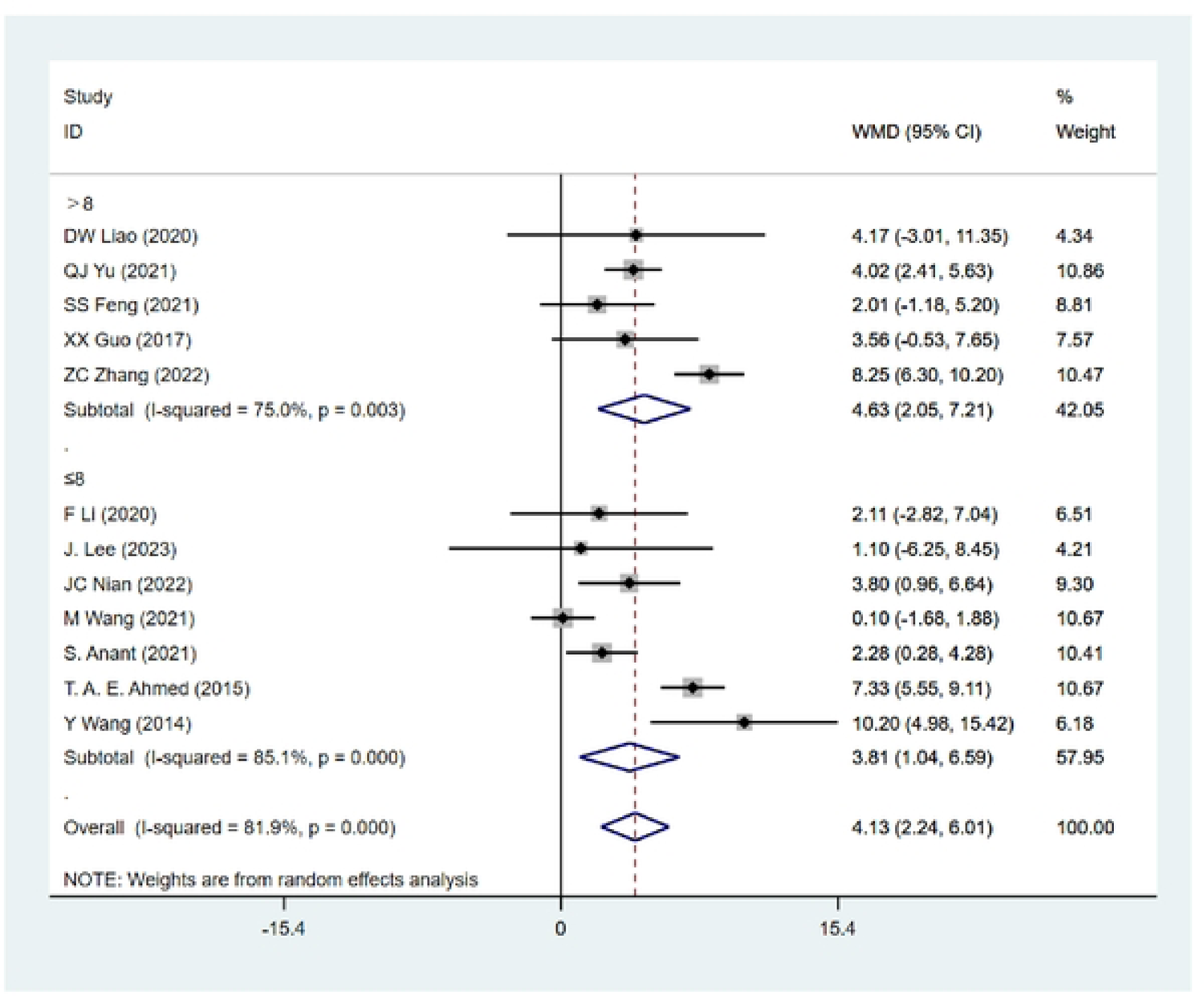
The effects of FT vs. TRT on muscle endurance (1 min SU) - Total weeks

**Figure 40:**
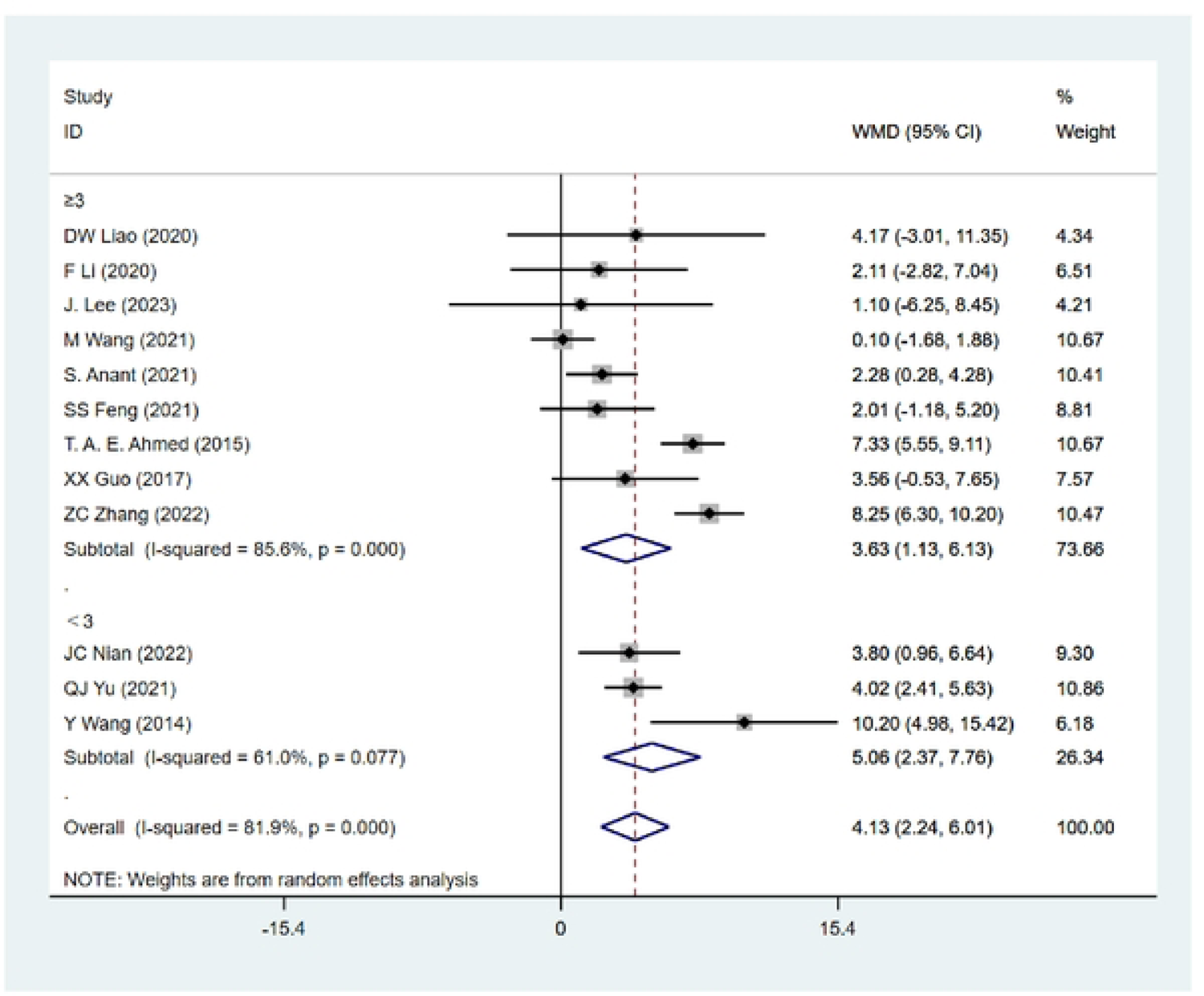
The effects of FT vs. TRT on muscle endurance (1 min SU) - Frequency.

**Figure 41:**
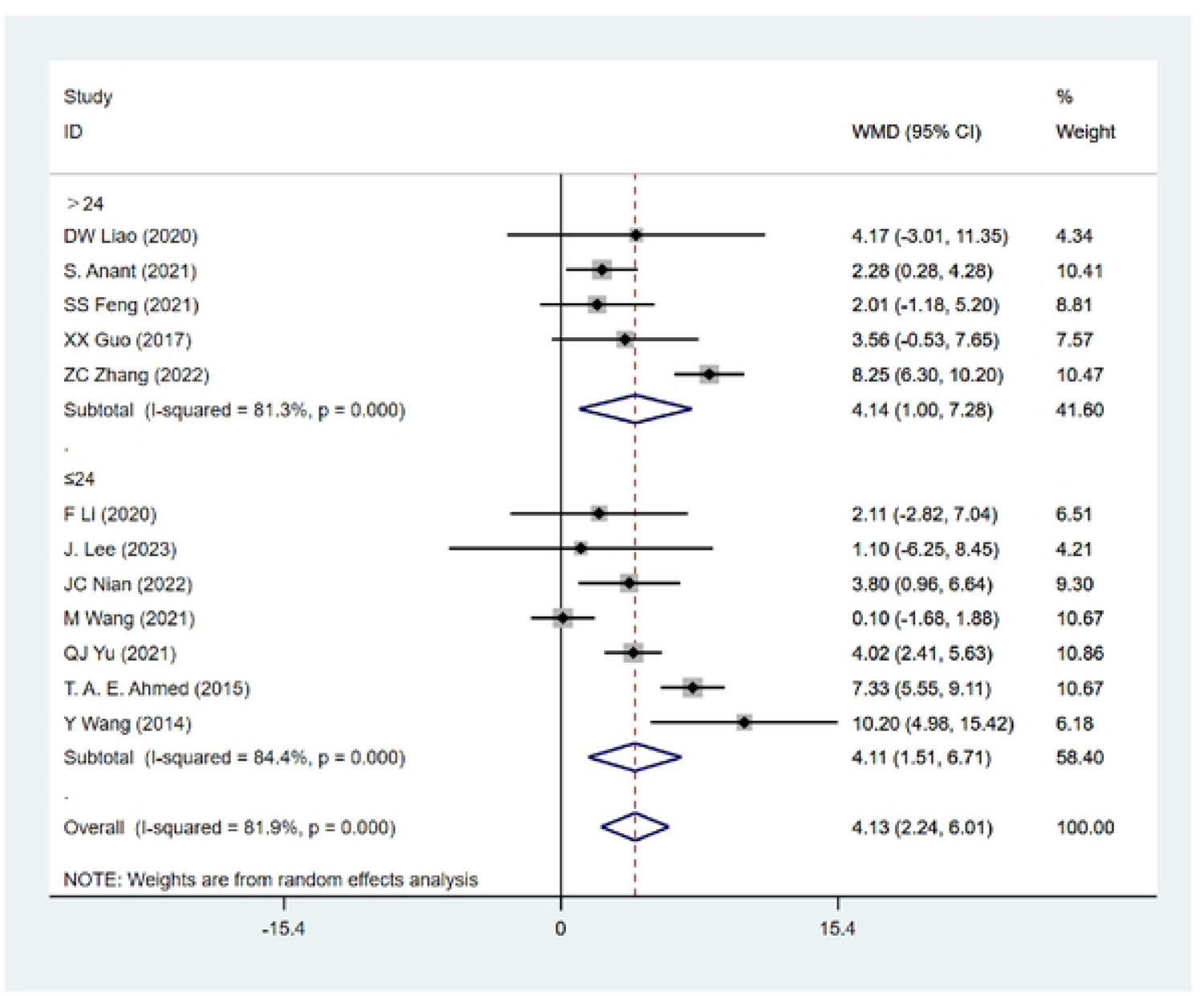
The effects of FT vs. TRT on muscle endurance (1 min SU) - Total sessions.

**Figure 42:**
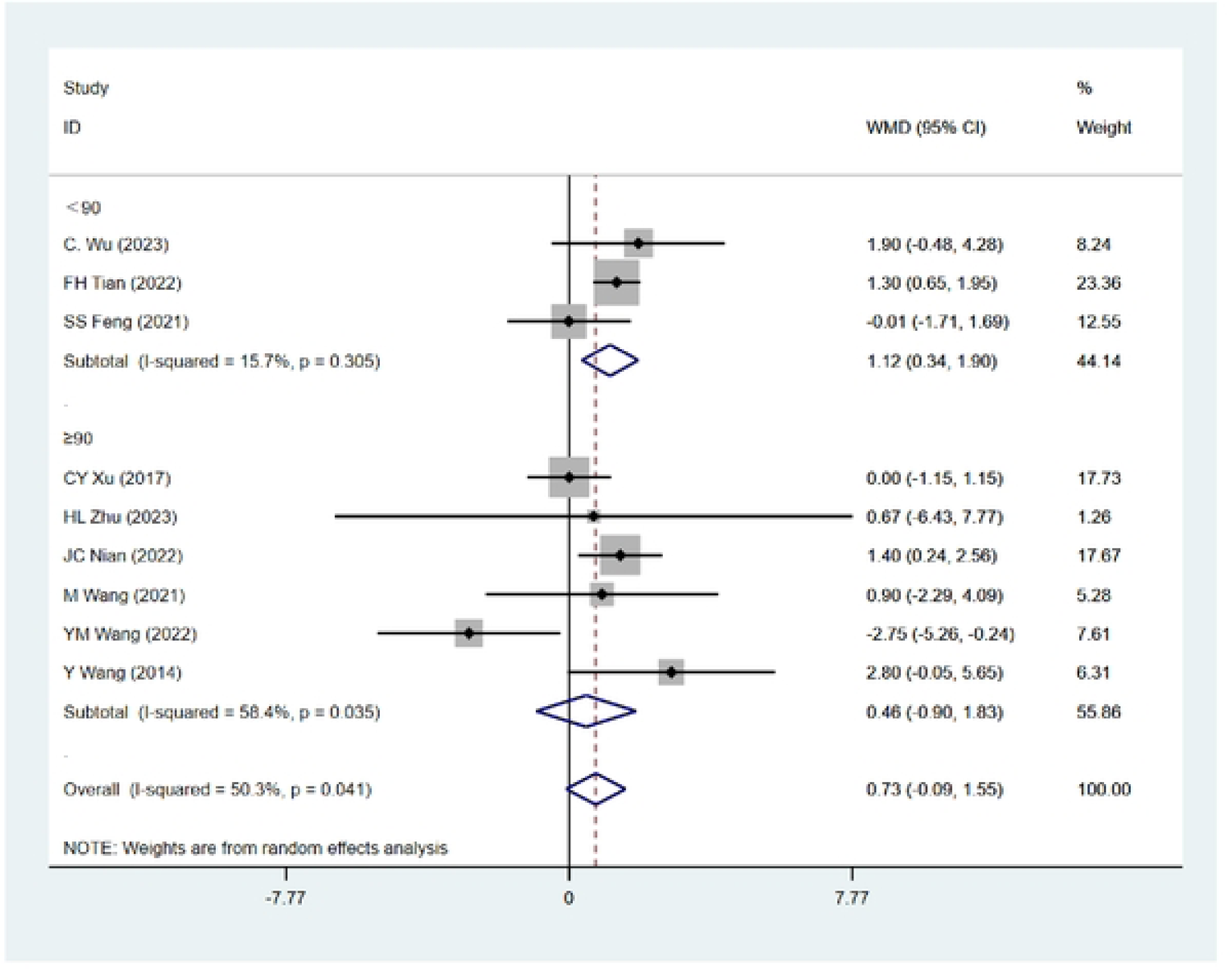
The effects of FT vs. TRT on muscle endurance (PLU) - Each session duration.

**Figure 43:**
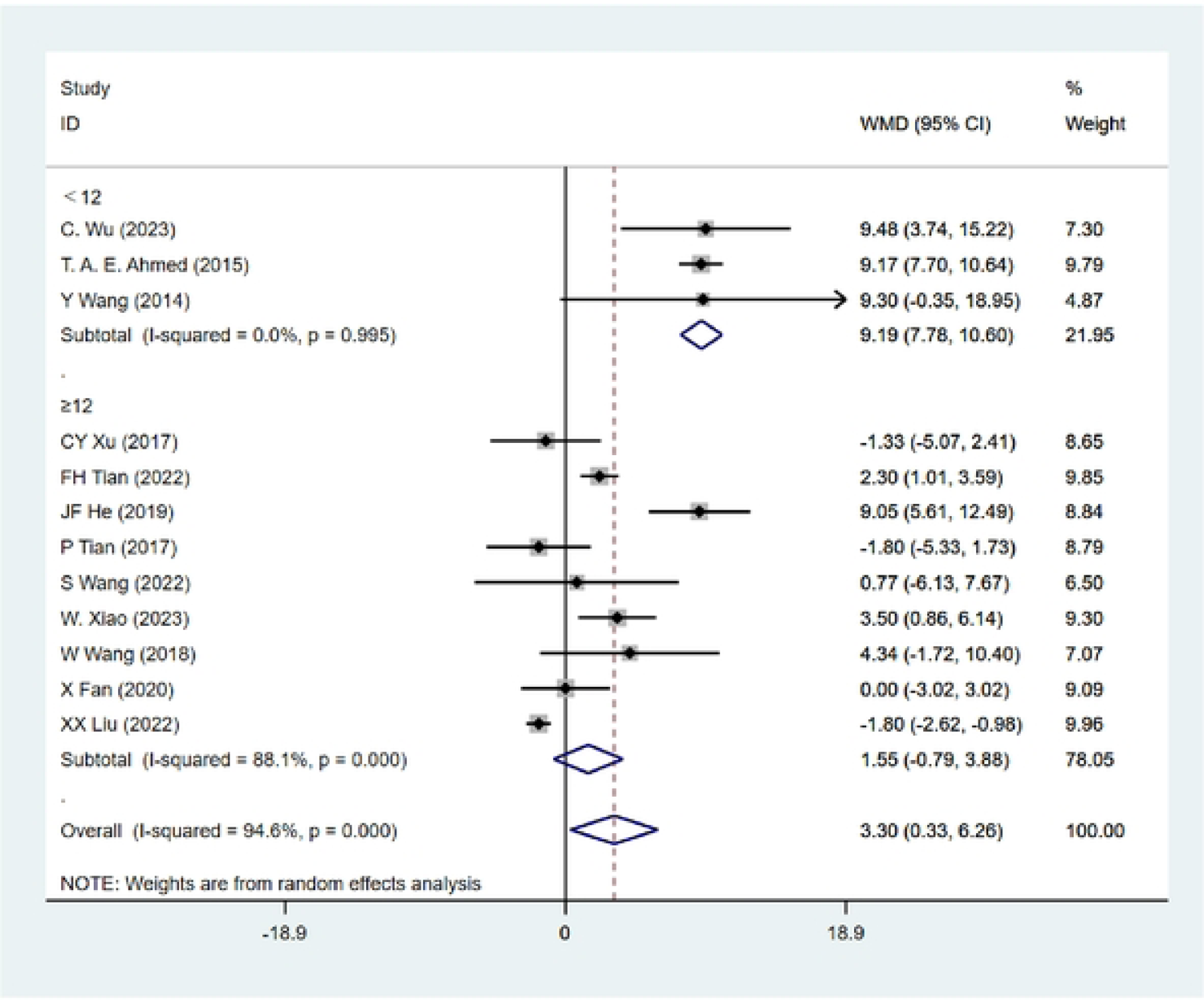
The effects of FT vs. TRT on muscle endurance (PSU) - Total weeks.

## References

1. Ahmed TAE. Improving musculoskeletal fitness and the performance enhancement of basketball skills through neuromuscular training program. Journal of Human Sport and Exercise 10: 795–804, 2015.

2. Alexander NB, Galecki AT, Grenier ML, et al. Task-specific resistance training to improve the ability of activities of daily living–impaired older adults to rise from a bed and from a chair. Journal of the American Geriatrics Society 49: 1418–1427, 2001.

3. Almuzaini KS, Fleck SJ. Modification of the standing long jump test enhances ability to predict anaerobic performance. The Journal of Strength & Conditioning Research 22: 1265–1272, 2008.

4. Alonso-Fernández D, Lima-Correa F, Gutierrez-Sánchez Á, Abadia-Garcia de Vicuna O. Effects of a high-intensity interval training protocol based on functional exercises on performance and body composition in handball female players. 2017.

5. Alonso-Fernández D, Lima-Correa F, Gutierrez-Sánchez Á, De Vicuña OA-G. Effects of a high-intensity interval training protocol based on functional exercises on performance and body composition in handball female players. Journal of Human Sport & Exercise 12: 1186–1198, 2017.

6. Anant S, Venugopal R. Effect of eight-week core muscles strength training on physical fitness and body composition variables in male players of team games. Revista Andaluza de Medicina del Deporte 14, 2021.

7. Anant SK, Venugopal R. Effect of eight-week core muscles strength training on physical fitness and body composition variables in male players of team games. Efecto del entrenamiento de fuerza muscular del Core de ocho semanas en las variables de aptitud física y composición corporal en jugadores masculinos de juegos de equipo 14: 17–23, 2021.

8. Andrade C. Mean difference, standardized mean difference (SMD), and their use in meta-analysis: as simple as it gets. The Journal of clinical psychiatry 81: 11349, 2020.

9. Aragón LF. Evaluation of four vertical jump tests: Methodology, reliability, validity, and accuracy. Measurement in physical education and exercise science 4: 215–228, 2000.

10. Balshaw TG, Massey GJ, Maden-Wilkinson TM, et al. Changes in agonist neural drive, hypertrophy and pre-training strength all contribute to the individual strength gains after resistance training. European journal of applied physiology 117: 631–640, 2017.

11. Barfield J, Channell B, Pugh C, Tuck M, Pendel D. Format of basic instruction program resistance training classes: Effect on fitness change in college students. Physical educator 69: 325, 2012.

12. Bashir M, Soh KG, Samsudin S, et al. Effects of functional training on sprinting, jumping, and functional movement in athletes: A systematic review. Frontiers in physiology 13:1045870, 2022.

13. Baumgartner TA. Modified pull-up test. Research Quarterly American Alliance for Health, Physical Education and Recreation 49: 80–84, 1978.

14. Baumgartner TA, Oh S, Chung H, Hales D. Objectivity, reliability, and validity for a revised push-up test protocol. Measurement in Physical Education and Exercise Science 6: 225–242, 2002.

15. Behm D, Sale D. Velocity specificity of resistance training. Sports medicine 15: 374–388, 1993.

16. Behm DG. Neuromuscular implications and applications of resistance training. Journal of Strength and Conditioning Research 9: 264–274, 1995.

17. Behm DG, Young JD, Whitten JH, et al. Effectiveness of traditional strength vs. power training on muscle strength, power and speed with youth: a systematic review and meta-analysis. Frontiers in physiology 8: 423, 2017.

18. Bianco A, Lupo C, Alesi M, et al. The sit up test to exhaustion as a test for muscular endurance evaluation. Springerplus 4: 1–8, 2015.

19. Bobbert MF. Drop jumping as a training method for jumping ability. Sports medicine 9: 7–22, 1990.

20. Bompa TO. Entrenamiento de la potencia aplicado a los deportes: la pliometría para el desarrollo de la máxima potencia. Inde, 2004.

21. Bonato M, Benis R, La Torre A. Neuromuscular training reduces lower limb injuries in elite female basketball players. A cluster randomized controlled trial. Scandinavian journal of medicine & science in sports 28: 1451–1460, 2018.

22. Borenstein M, Hedges LV, Higgins JP, Rothstein HR. A basic introduction to fixed-effect and random-effects models for meta-analysis. Research synthesis methods 1: 97–111, 2010.

23. Bouteraa I, Negra Y, Shephard RJ, Chelly MS. Effects of Combined Balance and Plyometric Training on Athletic Performance in Female Basketball Players. J Strength Cond Res 34: 1967–1973, 2020.

24. Bouteraa I, Negra Y, Shephard RJ, Chelly MS. Effects of combined balance and plyometric training on athletic performance in female basketball players. The Journal of Strength & Conditioning Research 34: 1967–1973, 2020.

25. Boyle M. New functional training for sports. Human Kinetics, 2016.

26. Buckner SL, Mouser JG, Jessee MB, et al. What does individual strength say about resistance training status? Muscle & nerve 55: 455–457, 2017.

27. Cabrejas C, Solana-Tramunt M, Morales J, et al. The Effects of an Eight-Week Integrated Functional Core and Plyometric Training Program on Young Rhythmic Gymnasts’ power Strength. International journal of environmental research and public health 20: 1041, 2023.

28. Castro-Piñero J, Ortega FB, Artero EG, et al. Assessing muscular strength in youth: usefulness of standing long jump as a general index of muscular fitness. The Journal of Strength & Conditioning Research 24: 1810–1817, 2010.

29. Claudino JG, Gabbett TJ, Bourgeois F, et al. CrossFit overview: systematic review and meta-analysis. Sports medicine-open 4: 1–14, 2018.

30. Clemson L, Singh MAF, Bundy A, et al. Integration of balance and strength training into daily life activity to reduce rate of falls in older people (the LiFE study): randomised parallel trial. Bmj 345, 2012.

31. Coyne J, Tran TT, Secomb JL, et al. Reliability of pull up & dip maximal strength tests. J Aust Strength Cond 23: 21–27, 2015.

32. Cress ME, Conley KE, Balding SL, Hansen-Smith F, Konczak J. Functional training: muscle structure, function, and performance in older women. Journal of Orthopaedic & Sports Physical Therapy 24: 4–10, 1996.

33. Cronin JB, Hansen KT. Strength and power predictors of sports speed. The Journal of Strength & Conditioning Research 19: 349–357, 2005.

34. Cureton TK. Relationship of physical fitness to athletic performance and sports. Journal of the American Medical Association 162: 1139–1149, 1956.

35. De Lacey J, Brughelli ME, McGuigan MR, Hansen KT. Strength, speed and power characteristics of elite rugby league players. The Journal of Strength & Conditioning Research 28: 2372–2375, 2014.

36. de Sousa AF, dos Santos GB, dos Reis T, et al. Differences in Physical Fitness between Recreational CrossFit® and Resistance Trained Individuals. Journal of Exercise Physiology Online 19, 2016.

37. De Villarreal ES-S, Requena B, Newton RU. Does plyometric training improve strength performance? A meta-analysis. Journal of science and medicine in sport 13: 513–522, 2010.

38. De Vreede PL, Samson MM, Van Meeteren NL, Duursma SA, Verhaar HJ. Functional-task exercise versus resistance strength exercise to improve daily function in older women: a randomized, controlled trial. Journal of the American Geriatrics Society 53: 2–10, 2005.

39. Diener MH, Golding LA, Diener D. Validity and reliability of a one-minute half sit-up test of abdominal strength and endurance. Research in Sports Medicine: An International Journal 6: 105–119, 1995.

40. Ebenbichler Gr, Oddsson Li, Kollmitzer J, Erim Z. Sensory-motor control of the lower back: implications for rehabilitation. Medicine & Science in Sports & Exercise 33: 1889–1898, 2001.

41. Elsayed Ahmed TA. Improving musculoskeletal fitness and the performance enhancement of basketball skills through neuromuscular training program. Journal of Human Sport & Exercise 10: 795–804, 2015.

42. Erol S, Yapici A, Gumusdag H. The effect of functional training applied to soccer players on anaerobic performance. Journal of ROL Sport Sciences 4: 128–141, 2023.

43. Fernandez-Fernandez J, García-Tormo V, Santos-Rosa FJ, et al. The effect of a neuromuscular vs. dynamic warm-up on physical performance in young tennis players. The Journal of Strength & Conditioning Research 34: 2776–2784, 2020.

44. Foss KDB, Thomas S, Khoury JC, Myer GD, Hewett TE. A school-based neuromuscular training program and sport-related injury incidence: a prospective randomized controlled clinical trial. Journal of athletic training 53: 20–28, 2018.

45. Gaedtke A, Morat T. TRX suspension training: A new functional training approach for older adults–development, training control and feasibility. International journal of exercise science 8(3):224, 2015.

46. Gambetta V, Gray G. Following a functional path. Training & Conditioning 5: 25–30, 1995.

47. Ganley KJ, Paterno MV, Miles C, et al. Health-related fitness in children and adolescents. Pediatric Physical Therapy 23: 208–220, 2011.

48. Gee TI, Morrow RA, Stone MR, Bishop DC. A neuromuscular training program enhances dynamic neuromuscular control and physical performance in court-sport athletes. Translational Sports Medicine 3: 9–15, 2020.

49. Giné-Garriga M, Guerra M, Pagès E, et al. The effect of functional circuit training on physical frailty in frail older adults: a randomized controlled trial. Journal of aging and physical activity 18: 401–424, 2010.

50. Guilherme JPLF, Tritto ACC, North KN, Lancha Junior AH, Artioli GG. Genetics and sport performance: current challenges and directions to the future. Revista Brasileira de Educação Física e Esporte 28: 177–193, 2014.

51. Haff GG, Triplett NT. Essentials of strength training and conditioning 4th edition. Human kinetics, 2015.

52. Hashim A, Ariffin A, Hashim AT, Yusof AB. Reliability and validity of the 90° push-ups test protocol. International Journal of Scientific Research and Management 6: 10.18535, 2018.

52. Helbostad JL, Sletvold O, Moe-Nilssen R. Effects of home exercises and group training on functional abilities in home-dwelling older persons with mobility and balance problems. A randomized study. Aging clinical and experimental research 16: 113–121, 2004.

53. Hermassi S, Wollny R, Schwesig R, Shephard RJ, Chelly MS. Effects of in-season circuit training on physical abilities in male handball players. The Journal of Strength & Conditioning Research 33: 944–957, 2019.

54. Higashihara A, Ono T, Kubota J, Okuwaki T, Fukubayashi T. Functional differences in the activity of the hamstring muscles with increasing running speed. Journal of sports sciences 28: 1085–1092, 2010.

55. Higgins JP, Thompson SG, Deeks JJ, Altman DG. Measuring inconsistency in meta-analyses. Bmj 327: 557–560, 2003.

56. Italo S, Giacomo C, Rosario Do. The effects of 8 weeks of integrative neuromuscular pitch training on strength values and sprint performance in young élite soccer players. Journal of Physical Education & Sport 23, 2023.

57. Italo S, Giacomo C, Rosario DO. The effects of 8 weeks of integrative neuromuscular pitch training on strength values and sprint performance in young élite soccer players. Journal of Physical Education & Sport 23: 909–917, 2023.

58. Keiner M, Kadlubowski B, Sander A, Hartmann H, Wirth K. Effects of 10 months of speed, functional, and traditional strength training on strength, linear sprint, change of direction, and jump performance in trained adolescent soccer players. Journal of Strength and Conditioning Research 36: 2236–2246, 2022.

59. Khayambashi K, Ghoddosi N, Straub RK, Powers CM. Hip muscle strength predicts noncontact anterior cruciate ligament injury in male and female athletes: a prospective study. The American journal of sports medicine 44: 355–361, 2016.

60. Khazaei L, Parnow A, Amani-Shalamzari S. Comparing the effects of traditional resistance training and functional training on the bio-motor capacities of female elite taekwondo athletes. *BMC Sports Science*, Medicine and Rehabilitation 15: 139, 2023.

61. Khazaei L, Parnow A, Amani-Shalamzari S. Comparing the effects of traditional resistance training and functional training on the bio-motor capacities of female elite taekwondo athletes. BMC Sports Sci Med Rehabil 15: 139, 2023.

62. Kibler WB, Press J, Sciascia A. The role of core stability in athletic function. Sports medicine 36: 189–198, 2006.

63. King MB, Judge JO, Whipple R, et al. Reliability and responsiveness of two physical performance measures examined in the context of a functional training intervention. Physical therapy 80(1):8–16, 2000.

64. Komi PV, Gollhofer A. Stretch reflexes can have an important role in force enhancement during SSC exercise. Journal of applied biomechanics 13: 451–460, 1997.

65. Kondrič M, Zagatto AM, Sekulić D. The physiological demands of table tennis: a review. Journal of sports science & medicine 12: 362, 2013.

66. La Scala Teixeira CV, Evangelista AL, Novaes JS, Da Silva Grigoletto ME, Behm DG. “You’re only as strong as your weakest link”: a current opinion about the concepts and characteristics of functional training. Frontiers in physiology 8: 643, 2017.

67. Lee J, Yang G-S, Lee M. The Effects of Two Types of Training on the Physical Ability of University Baseball Players. Annals of Applied Sport Science 11, 2023.

68. Lee J, Yang G-S, Lee M. The Effects of Two Types of Training on the Physical Ability of University Baseball Players. Annals of Applied Sport Science: 0–0, 2023.

69. Lehance C, Binet J, Bury T, Croisier J-L. Muscular strength, functional performances and injury risk in professional and junior elite soccer players. Scandinavian journal of medicine & science in sports 19: 243–251, 2009.

70. Liu C-j, Shiroy DM, Jones LY, Clark DO. Systematic review of functional training on muscle strength, physical functioning, and activities of daily living in older adults. European review of aging and physical activity 11: 95–106, 2014.

71. MacArthur DG, North KN. Genes and human elite athletic performance. East African Running: 241–257, 2007.

72. Manini T, Marko M, VanArnam T, et al. Efficacy of resistance and task-specific exercise in older adults who modify tasks of everyday life. The Journals of Gerontology Series A: Biological Sciences and Medical Sciences 62: 616–623, 2007.

73. Markovic G, Dizdar D, Jukic I, Cardinale M. Reliability and factorial validity of squat and countermovement jump tests. The Journal of Strength & Conditioning Research 18: 551–555, 2004.

74. Markovic G, Mikulic P. Neuro-musculoskeletal and performance adaptations to lower-extremity plyometric training. Sports medicine 40: 859–895, 2010.

75. Masley JW, Hairabedian A, Donaldson DN. Weight training in relation to strength, speed, and co-ordination. *Research Quarterly American Association for Health*, Physical Education and Recreation 24: 308–315, 1953.

76. Michell TB, Ross SE, Blackburn JT, Hirth CJ, Guskiewicz KM. Functional balance training, with or without exercise sandals, for subjects with stable or unstable ankles. Journal of Athletic Training 41: 393, 2006.

77. Moeskops S, Oliver JL, Read PJ, et al. Effects of a 10-Month Neuromuscular Training Program on Strength, Power, Speed, and Vault Performance in Young Female Gymnasts. Med Sci Sports Exerc 54: 861–871, 2022.

78. Moher D, Shamseer L, Clarke M, et al. Preferred reporting items for systematic review and meta-analysis protocols (PRISMA-P) 2015 statement. Systematic reviews 4: 1–9, 2015.

79. Mohr M, Krustrup P, Bangsbo J. Match performance of high-standard soccer players with special reference to development of fatigue. Journal of sports sciences 21: 519–528, 2003.

80. Montgomery PG, Pyne DB, Minahan CL. The physical and physiological demands of basketball training and competition. International journal of sports physiology and performance 5: 75–86, 2010.

81. Nadler SF, Malanga GA, DePrince M, Stitik TP, Feinberg JH. The relationship between lower extremity injury, low back pain, and hip muscle strength in male and female collegiate athletes. Clinical Journal of Sport Medicine 10: 89–97, 2000.

82. Nawed A, Khan IA, Jalwan J, Nuhmani S, Muaidi QI. Efficacy of FIFA 11 + training program on functional performance in amateur male soccer players. Journal of Back & Musculoskeletal Rehabilitation 31: 867–870, 2018.

83. Nawed A, Khan IA, Jalwan J, Nuhmani S, Muaidi QI. Efficacy of FIFA 11+ training program on functional performance in amateur male soccer players. Journal of back and musculoskeletal rehabilitation 31: 867–870, 2018.

84. Okada T, Huxel KC, Nesser TW. Relationship between core stability, functional movement, and performance. The Journal of Strength & Conditioning Research 25: 252–261, 2011.

85. Orwin RG. A fail-safe N for effect size in meta-analysis. Journal of educational statistics 8: 157–159, 1983.

86. Peters JL, Sutton AJ, Jones DR, Abrams KR, Rushton L. Comparison of two methods to detect publication bias in meta-analysis. Jama 295: 676–680, 2006.

87. Phomsoupha M, Laffaye G. The science of badminton: game characteristics, anthropometry, physiology, visual fitness and biomechanics. Sports medicine 45: 473–495, 2015.

88. Pienaar C, Coetzee B. Changes in selected physical, motor performance and anthropometric components of university-level rugby players after one microcycle of a combined rugby conditioning and plyometric training program. J Strength Cond Res 27: 398–415, 2013.

89. Pienaar C, Coetzee B. Changes in selected physical, motor performance and anthropometric components of university-level rugby players after one microcycle of a combined rugby conditioning and plyometric training program. The Journal of Strength & Conditioning Research 27: 398–415, 2013.

90. Puthucheary Z, Skipworth JR, Rawal J, et al. Genetic influences in sport and physical performance. Sports medicine 41: 845–859, 2011.

91. Radcliffe JC. Functional training for athletes at all levels: workouts for agility, speed and power. Simon and Schuster, 2007.

92. Ramirez-Campillo R, Álvarez C, García-Hermoso A, et al. Methodological characteristics and future directions for plyometric jump training research: a scoping review. Sports Medicine 48: 1059–1081, 2018.

93. Ratamess N. ACSM’s foundations of strength training and conditioning. Lippincott Williams & Wilkins, 2021.

94. Reis I, Rebelo A, Krustrup P, Brito J. Performance enhancement effects of Federation Internationale de Football Association’s “The 11+” injury prevention training program in youth futsal players. Clinical journal of sport medicine 23: 318–320, 2013.

95. Riva D, Bianchi R, Rocca F, Mamo C. Proprioceptive training and injury prevention in a professional men’s basketball team: a six-year prospective study. Journal of strength and conditioning research 30: 461, 2016.

96. Root H, Trojian T, Martinez J, Kraemer W, DiStefano LJ. Landing technique and performance in youth athletes after a single injury-prevention program session. Journal of athletic training 50: 1149–1157, 2015.

97. Roso-Moliner A, Mainer-Pardos E, Carton-Llorente A, et al. Effects of a neuromuscular training program on physical performance and asymmetries in female soccer. Frontiers in Physiology 14, 2023.

98. Roso-Moliner A, Mainer-Pardos E, Cartón-Llorente A, et al. Effects of a neuromuscular training program on physical performance and asymmetries in female soccer. Frontiers in Physiology 14: 1171636, 2023.

99. Rothwell PM. Subgroup analysis in randomised controlled trials: importance, indications, and interpretation. The Lancet 365: 176–186, 2005.

100. Sakata J, Nakamura E, Suzuki T, et al. Efficacy of a prevention program for medial elbow injuries in youth baseball players. The American journal of sports medicine 46: 460–469, 2018.

101. Sale D, MacDougall D. Specificity in strength training: a review for the coach and athlete. Canadian journal of applied sport sciences Journal canadien des sciences appliquées au sport 6: 87–92, 1981.

102. Sattler T, Sekulic D, Hadzic V, Uljevic O, Dervisevic E. Vertical jumping tests in volleyball: reliability, validity, and playing-position specifics. The Journal of Strength & Conditioning Research 26: 1532–1538, 2012.

103. Sawczuk M, Maciejewska A, Cięszczyk P, Eider J. The role of genetic research in sport. Science & sports 26: 251–258, 2011.

104. Sawilowsky SS. New effect size rules of thumb. Journal of modern applied statistical methods 8: 26, 2009.

105. Sartor CD, Hasue RH, Cacciari LP, et al. Effects of strengthening, stretching and functional training on foot function in patients with diabetic neuropathy: results of a randomized controlled trial. BMC musculoskeletal disorders 15:1–13, 2014.

106. Schwarzer G. meta: An R package for meta-analysis. R news 7: 40–45, 2007.

107. Siff MC. Biomechanical foundations of strength and power training. Biomechanics in sport: performance enhancement and injury prevention: 103–139, 2000.

108. Silva-Grigoletto MED, Resende-Neto AGd, Teixeira CVLS. Functional training: a conceptual update. Revista Brasileira de Cineantropometria & Desempenho Humano 22, 2020.

109. Skelton DA, McLaughlin AW. Training functional ability in old age. Physiotherapy 82: 159–167, 1996.

110. Song H-S, Woo S-S, So W-Y, et al. Effects of 16-week functional movement screen training program on strength and flexibility of elite high school baseball players. Journal of exercise rehabilitation 10: 124, 2014.

111. Spennewyn KC. Strength outcomes in fixed versus free-form resistance equipment. The Journal of Strength & Conditioning Research 22: 75–81, 2008.

112. Sterne JA. Meta-analysis in Stata: an updated collection from the Stata Journal. StataCorp LP, 2009.

112. Sterne JA, Harbord RM. Funnel plots in meta-analysis. The stata journal 4: 127–141, 2004.

113. Sterne JA, Savović J, Page MJ, et al. RoB 2: a revised tool for assessing risk of bias in randomised trials. bmj 366, 2019.

114. Stockbrugger BA, Haennel RG. Validity and reliability of a medicine ball power power test. The Journal of strength & conditioning research 15: 431–438, 2001.

115. Stone MH. Position statement: power exercise and training. Strength & Conditioning Journal 15: 7–15, 1993.

116. Suchomel TJ, Nimphius S, Bellon CR, Stone MH. The importance of muscular strength: training considerations. Sports medicine 48: 765–785, 2018.

117. Suchomel TJ, Nimphius S, Stone MH. The importance of muscular strength in athletic performance. Sports medicine 46: 1419–1449, 2016.

118. Taube W, Leukel C, Gollhofer A. How neurons make us jump: the neural control of stretch-shortening cycle movements. Exercise and sport sciences reviews 40: 106–115, 2012.

119. Thompson BJ. Effect of surface stability on core muscle activity during dynamic resistance exercises. Utah State University, 2009.

120. Thompson SG, Higgins JP. How should meta-regression analyses be undertaken and interpreted? Statistics in medicine 21: 1559–1573, 2002.

121. Thompson CJ, Cobb KM, Blackwell J. Functional training improves club head speed and functional fitness in older golfers. The journal of strength & conditioning research 21(1):131–137, 2007.

122. Tomljanović M, Spasić M, Gabrilo G, Uljević O, Foretić N. Effects of five weeks of functional vs. traditional resistance training on anthropometric and motor performance variables. Kinesiology 43: 145–154, 2011.

123. Trakis JE, McHugh MP, Caracciolo PA, et al. Muscle strength and range of motion in adolescent pitchers with throwing-related pain: implications for injury prevention. The American journal of sports medicine 36: 2173–2178, 2008.

124. Turna B, Alp M. The Effects of Functional Training on Some Biomotor Abilities and Physiological Characteristics in Elite Soccer Players. Journal of Education and Learning 9: 164–171, 2020.

125. Usgu S, Yakut Y, Kudaş S. Effects of functional training on performance in professional basketball players. Spor Hekimliği Dergisi 55: 321–331, 2020.

126. Vandorpe B, Vandendriessche J, Vaeyens R, et al. Factors discriminating gymnasts by competitive level. International journal of sports medicine: 591–597, 2011.

127. Verdijk LB, Van Loon L, Meijer K, Savelberg HH. One-repetition maximum strength test represents a valid means to assess leg strength in vivo in humans. Journal of sports sciences 27: 59–68, 2009.

128. Verstegen M, Williams P. Core performance essentials: the revolutionary nutrition and exercise plan adapted for everyday use. Rodale Books, 2006.

129. Vitale JA, La Torre A, Banfi G, Bonato M. Effects of an 8-Week Body-Weight Neuromuscular Training on Dynamic Balance and Vertical Jump Performances in Elite Junior Skiing Athletes: A Randomized Controlled Trial. J Strength Cond Res 32: 911–920, 2018.

130. Vitale JA, La Torre A, Banfi G, Bonato M. Effects of an 8-week body-weight neuromuscular training on dynamic balance and vertical jump performances in elite junior skiing athletes: A randomized controlled trial. The Journal of Strength & Conditioning Research 32: 911–920, 2018.

131. Walsh M, Arampatzis A, Schade F, Brüggemann G-P. The effect of drop jump starting height and contact time on power, work performed, and moment of force. The Journal of Strength & Conditioning Research 18: 561–566, 2004.

132. Willardson JM. Core stability training: applications to sports conditioning programs. The Journal of Strength & Conditioning Research 21: 979–985, 2007.

133. Wu C, Cheong M, Wang Y, et al. Impact of Functional Training on Functional Movement and Athletic Performance in College Dragon Boat Athletes. International Journal of Environmental Research and Public Health 20: 3897, 2023.

134. Xiao W, Bai X, Geok SK, Yu D, Zhang Y. Effects of a 12-Week Functional Training Program on the Strength and Power of Chinese Adolescent Tennis Players. Children 10: 635, 2023.

135. Yildiz S, Pinar S, Gelen E. Effects of 8-week functional vs. traditional training on athletic performance and functional movement on prepubertal tennis players. The Journal of Strength & Conditioning Research 33: 651–661, 2019.

136. Zırhlı O, Demirci N. The Influence of functional training on biomotor skills in girl tennis players aged 10–12. Baltic Journal of Health and Physical Activity 12: 4, 2020.

137. Duan Yuanpeng. A Practical Study of Functional Training in Tianjin Primary School Football Team. Master’s thesis, Tianjin Sports Institute, Tianjin, 2020.

138. Fan Xiao. Research on the effects of functional training and traditional physical training on the athletic quality index of Fujian men’s basketball reserve team. Master’s thesis, Chengdu Institute of Physical Education, Chengdu 2020.

139. Feng Shasha. A study on the effect of functional training on the physical characteristics of U15 male soccer players. Master’s thesis, Liaoning Normal University, Dalian, 2021.

140. Guo Haixia. Research on the Effect of Physical Function Training on Physical Quality of Badminton Players in General Colleges and Universities. Master’s Thesis, Northwest Normal University, Lanzhou, 2020.

141. Guo Xuanxuan. An experimental study on functional training of child and adolescent badminton players. Master’s thesis, Beijing Sport University, Beijing, 2017.

142. Han Lei. A study on the effect of functional training based on FMS test on children’s basketball offensive skills. Master’s thesis, Xinjiang Normal University, Urumqi 2022.

143. He Jianfei. Experimental Study on the Effect of Functional Training on Shooting Percentage of 14--16 Years Old Male Basketball Players--Taking Yulin Blue Dream Basketball Training Camp as an Example. Master’s thesis, Xi’an Institute of Physical Education,Xi’an, 2019.

144. Hu Sheng. Research on the Application of Functional Training in Physical Training of College Men’s Volleyball Team. Master Thesis, Hunan Normal University, Changsha 2016.

145. Janot JM, Weiss T, Kreitinger J, et al. Effect of Functional Resistance Training on Muscular Fitness Outcomes in Young Adults. Medicine and Science in Sports and Exercise 42(5):296–296, 2010.

146. Jia Haiyuan. Effects of functional strength training on the bouncing quality of high school male basketball players. Master’s thesis, Shanxi Normal University, Taiyuan, 2018.

147. Jiang Yang, Huang Baohong, Zhou Zhixiong. A study on the effect of physical function training on FMS and motor quality of female volleyball players. Journal of Jilin Institute of Physical Education 30: 60–64, 2014.

148. JU Yucheng. An empirical study on functional training to improve the physical quality of badminton specialization. Master’s thesis, Shudu Institute of Physical Education, Beijing, 2023.

149. Li Buchun. An Experimental Study on the Effect of Functional Training on Badminton Technique. Master’s thesis, Xinjiang Normal University, Urumqi, 2022.

150. Li F. An experimental study on functional training to improve landing stability of age-appropriate male gymnasts. Master’s thesis, Shandong Institute of Physical Education, Jinan, 2020.

151. Li Fengyan. Research on the Effect of Physical Function Training on Lower Limb Stability of Secondary School Soccer Players. Master’s thesis, Shandong Institute of Physical Education, Jinan, 2019.

152. Liao Dawei. An Experimental Study of Functional Training for Baseball Players. Master’s thesis, Shanghai International Studies University, Shanghai, 2020.

153. Ling Xin. Research on the influence of functional training on the effect of batting quality of tennis specialization students. Master’s thesis, Tianjin Sports Institute, Tianjin, 2023.

154. Liu Xixiao. The Role of Functional Fitness Training in Enhancing Physical Fitness of Youth Basketball Players. Journal of Anyang Institute of Technology 21: 118–122, 2022.

155. Liu Yiwei. Research on the effect of functional training on the stability of college soccer players’ special physical quality and shooting technique. Master’s thesis, Jilin University, Changchun, 2021.

156. Luo Li. Research on the Effect of Functional Training on Physical Fitness of Men’s Track and Field Athletes in Hunan Institute of Technology. Master’s thesis, Hunan Institute of Technology, Yueyang, 2022.

157. Luo Zhenyu. Research on the Effect of Functional Training on the Strength of Shoulder Complex of College High-level Tennis Players. Master’s Thesis, Southwest University of Finance and Economics, Chengdu, 2020.

158. Nian, Jianchun. An empirical study of functional training on improving strength quality of college basketball specialization students. Master’s thesis, Harbin Institute of Physical Education, Harbin, 2022.

159. Pan Q. A Study on Functional Training to Improve Sports Performance and Balance Ability of Male Freestyle Wrestlers. Master’s Thesis, Guangzhou Sports Institute, Guangzhou, 2020.

160. Qi Yifei. Experimental Study on the Effect of Functional Training on Lower Limb power Strength of Youth Taekwondo Athletes. Master’s thesis, Shandong Institute of Physical Education, Jinan, 2022.

161. Song Chundi. Applied Research on Functional Physical Training to Improve the Specialized Sports Quality of Classical Wrestlers. Master’s thesis, Shandong Institute of Physical Education, Jinan, 2021.

162. Tian Fanghe. An experimental study of functional upper limb strength training on the quality of technical movement completion of students specializing in gymnastics. Master’s thesis, Tianjin Sports Institute, Tianjin, 2022.

163. Tian Peng. Application of Functional Fitness Training in Physical Training of Youth Basketball Players. Master’s thesis,Shandong Institute of Physical Education, Jinan, 2017.

164. Wang Mu. Research on the Effect of Functional Training on the Sports Performance of Young Swimmers. Master’s Thesis, Southeast University, Nanjing, 2021.

165. Wang Shuai, Guo Liya. Research on the innovative application of body functional training in college tennis training. Sports Goods and Technology: 204–206, 2022.

166. Wang W. An experimental study on the effect of functional training of boxers. Master’s Thesis, Xi’an Institute of Physical Education, Xi’an, 2018.

167. Wang Yan. Effects of Functional Training on Swimmers’ Performance. Master’s thesis, Beijing Sport University, Beijing 2014.

168. Wang Yiming. Experimental Research on Functional Training in Specialized Physical Quality Training of College Men’s High Level Soccer Team. Master’s thesis, Wuhan Institute of Physical Education, Wuhan, 2022.

169. Wang Z. Effects of functional training on the specialized qualities of junior table tennis players. Master’s thesis,Nanjing Normal University, Nanjing, 2020.

170. Wei ZF. Exercise biomechanics analysis and body movement function training program research on special movement patterns of adolescent male volleyball players. Doctoral dissertation, First Institute of Physical Education, Beijing, 2023.

171. Xiao Haobo. An experimental study on the effect of functional strength training on the specialization quality of 7-12 years old badminton amateur gymnasium athletes. Master’s Thesis, Shandong Institute of Physical Education, Jinan, 2022, China.

172. Xu Haokai. Research on the effect of functional strength training on jump shot hitting rate and bouncing quality of college basketball players. Master’s thesis, Jilin University, Changchun, 2023.

173. Xu Jianchong. An empirical study on the effect of functional strength training on the rapid change of direction movement ability of volleyball specialized students. Master’s thesis, Beijing Sport University, Beijing, 2018.

174. Xu Changyong. Application and Empirical Research of Functional Training in High-level Sparring Athletes. Master’s thesis, Beijing Sport University, Beijing, 2017.

175. Yang Yang. An experimental study on the effect of functional strength training on speed and strength qualities of badminton players. Master’s thesis, Beijing Sport University, Beijing, 2015.

176. Yu Qingji. Research on the effect of functional strength training method on the bouncing force of middle school basketball players. Master’s thesis, Hunan University of Science and Technology, Xiangtan, 2021.

177. Zhang Zhenchao. An experimental study on the effect of functional training on the serving effect of junior male tennis players. Master’s thesis, Guangzhou Sports Institute, Guangzhou, 2022.

178. Zhao Jinwei. Research on the effect of functional training on table tennis technique. Master Thesis, Zhengzhou University, Zhengzhou, 2020.

179. Zhou Xiao. Empirical Study on Functional Training Based on FMS in Physical Training of Players in Soccer Special Schools. Master’s thesis, Shandong Institute of Physical Education, Jinan, 2020.

180. Zhu Huali. Research on the Effect of Functional Training on the Basic Physical Fitness of Windsurfing Athletes in Shandong Province. Master’s Thesis, Shandong Institute of Physical Education, Jinan, 2023.

181. Zhu Y. An experimental study of functional training in soccer-specific training for physical education majors. Master’s thesis, Jiangxi Normal University, Nanchang, 2022.

